# Mosaic integration of spatial multi-omics with SpaMosaic

**DOI:** 10.1101/2024.10.02.616189

**Authors:** Xuhua Yan, Zhaoyu Fang, Kok Siong Ang, Lynn van Olst, Alex Edwards, Thomas Watson, Ruiqing Zheng, Di Zhang, Rong Fan, Min Li, David Gate, Jinmiao Chen

## Abstract

With the advent of spatial multi-omics, mosaic integration of diverse datasets with partially overlapping modalities enables construction of comprehensive multi-modal spatial atlases from heterogeneous sources. Here, we present SpaMosaic, a tool that employs contrastive learning and graph neural networks to build a modality-agnostic and batch-corrected latent space for spatial domain identification and missing modality imputation. We systematically benchmarked SpaMosaic against existing integration methods using simulated data and experimentally acquired datasets spanning RNA and protein abundance, chromatin accessibility, and histone modifications from brain, embryo, tonsil, and lymph node tissues. SpaMosaic consistently outperformed other methods in identifying coherent spatial domains by reducing noise and mitigating batch effects across diverse technologies and developmental stages. Computationally, SpaMosaic is highly scalable, capable of integrating over 100 sections and processing a single section with more than 800,000 spots. Beyond robust integration, the unified latent space generated by SpaMosaic enables accurate imputation of missing modalities. In a mosaic mouse brain dataset, the imputed histone modifications not only recapitulated expected transcriptome-epigenome correlations but also uncovered more region-specific regulatory links compared to the measured chromatin accessibility data, demonstrating the ability to infer relationships between modalities without co-profiling. In summary, SpaMosaic provides a versatile framework for unifying the rapidly accumulating heterogeneous spatial omics data into comprehensive biological atlases.

## Introduction

Spatial omics are powerful techniques for studying cellular heterogeneity in its spatial context within tissues^1^. Current developments in spatial multi-omics are expanding the scope to simultaneously acquire multiple omics from a single tissue section^2^. For example, spatial ATAC-RNA-seq captures chromatin accessibility and RNA expression^2^, spatial CUT&Tag-RNA-seq captures histone modifications and RNA expression^2^, and SPOTS measures RNA and protein abundance^3^. Recent developments such as spatial ATAC-RNA-Protein-seq and spatial CUT&Tag-RNA-Protein-seq^4^ have enabled tri-omics data acquisition, while Spatial-Mux-seq^5^ can simultaneously profile up to five different modalities. By capturing different omics profiles of the same cell, these complementary views can be used to construct more comprehensive pictures and better dissect phenotypic niches of samples^6^. However, spatial omics experiments are technically challenging, involving trade-offs between resolution, sensitivity and throughput, and acquiring multiple omics concurrently from the same section exacerbates these challenges. At present, the most commonly available data is from single-omics modality experiments on different and sometimes adjacent tissue sections. Thus, spatial multi-omics with shared but also complementary omes can be computationally integrated to obtain a more coherent, complete, and higher-dimensional picture of tissue biology. Such integration, termed mosaic integration, can help consensus domain identification, enhance signal to noise ratio, impute missing assays in different sections, and dissect cross-linking between omics, which will be crucial for constructing a multi-omics spatial tissue atlas.

In essence, mosaic integration seeks to integrate multiple batches with different multi-modal compositions^7^. However, unlike diagonal integration where there are no shared batches or modalities, mosaic integration leverages a mosaic of shared modalities between batches to serve as bridges to accomplish data integration. Therefore, mosaic integration needs to accomplish both vertical integration of harmonizing modality heterogeneity for the same cells and horizontal integration of reducing technical variations in the same modality across batches^8^. In the context of mosaic integrating spatial omics data, the spatial context is an additional layer of information that should be utilized to accomplish a more coherent and accurate integration.

There are currently several single-cell mosaic integration methods available, such as Cobolt^9^, scMoMaT^8^, StabMap^10^ and MIDAS^11^. As these methods are tailored for single-cell mosaic integration, they do not consider the spatial context and thus may struggle with mosaic spatial multi-omics data. Spatially aware methods include neighborhood-aware clustering approaches like BANKSY^12^, as well as integration methods such as SEDR^13^, GraphST^14^, STAligner^15^ and STitch3D^16^ that are designed for horizontal integration, while SpatialGLUE^6^ performs vertical integration. Due to the respective assumptions on data availability for these integrations, none of these methods are usable for spatial mosaic integration. CellCharter^17^ and PRESENT^18^, are designed for spatial vertical integration and horizontal integration, but it is not able to handle the general case of spatial mosaic integration.

In this work, we present SpaMosaic for spatial mosaic integration. Using a contrastive learning framework^19^ with graph neural networks (GNNs), SpaMosaic projects each section into a modality-agnostic latent space in a spatially aware manner while minimizing batch variations. In particular, we adopted a weighted light graph convolution network to encode the similarity between spots. The output latent embedding is usable for imputation of missing data modalities and downstream analyses like spatial domain identification and joint analysis of multiple sections. We first employed simulated datasets to demonstrate SpaMosaic’s capabilities at integrating mosaic datasets while reducing technical noise and recovering hidden spatial cluster patterns. We then benchmarked SpaMosaic with experimentally acquired data of both human and mouse tissues with data modalities encompassing RNA, ATAC, histone modifications, and protein. Compared to other mosaic integration methods, SpaMosaic showed superior performance in terms of modality alignment, batch integration and spatial domain identification. Furthermore, by benchmarking with multiple complex real-world mosaic datasets from diverse biological contexts, we further demonstrated SpaMosaic’s versatility in integrating spatial omics sections that vary in modalities, resolutions, section sizes, and technologies, thereby enabling comprehensive and coherent analysis of heterogeneous spatial omics data.

## Results

### The SpaMosaic model

SpaMosaic is designed to integrate mosaic spatial omics data acquired from multiple tissue sections to output a joint latent space for downstream analyses (**Fig. 1a**). For input, SpaMosaic takes in the mosaic feature-by-spot (or cell) count matrices of different data modalities from multiple sections and associated spatial coordinates (**Fig. 1b**). The sections are acquired from the same tissue type but have no other restrictions. SpaMosaic is acquisition technology agnostic so the modalities can be acquired with different technologies, and it is capable of integrating sections of different sizes and spatial resolutions. To capture the links between modalities, SpaMosaic requires the modalities to be directly or indirectly connected through one or more sections. A direct connection appears in the same section, while an indirect connection requires multiple intermediary direct connections. Within each section and each modality, SpaMosaic first performs dimension reduction on the highly variable features with SVD to obtain a low-dimensional representation. We then use horizontal integration methods, such as Harmony^20^ and Seurat^21^, to remove the batch effects within each modality. Thereafter, we construct a heterogeneous adjacency graph for each modality. Each adjacency graph records two types of adjacencies, spatial distance-based adjacency and omics feature-based adjacency. The spatial distance-based adjacency denotes adjacency of spots with their nearest spatial neighbors from the same section, while the omics feature-based adjacency denotes adjacency between spots and their mutual nearest neighbors (MNNs)^22^ in other sections. To improve robustness, we leverage the spatial context and incorporate outlier detection algorithms to filter out high-noise MNN pairs.

**Fig. 1.**
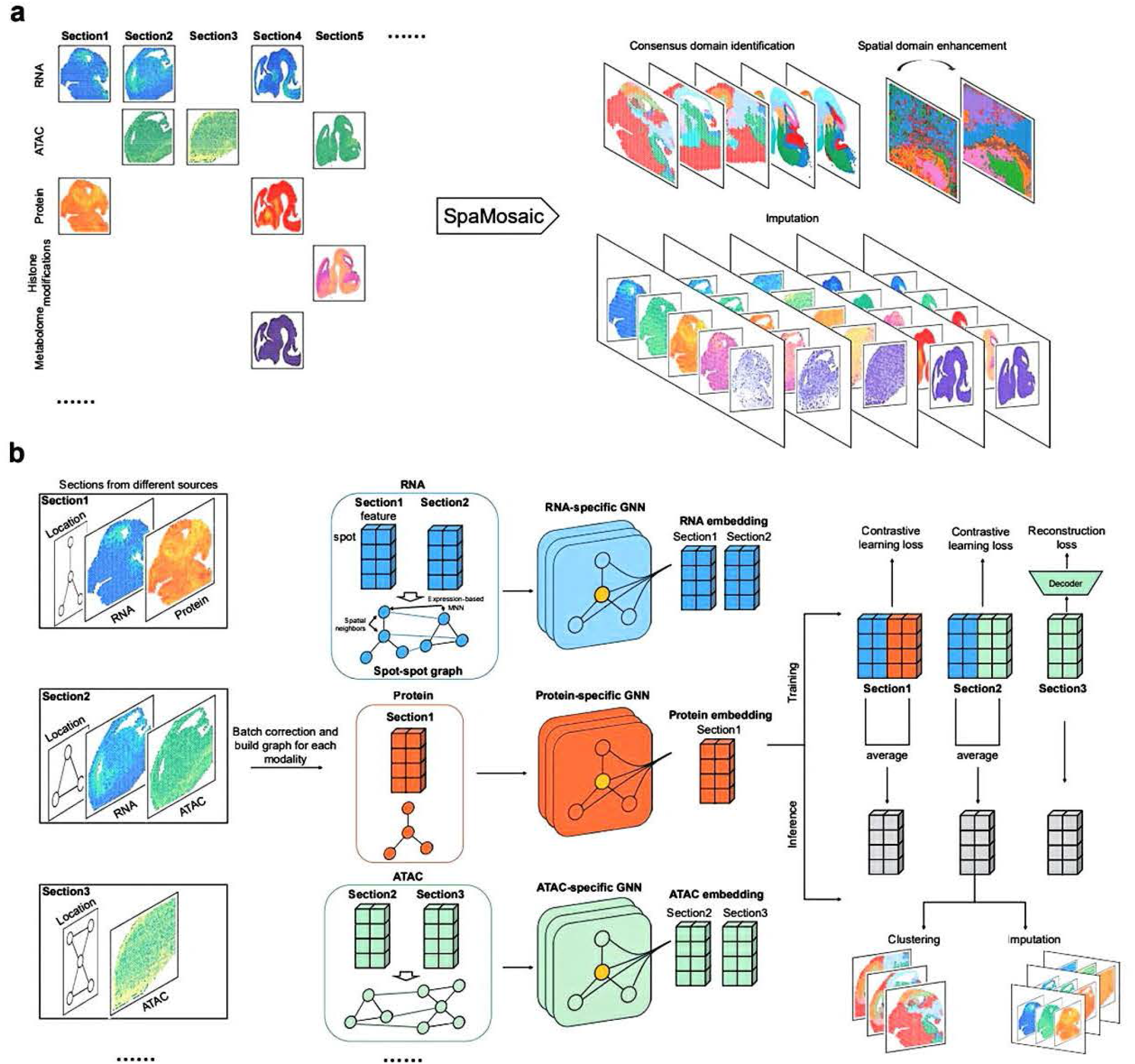
Overview of SpaMosaic framework. **a** SpaMosaic integrates multiple tissue sections with diverse modality compositions into a unified representation. It enables the identification of consensus spatial domains across sections, imputation of missing modalities, and enhancement of spatial domain resolution by revealing coherent anatomical structures that are obscured in single-modality analyses. The left panel presents an abstract representation of the general mosaic integration task, where sections with different modality compositions can be integrated. **b** Illustration of the SpaMosaic workflow using a specific example of spatial multi-omics mouse brain dataset integration. SpaMosaic first reduces data dimensionality and then performs batch effect correction to generate batch-corrected low-dimensional representations for each modality. Thereafter, a heterogeneous adjacency graph is constructed between spots within each modality. SpaMosaic employs graph neural networks to embed the modality-specific representations into modality-specific embeddings. The paired multi-modal embeddings are input for contrastive learning which aims to align embeddings from different modalities and constructs a modality-agnostic latent space. Finally, SpaMosaic infers the aligned embeddings for each section for downstream analysis (such as spatial domain identification) and imputation.

In the next step, SpaMosaic builds a modality-agnostic latent space using graph neural networks (GNNs) in a contrastive learning framework. For each modality, a GNN is used to encode modality-specific information from the modality inputs and adjacency graph into a set of embeddings. To encode the heterogeneous graphs, we explored four graph network architectures, the graph attention network (GAT)^23,24^, heterogeneous graph transformer (HGT)^25^ and two variants of light graph convolution network (LGCN)^26^. We selected the weighted LGCN (**Supp. Fig. S1a**) for the final model due to its efficiency and robust comprehensive performance (**Supplementary Information**). We then employ contrastive learning to align modality-specific embeddings. The training pairs are constructed from multi-modal sections as follows: embeddings from different modalities at the same spot form positive pairs, while embeddings from different spots within the same section form negative pairs, irrespective of their modalities. In conjunction, the contrastive learning objective is formulated to minimize the distance between positive pairs and maximize the distance between negative pairs. This seeks to align the different modalities while preserving variation between spots. After aligning the modalities, the average of the modality-specific embeddings will be used as the final embedding for sections with more than one modality. For sections with a single modality, we employ a feature reconstruction objective to preserve inter-spot variations. SpaMosaic outputs the integrated spot embeddings across sections with imputed feature matrices for the missing omics. This output is usable in downstream analyses such as spatial domain identification and analysis of feature correlations across modalities. To impute data for missing modalities, SpaMosaic uses omics feature information from the nearest neighbors identified in the modality-aligned latent space (details are provided in the Methods section and **Supp. Fig. S1b**).

### Benchmarking on simulated data

In this first example, we employed simulated data to evaluate the performance of SpaMosaic, an alternate version of SpaMosaic (non-spatial) that does not use spatial information, and five state-of-the-art single-cell mosaic integration methods, namely Cobolt^9^, scMoMaT^8^, StabMap^10^, CLUE^27^ and MIDAS^11^. We also evaluated BANKSY^12^ and CellCharter^17^ which are spatial domain (niche) identification methods. BANKSY and CellCharter are designed to integrate multiple sections within a single modality and hence considered to be indirect competitors with mosaic integration methods. Therefore, we included them as representative spatial clustering approaches to serve as performance reference for spatial omics approaches on this task. To create the simulated data, we followed the procedure of Townes et al.^28^, using five different sets of simulation parameters to each generate three replicates of spatial multi-omics datasets with different random seeds (**Supp. Fig. S2a**). For the different datasets, we varied the erosion parameter *s*^*rna*^ and dropout probability *p*^*rna*^ to modify the RNA expression data distribution and varied the background noise parameter *b*^*rna*^ to modify the protein expression data distribution (Methods). Each dataset has three sections where the first section consists of RNA and protein modalities while the second only has RNA expression and the third only has protein expression (**Supp. Fig. S2b**). We kept all modalities at the same dimensions, namely 1,296 spots arranged in a 36×36 grid. The simulated data has a ground truth of four spatial factors and a background (**Fig. 2a**). For all datasets, we reduced the RNA expression of factor 1 and protein expression of factor 4 to simulate a scenario where these factors were missing in their modalities. This aims to test the ability to recover all factors from integrating the two modalities. We also simulated batch effects between sections 1 and 2 in the RNA modality and sections 1 and 3 in the protein modality by varying the feature expression mean values in sections 2 and 3 (Methods, **Supp. Fig. S2c**). We then evaluated the methods in three aspects: preservation of spatial factors using the Adjusted Rand Index (ARI), percentage of abnormal spots (PAS)^29^ and spatial chaos score (CHAOS)^29^ metrics, batch mixing using Integration Local Inverse Simpson’s Index (iLISI)^30^, and modality alignment using the F1 score computed for label transfer accuracy^11^, which we computed for the second and the third sections. Higher scores indicate better performance for all metrics, except for PAS and CHAOS, where lower scores denote better performance.

**Fig. 2.**
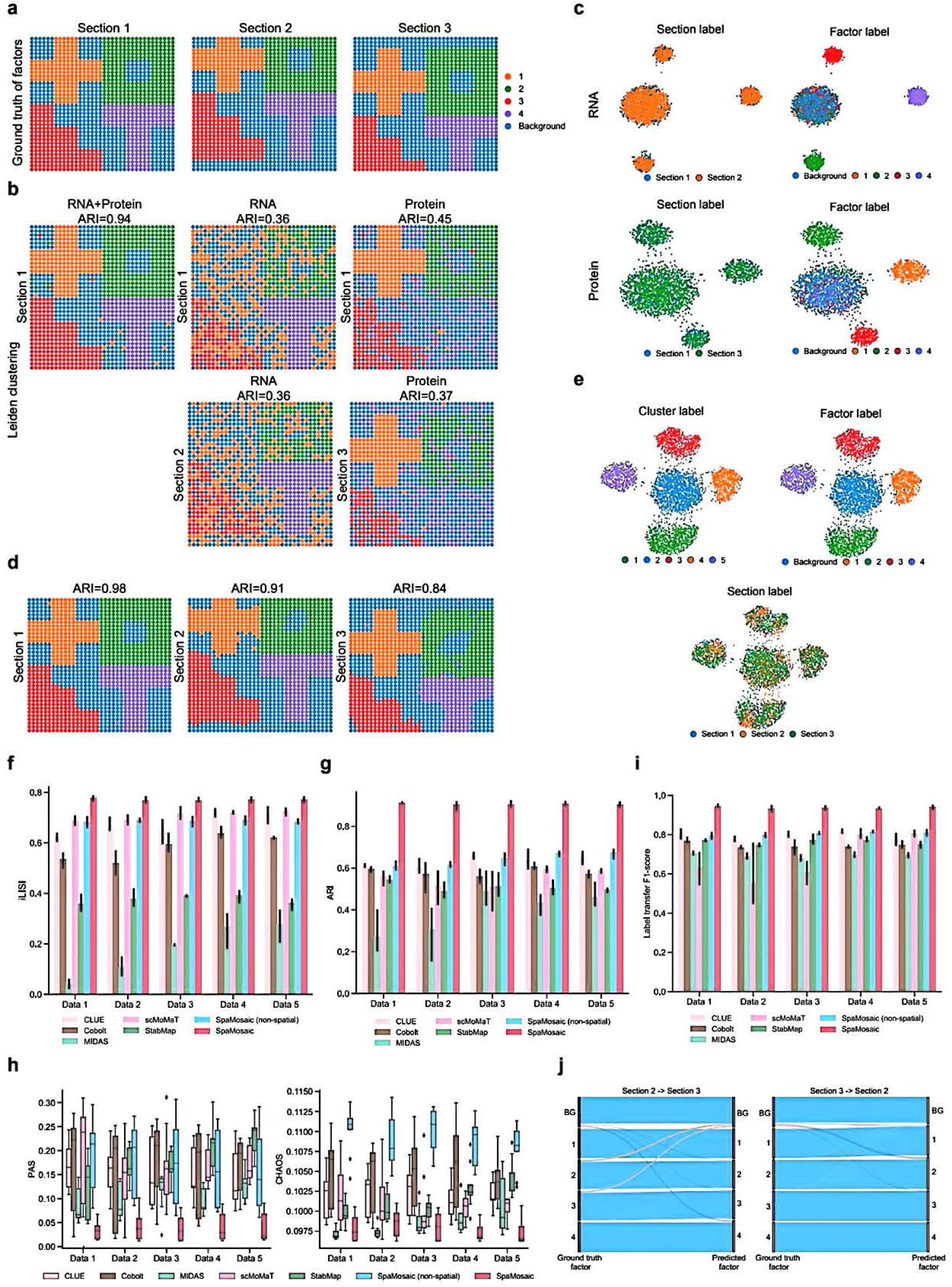
Benchmarking on simulated datasets. **a** Ground truth of spatial factors and background of the three sections. **b** Clustering patterns obtained with the concatenated RNA and protein profiles, RNA profiles and protein profiles, respectively. RNA features from sections 1 and 2 were jointly clustered using the Leiden algorithm, as were ADT features from sections 1 and 3. Multimodal clustering of section 1 was performed on concatenated RNA and ADT features using the Leiden algorithm. ARI scores were evaluated by comparing the clustering with the ground truth. **c** UMAP plots of batch-corrected RNA (sections 1 and 2) and protein (sections 1 and 3) profiles. Spots are colored by section (left) or factor (right). **d** Clustering patterns obtained using SpaMosaic’s embeddings for all three sections. **e** UMAP plots of SpaMosaic’s embeddings. Spots are colored by clustering, factor or section. **f** iLISI scores of benchmarked methods across five simulation data sets. The error bars represent the 950fo confidence interval around the score means. **g** ARI scores of benchmarked methods across the five simulation sets. The error bars denote the 950fo confidence interval around the mean of the scores. **h** PAS and CHAOS scores of benchmarked methods across the five simulation sets. In the box plot, the center lines indicate the median, boxes indicate the interquartile range, and whiskers indicate I.5x interquartile range. **i** Label transfer Fl-scores of benchmarked methods. **j** Sankey plots showing the comparison of prediction and ground truth of factors on the first replicate of the first simulation setting. BG denotes background.

We first examined the results of single modality clustering/ domain identification methods for one replicate from dataset 1 as an example. As the baseline, we used Leiden^31^ clustering with Harmony for batch integration (**Fig. 2b**). For section 1, Leiden clustering captured spatial factors 2, 3 and 4 in the RNA modality, factors 1, 2 and 3 in the protein modality, and all four factors from clustering on the concatenated RNA and protein data. For sections 2 (RNA only) and 3 (protein only), the clustering also captured the same respective factors in section 2 (factors 2, 3 and 4) and section 3 (factors 1, 2 and 3). The captured factors were not of equal quality as factors 1 and 4 were captured clearly in the protein and RNA modalities, respectively; meanwhile factors 2 and 3 were better captured in the protein modality but their identified clusters in the RNA modality were less spatially smooth. The UMAP^32^ plots (**Fig. 2c**) were consistent with factor 1 (RNA) and factor 4 (protein) merging into their respective modalities’ background. Next, we considered the spatial domain (niche) identification methods BANKSY and CellCharter. Specifically, we applied BANKSY to the RNA modality and CellCharter to both modalities. For sections 1 and 2 in the RNA modality, BANKSY and CellCharter both captured spatial factors 2, 3, and 4 (**Supp. Fig. S2d**). However, the clusters identified by BANKSY were noisier than CellCharter’s, especially factors 2 and 3, which are similar to the Leiden clustering. Comparatively, CellCharter captured more coherent spatial factors but split factor 2 into two clusters (**Supp. Fig. S2d**). For sections 1 and 3 in the protein modality, CellCharter identified spatial factors 1, 2, and 3, but was unable to clearly delineate factor 4 (**Supp. Fig. S2d**). The UMAP visualization further supported these observations by revealing overlaps between the background factor and other spatial factors (**Supp. Fig. S2e**). While it was not unexpected that BANKSY and CellCharter could not recover factors 1 (RNA) and 4 (protein), their recovery of the other factors suffered from the noise present.

We next considered the spatial clustering results obtained with SpaMosaic and competing mosaic integration methods. In SpaMosaic’s clustering results, all four factors were recovered from all three sections (**Fig. 2d**). The recovered factors also featured less noise and more accurate in terms of shape than the output of Leiden, BANKSY or CellCharter. In contrast, all competing mosaic integration methods except MIDAS successfully delineated the four factors in section 1, while MIDAS failed to capture the basic shape of factor 3 (**Supp. Fig. S3a**). For section 2, all competing methods captured factor 4 clearly, but factors 2 and 3 were captured with substantial noise, while factor 1 was only effectively captured by CLUE. For section 3, factor 1 was clearly captured, factors 2 and 3 were captured with substantial noise, and factor 4 was captured by CLUE; Cobolt and scMoMaT were only partially successful at capturing factor 4. Visualizing the SpaMosaic’s embeddings with UMAPs (**Fig. 2e**), we see the integration of all three sections within each factor’s cluster (bottom) while the spatial clusters show clear separation (upper left), and this compares well with the ground truth (upper right). However, the competing methods achieved only varying degrees of data integration (**Supp. Fig. S3b, bottom row**).

Using quantitative metrics, we compared SpaMosaic and competing methods on all five sets of simulated data with replicates. In terms of iLISI (batch integration), SpaMosaic was the best method for all five datasets and scMoMaT was in second place (**Fig. 2f**). The non-spatial form of SpaMosaic’s performance was on par with scMoMaT on datasets 1 and 2, and slightly behind for 3, 4 and 5. CLUE also performed close to scMoMaT on datasets 2, 4 and 5. For the ARI metric, SpaMosaic was the top performer for all datasets while CLUE and non-spatial SpaMosaic were second (**Fig. 2g**). Cobolt and scMoMaT also performed close to CLUE on datasets 1, 2 and 4, but fell behind for 3 and 5. For the PAS metric, SpaMosaic was again the top performer across all datasets (**Fig. 2h**), while it scored first in the CHAOS metric for datasets 4 and 5, and second to MIDAS for datasets 1, 2 and 3. We also computed the average ARI, PAS, and CHAOS scores for each section to observe section-specific performance. SpaMosaic was top for all sections for ARI and PAS, especially sections 2 and 3 (**Supp. Fig. S3c, d**). In terms of CHAOS, it was also the best on sections 1 and 2, and second for section 3 (**Supp. Fig. S3d**). Taken together, these results quantified SpaMosaic’s capture of spatially coherent and accurate factors, which we attributed to its incorporation of spatial information and batch effect removal capabilities, and SpaMosaic’s ability to integrate information from all three sections to recover the missing factors in sections 2 and 3.

Finally, to evaluate the cross-modal alignment capability of the integration methods, we performed label transfer assessment. In this assessment, we used the labels from one single-modality section to predict the corresponding labels in another single-modality section, testing whether each method could effectively align and transfer information across different modalities^11^. The F1-score metric showed SpaMosaic outperforming all other methods with the highest median score for all datasets (**Fig. 2i**). Non-spatial SpaMosaic and CLUE consistently ranked second on datasets 1-4, while on dataset 5, scMoMaT and CLUE were second ranked. We further investigated SpaMosaic’s label transfer by comparing the assigned factors to the ground truth with a Sankey plot for the first replicate of dataset 1 (**Fig. 2j**). Here we found that most of the incorrect assignments were spots of the spatial factors (1 to 4) being assigned to the background or vice versa, which was consistent with the spatial clustering obtained (**Fig. 2d, e**). Overall, SpaMosaic achieved the best performance in both visual comparisons and benchmarking metrics.

### Integration of intra-assay embryonic mouse brain sections across developmental stages

We next benchmarked SpaMosaic and competing methods to test their capabilities to handle experimentally acquired data. Here we used an experimental dataset consisting of mouse embryonic brain sagittal section data profiled with Misar-seq^33^ that jointly captures gene expression (RNA) and chromatin accessibility (ATAC). We selected the E13.5, E15.5 and E18.5 sections, and removed the ATAC modality of E13.5 and RNA modality of E18.5 to simulate mosaic data availability (**Fig. 3a**). Examination of the UMAPs of each modality showed the presence of strong batch effects between the different time points in both modalities (**Supp. Fig. S4a**). Following the same procedure in the previous example, we compared SpaMosaic with the other single-cell mosaic integration methods as well as spatial domain (niche) identification methods. In addition, we included MultiVI^34^, a framework designed for mosaic integration of RNA and epigenomic modalities.

**Fig. 3.**
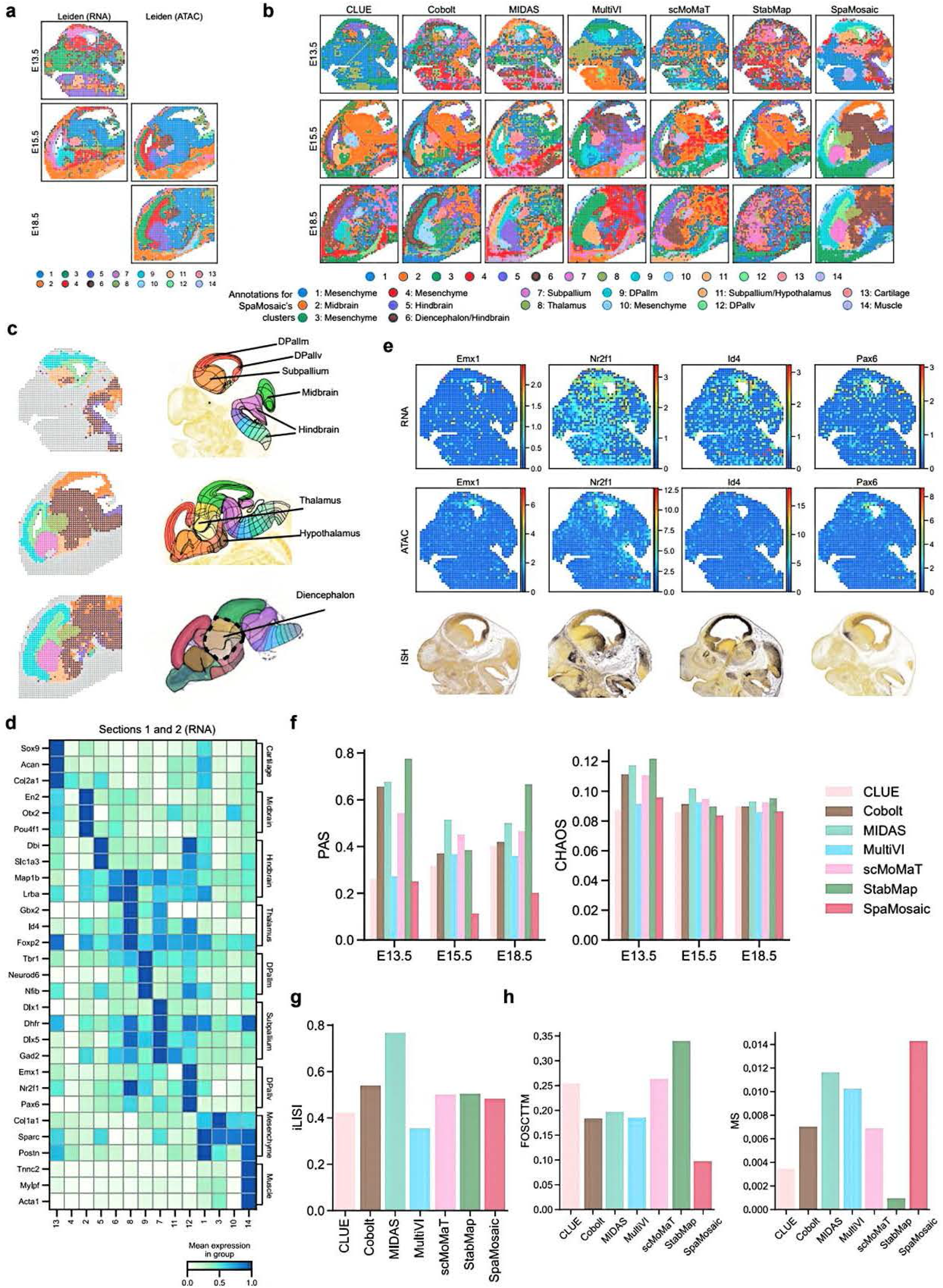
Downstream analysis results obtained with SpaMosaic and quantitative benchmarking on an embryonic mouse brain (Misar) dataset. **a** Mosaic dataset and clustering patterns for all three sections: El3.5 without the ATAC modality, EI5.5 with both RNA and ATAC expression profiles, and EI8.5 without the RNA modality. Modality-specific clustering was jointly performed across relevant sections using the Leiden algorithm on batch-corrected representations. **b** Spatial plots of the clusters obtained from mosaic integration methods for the three sections. Manual annotation was performed to identify SpaMosaic’s clusters. **c** Comparison of SpaMosaic clustering results with reference annotations from the Allen Reference Atlas for the corresponding three developmental stages. Reference images were cropped from the reference atlas (https://atlas.brain-map.org/). **d** RNA expression of marker genes specific to the brain regions captured by SpaMosaic’s clusters. **e** Gene activity scores of DPallv-specific marker genes *(Emxl, Emx2, ld4, Nr2fl*, and *Pax6)* in SpaMosaic’s clusters. The bottom row shows the ISH images from the Allen Brain Atlas for the corresponding markers. Reference images were cropped from the Allen Brain Atlas (https://developingmouse.brain-map.org/). **f** PAS and CHAOS scores of benchmarked methods across the three sections. **g** iLISI scores of benchmarked methods. **h** Modality alignment scores (FOSCTTM and MS) of benchmarked methods.

We first considered the single modality clustering methods. The baseline Leiden clustering successfully identified major brain regions like the mantle zone of dorsal pallium (DPallm; cluster 7 in RNA, 3 in ATAC), ventricular zones of dorsal pallium (DPallv; cluster 10 in RNA, 4 in ATAC), hindbrain (cluster 1 in both modalities), and subpallium (cluster 9 in RNA, 5 in ATAC) across sections, but showed noisy cluster boundaries. For CellCharter and BANKSY that are tailored for spatial omics, their clusters showed improved spatial coherence compared to the Leiden clustering (**Supp. Fig. S5a and S4b**). However, the two methods showed markedly different performance in distinguishing known anatomical regions. BANKSY’s performance was largely poor, merging distinct brain regions with other structures and showing inconsistent cluster assignments for most anatomical regions across sections. For example, BANKSY merged DPallv and DPallm with the hindbrain and midbrain regions in the E13.5 and E15.5 sections. In contrast, CellCharter achieved more biologically relevant segmentation of major brain structures. It successfully identified the DPallm and DPallv in both modalities of the E15.5 section, and accurately identified the subpallium (cluster 10), thalamus (cluster 6), and hypothalamus (cluster 7) in the ATAC modality. However, CellCharter’s limitation in handling mosaic datasets led to misaligned clusters between RNA and ATAC within the same section, making it difficult to achieve unified clustering across the entire dataset.

Next, we examined the mosaic integration methods’ output. Among the mosaic integration methods, SpaMosaic’s spatial clusters had the least noise and best matched the Allen Brain Atlas reference^35^ in terms of the brain regions captured (**Fig. 3b, c**). Across all developmental stages, SpaMosaic successfully identified the most comprehensive set of brain regions with well-defined boundaries, including the midbrain, hindbrain, diencephalon, subpallium, hypothalamus, thalamus, DPallm and DPallv (detailed cluster assignments shown in **Fig. 3b**). In comparison, the other mosaic integration methods showed varying degrees of limitations. MultiVI identified the subpallium across all three sections and thalamus in E15.5 and E18.5, but frequently merged other distinct brain regions (e.g., DPallm with DPallv, hindbrain with midbrain). The performance of CLUE, Cobolt and MIDAS were also poor, only capturing certain major brain regions like the DPallm, DPallv, thalamus and subpallium. Their remaining clusters were highly noisy and could not be matched to the reference structures. Furthermore, these methods produced inconsistent cluster assignments across sections. For example, CLUE assigned the hindbrain region to clusters 1, 2, and 7 in the three sections respectively. scMoMaT successfully captured the DPallm, DPallv, subpallium and thalamus in the E15.5 and E18.5 sections but failed to delineate the DPallm and DPallv in E13.5. StabMap identified the DPallm and DPallv in the E15.5 section, while its clusters in the E13.5 and E18.5 sections could not be matched to known anatomical references.

Our visual comparison of SpaMosaic’s clusters with the Allen Brain Atlas’ reference structures showed high concordance but we followed up with further validation using known marker genes (**Fig. 3d**). For instance, the cartilage marker *Sox9*^*36*^ was highly expressed in cluster 13 (RNA, sections 1 and 2), while the midbrain markers *En2*^*33*^ and *Otx2*^*37*^ were enriched in cluster 2 (RNA, sections 1 and 2). Among the clusters, 9 and 12 in E13.5 were particularly notable. Comparison with the Allen Brain Atlas revealed that these clusters corresponded well to the DPallm and DPallv regions (**Fig. 3c**), which were neither captured by CellCharter and BANKSY, nor annotated in the original publication (**Supp. Fig. S5b**). Among the other mosaic integration methods, CLUE, Cobolt, and MIDAS could also demarcate these regions. To confirm their identities, we visualized the expression of markers *Emx1, Id4, Nr2f1* and *Pax6* that regulate mouse neocortex development^38-40^ and are highly expressed in the DPallv or pallium region. We find them highly expressed in cluster 12, both in RNA count and gene activity scores derived from the ATAC modality (**Fig. 3d, e**). Comparison with ISH staining samples from the Allen Brain Atlas^41^ also showed matching locations, confirming cluster 12’s accuracy in capturing the DPallv (**Fig. 3e**).

Finally, we quantitatively evaluated the mosaic integration methods’ performance in three aspects: spatial domain identification (PAS and CHAOS), batch mixing (iLISI) and modality alignment (Fraction of Samples Closer than True Match, FOSCTTM^42^ and Matching Score, MS^43^). Higher scores indicate better performance for iLISI and MS, while the reverse is true for FOSCTTM, PAS and CHAOS. To calculate the FOSCTTM and MS metrics, we created a new dataset using the E15.5 and E18.5 sections and split the RNA and ATAC modalities of the E18.5 section to simulate two new sections. The ground-truth pairing between the RNA and ATAC modalities of the E18.5 section was used to evaluate modality alignment. In terms of continuity of identified spatial domains (PAS and CHAOS), SpaMosaic scored the lowest for all sections in PAS, and for E15.5 and E18.5 in CHAOS (**Fig. 3f**). For batch mixing (iLISI), MIDAS was first, while Cobolt was second and scMoMaT and StabMap closely behind at third place. SpaMosaic was ranked fourth but close to scMoMaT and StabMap in performance (**Fig. 3g**). These results prompted us to investigate the batch mixing by examining the UMAPs colored by section and clusters (**Supp. Fig. S4c**). MIDAS indeed mixed the three sections very evenly, but the clusters also showed substantial overlapping. The remaining methods had better cluster separation but lower batch mixing. For modality alignment, SpaMosaic was the best at both FOSCTTM and MS metrics (**Fig. 3h**), highlighting its capabilities in cross-modality alignment.

### Integration of postnatal mouse brain sections with section-size imbalance

In our next scenario, we employed a postnatal mouse brain dataset (spatial transcriptome and histone modification co-sequencing^2^ + spatial Cut&Tag-seq^1^; spRH+spC&T), which contains data of RNA and a different epigenomic modality, the H3K4me3 histone modification. This compiled dataset consists of three sections acquired from different studies (**Fig. 4a**). The first section consists of the transcriptome and histone modification (H3K4me3) profiles of a P22 mouse brain section in a capture area with 100×100 pixels, each 20 µm in diameter^2^. For the second section, the original experiment acquired the spatial transcriptome and histone modification (H3K27ac) of a P22 mouse brain section with a capture array of 100×100 pixels, each 20 µm in diameter^2^, of which we only retained the spatial transcriptome. For the third section, we selected the spatial histone modification (H3K4me3) profile of a P21 mouse brain section, acquired on a 50×50 array with 20 µm pixels^1^. These three datasets were acquired from mouse brain sections at different ages and with different capture area sizes, which posed additional challenges for data integration. From the UMAP plots, we can easily observe batch effect present, especially between section 1 and section 3 in their histone modification profiles (**Supp. Fig. S6a**). Again, we benchmarked SpaMosaic alongside the other state-of-the-art integration and spatial domain (niche) identification methods. Although Cobolt was originally only demonstrated on RNA and ATAC modalities, there is no methodological restrictions on its generalizability towards other omics modalities. Therefore, we tested it alongside the other integration algorithms.

**Fig. 4.**
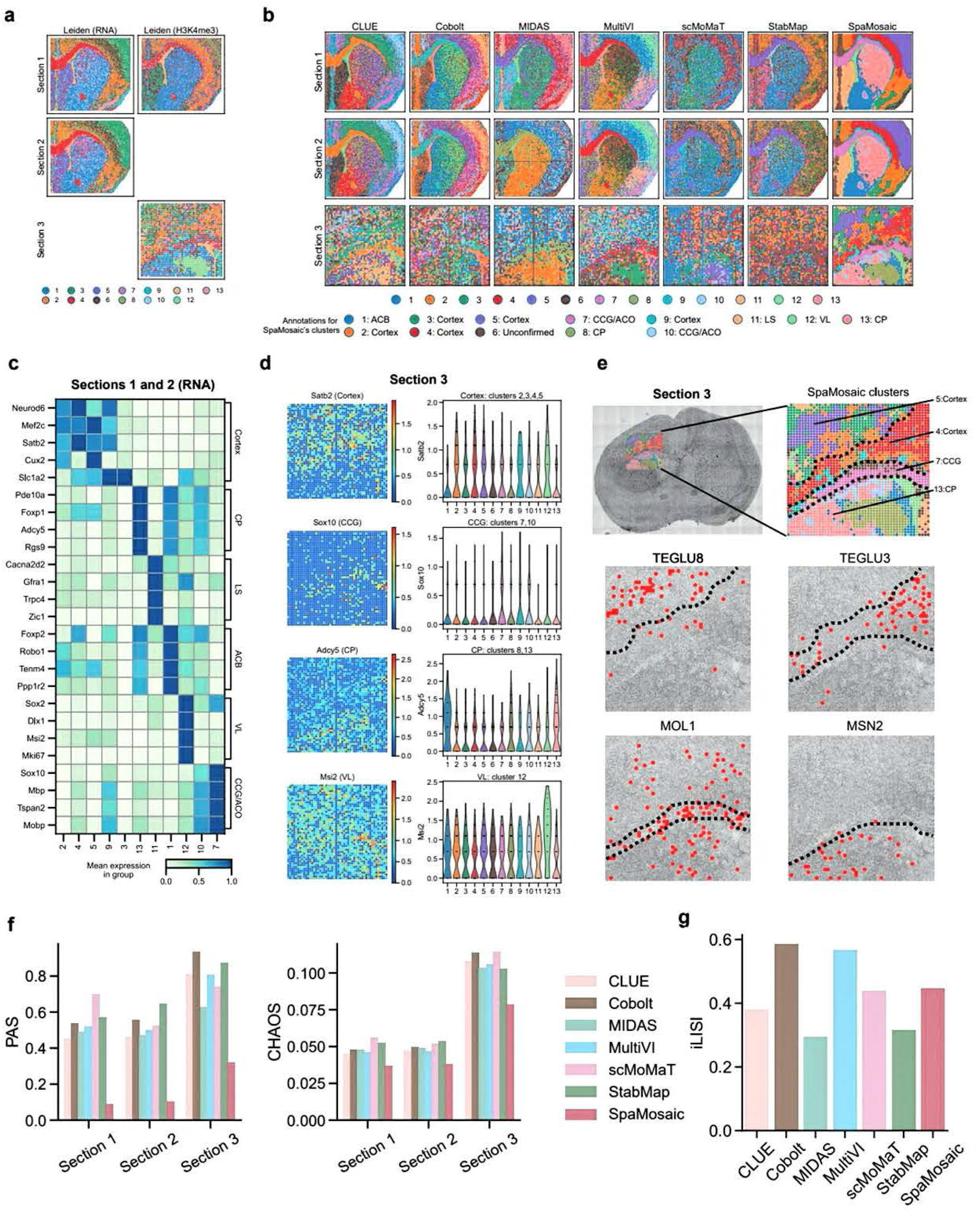
Downstream analysis results obtained with SpaMosaic and quantitative benchmarking on a postnatal mouse brain (spRH+spC&T) dataset. **a** Mosaic dataset configuration and clustering patterns obtained for all three sections. Modality-specific clustering was jointly performed across relevant sections using Leiden clustering on the batch-corrected representations. **b** Spatial plots of the clusters obtained from benchmarked methods for all three sections. Manual annotation was performed to identify SpaMosaic’s clusters. **c** RNA expression of marker genes specific to the six brain regions captured by SpaMosaic’s clusters. **d** Spatial heatmap and violin plots of gene activity scores (GAS) for region specific markers on the third section and in SpaMosaic’s clusters, respectively. **e** Spa.Mosaic’s clusters overlaid on the image of the third section and cell types identified by label transfer. **f** PAS and CHAOS scores of benchmarked methods across the three sections. **g** iLISI scores of benchmarked methods.

Using the Allen Brain Atlas^35^ as reference, we found that SpaMosaic produced clustering results that aligned more closely with known anatomical structures compared to the other mosaic integration methods. (**Fig. 4b and Supp. Fig. S7a**). In sections 1 and 2, we identified the cortex (clusters 2, 3, 4, 5, and 9), lateral ventricle (VL) (cluster 12), nucleus accumbens (ACB, cluster 1), genu of corpus callosum (CCG) and anterior commissure of olfactory limb (ACO) in cluster 7, caudoputamen (CP, cluster 13), and cluster 11 capturing the lateral septal complex (LS). In section 3, we could identify the VL (cluster 12), CCG (cluster 7), cortex (clusters 2, 3, 4, and 5), and CP (clusters 8 and 13). In contrast, the competing methods produced noisier clusters and were comparatively less successful at capturing the brain structures. All methods, except scMoMaT and StabMap, captured the cortex layers in sections 1 and 2; however, none of them were able to explicitly separate the ACB and CP. Notably, Cobolt identified cortical layers as clusters 4, 5, and 11, demonstrating finer cortical stratification compared to SpaMosaic. For section 3, we could not unambiguously match the competing methods’ output clusters to the reference. We also tested BANKSY and CellCharter, which achieved more coherent spatial clustering compared to the other mosaic integration methods (**Supp. Fig. S7b**). However, BANKSY identified only a single cortex layer and did not distinguish between the ACB and CP. CellCharter’s output revealed spatial patterns similar to SpaMosaic in the RNA modality of sections 1 and 2, but it split the CP into two clusters and failed to differentiate the ACB and CP in section 1’s H3K4me3 modality. On section 3, CellCharter produced more coherent spatial clustering than SpaMosaic. However, the cortex was assigned to different clusters in sections 1 and 3, preventing proper cross-section correspondence. The UMAP plots confirmed this misalignment (**Supp. Fig. S6b**).

In addition to referencing the Allen Brain Atlas, RNA expression of marker genes for the six brain regions in sections 1 and 2 was visualized (**Fig. 4c**). The expected markers were highly expressed in the corresponding SpaMosaic’s clusters, affirming their identities. For instance, the striatum markers *Pde10a*^*2*^, *Adcy5*^*6*^ and *Foxp1*^*44*^ were highly expressed in cluster 13 (CP), which is part of the striatum. We also observed the markers of CCG, *Sox10*^*2*^, *Mbp*^*2*^ and *Tspan2*^*2*^ to be highly expressed in cluster 7 that contains the CCG structure. For section 3 that lacked RNA modality, we visualized the spatial plots and violin plots of gene activity scores derived from the histone marks of relevant marker genes (**Fig. 4d**). The gene activity scores showed high levels of noise, but we could observe region-specific expression patterns of key markers, such as *Satb2*^*2*^ in the cortex and *Adcy5* in the CP/striatum. Violin plots also illustrated that gene activity patterns aligned with the identified spatial domains. To further verify the identified brain structures, we used Seurat^21^ to integrate section 3’s gene activity scores (GAS) with a published scRNA-seq mouse nervous system dataset^45^ and transferred the annotated cell types to section 3. We then visualized the spatial distribution of four cell types, each with a unique spatial distribution (**Fig. 4e**). We found the mature oligodendrocytes (MOL1) enriched in the identified CCG region and the medium spiny neurons (MSN2) located only within the CP/striatum, which is consistent with existing studies^1,45^. We also observed that the TEGLU8 excitatory neurons were enriched in cluster 5 (cortex) that matched the upper cortical layer, and the TEGLU3 excitatory neurons were enriched in cluster 4 (cortex) that encompassed the deep cortical layer^45^.

We next employed the metrics PAS and CHAOS to assess the spatial continuity of the identified domains, and iLISI to evaluate batch integration. Quantitative evaluation of modality alignment was not possible due to the lack of another section with paired RNA and H3K4me3 histone modification measurements. For spatial domain identification, SpaMosaic clearly outperformed the competing methods for all three sections (**Fig. 4f**). In terms of batch mixing, Cobolt and MultiVI ranked as the top two methods, followed by SpaMosaic, which slightly outperformed scMoMaT (**Fig. 4g**). We further examined the UMAP visualizations colored by the clustering labels and section labels (**Supp. Fig. S6c**). For the cluster labelled plots, the consistent grouping and clear separation of the SpaMosaic’s clusters corroborated with its PAS and CHAOS scores, especially for sections 1 and 2. In contrast, Cobolt’s clusters showed observable mixing while scMoMaT’s output showed intra-cluster separations. In the visualization with section labels, the even mixing achieved by Cobolt and MultiVI matched their iLISI scores, while SpaMosaic, StabMap, and CLUE achieved better mixing than the remaining methods albeit with small local clusters of batch specific spots still visible.

### Integration of a mosaic dataset of the postnatal mouse brain with diverse epigenomic layers

Here we consider another scenario with a postnatal mouse brain dataset (spatial RNA-ATAC-seq + spatial transcriptome and histone modification co-sequencing; spRA+spRH) consisting of four sections with diverse epigenomic measurements including ATAC and different histone modifications, in addition to the RNA modality bridging all samples^2^. All sections have a capture area of 100×100 pixels, each 20 µm in diameter (**Fig. 5a**). In addition to the transcriptome, the first three sections measured the H3K27me3, H3K4me3 and H3K27ac histone modifications, respectively, and the fourth section captured ATAC-seq profiles. For method evaluation, we included UINMF^46^ which can handle this type of dataset with a common bridging modality, but excluded MIDAS as it cannot simultaneously integrate ATAC with other epigenomic modalities. In our pre-integration analysis, we observed batch effects present among the RNA profiles (**Supp. Fig. S8a**).

**Fig. 5.**
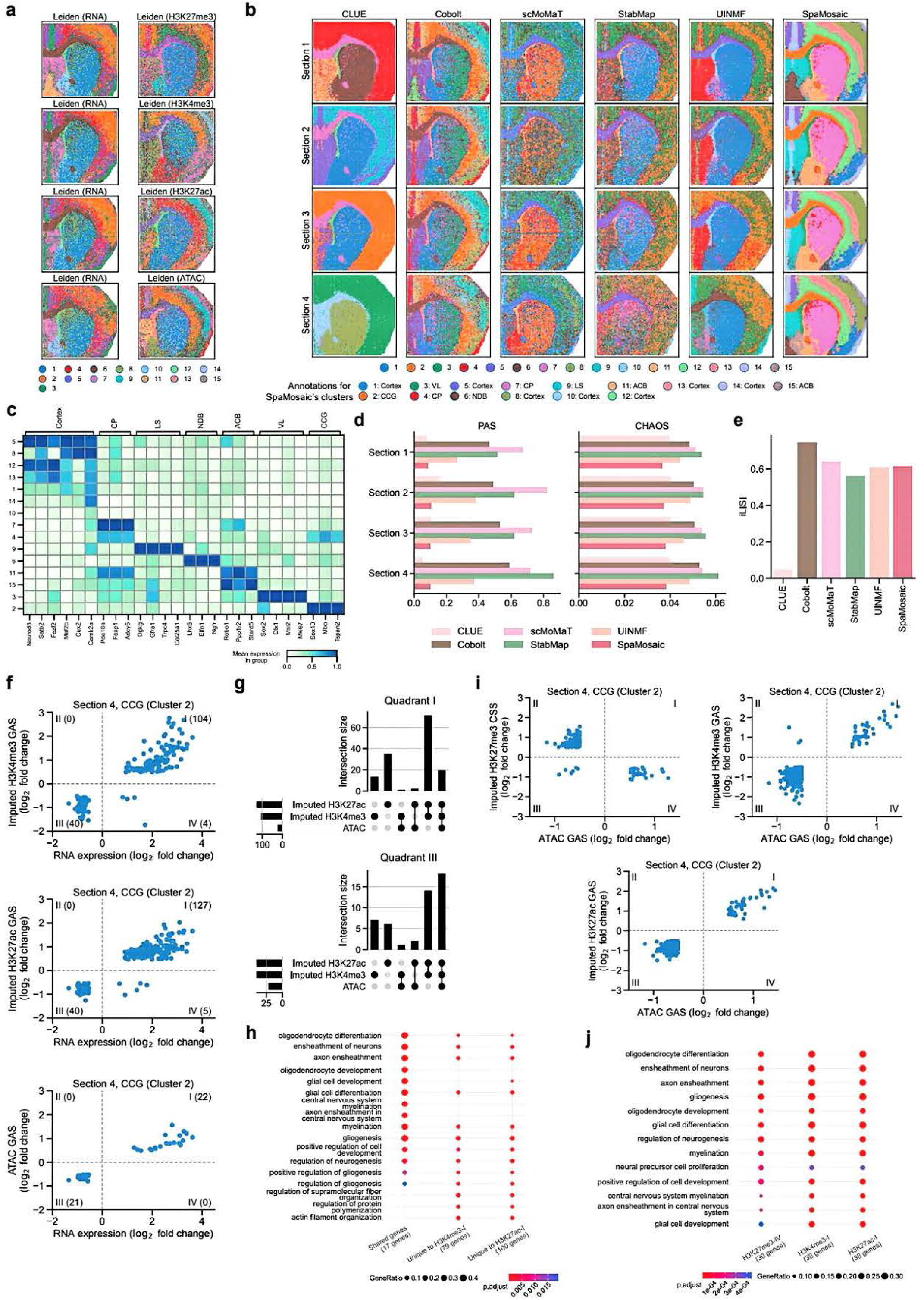
Application of SpaMosaic to postnatal mouse brain dataset (spRA+spRH). **a** Mosaic dataset configuration and clustering patterns of the four sections. The RNA cluster labels were obtained by performing Leiden clustering on the batch-corrected low-dimension embeddings, and each epigenomic modality’s cluster labels were obtained by performing Leiden clustering on the principal components of the epigenomics profiles. **b** Spatial plots of clusters obtained from the methods for all four sections. Manual annotation was performed to identify SpaMosaic’s clusters. **c** RNA expression of brain region markers in SpaMosaic’s clusters. **d** PAS and CHAOS scores of benchmarked methods across the four sections. **e** iLISI scores of benchmarked methods. **f** Relationships between the measured ATAC GAS, imputed H3K4me3 GAS and imputed H3K27ac GAS with the RNA in the CCG region of the fourth section. Only region-specific genes were kept for all modalities if their adjusted *p* value was less than le-5 and the absolute value of log2 fold change was larger than 0.5. **g** Upset plot of genes in quadrant I and quadrant III of all comparisons in **f. h** Enriched GO terms for genes shared across quadrant I of all comparisons, as well as for genes unique to quadrant I of the H3K27ac and H3K4me3 comparison in **f. i** Relationships between the imputed H3K27me3 GAS, imputed H3K4me3 GAS and imputed H3K27ac GAS with measured ATAC GAS in the CCG region of the fourth section. Only region-specific genes were kept if their adjusted *p* value was less than le-5 and the absolute value of log2 fold change was larger than 0.5. **j** Enriched GO terms for the genes in quadrant IV of H3K27me3 vs ATAC, quadrant I of H3K4me3 vs ATAC and quadrant I of H3K27ac vs ATAC comparisons in **i**.

We first visually examined the clusters obtained from the mosaic integration methods (**Fig. 5b**). SpaMosaic’s clusters clearly captured the different brain regions that we could match to the reference Allen Brain Atlas^35^ (**Supp. Fig. S8b**). For example, we identified the diagonal band nucleus (NDB, cluster 6), CCG and ACO (cluster 2), CP (clusters 4 and 7), LS (cluster 9), ACB (cluster 11) and VL (cluster 3). Plotting previously described brain markers, we found them highly expressed in their respective annotated clusters, such as *Mbp* and *Tspan2* in the CCG, *Stard5*^*47*^ in the ACB, and *Trpc4* and *Dgkg*^*48*^ in the LS (**Fig. 5c**). For the cortex layers, we found *Fezf2*^*2*^ highly expressed in the deep cortical layers (clusters 12 and 13) and *Cux2*^*2*^ in the upper layers (cluster 8). In general, the competing methods captured noisier clusters, except for CLUE. However, CLUE’s clusters captured fewer brain regions than SpaMosaic, being unable to separate the CP and ACB or even distinguish the cortex layers. For Cobolt, scMoMaT, StabMap and UINMF, they produced noisier clusters, and none could separate the CP and ACB. While we could discern cortex layers in Cobolt’s and UINMF’s clusters, scMoMaT’s and StabMap’s clusters were even noisier with no cortex layers identifiable. We next examined the outputs of BANKSY and CellCharter. Both identified more continuous spatial domains compared to the mosaic integration methods, especially for the RNA modality (**Supp. Fig. S9**). However, BANKSY captured only a single cortex layer and failed to distinguish between regions such as ACB and CP, or LS and NDB. CellCharter (RNA) recovered spatial structures broadly consistent with the Allen Brain reference, including the cortex layers (clusters 1, 4, 13, and 14), CCG and ACO (cluster 9), CP (cluster 15), VL (cluster 6), and LS (cluster 2), but mixed NDB with the cortex layers (cluster 1). Additionally, the cortical regions were assigned to different clusters by CellCharter (RNA) in section 4 compared to the other three sections, which is consistent with the UMAP visualizations, despite the inputs having been batch corrected with scVI^49^ (**Supp. Fig. S10a**). For the epigenomic modalities, CellCharter produced considerably noisier spatial domains, likely due to high intrinsic data noise and the absence of cross-section and cross-modality integration.

We next quantified spatial domain identification performance with the PAS and CHAOS metrics and batch integration with iLISI. For PAS, SpaMosaic scored a joint first with CLUE on sections 1 and 3, first on section 2, and second on section 4 (**Fig. 5d**). Taken together, SpaMosaic and CLUE performed similarly but CLUE identified fewer brain regions as discussed in our earlier analysis. For CHAOS, SpaMosaic attained the lowest scores for all sections. For batch mixing assessed with iLISI, Cobolt was top with scMoMaT second and SpaMosaic a close third with UINMF (**Fig. 5e**). We also visually inspected the UMAPs (**Supp. Fig. S10b**), where we see that the batch mixing broadly reflects the metrics. For example, the batches were well mixed for Cobolt’s output, while there was clear batch specific separation for CLUE.

In addition to data integration and learning modality agnostic latent embeddings in a spatially aware manner, SpaMosaic can also impute the missing data modalities. We first evaluated SpaMosaic’s imputation performance by benchmarking it together with MIDAS, BABEL^50^, TotalVI^51^ and MultiVI on four datasets, and measured their performance using the Pearson correlation coefficient, correlation matrix distance^52,53^, and area under the receiver operating characteristic curve (AUROC). We found that SpaMosaic outperformed other methods in preserving spot-spot correlations and effectively maintained feature-feature correlations. Full details on the employed datasets, metrics, and results are provided in the **Supplementary Information**. Next, we used the five-modal dataset of mouse brain sections to further demonstrate the imputation capabilities of SpaMosaic. Specifically, we used SpaMosaic to impute the gene activity scores (GAS) or chromatin silencing scores (CSS)^54^ of the missing epigenomic profiles. We then compared the correlation between RNA expression and measured or imputed GAS or CSS of each section in a cluster-specific manner. Prior to analysis, we applied filtering thresholds to retain only region-specific genes with an adjusted *p*-value (*p*-adj) less than 10^-5^ and absolute log_2_ fold change (l2fc) greater than 0.5, as shown in **Supp. Fig. S11a-d**. Taking the CCG as an example, the measured H3K27me3 CSS had anticorrelation with the RNA modality on section 1 as H3K27me3 is a suppressor, and the imputed H3K27me3 CSS also showed similar anticorrelation with the RNA modality of the other sections (**Supp. Fig. S11a**). For section 2, the measured H3K4me3 GAS was positively correlated with the RNA modality, which matches H3K4me3’s functionality as a promoter. The imputed H3K4me3 GAS of the other sections likewise showed positive correlations. Similarly, the imputed H3K27ac and ATAC modalities also showed positive correlations with the RNA. We also examined the correlations of the measured and imputed epigenomic profiles with the RNA in other clusters like the VL (cluster 3), CP (cluster 7) and cortex (cluster 8) (**Supp. Fig. S11b-d**), which we found to be consistent. This showed that SpaMosaic’s latent representation captured the relationships between modalities and preserved them in the imputation.

To further investigate the imputation results of the CCG cluster, we selected section 4 to compare the correlations between the measured RNA with GAS computed from the measured ATAC modality and the imputed histone modification modalities. Using the same filtering thresholds (*p*-adj < 10^-5^, l2fc > 0.5) as above, we observed that the imputed histone modifications’ (H3K4me3 and H3K27ac) GAS of the fourth section had more genes correlated with the RNA modality compared to the measured ATAC GAS (**Fig. 5f**). We then compared the genes and found that the correlated genes (in quadrant I and III) for the measured ATAC assay were also found in the imputed histone modification modalities (**Fig. 5g**). This showed that the imputed GAS preserved their relationships with the measured gene signals. It is also worth noting that the overall stronger correlation between the imputed histone modifications and RNA compared to ATAC-RNA correlation stems from the fact that RNA serves as the bridge modality that is explicitly aligned with all other modalities, resulting in imputed profiles that inherit stronger associations with the RNA expression patterns. We then investigated the other genes in the imputed histone modification modalities that were not found in the measured ATAC assay. Among the quadrant I correlated genes, we selected those unique to the imputed H3K4me3 or H3K27ac for GO enrichment analysis with clusterProfiler^55^, and found that these genes were associated with oligodendrocyte differentiation and myelination (**Fig. 5h**), which is in accordance with previous findings^2^. Across the three epigenomic modalities, 17 genes in quadrant I were shared, and they are highly expressed by oligodendrocytes that are enriched in the mouse brain’s corpus callosum. These genes include key transcriptional regulators involved in oligodendrocyte development and differentiation (*Sox2*^*56*^, *Sox10*^*57*^, *Olig1*^*58*^, and *Zeb2*^*59*^) and genes encoding major myelin structural proteins (*Mobp*^*58*^, *Plp1*^*58*^, *Mag*^*58*^, and *Mal*^*58*^). These genes have also been shown to be promoted by H3K4me3 for expression^60,61^. Additional oligodendrocyte-enriched genes unique to H3K27ac or H3K4me3 quadrant I include genes encoding myelin structural proteins (*Ermn*^*62*^, *Cnp*^*58*^, *Mog*^*58*^, *Opalin*^*58*^) and metabolism-related proteins (*Enpp6*^*63*^, *Sirt2*^*64*^). We also found genes in quadrant IV which have an inverse relationship between RNA expression and imputed histone mark’s GAS. Between the RNA and H3K4me3 modalities, there were four genes (*Kif5a, Mt3, Mir682*, and *Clu*), and for RNA and H3K27ac, five genes (*Camk2n1, Igfbp5, Rps23, Rpl26, Lrrc17*), which have not been reported to be under the control of the respective histone marks. Moreover, these histone marks are associated with activation. Thus, we have been unable to ascertain any mechanistic explanations behind these observations.

Spatial mapping of the imputed histone modification GAS of CCG marker genes indicated that the imputed values to clearly demarcate the CCG region, which was more spatially coherent than clusters of the measured ATAC GAS (**Supp. Fig. S12**). We believe that this was due to the information aggregation from the other sections, as well as the incorporation of spatial information in the imputation process, which fuses gene signals within each region and enhances them. In addition to the CCG cluster, we also examined the correlations for other brain regions in section 4, namely the VL (cluster 3), CP (cluster 7) and cortex (cluster 8), and we likewise observed similar signal enhancement (**Supp. Fig. S11e**).

In addition to the correlations between the transcriptomic and epigenomic modalities, we also investigated the relationships among different epigenomic modalities that were not co-measured from the same section (e.g., imputed histone modifications versus measured ATAC). We again analyzed the CCG cluster in section 4, focusing on the correlations between the imputed histone modification assays and the measured ATAC GAS. Using the same filtering thresholds (*p*-adj < 10^-5^, l2fc > 0.5) as described above, the imputed histone mark GAS or CSS showed the expected correlations with the measure ATAC GAS (anticorrelated with repressor H3K27me3, correlated with promoters H3K27ac and H3K4me3) (**Fig. 5i)**. We selected the anticorrelated genes in quadrant IV (H3K27me3) and correlated genes in quadrant I (H3K4me3 and H3K27ac) and performed GO enrichment for each set. The enriched processes included oligodendrocyte differentiation, oligodendrocyte development and myelination, which were shared by all three sets (**Fig. 5j**). We further focused on the 29 genes that were shared among quadrant IV of H3K27me3 and quadrant I of both H3K4me3 and H3K27ac. These included genes encoding oligodendrocyte transcription factors (*Sox10, Sox2, Olig1*, and *Olig2*^*58*^) and constituents of myelin sheaths (*Cldn11*^*58*^, *Fa2h*^*65*^, and *Ugt8a*^*66*^). Interestingly, *Mbp* and *Plp1* that encode major myelin structural proteins were absent from this shared gene set, suggesting their regulation may involve other epigenetic mechanisms, such as DNA methylation^67^. This analysis was also extended to the CP (cluster 7) and cortex (cluster 8) which showed similar correlation patterns, and we again observed much overlap in the GO terms obtained between the different sets of genes (**Supp. Fig. S13**). These results give us confidence in the imputation’s accuracy.

### Cross-technology integration of mouse brain data across heterogeneous sizes and resolutions

We further benchmarked SpaMosaic using two mosaic integration scenarios of mouse brain acquired with different technologies, size and resolutions (**Supp. Fig. S14a**). Scenario 1 uses an adult mouse brain dataset composed of three sections profiled by distinct technologies: two RNA only sections acquired with the 10x Genomics Visium HD^68^ and 10x Genomics Visium platforms^69^, respectively, and a third section captured by Spatial Mux-seq with paired RNA and ATAC data^5^ (**Figure 6a**). For Visium HD section with a pixel diameter of 2 μm, we performed 8×8 binning to obtain a 16 μm resolution for subsequent analysis. In Scenario 2, we employed a set of postnatal mouse brain data consisting of four sections profiled by three technologies: one section acquired with DBiT-ARP-seq (DBiT) for paired RNA and ATAC data^4^, two Spatial RNA-ATAC-seq (spRA) sections with paired RNA and ATAC data^2^, and one Spatial ATAC-seq (spA) section with ATAC only data^70^ (**Figure 6b**). The number of spots per section varied substantially, ranging from 2,707 to 98,917 in scenario 1 and from 2,497 to 9,787 in scenario 2 (**Supp. Fig. S14b**). UMAP visualizations of both settings revealed strong batch effects across technologies and sections (**Supp. Fig. S15a-b**).

**Fig. 6.**
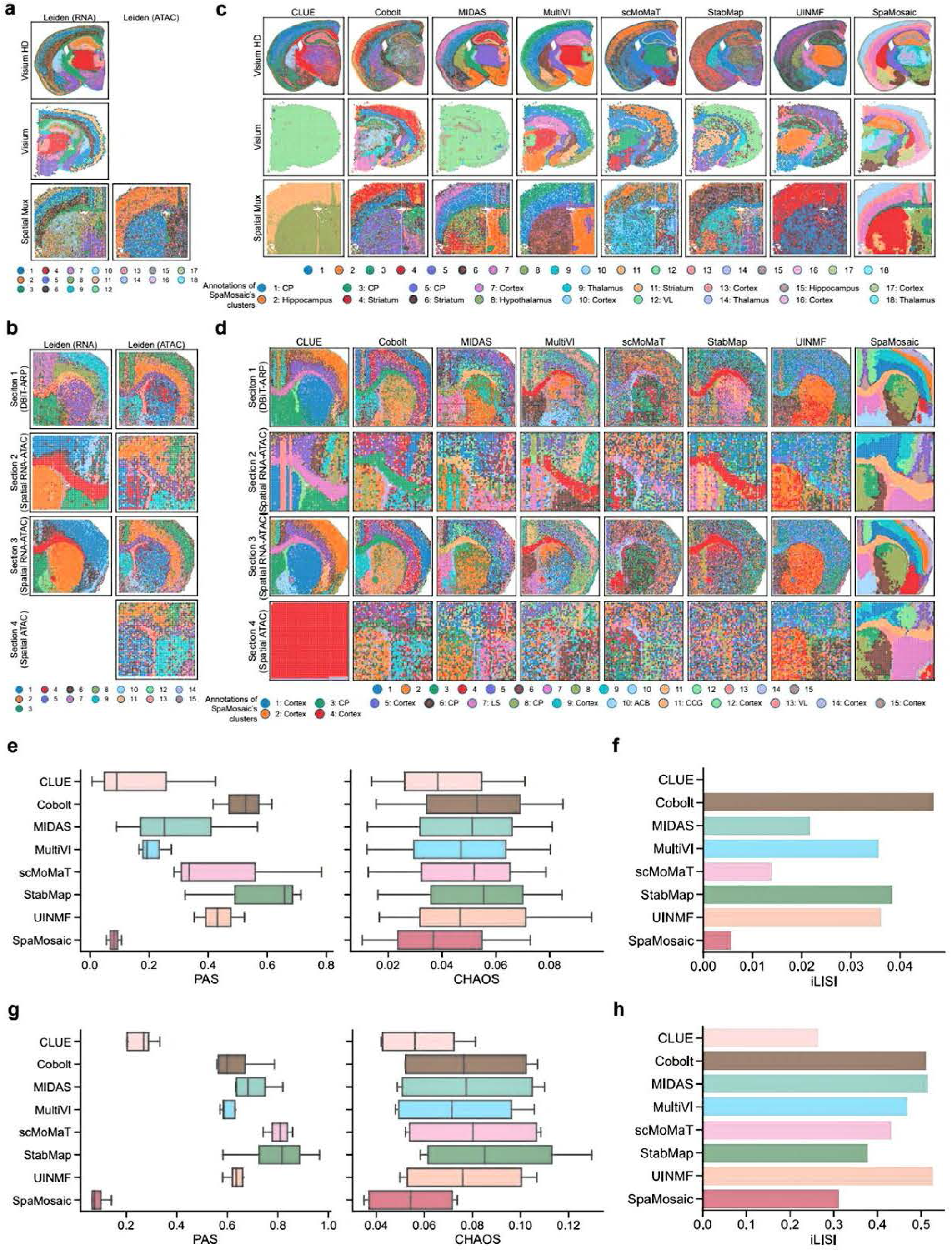
Cross-technology integration of mouse brain data across heterogeneous sizes and resolutions. **a** Mosaic dataset configuration and clustering patterns obtained for all three sections in scenario I. Modality-specific clustering was performed jointly across relevant sections using the Leiden algorithm on batch-corrected representations. **b** Spatial plots of the clusters obtained from benchmarked methods for all three sections in scenario I. Manual annotation was performed to identify SpaMosaic’s clusters. **c** Mosaic dataset configuration and clustering patterns obtained for all four sections in scenario 2. Modality-specific clustering was performed jointly across relevant sections using the Leiden algorithm on batch-corrected representations. **d** Spatial plots of the clusters obtained from benchmarked methods for all four sections in scenario 2. Manual annotation was performed to identify SpaMosaic’s clusters. **e** PAS and CHAOS scores across the three sections in scenario I. **f** iLISI scores in scenario 1. **g** PAS and CHAOS scores across the four sections in scenario 2. **h** iLISI scores in scenario 2.

In scenario 1, by comparing the clustering results with the Allen Brain Atlas reference^35^ (**Supp. Fig. S16a**), we observed that SpaMosaic accurately recapitulated key brain structures across the sections with different resolutions (16μm, 50μm and 55μm), including the cortex, hippocampus, striatum, hypothalamus, VL, and thalamus (detailed cluster assignments shown in **Figure 6c**). SpaMosaic also resolved multiple cortical layers across all three sections. To further validate these anatomical assignments, we analyzed the expression patterns of region-specific marker genes across all sections (**Supp. Fig. S17a**). Given the large difference in spot counts between sections, we analyzed the Visium and Spatial Mux-seq sections separately (**Supp. Fig. S17b-c**). Overall, the marker gene expression patterns consistently corroborated the anatomical region assignments. For the competing methods, their clustering results on the Visium HD section broadly matched the reference anatomical structures, albeit with fewer regions demarcated than SpaMosaic and with higher noise. However, these methods showed reduced performance on the lower-resolution Visium and Spatial Mux-seq sections. Specifically, for the Visium section, CLUE failed to identify anatomical structures, while MIDAS only captured the hippocampus (cluster 15). StabMap, scMoMaT, and UINMF showed improved brain structure recognition over CLUE and MIDAS, but exhibited high noise levels and lacked cortical layer differentiation. Cobolt and MultiVI separated cortical layers, though they resolved fewer layers than SpaMosaic. Both methods also identified a distinct dentate gyrus region (Cobolt: cluster 14; MultiVI: cluster 16), whereas SpaMosaic merged this region with cluster 2. However, Cobolt and MultiVI exhibited higher noise levels overall, resulting in less sharply defined spatial domains.. For the Spatial Mux-seq section, scMoMaT’s clusters showed poor correspondence with the reference structures due to substantial noise. CLUE and UINMF could only distinguish between the cortical and the non-cortical regions, and the cortical regions showed inconsistent cluster assignments across different sections. Cobolt, MIDAS, and MultiVI recovered anatomical structures comparable to those identified by SpaMosaic, albeit with high noise and reduced cortical layer differentiation. The spatial clustering method BANKSY also produced noisy spatial domains across all sections, particularly on the Spatial Mux-seq section (**Supp. Fig. S18**). It could not segregate the thalamus and hypothalamus and failed to resolve the cortical layers in the Visium HD section. CellCharter performed better by recapitulating coherent clusters on the Visium HD and Spatial Mux-seq (RNA) section that aligned with anatomical references. However, CellCharter failed to distinguish the hippocampus from surrounding regions in the Visium section and produced highly noisy clustering results on the Spatial Mux-seq ATAC modality.

In scenario 2, SpaMosaic’s clusters corresponded well to the Allen Brain Atlas reference^35^ (**Supp. Fig. S16b**), accurately capturing major brain regions including the cortex, CP, VL, and ACB across four sections (detailed cluster assignments shown in **Figure 6d**). The marker gene expression and gene activity score patterns consistently supported the anatomical region assignments (**Supp. Fig. S19a, b**). Notably, SpaMosaic resolved more detailed cortical layers in sections 1 and 3 compared to other methods. Layer-specific marker gene expression was visualized to confirm the biological identity of these clusters (**Supp. Fig. S19c**). Specifically, the deeper layer markers^71^ (*Hs3st4, Tox, Sdk2*) were predominantly expressed in layers 5 and 6 (clusters 1 and 9), while the upper cortical markers^72^ (*Rorb, Rab3c, Cux2*) showed enriched expression in layers 2/3 and 4 (clusters 2 and 14), with spatial distributions aligning well with the cluster boundaries. Among the competing methods, CLUE and Cobolt captured the CCG, CP, and several cortical layers across the first three sections. Comparatively, the other methods’ output showed limited correspondence with reference anatomical structures, with only a few identifiable regions such as the CCG being captured by MIDAS, MultiVI, and StabMap. On the more challenging ATAC-only section 4, all competing methods showed substantially noisy outputs, with CLUE failing to capture any meaningful spatial domains. For the spatial domain (niche) identification methods, BANKSY and CellCharter, identified more coherent domains than the other integration methods, but BANKSY was unable to resolve the cortical layers (**Supp. Fig. S20**). Furthermore, BANKSY and CellCharter produced inconsistent clustering of the cortical and CP regions across sections 1–3, suggesting suboptimal integration.

Finally, we applied PAS, CHAOS and iLISI for quantitative performance assessment. In scenario 1, SpaMosaic outperformed all other methods in spatial domain identification with the lowest PAS and CHAOS scores (**Fig. 6e**). In terms of batch integration, all methods achieved low iLISI scores (below 0.05) due to the substantial imbalance in spot numbers. Nevertheless, UMAP visualizations revealed that all methods except CLUE achieved partial mixing of sections, though the extreme numerical imbalance prevented uniform integration (**Supp. Fig. S15c**). In scenario 2, SpaMosaic also achieved the lowest PAS and CHAOS scores for spatial domain identification (**Fig. 6g**). Regarding batch integration, UINMF obtained the highest iLISI score, closely followed by MIDAS and Cobolt (**Fig. 6h**). The UMAP visualizations revealed a trade-off between section integration and cluster preservation: methods like MIDAS, Cobolt, and MultiVI achieved better cross-section mixing but at the cost of cluster boundaries becoming blurred (**Supp. Fig. S15d**). In contrast, SpaMosaic showed less uniform section mixing but maintained clearer cluster separation.

### Cross-platform integration of embryonic mouse brain data across six developmental stages

For this scenario, we considered a larger set of mosaic data acquired with different technologies (embryonic mouse brain dataset, Misar+Stereo). The data consisted of three mouse embryonic brain sections (E13.5, E15.5 and E18.5) with RNA and ATAC modalities acquired using Misar-seq^33^, and three additional samples (E12.5, E14.5 and E16.5) with RNA profiles obtained using Stereo-seq^36^ (**Fig. 7a**). These two spatial omics technologies differ markedly in resolution, with Stereo-seq having a center-to-center spot spacing of 0.5 μm (corresponding to an effective resolution of 25 μm after binning 50) and Misar-seq at 50 μm. Moreover, each processed Stereo-seq section contained approximately 15,000 bins, compared to about 2,000 grids per Misar-seq section (**Fig. 7b**). Using UMAP visualization, we could observe strong batch effects across technologies and time points, which posed a major challenge for integration (**Supp. Fig. S21a**). Alongside SpaMosaic, we also tested CLUE, Cobolt, MIDAS, StabMap, scMoMaT, MultiVI and UINMF.

**Fig. 7.**
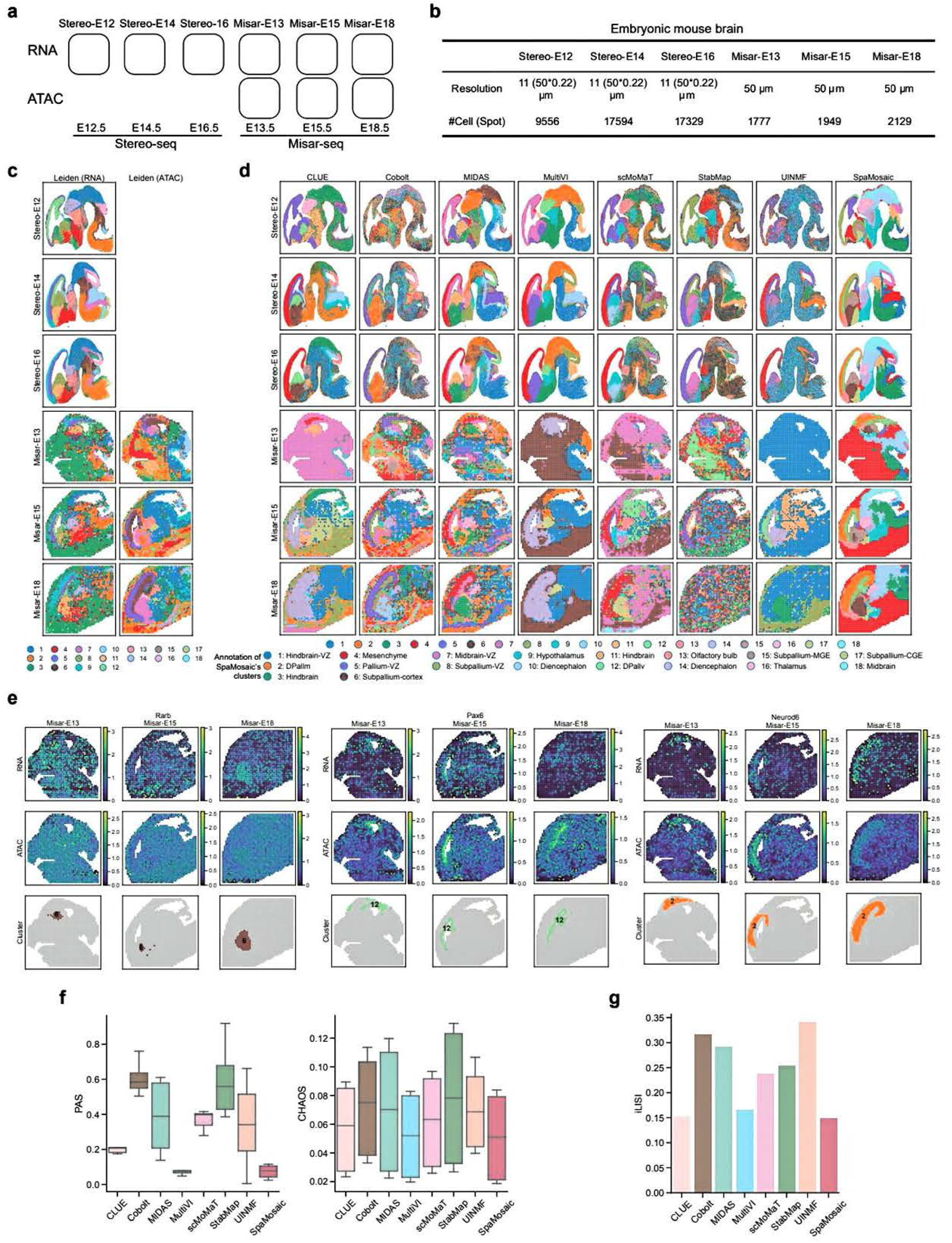
Downstream analysis results obtained with SpaMosaic and quantitative benchmarking on an embryonic mouse brain (Misar+Stereo) dataset. **a** Schematic of the dataset composition. Mouse embryonic brain sections at developmental stages El3.5, El5.5, and El8.5 were profiled for the RNA and ATAC modalities using Misar-seq, and sections at El2.5, El4.5, and El6.5 were profiled for RNA expression using Stereo-seq. **b** Comparison of resolution and cell numbers between Stereo-seq (**11**µm; 15,000 cells per section) and Misar-seq (50 µm; 2,000 cells per section). **c** Spatial plots of modality-specific clustering results obtained by applying the Leiden algorithm to batch-corrected representations, jointly across relevant sections. **d** Spatial plots of the clusters obtained from mosaic integration methods for the six sections. Manual annotation was performed to identify SpaMosaic’s clusters. **e** RNA expression and ATAC-derived gene activity maps for *Rarb, Pax6*, and *Neurod6* in the three Misar-seq sections. **f** PAS and CHAOS scores of benchmarked methods across the six sections. **g** iLISI scores of benchmarked methods.

Among all test methods, SpaMosaic produced coherent spatial domains with markedly higher consistency in inter-section cluster assignment (**Fig. 7c-d**). With reference to the Allen Brain Atlas^35^ (**Supp. Fig. S22a**), we could identify the major brain regions, including the dorsal pallium (DPall), olfactory bulb (OB), subpallium, hypothalamus, diencephalon, midbrain and hindbrain (detailed cluster assignments shown in **Fig. 7d**). Expression of relevant brain region markers was visualized to confirm the identities of SpaMosaic’s clusters (**Supp. Fig. S22b**). For example, SpaMosaic subdivided the dorsal pallium into two distinct clusters (DPallm and DPallv), with DPallm (cluster 2) exhibiting higher expression of *Mef2c*^*33*^ and *Tbr1*^*73*^. In the subpallium, three transcriptionally distinct clusters were identified: cluster 6 (subpallium-cortex) was marked by *Foxp1*, a cortical excitatory neuron marker^44^; cluster 15 (subpallium-medial ganglionic eminence; MGE) which was enriched for *Dlx1*, a key transcription factor for GABAergic interneuron development^74^; and cluster 17 (subpallium-caudal ganglionic eminence; CGE) that was characterized by *Lhx8* and *Adarb2*, markers of CGE-derived inhibitory interneurons^75^. Cluster 16 (thalamus) was marked by transcription factors *Tcf7l2* and *Lhx9*, which are essential for thalamic and excitatory neuronal development^76^. Within the diencephalon, SpaMosaic resolved two clusters: cluster 10 which was enriched for *Ebf3* and *Gata3*, associated with neuronal differentiation and cholinergic/inhibitory lineages^77,78^; and cluster 14 that was marked by *Dlx1* and *Arx*, characteristic of GABAergic interneurons^74^. In the hindbrain, two clusters were resolved: cluster 3 that expressed *Meis1* and *Mab21l2*, genes involved in neuronal differentiation and patterning^79^; and cluster 11 that was enriched for *Pax5* and *En1*, regulators of the midbrain–hindbrain boundary and progenitor maintenance^80^. Beyond these major brain regions, SpaMosaic also resolved the ventricular zones (VZ) into transcriptionally distinct clusters: pallium-VZ (cluster 5), midbrain-VZ (cluster 7), subpallium-VZ (cluster 8) and hindbrain-VZ (cluster 1), each identified by region-specific marker gene expression. We next analyzed the Misar-seq samples to demonstrate how SpaMosaic integrates multimodal information. We visualized the expression of *Rarb, Pax6* and *Neurod6*, which serve as marker genes for subpallium-cortex, DPallv, and DPallm clusters, respectively^33,81^ (**Fig. 7e**). These genes displayed distinct expression patterns across the RNA and ATAC modalities: *Rarb* showed enhanced RNA expression in the subpallium, while its ATAC-derived gene activity was uniformly distributed without evident spatial specificity. Conversely, *Pax6* and *Neurod6* showed more pronounced region-specific activity in ATAC data compared to lower specificity in their RNA expression. Together, these results illustrate how SpaMosaic leverages complementary RNA and ATAC signals to achieve integrative spatial characterization.

By contrast, most competing methods produced noisy clustering outputs that poorly matched known anatomical structures, particularly for the Misar-seq sections (**Fig. 7d)**. For example, CLUE merged the distinct hindbrain and midbrain regions on Stereo-seq (E12.5) and Misar-seq (E13.5) sections and failed to distinguish DPallm from DPallv regions within most sections. MIDAS successfully identified major brain regions (DPallm, DPallv, hindbrain, midbrain, subpallium, etc.) in the Stereo-seq sections, but for the Misar-seq sections, it delineated only DPallm and DPallv while the remaining clusters appeared dispersed. MultiVI produced coherent spatial domains across the six sections and successfully captured major brain regions on the Stereo-seq sections, such as the hindbrain, midbrain and subpallium. However, MultiVI recovered only a few structures in the Misar-seq sections and these cluster assignments were inconsistent with those in the Stereo-seq sections, indicating suboptimal integration between the two platforms. For the remaining methods, Cobolt, StabMap, scMoMaT and UINMF, their clusters were noisy and few could be matched to the reference anatomical structures. Finally, we examined the clustering results from BANKSY and CellCharter. Despite being tailored for spatial transcriptomics clustering, BANKSY produced noisy clusters and failed to achieve consistent cluster assignments between the two groups of sections acquired with different technologies (**Supp. Fig. S23**). CellCharter generated more spatially coherent clusters but also failed to provide consistent cluster labels between Stereo-seq and Misar-seq sections in the RNA modality.

We next quantitatively compared SpaMosaic to the other mosaic integration methods on two aspects: spatial domain identification (PAS and CHAOS) and batch integration (iLISI). In terms of spatial domain identification, MultiVI and SpaMosaic achieved similar PAS and CHAOS scores that consistently outperformed other methods (**Fig. 7f**). However, our earlier analysis demonstrated that SpaMosaic accurately captured more anatomical brain structures than MultiVI. For batch mixing, UINMF achieved the highest iLISI score, followed by Cobolt (**Fig. 7g**). The UMAP visualizations showed that while SpaMosaic had a lower iLISI score, it maintained clear cluster boundaries with reasonable section mixing, whereas Cobolt produced more diffused clusters with overlap (**Supp. Fig. S21b**).

### Comprehensive integration of mouse embryo data

In this final example, we applied SpaMosaic and competing methods to a comprehensive integration scenario of seven mouse embryo sections, which differed in developmental stages, sequencing technologies and anatomical regions. This example used tissue sections from E11 to E13 embryos, profiled by four distinct spatial multi-omics techniques: Spatial RNA-ATAC-seq^2^ (section 1, E13), Spatial Mux-seq^5^ (sections 2 and 3, E13), Spatial ATAC-seq^70^ (section 4, E13), and Spatial Cut&Tag-seq^1^ (sections 5 to 7, with section 5 from E13 and sections 6 and 7 from E11). Each section contained a unique combination of molecular modalities: section 1 contained RNA and ATAC, section 2 had H3K27ac and H3K27me3, section 3 encompassed RNA, ATAC, H3K4me3, and H3K27ac, while sections 4 to 7 consisted of single-modality ATAC and histone modification assays without accompanying RNA profiles (**Fig. 8a**). All sections shared the same spatial resolution, captured on a 50×50 array with 50□µm pixel diameters (**Fig. 8b, c**). UMAP visualizations of each modality revealed pronounced batch effects between sections (**Supp. Fig. S24a**). Several existing mosaic integration methods, Cobolt, MIDAS, MultiVI, and UINMF, were unable to process such a complex scenario, and hence we only tested SpaMosaic, StabMap, scMoMaT and CLUE.

**Fig. 8.**
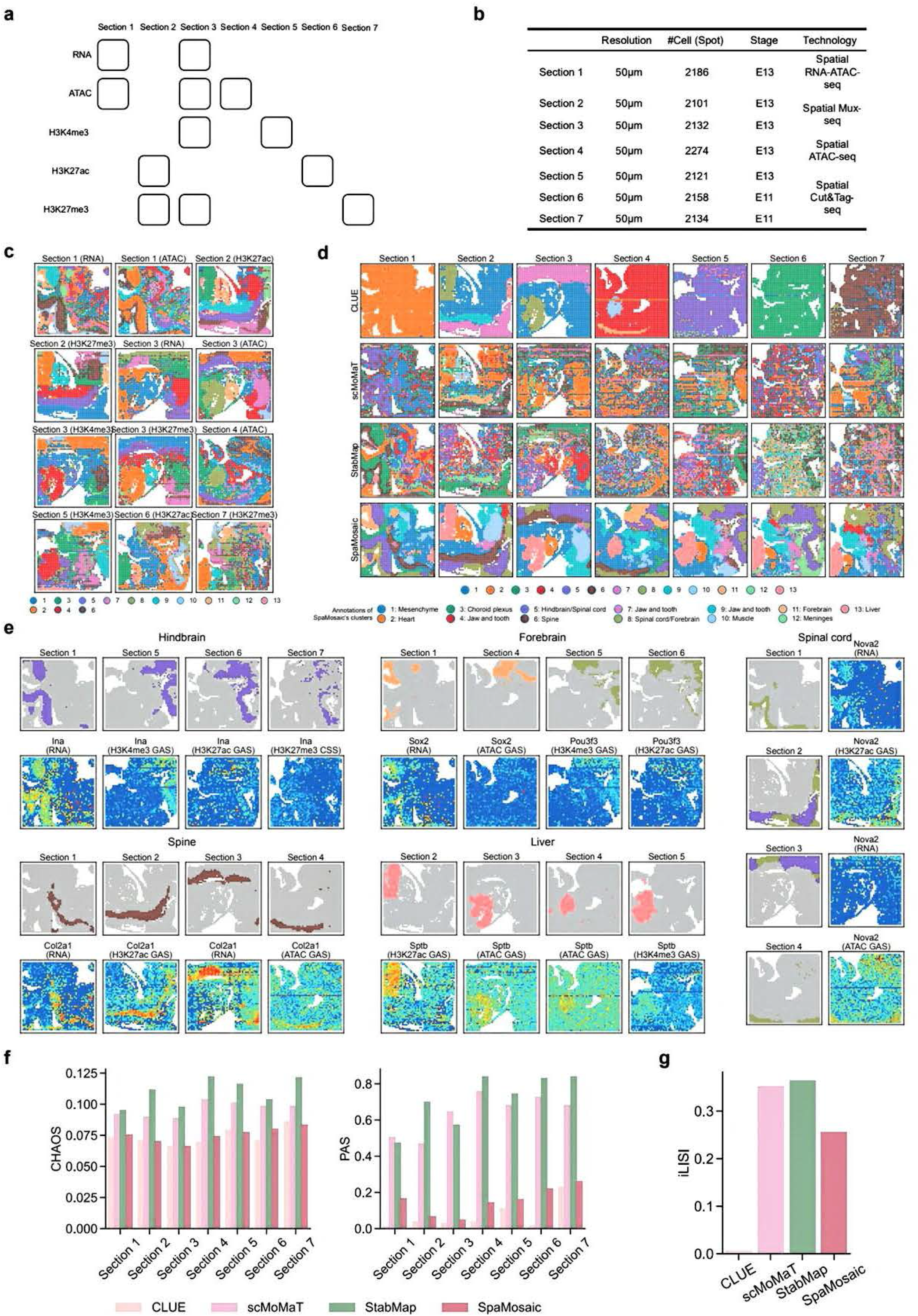
Application of SpaMosaic to mouse embryo data integration. **a** Layout of input data matrices in mouse embryo dataset. **b** Metadata for each section in the mouse embryo dataset. **c** Modality-specific clustering was performed jointly across relevant sections using the Leiden algorithm on batch-corrected representations. **d** Spatial plots of clusters obtained from the methods for all seven sections. Manual annotation was performed to identify SpaMosaic’s clusters. **e** Spatial heatmap of RNA expression, gene activity scores (GAS), and chromatins silence scores (CSS) for region-specific markers in SpaMosaic’s clusters. **f** PAS and CHAOS scores of benchmarked methods across seven sections. **g** iLISI scores of benchmarked methods.

Among the tested mosaic integration approaches, SpaMosaic demonstrated superior ability in identifying consistent spatial domains across all seven sections, despite differences in the sections’ orientation and modality composition (**Fig. 8d**). We visualized the respective marker gene expression to annotate and confirm that SpaMosaic successfully recovered key tissues and organs (**Fig. 8e**). For instance, the hindbrain marker *Ina*^*36*^ was highly expressed in the RNA modality (cluster 5, section 1), and showed elevated H3K4me3 and H3K27ac GAS (cluster 5, sections 5 and 6) and reduced H3K27me3 CSS (cluster 5, section 7). The forebrain markers^36^ *Sox2* and *Pou3f3* showed high RNA expression (cluster 11, section 1) and increased ATAC or histone modification GAS (cluster 11 in section 4, cluster 8 in sections 5 and 6). The spine marker *Col2a1*^*2*^ had high RNA expression (cluster 6, sections 1 and 3) and exhibited elevated ATAC and H3K27ac GAS (cluster 6, sections 2 and 4). The liver marker *Sptb*^*70*^ had consistently high H3K27ac, ATAC, and H3K4me3 GAS (cluster 13, sections 2–5). The spinal cord marker *Nova2*^*82*^ was enriched in the RNA modality (clusters 8 and 5, sections 1 and 3) and showed increased H3K27ac and ATAC GAS (clusters 8 and 5, sections 2 and 4). We also visualized the markers for other anatomical features, namely the heart, jaw and teeth (mandibule), mesenchyme, meninges, muscle and choroid plexus, which displayed high RNA expression or gene activity scores in their corresponding clusters (**Supp. Fig. S25**).

Among the competing methods, CLUE was unsuccessful at identifying any spatial domains in sections 1, 5, 6 and 7. It captured several anatomically relevant clusters in sections 2, 3 and 4, but fewer than the expected anatomical features. For scMoMaT and StabMap, their clusters were spatially discontinuous and only a few could be identified as relevant anatomical structures. For the single-modality spatial clustering algorithms, BANKSY and CellCharter, more coherent and anatomically relevant clusters were identified (**Supp. Fig. S26a**). For example, BANKSY detected the hindbrain (cluster 2) and forebrain (cluster 4) in section 1 and the spine (cluster 3), but did not capture the heart or liver in section 3. CellCharter (RNA) delineated the forebrain (clusters 5 and 13), hindbrain (cluster 3) and spinal cord (cluster 7) in section 1, but detected only four clusters in section 3 while missing major anatomical domains such as the heart and liver. For the other modalities, CellCharter showed inconsistent cluster assignments for the same structures across sections. The UMAP plots further highlighted that both CellCharter and BANKSY did not mix the sections effectively (**Supp. Fig. S26b**).

As our final assessment component, we quantitatively benchmarked performance in spatial domain identification (CHAOS and PAS) and batch integration (iLISI). In terms of CHAOS scores, CLUE and SpaMosaic were close competitors as the top two methods; SpaMosaic was top on sections 2, 3, 5, and 7, while CLUE was top on the rest (**Fig. 8f**). For the PAS metric, CLUE was top and SpaMosaic second for all sections. However, we note here that while CLUE’s spatial domains scored highly in the spatial coherence metrics, they showed limited ability in capturing embryonic anatomical structures. In terms of batch integration, StabMap attained the highest iLISI score with scMoMaT coming in as a close second and SpaMosaic in the third place (**Fig. 8g**). In contrast to its strong performance on CHAOS and PAS, CLUE showed poor integration performance with a score close to zero. These integration results were consistent with the UMAP visualizations (**Supp. Fig. S27**).

## Discussion

Spatial omics technologies are powerful techniques for interrogating tissue sections. By measuring different omes, we can obtain complementary views of the tissue. Coupled with the spatial information, such data can enable greater insights into the underlying biology. However, technological limitations and high costs pose challenges to acquiring multiple high quality omics profiles from a single section. Therefore, an alternative approach is to integrate data acquired of different omes from different sections. In this work, we presented SpaMosaic for such spatial mosaic integration. It leverages a contrastive learning framework to construct modality-agnostic latent spaces and employs graph neural networks to encode spatial proximity and omics feature similarity into the embeddings, achieving spatially aware integration. The contrastive learning framework bridges the information gap between modalities, which has been validated as a simple but effective strategy for modality alignment and semantic preservation in various domains^83-86^. Moreover, it is also highly flexible and allows easy scaling to include more data modalities. As shown in our last example, SpaMosaic can perform five-modal mosaic integration.

SpaMosaic’s latent embedding output enables accurate and robust consensus spatial clustering, outperforming other methods in our benchmarking with different tissue types and different technology platforms. By integrating information from different sections and different omics modalities, SpaMosaic reduces noise in the sections to enhance spatial clustering results. SpaMosaic is not restricted to integrating data from adjacent sections; we demonstrated that it could integrate data from different time points, different studies, or even different technology platforms. Alongside cross-modality integration, it simultaneously achieves cross-batch integration to remove technical effects while preserving biological variations. This enables the identification of consensus cell types and tissue structures across samples, as well as the capture of subtle differences between sample groups. In addition to data integration, SpaMosaic can impute the missing molecular layers within a mosaic dataset. To accomplish imputation, SpaMosaic employs a *k*NN-based approach to estimate the measurements in the missing modalities, avoiding the direct use of predictions from neural networks and thereby enhancing its interpretability.

Due to the high cost and difficulty of simultaneously profiling multiple omics from the same tissue section, most existing data have been generated with different single-omics modalities from separate sections. In this context, diagonal integration is necessary, and image registration is often required to align the sections prior to data integration with most currently available integration algorithms. However, aligning sections can be challenging and even impossible in cases where the sections are highly distinct. We propose a possible workaround here with the addition of relevant available spatial multi-omics data as bridges between the different omics modalities, thereby converting into a mosaic integration problem. With SpaMosaic capable of alignment-free mosaic integration, it can accomplish the integration as it does not require the sections to be adjacent. Such data integration will enable us to construct spatial multi-omics atlases from diverse datasets with varying modality compositions. Furthermore, SpaMosaic’s cross-modality integration and imputation also enable us to dissect the relationships between omes that cannot be co-profiled, potentially revealing new connections between molecular layers that would otherwise remain undetectable.

Future developments of SpaMosaic include enhancing its scaling with large datasets and avoiding model retraining with new data. By enhancing SpaMosaic’ scaling, it can handle larger datasets with more tissue sections, which is critical for constructing spatial multi-omics atlases. As graph neural networks are more computationally expensive than traditional multiple layer perceptron (MLP) networks, we propose to adopt techniques such as graph sampling and caching of intermediate outputs to improve performance^87^. We also plan to investigate continual learning^88^ and transfer learning^89^ approaches to enable SpaMosaic to analyze new sections without the need to retrain on the entire dataset. Another benefit is the offline analysis of private datasets with a relevant pretrained SpaMosaic model, thus addressing potential data privacy issues. In the future, we will also incorporate accompanying image data such as staining images as an additional modality to enhance the accuracy of cell type and tissue domain identification.

## Methods

### Data Preprocessing

For each spot (cell) *i* ∈ {1,…,*N*_*b*_} from section id *b* ∈ {1,…,*B*}, let 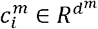 of size *d*^*m*^ be the omics feature count vector of spot *i* from modality *m* ∈ *U* ={*m*_1_,…,*m*_*M*_} where *U* is the set of all RNA, epigenomic and protein modalities. For each modality, we first normalized the count matrix using log-normalization for RNA, TF-IDF transformation for epigenomic modality, and CLR normalization for protein. Thereafter, we selected the top 5,000 highly variable features for RNA and top 100,000 for epigenome, followed by SVD decomposition into low-dimensional representations. To correct batch effects within a modality, we used Harmony or Seurat to integrate these low-dimensional representations. For spot *i* with modality *m*, the batch-corrected representation 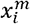, which was then used as input for model training.

### Construction of the spot-spot graph

Here we propose to encode both spatial and omics feature similarity of spots in each modality with a spot-spot graph. Each modality specific graph contains two types of edges: spatial-based edges and omics feature-based edges. Spatial-based edges connect spots within the same section while omics feature-based edges connect spots across different sections. For spatial-based edges, we first compute the Euclidean distance between spots of the same section based on their spatial coordinates. We then identify each spot’s *k* nearest neighbors (*k*NN) using the Annoy python package (https://github.com/spotify/annoy) for efficient (*k*NN) searching. The neighbors with the shortest distance are considered spatial neighbors. Finally, we construct an undirected spatial neighbor graph of all spots in modality *m* (all modalities share the same spatial neighbor graph) in the form of an adjacency matrix *A*^*intra-m*^, where 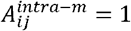 if spots *i* and *j* are spatial neighbors.

For the omics feature-based edges, we use the batch-corrected low-dimensional representation to find the mutual nearest neighbors (MNNs)^22^ between sections. Specifically, for each spot *i* in section *b*_1_, we find its *k*_1_ nearest neighbors in section *b*_2_; for each spot *j* in *b*_2_, we find its *k*_2_nearest neighbors in *b*_1_. If spots *i* and *j* are in each other’s nearest neighbors set, then (*i,j*) form a pair of MNNs. We then construct a neighbor graph for each modality *m* denoted by an adjacency matrix *A*^*inter-m*^, where 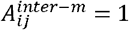 if *i* and *j* belong to a pair of MNNs. For each modality *m*, we now combine the *A*^*intra-m*^ and *A*^*inter-m*^ into one graph:

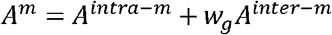

where *w*_*g*_ is a positive constant, denoting the weight for *A*^*inter-m*^.

We found that when a pair of sections have highly imbalanced spot counts, using equal neighbor counts (*k*_1_= *k*_2_) tends to bias the MNN set. Specifically, most spots from the smaller section are selected, while only a small fraction of spots from the larger section are used. This is analogous to scaling up the small section to match the large one, even though the small section may only occupy a fraction of the large section’s area. To mitigate this, we adopt asymmetric neighbor sizes when section sizes are substantially imbalanced. For a base neighbor count *k*_0_, if the imbalance exceeds a threshold *θ* (i.e., *n*_1_/*n*_2_< *θ*, where *n*_1_ and *n*_2_ represent the spot counts of the two sections), the smaller section uses *k*_0_, and the larger section uses max (1, ⌊*k*_0_ ·(n_1_/n_2_) ⌋). Otherwise, both sections use *k*_0_. To further prune potential outliers, we apply an isolation forest^90^ to remove 50% of MNN pairs with the highest anomaly scores. The input features to the isolation forest model consist of spatial offsets (absolute coordinate differences between paired spots) concatenated with their coordinates. Both the asymmetric neighbor sizing and outlier removal steps are configurable options within our pipeline.

### Graph embedding

The spot-spot graph encodes information about the spatial proximity and omics feature similarity between spots which we integrate with the omics profiles into spot embeddings via graph neural networks. However, our spot-spot graph contains two types of edges, each with different semantics. This differs from the traditional homogeneous graphs and thus we explored various approaches to address this situation, namely the graph attention network (GAT)^23,24^, heterogeneous graph transformer (HGT)^25^ and two variants of light graph convolution network (LGCN)^26,91^ (**Supplementary Information**). In our final model, we adopt the weighted LGCN, which is defined as:

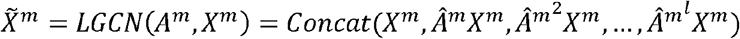

where *m* denotes the modality and *l* denotes the number of layers. 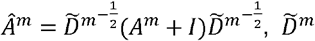 is the diagonal degree matrix of *A*^*m*^+ *I*. The embeddings are then transformed by a multiple layer perceptron (MLP):

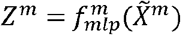

where *m* denotes the modality. Finally, the graph embedding function is defined as *g*:

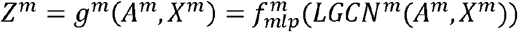

By setting *w*_*g*_, we make *A*^*m*^ a weighted graph to control the importance between spatial proximity and omics feature similarity. From our testing, we decided on *w*_*g*_ = 0.8 for our benchmarking examples, giving more weight to spatial proximity than omics feature similarity, as we consider the former to be more reliable due to measurement noise present in the omics profiles.

### Contrastive learning

Within SpaMosaic, the model’s goal is to learn a set of modality-agnostic low-dimensional embeddings, thereby aligning multiple sections with different modalities. SpaMosaic employes contrastive learning to leverage the input data’s multi-modality nature to achieve this goal. In the contrastive learning framework, two views of one sample constitute a positive pair while views from different samples constitute negative pairs^19^. By pushing positive pairs together while separating negative pairs in the latent space, contrastive learning achieves alignment between the different views and preserve variation between samples. For SpaMosaic, two modalities of one spot constitute a positive pair and modalities from different spots constitute negative pairs. Thus, for two modalities *m*_1_ and *m*_2_, there is a sampled mini-batch of spots 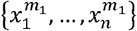 and 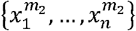 that form positive pairs and the associated contrastive learning objective is:

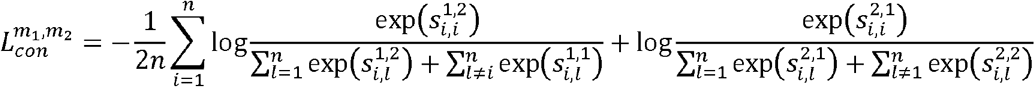

where

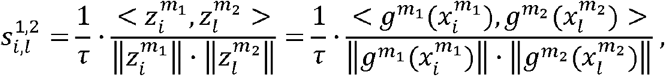

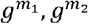 denote the modality-specific graph encoder network, 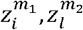denote the modality-specific embeddings, <, > denotes inner product of two vectors and *τ* is a positive constant. The numerator in the contrastive learning objective aims to bring closer the embeddings of each spot’s two modalities, while the first term in the denominator aims to separate the embeddings of two modalities from different spots and the second term aims to separate the embeddings within the same modality but from different spots.

Combining embeddings from the measured modalities, the unique representation for spot *l* in section *b* is defined as:

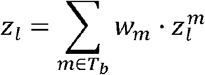

where *T*_*b*_ denotes the set of modalities that section *b* is measured with, *w*_*m*_ ∈ (0,1) denotes the weight for modality *m* and 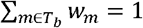.

### Feature reconstruction

Based on our definition, contrastive learning can only be applied to spots with multimodal profiles. For the remaining spots without multimodal profiles, we add a feature reconstruction objective to include them in the model training. We employ a modality-specific MLP decoder, *r*^*m*^, to reconstruct the inputs 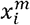 based on the embeddings 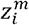. The reconstruction loss is defined as:

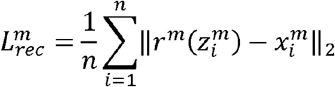

Thus, final loss function for the whole model is defined as:

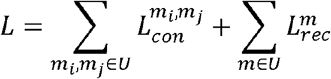

where *U*={*m*_1_,…,*m*_*M*_}.

### Implementation details

For each modality, the graph encoder network comprises two weighted-LGCN layers, followed by a fully connected layer with a dimension of 128, leaky ReLU activation (slope=0.2)^92^, batch normalization^93^, dropout(*p* = 0.2)^94^, and a final fully connected layer with a dimension of 32. The decoder comprises one fully connected layer with linear activation. In our implementation, the graph neural networks uses the PyTorch geometric library^95^. To train the model, we used the Adam optimizer^96^ with a learning rate of 0.01. The weight decay was set to 5e-4, the number of training epochs was set to 100 and the temperature parameter *τ* was set to 0.01. The parameter *k* used to calculate the spatial adjacency graph was set to 10 across all experiments. For MNN calculation, we set *k*_0_ = 10 across all experiments. When asymmetric neighbor sizing was applied, the imbalance threshold was set to θ = 0.8. For outlier removal with isolation forest, we used scikit-learn’s default parameters^97^. Our implementation supports both GPU and CPU execution, with GPU acceleration recommended for improved computational efficiency.

### Imputation

Let 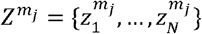 denote the graph embeddings for all spots that were measured in modality *m*_*j*_. Alongside, 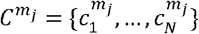 denotes the feature profiles (counts, gene activity scores, etc.) for all spots that were measured with modality *m*_*j*_. Let *T*_*b*_ denote the set of modalities that section *b* was measured with and 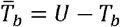 denotes the set of modalities that are missing in section *b*. To impute the missing modality 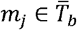 for *b*, we first perform cross-modal matching between 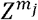 and 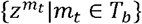. Specifically, for each 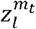 in section *b*, we find the indices of its *k*_2_ nearest neighbors from, 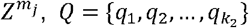. The profiles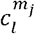 imputed from modality *m*_*t*_ are thus defined as:

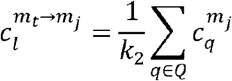

where *m*_*t*_ → *m*_*j*_ denotes the profiles imputed based on modality *m*_*j*_. For each modality in *T*_*b*_, we repeat the above imputation process and the final imputed profiles for *m*_*j*_ in section *b* are given by:

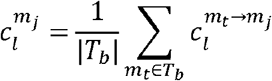

where |*T*_*b*_ | denotes the number of elements in *T*_*b*_.

### Data simulation

Following the procedure of Townes et al.^28^, we simulated multivariate count data with the ‘ggblocks’ spatial pattern. The ground true spatial factors comprised four simple shapes in different spatial regions. For each section created, the spots were arranged in a 36×36 grid for 1296 spots. The number of RNA features was set to *J*^*rna*^= 200, and the number of protein features was set to *J*^*pro*^=60. Each feature was randomly assigned to one of the four factors with uniform probability. For the RNA modality, its count matrix was drawn from a zero-inflated negative binomial (ZINB) distribution with a mean *M*^*rna*^= *b*^*rna*^ + *S*^*rna*^ *F*^*rna*^ *W*^*rna*^ where *S*^*rna*^ ∈ *R*^1296×1296^, *F*^*RNA*^ ∈ *R*^1296×4^and *W*^*rna*^ ∈ *R*^4×200^. 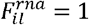 if spatial factor was active in spatial location *i* and 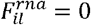 otherwise. By default, 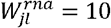 if feature *j* was assigned to spatial factor *l* and 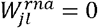 otherwise. *b*^*rna*^ represents the background expression mean value. To simulate a spatial factor *l* mixing with the background factor, we multiplied *F*^*rna*^ by a diagonal matrix *S*^*rna*^. For those entries corresponding to factor 1 we set them to *s*^*rna*^, while for factors 2 and 3, we randomly selected 40% entries to *s*^*rna*^ and the other entries were set to 1. The shape parameter for the negative binomial component was set to 20 and the dropout probability was set to *p*^*rna*^. For the protein modality, its count matrix was drawn from a negative binomial distribution with a mean *M*^*pro*^= *b*^*pro*^ + *S*^*pro*^ *F*^*pro*^ *W*^*pro*^, where *S*^*pro*^ ∈ *R*^1296×1296^, *F*^*pro*^ ∈ *R*^1296×4^and *W*^*pro*^ ∈ *R*^4×60^. 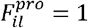 if spatial factor *l* was active in location *i* and 0 otherwise. By default, 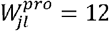 if feature *j* was assigned to pattern *l* and 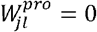 otherwise. *S*^*pro*^ is a diagonal matrix and for those entries corresponding to factor 4 we set them to *S*^*pro*^. For factors 2 and 3, we randomly selected 40% entries to be assigned *S*^*pro*^, and the other entries were set to 1. The shape parameter was set to 10.

To simulate batch effects between sections, we changed the *W*^*rna*^ and *W*^*pro*^ for the second and the third section, respectively. Specifically, if the RNA feature set 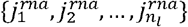 was assigned to spatial factor *l*, we randomly picked half of these features and increased each 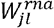 by a factor of 3. Similarly, if protein feature set 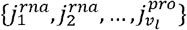 was assigned to spatial factor *l*, we randomly picked half of these features and increased each corresponding 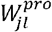 by a factor of 1.1, while for the remaining features, we increased each 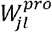 by a factor 3.

We designed five sets of simulation parameters (**Supp. Fig. S2a**) and we created three datasets with different random seeds for each parameter set. Each dataset consisted of three sections sharing the same simulation parameters. The first section contained RNA and protein expression, the second section only contained RNA expression, and the third section contained only protein expression (**Supp. Fig. S2b**).

### Benchmarking metrics for integration

The benchmarking employed in this work encompassed three aspects: spatial domain identification, batch correction and modality alignment. Spatial domain identification was assessed using the adjusted rand index (ARI), PAS and CHAOS^29^, batch correction was assessed using the graph integration local inverse Simpson’s index (iLISI) ^30^, and modality alignment was assessed using Fraction of Samples Closer than True Match (FOSCTTM)^42^, matching score (MS)^43^ and label transfer F1 score^11^.

*CHAOS* CHAOS evaluates the spatial continuity of spatial clusters. To calculate CHAOS, we first construct a 1-nearest neighbor (1-NN) graph for each cluster based on spatial coordinates. Specifically, for each spot *i* in cluster *k*, we identify the nearest spot within the same cluster *k* based on Euclidean distance, obtaining the distance *d*_*ki*_. We then compute the average of these minimum distances across all spots:

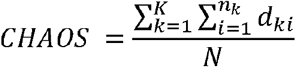

where *K* represents the number of clusters and *n*_*k*_ represents the number of spots within cluster *k*. A lower CHAOS score indicates better continuity of the spatial domains.

*PAS* PAS is also used to evaluate the spatial continuity of output from spatial domain identification methods. In brief, the PAS score is calculated as the percentage of spots whose cluster label differs from that of at least six of their ten spatial neighbors.

*Graph iLISI* The original iLISI^20^ was initially conceived to assess the degree of local batch mixing in single-cell data, and Graph iLISI is an extension for assessing graph-based outputs. Graph iLISI uses a graph-based distance metric to determine the nearest neighbor list and avoids skews in graph-based integration outputs^11^. Its scaled output ranges from 0 to 1, where 0 indicates strong separation of batches while 1 indicates perfect batch mixing.

*FOSCTTM* FOSCTTM quantifies the degree of alignment between embeddings of different modalities for the same location. For each modality pair(*m*_1_,*m*_2_), FOSCTTM is defined as:

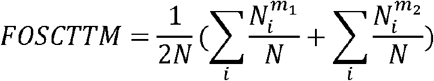

where *N* is the number of spots that are simultaneously measured in modality *m*_1_ and *m*_2_, 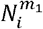 denotes the number of spots in modality *m*_1_ that are closer to 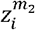 than 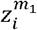 to 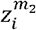, and 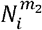 is the number of spots in modality *m*_2_ that are closer to 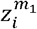 than 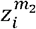 to 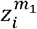. The computed value ranges from 0 to 1, where 1 indicates perfect modality matching.

*MS* Like FOSCTTM, MS quantifies the degree of modality alignment and is calculated using spots with multimodal measurements. For two modalities, *m*_1_ and *m*_2_, a cross-modality matching matrix *P* is first constructed by computing the Jaccard index of cross-modality nearest neighbors in the modal-aligned embedding space^27^:

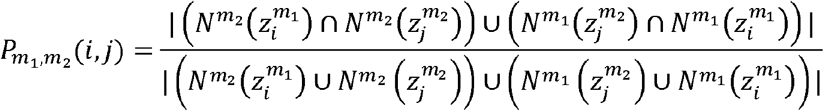

where 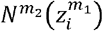 denotes the set of spot 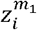 ‘s nearest neighbors in modality *m*_2_, 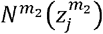 denotes the set of spot 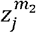 ’s nearest neighbors in modality *m*_2_ (including itself), and vice versa for 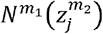 and 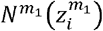. Following the approach of CLUE, the number of nearest neighbors is set to 1 (i.e., *p*(*i,j*)=1 if spot *i* and *j* are mutual nearest neighbors). Finally, MS is computed by:

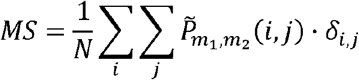

where 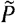denotes the row-normalized *P, δ*_*i,j*_=1 if spot *i* and *j* represent the same spot and 0 otherwise, and *N* is the number of spots. The computed MS ranges between 0 and 1 where 1 indicates perfect alignment.

*Label transfer F1 score* This metric assesses the alignment of labeled cell types or spatial domains between sections and is not restricted to spots with multimodal measurements. Therefore, it can test for the transfer accuracy of cell type or spatial domain labels from one section to another. For each pair of modalities, we first built a *k*NN graph between their embeddings and the transferred labels from one to another based on the *k*NN classifier (*k*=10). Thereafter, we calculated the F1 score by comparing the transferred labels with the ground truth. For each pair of modalities, we then averaged the F1 scores from both transfer directions.

### Benchmarking methods for mosaic integration

In our benchmarking, we tested SpaMosaic and other state-of-the-art single-cell mosaic integration methods, CLUE, Cobolt, scMoMaT, StabMap, MIDAS, UINMF, and MultiVI. We also tested a non-spatial version of SpaMosaic. Here we describe the detailed settings used for each method.

#### CLUE

We employed CLUE from the scglue package (version 0.1.1)^98^ in Python. We followed the processing pipeline as described by the authors (https://github.com/openproblems-bio/neurips2021_multimodal_topmethods/tree/main/src/match_modality/methods/clue). Raw count matrices of each modality were used as input, which were then transformed by the customized preprocessing steps in CLUE. Thereafter, we applied the default model training pipeline. We set the *shared_batches = False* and used the recommended training parameters for different types of input data, namely RNA+ATAC and RNA+protein.

#### Cobolt

Although originally designed for RNA and ATAC-seq integration, Cobolt’s statistical framework is generalizable to other omics modalities through its multinomial modeling approach^9^. We ran the Cobolt package (version 1.0.1) following its tutorial (https://github.com/epurdom/cobolt/blob/master/docs/tutorial.ipynb). We first filtered for the highly variable features (top 5,000 for RNA, top 100,000 for epigenome) from the count data matrices before input to the model for training. Most of training hyperparameters were set to default values: training iteration=100, latent dimension=10, batch size=128. However, we modified the learning rate from 0.005 to 0.001 to avoid the numerical issues encountered with our datasets. When running the ‘calc_all_latent’ function for the postnatal mouse brain dataset (spRA+spRH), we set the parameters *target = [True,False,False*,*False*,*False]* for successful execution.

*scMoMaT* We employed the Python package scMoMaT (version 0.2.2) by following its tutorial (https://github.com/PeterZZQ/scMoMaT/blob/main/demo_scmomat.ipynb). We first selected the highly variable features (top 5,000 for RNA, top 100,000 for epigenome) in the count data matrices as input for model training. The model parameters were kept at their defaults except that we disabled the generation of pseudo-count RNA matrix from ATAC data for better performance with our examples.

*StabMap* We employed the StabMap package (version 1.3.0) in R, following its tutorial (https://www.bioconductor.org/packages/devel/bioc/vignettes/StabMap/inst/doc/stabMap_PBMC_Multiome.html). We first performed data normalization using ‘logNormCounts function for all modalities. Thereafter, we selected the highly variable features using ‘modelGeneVar’ function with modality-specific filtering thresholds (mean>0.01 and p<=0.05 for RNA and protein, mean>0.05 and p<0.1 for epigenomic data). The model was trained using default parameters for all runs.

*MIDAS* We ran the MIDAS package following its tutorial (https://sc-midas-docs.readthedocs.io/en/latest/mosaic.html). The highly variable features (top 5,000 for RNA, top 100,000 for epigenome) were first selected, and count matrices were then used for training. For epigenomic input, the features were split by chromosome id to reduce model complexity. For all runs, the number of training epoch was set to 2,000.

*UINMF* We ran the UINMF package (version 1.0.1) following the official tutorials (https://github.com/welch-lab/liger/tree/v1.0.1/vignettes) and a benchmarking tutorial for single-cell multi-omics integration methods (https://github.com/QuKunLab/MultiomeBenchmarking/tree/main/code/Integration/pipeline/Mosaic). Specifically, we first normalized the raw count matrices using the ‘normalize’ function for all modalities. Then, we selected highly variable features using the ‘selectGenes’ function with modality-specific variance thresholds (var.thresh=0.00001 for epigenomics data and var.thresh=0.1 for RNA data). Data scaling was then performed without zero-centering, and the scaled matrices were subjected to matrix factorization with a fixed dimensionality of 30 factors across all analyses.

*MultiVI* We ran MultiVI by following the pipeline described in its tutorial (scvi version 0.19.0) (https://docs.scvi-tools.org/en/stable/tutorials/notebooks/multimodal/MultiVI_tutorial.html). Raw count matrices were used as input and the training epoch was set to 100 while the other parameters were left at their default values.

#### SpaMosaic (non-spatial)

SpaMosaic (non-spatial) is a modified version of SpaMosaic that uses an MLP as the encoder network without considering spatial information. The MLP encoder consists of two fully connective layers where the first layer is followed by a dropout layer (p=0.2) and non-linear activation ELU^99^. The rest of the model components remain unmodified, and we ran it with the same settings as those for SpaMosaic.

#### CellCharter

We ran the CellCharter package (version 0.3.5) following its official tutorial (https://cellcharter.readthedocs.io/en/latest/). For RNA-seq data, dimensionality reduction and batch effect correction were performed using scVI with a zero-inflated negative binomial (ZINB) likelihood. ATAC-seq data were processed with scVI using a Poisson likelihood, while protein data were analyzed using a modified CellCharter-scArches^100^ framework, in which the default negative binomial loss was replaced with a mean squared error (MSE) loss. The number of nearest neighbors for constructing spatial neighborhoods was determined according to the sequencing technology. For instance, for Visium and Misar-seq acquired sections with regular spatial layouts, we set the number of nearest neighbors to 6 and 8, respectively.

#### BANKSY

We ran the BANKSY package (version 1.2.1) following the official python tutorial (https://github.com/prabhakarlab/Banksy_py/blob/Bansky_1.2.1/DLPFC_harmony_multisample.ipynb). We first selected the top 2,000 highly variable genes. Principal component analysis was next performed on the BANKSY-generated spatially-smoothed embeddings from individual tissue sections, followed by Harmony-based batch correction for dataset integration.

### Clustering

For SpaMosaic, we applied model-based clustering using the Mclust package^101^ across all tests, with the parameter modelNames=‘EEE’. For the other methods, we adopted the recommended clustering algorithms in their original publication or the official tutorials. Specifically, for UINMF, Cobolt, and MIDAS, the Louvain^102^ clustering algorithm was used. For BANKSY, CLUE, MultiVI, scMoMaT, and StabMap, Leiden clustering was applied. For SpaMosaic (non-spatial), Mclust was applied. For CellCharter, we used its built-in Gaussian Mixture Model (GMM)-based clustering approach as implemented in the official repository with all parameters setting as default.

To determine the target number of clusters for each dataset, we first consulted the original publications or data portals for their number of clusters identified in each individual section. For each dataset, we selected the maximum number of clusters across all sections as the target number of clusters. The only exception was the postnatal mouse brain (DBiT+spRA+spA) dataset where the four sections differ substantially in the size of their corresponding anatomical regions. In this case, we used the median number of clusters across the four sections as the final target. To achieve the target number of clusters with Leiden and Louvain algorithms, we performed clustering at different resolutions ranging from 0 to 2 (with step size 0.01) and selected the resolution that yielded the cluster count closest to the target value.

### Differential expression analysis and GO enrichment analysis

We used the Wilcoxon test as implemented in the rank_genes_groups function of the Scanpy package (version 1.9.6)^103^ to identify DEGs of spatial clusters. We performed the GO enrichment analysis on cluster specific DEGs using the R package clusterProfile (version 4.6.2)^55^.

## Supporting information

supplementary notes for the main text

## Data availability

All datasets used in this study were obtained from public repositories, except for the in-house generated human lymph node and tonsil datasets. The Misar-seq sections were downloaded from https://www.biosino.org/node/project/detail/OEP003285 and https://pan.baidu.com/s/15i0R5na2Jud_7EpkWSaZpA?pwd=oipj. The Stereo-seq sections were obtained from https://db.cngb.org/stomics/mosta/download/. The spatial RNA-Epigenomics seq sections were accessed via https://www.ncbi.nlm.nih.gov/geo/query/acc.cgi?acc=GSE205055 and https://cells.ucsc.edu/?ds=brain-spatial-omics. The Spatial ATAC-seq, Spatial Cut&Tag-seq, Spatial Mux-seq, and DBiT-ARP-seq sections were retrieved from NCBI GEO under the accession numbers GSE171943, GSE165217, GSE263333, and GSE308623, respectively. The mouse brain 10x Visium and Visium HD datasets were obtained from the publicly available 10x Genomics resources (https://www.10xgenomics.com/datasets/mouse-brain-section-coronal-1-standard and https://www.10xgenomics.com/datasets/visium-hd-cytassist-gene-expression-libraries-of-mouse-brain-he). All processed datasets are publicly available at https://zenodo.org/records/18946723.

## Code availability

The SpaMosaic open-source Python package is available at https://github.com/JinmiaoChenLab/SpaMosaic/tree/main, and all code for reproducing the benchmarking results presented in this study can be accessed at https://github.com/JinmiaoChenLab/SpaMosaic/tree/SpaMosaic-reproduce and archived at Zenodo (https://doi.org/10.5281/zenodo.18402871).

## Authors’ contributions

J.C. and M.L. initiated the project and provided funding support. J.C. conceived the idea. X.Y., J.C.,Z.F., and R.Z. developed the method and designed the experiments. X.Y., Z.F., J.C., and K.S.A. performed the data analysis and wrote the manuscript. L.V.O., A.E., T.W. and D.G. provided the lymph node and tonsil datasets, and manually annotated the tissue structures. D. Z. and R.F. provided the postnatal mouse brain and mouse embryo datasets.

## Acknowledgements

The research was supported by A*STAR under its BMRC Central Research Fund (CRF, UIBR) Award; AI, Analytics and Informatics (AI3) Horizontal Technology Programme Office (HTPO) seed grant (Spatial transcriptomics ST in conjunction with graph neural networks for cell–cell interaction #C211118015) from A*STAR, Singapore; Open Fund Individual Research Grant (Mapping hematopoietic lineages of healthy and high-risk acute myeloid leukemia patients with FLT3-ITD mutations using single-cell omics #OFIRG18nov-0103) from Ministry of Health, Singapore; National Research Foundation (NRF), Award no. NRF-CRP26-2021-0001; the National Research Foundation, Singapore, and Singapore Ministry of Health’s National Medical Research Council under its Open Fund-Large Collaborative Grant (“OF-LCG”) (MOH-OFLCG18May-0003); Singapore National Medical Research Council (#NMRC/OFLCG/003/2018). M.L. is supported in part by the National Natural Science Foundation of China under Grant (No.62225209 to M.L.). We thank Dr. Leslie A Rubio Rodríguez-Kirby and Dr. Gonçalo Castelo-Branco from Karolinska Institute for generously sharing the DBiT-ARP-seq data.

---

**Supp. Fig. S1.**
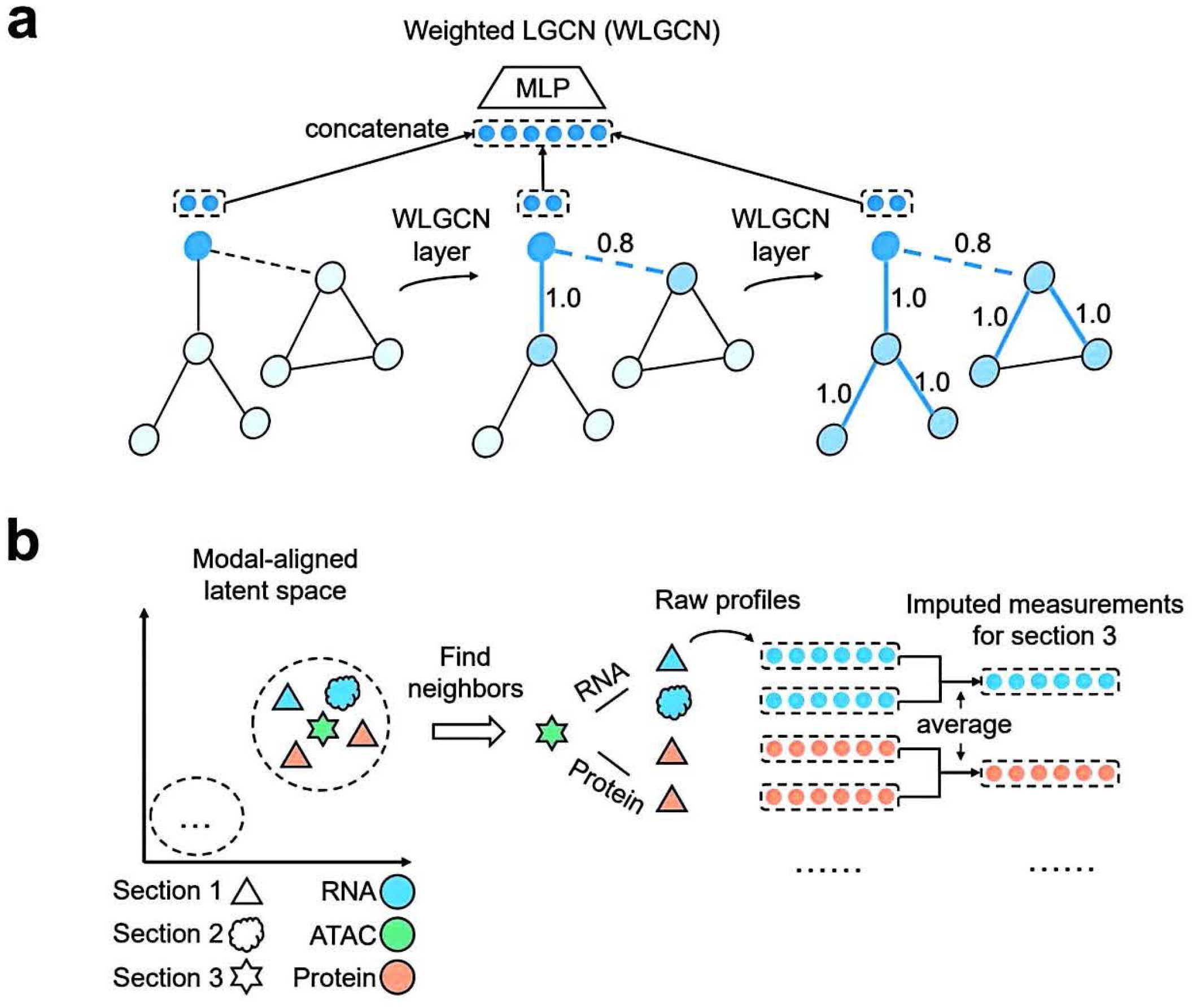
Graph neural network architecture and imputation strategy. **a** Aggregation and propagation processes of weighted LGCN. The weight for spatial neighbors is set to I while the weight for expression neighbors is set to 0.8. In each layer, each node aggregates the information from its neighbors using the weighted average. After two layers, the information from each node is concatenated across three output layers and then processed by a multiple layer perceptron. **b** Imputation process. Taking section 3 as an example, which only has ATAC modality, we aim to impute its RNA and protein profiles. To impute the RNA profile of each spot in section 3, we find its RNA-specific neighbors from the other sections and then compute the average target expression profiles of these neighbors. The same approach is also taken for other modalities.

**Supp. Fig. S2.**
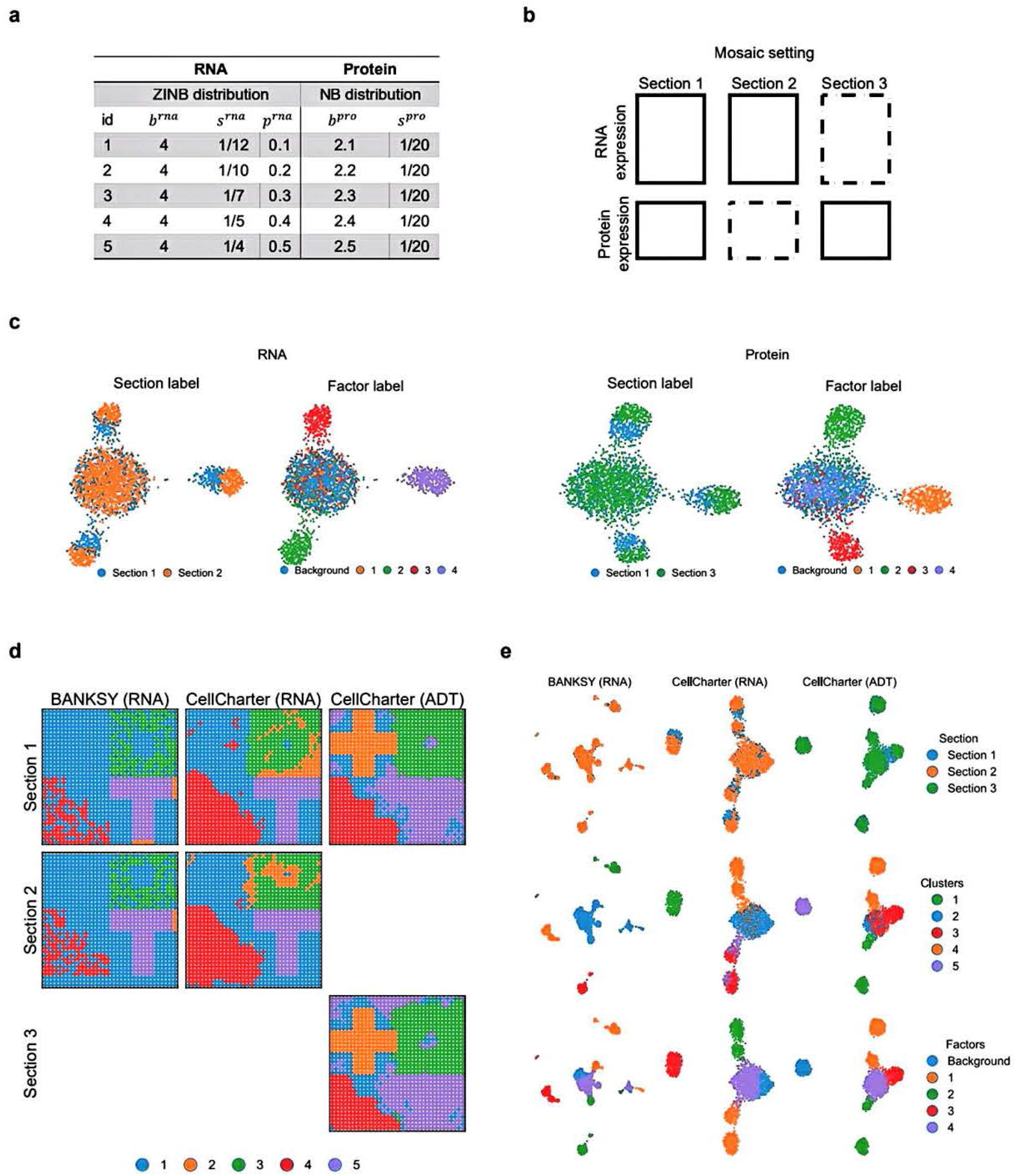
Simulation settings and plots of results from baseline methods. **a** Five sets of simulation parameters used for data generation. **b** Mosaic configuration of each dataset. A dotted box denotes that the modality is missing. **c** UMAP plots of RNA and protein profiles before batch correction. Spots are colored by section and factor labels for each modality. **d** Spatial plots of clustering results from BANKSY and CellCharter applied to each modality. **e** UMAP plots of embeddings from BANKSY and CellCharter applied to each modality.

**Supp. Fig. S3.**
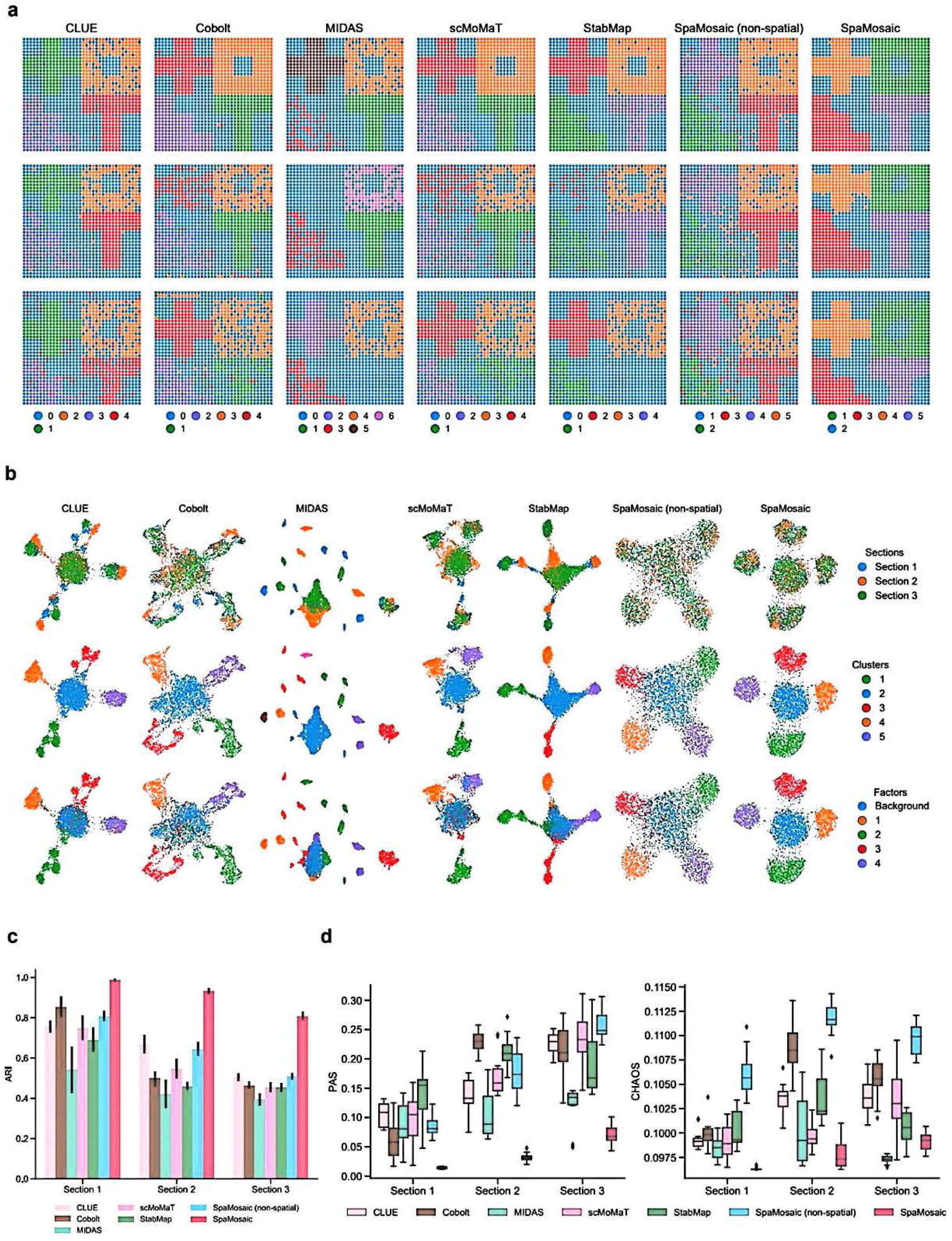
Quantitative evaluation of individual sections and plots of results from benchmarked methods. **a** Spatial plotsof the clustering results of benchmarked methods. **b** UMAP plots of embeddings from benchmarked methods. From top to bottom, spots are colored by section, cluster, and factor labels. **c** ARI scores of benchmarked methods on individual sections across five simulation settings. **d** PAS and CHAOS scores of benchmarked methods on individual sections across five simulation settings.

**Supp. Fig. S4.**
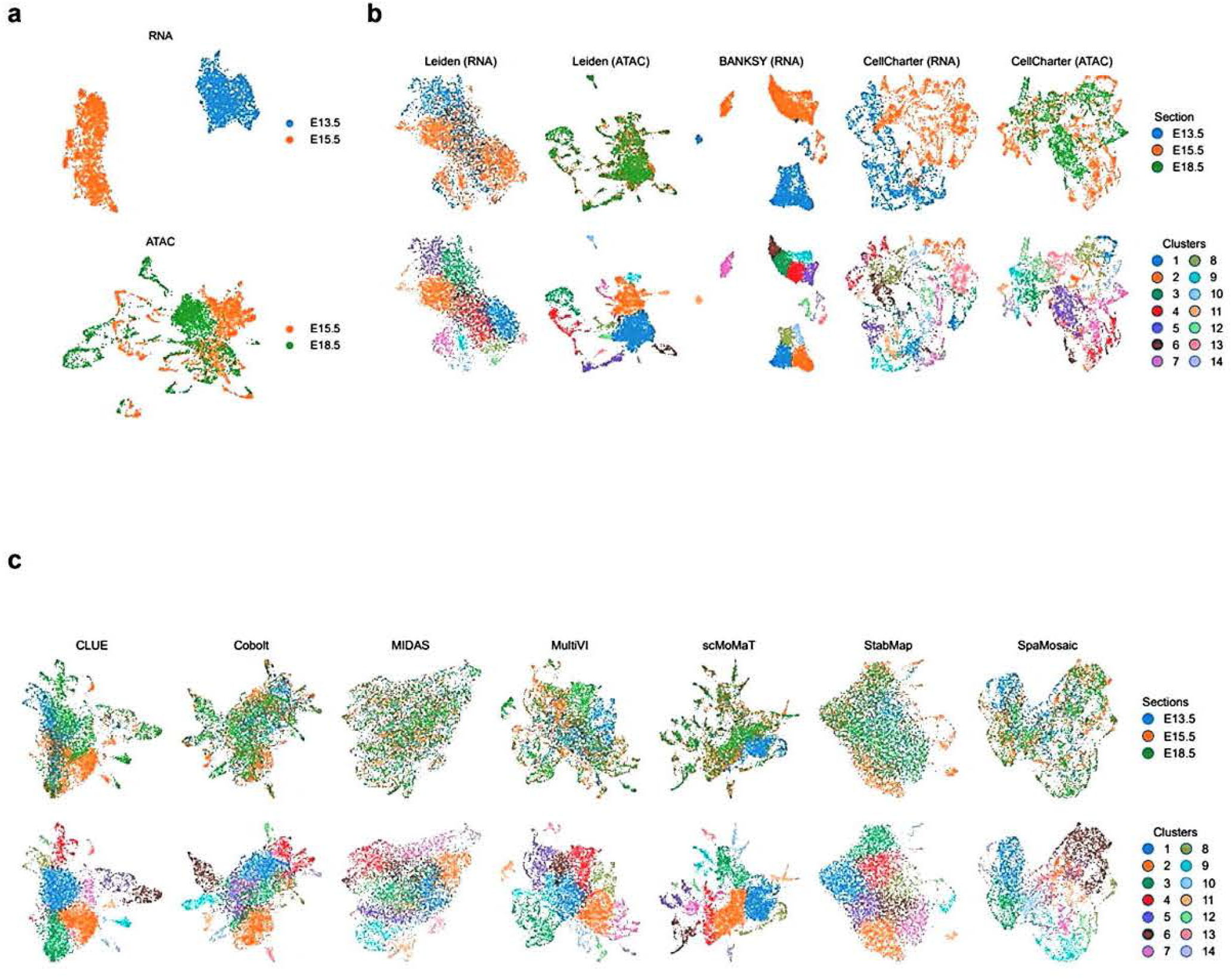
UMAP plots of embryonic mouse brain (Misar) dataset integration analysis. **a** UMAP plots of the unintegrated RNA and ATAC profiles. Spots are colored by section labels. **b** UMAP plots of the embeddings from baseline methods. Spots in the first row are colored by section and the second row by cluster. **c** UMAP plots of the embeddings from benchmarked methods.

**Supp. Fig. S5.**
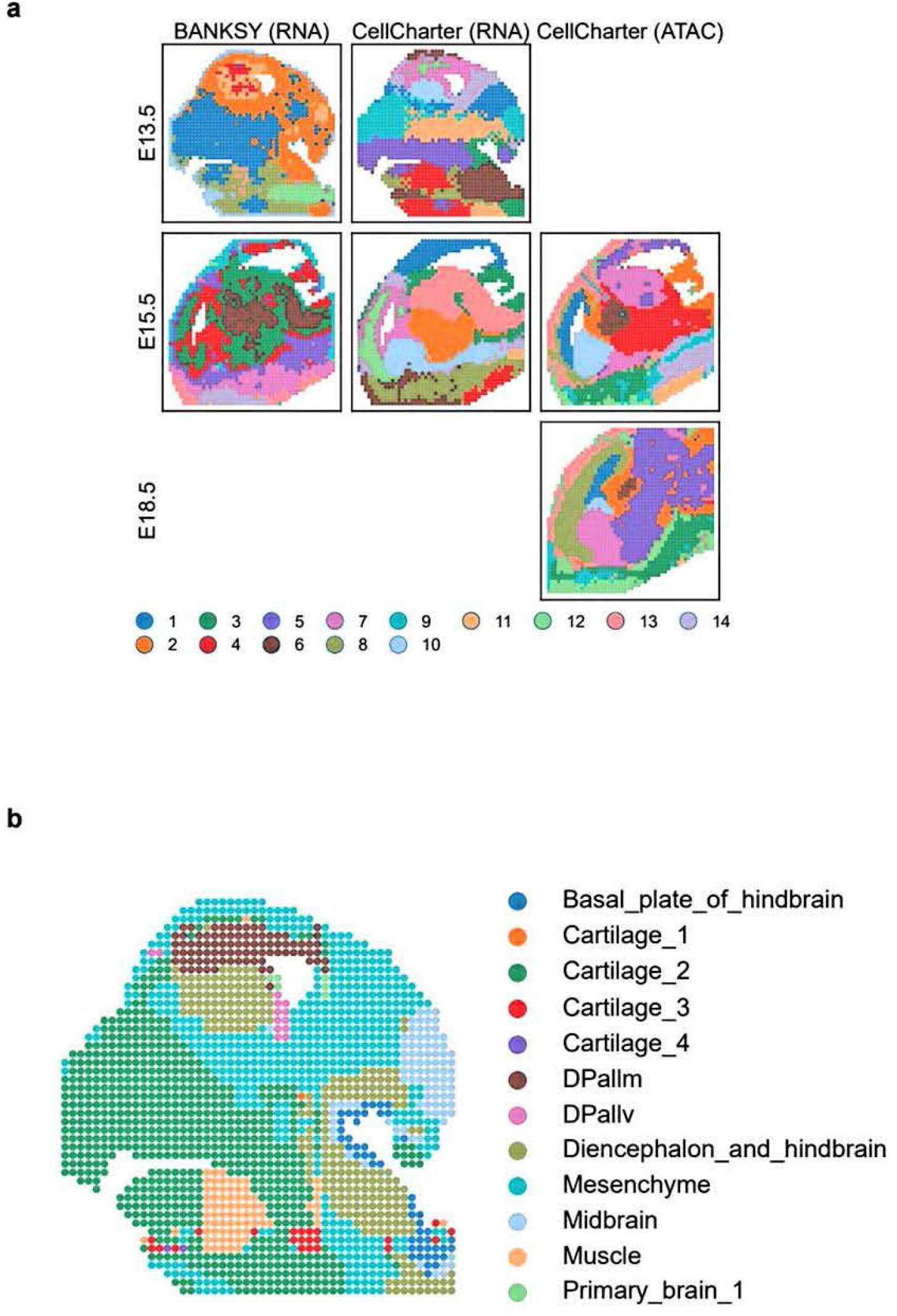
Spatial plots of embryonic mouse brain (Misar) dataset integration analysis. **a** Spatial plots of clustering results from baseline methods. **b** Reference annotation for the El3.5 section from the original publication.

**Supp. Fig. S6.**
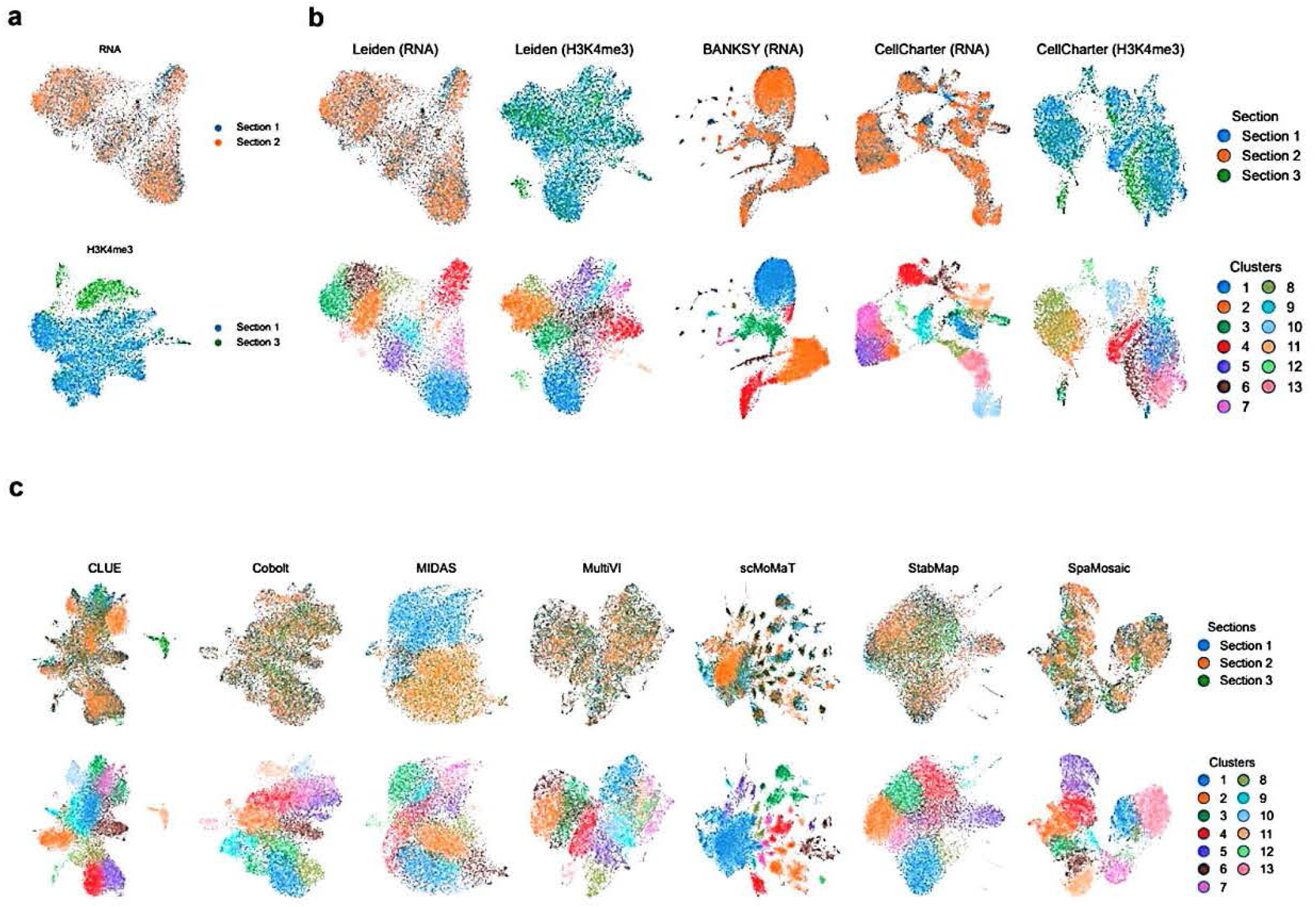
UMAP plots of postnatal mouse brain dataset (spRH+spC&T) integration analysis. **a** UMAP plots of the unintegrated RNA and H3K4me3 histone modification profiles. Spots are colored by section labels. **b** UMAP plots of the embeddings from baseline methods. Spots in the first row are colored by section and the second row by cluster. **c** UMAP plots of the embeddings from benchmarked methods.

**Supp. Fig. S7.**
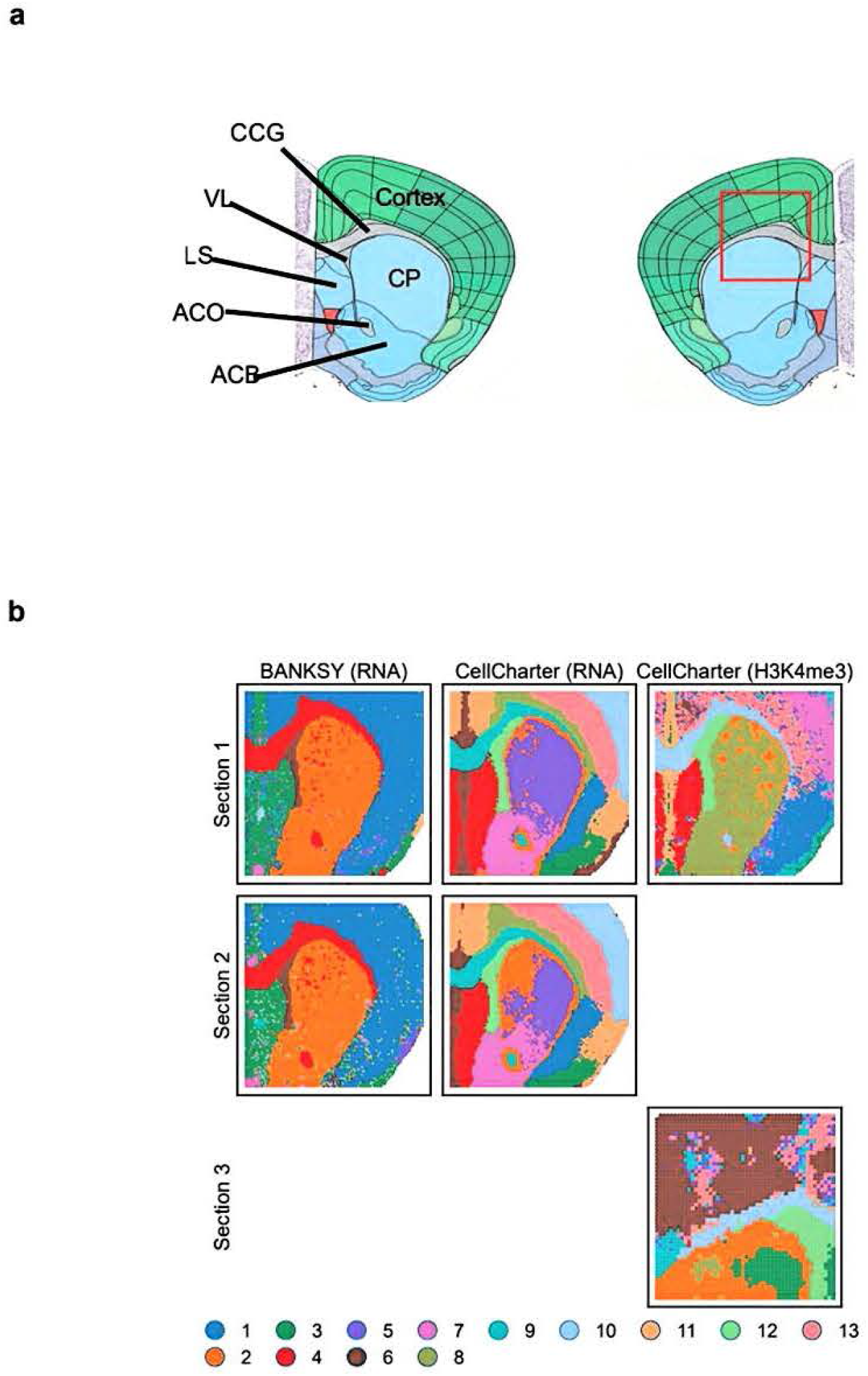
Reference atlas and spatial plots of clustering results from baseline methods on the postnatal mouse brain (spRH+spC&T) dataset. **a** Postnatal mouse brain structure reference atlas. Images were taken from the Allen Reference Atlas (https://atlas.brain-map.org/). **b** Spatial plots of clustering results from baseline methods.

**Supp. Fig. S8.**
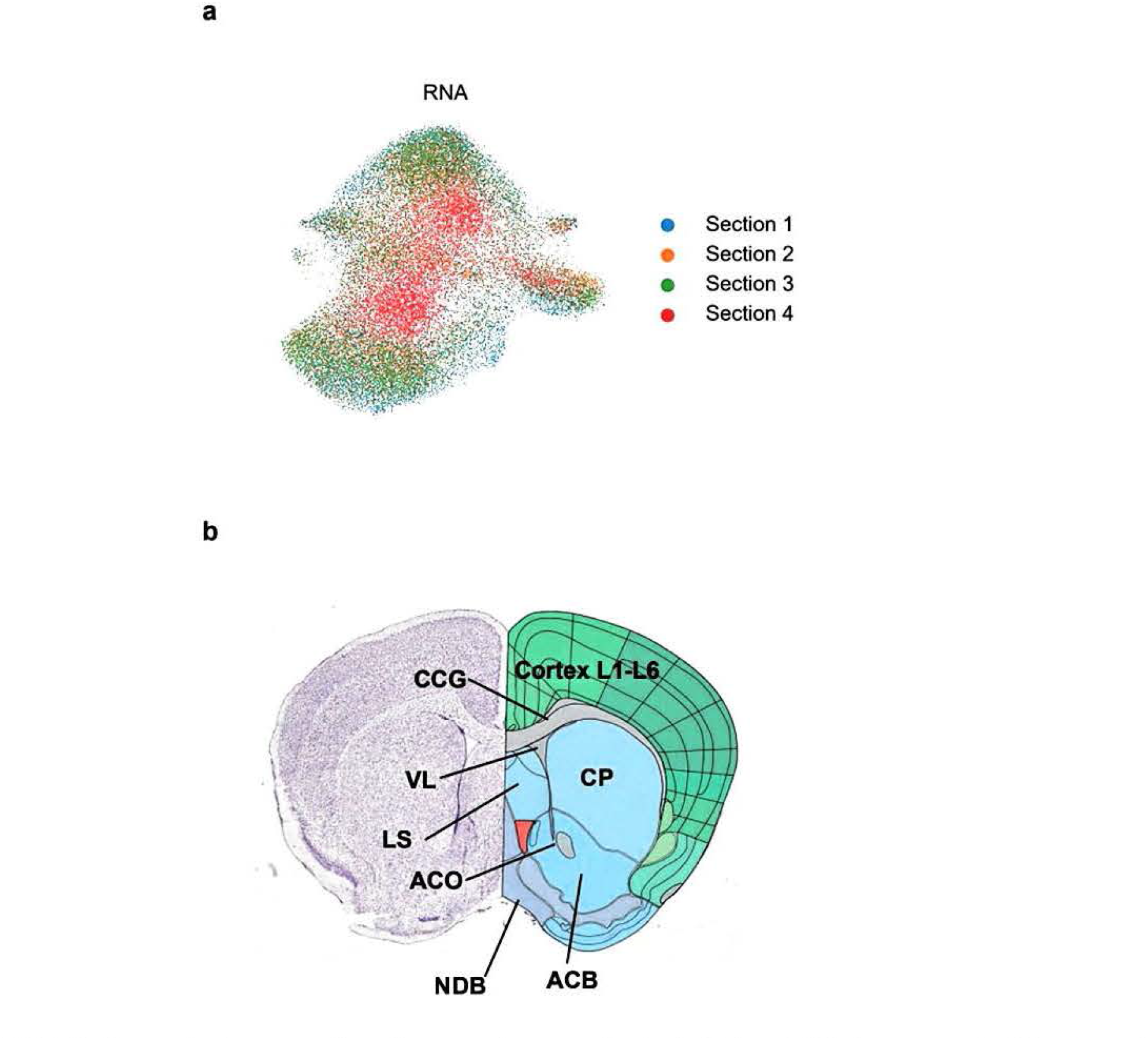
UMAP plot and reference atlas of postnatal mouse brain (spRA+spRH) dataset. **a** UMAP plots of raw RNA profiles. Spots are colored by section labels. **b** Reference atlas of postnatal mouse brain structures. Images were from the Allen Reference Atlas (https://atlas.brain-map.org/).

**Supp. Fig. S9.**
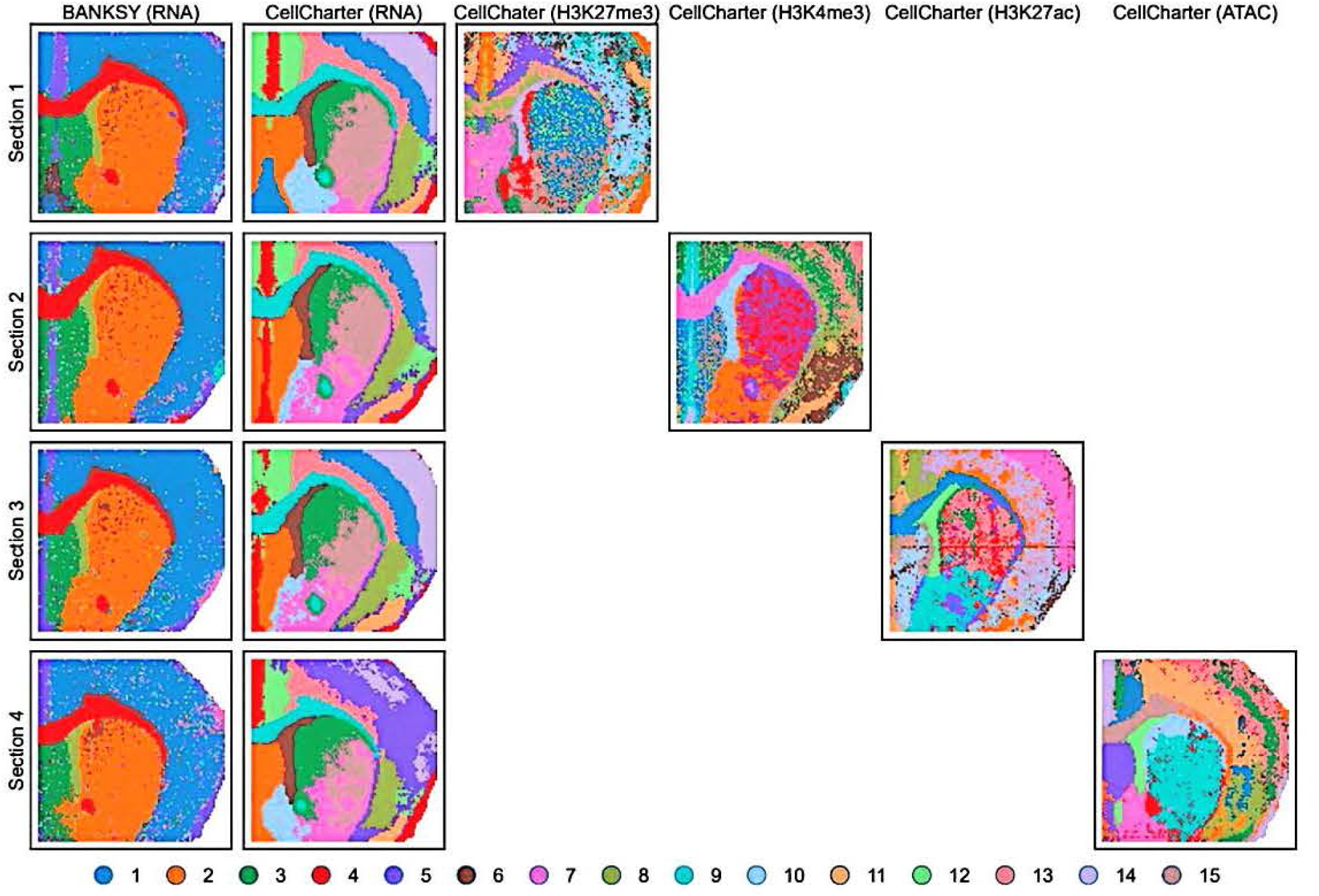
Spatial plots of clustering results from baseline methods on the postnatal mouse brain (spRA+spRH) dataset.

**Supp. Fig. S10.**
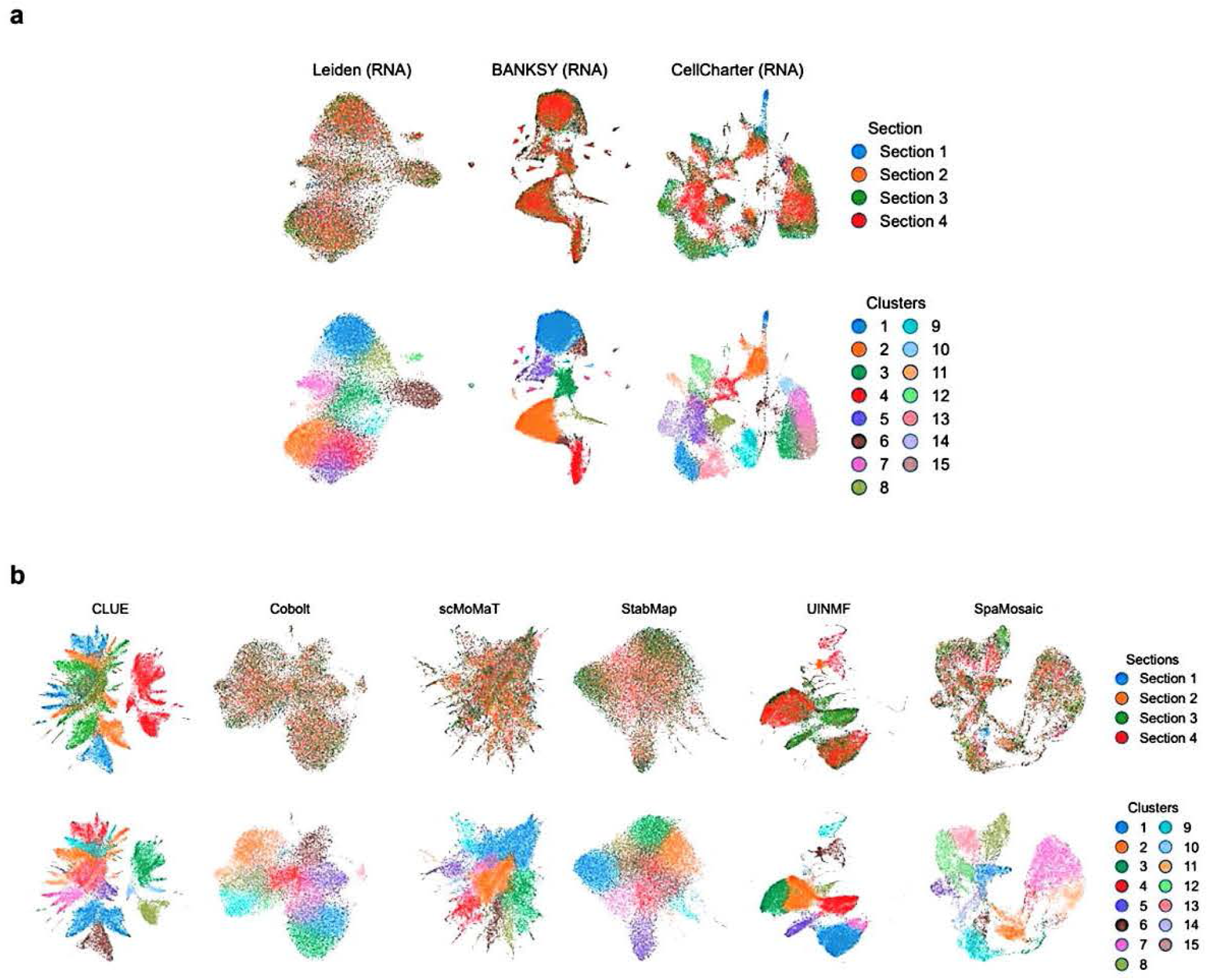
UMAP plots of postnatal mouse brain dataset (spRA+spRH) integration analysis. **a** UMAP plots of the embeddings from baseline methods. Spots in the first row are colored by section and the second row by cluster. **b** UMAP plots of the embeddings from benchmarked methods.

**Supp. Fig. S11.**
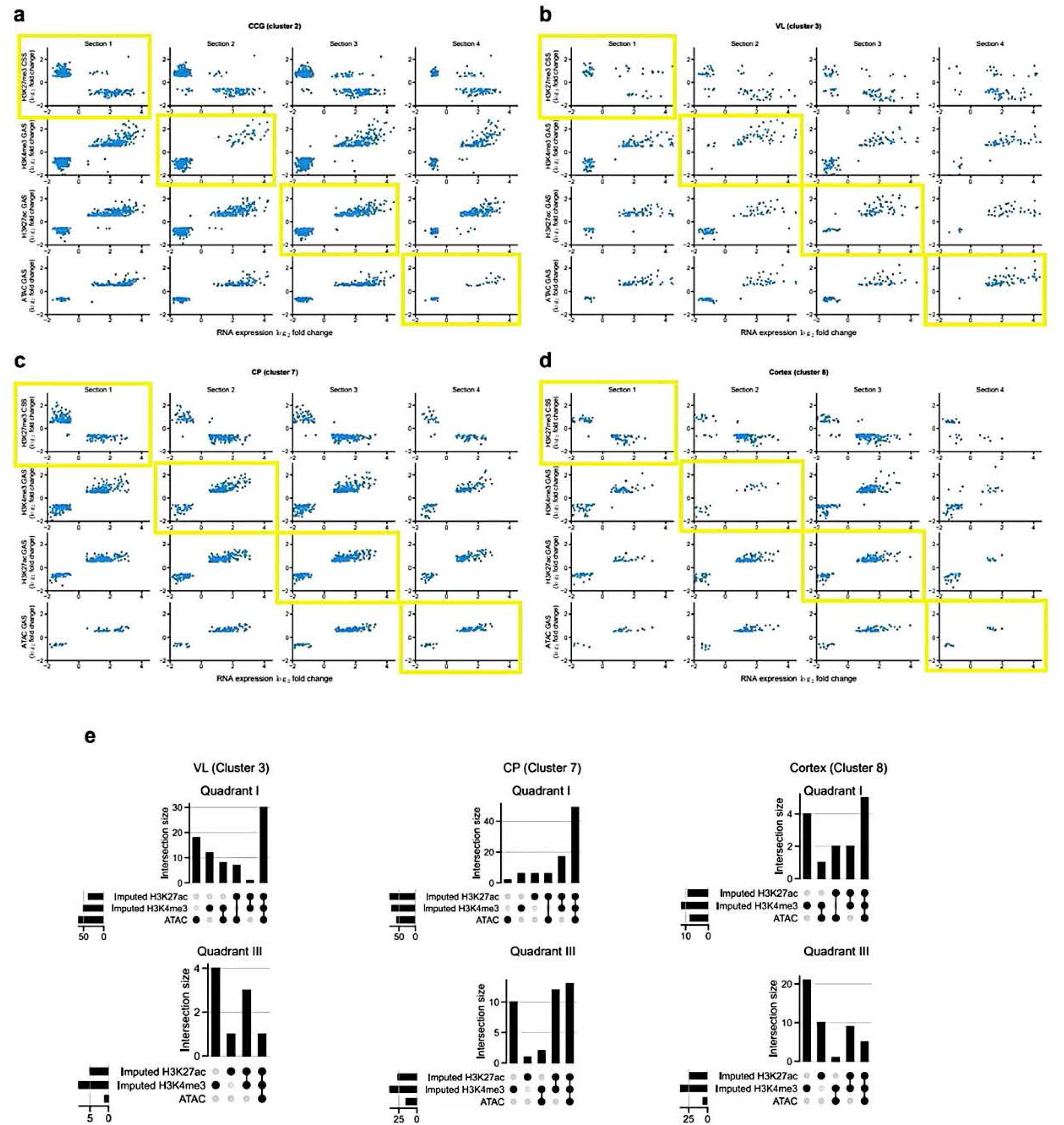
Analyzing the correlation between RNA and GAS/CSS in selected brain regions. All genes plotted have an adjusted p value less than le-5 and absolute value of log2 fold change larger than 0.5. The yellow boxes in **a, b, c, d** denoted the relationship between the measured in each section. **a** Correlation between RNA expression and GAS/CSS for CCG-specific (SpaMosaic’s cluster 2) genes across the four sections. **b** Correlation between RNA expression and GAS/CSS for VL-specific (SpaMosaic’s cluster 3) genes across all sections. **c** Correlation between RNA expression and GAS/CSS for CP-specific (SpaMosaic’s cluster 7) genes across all sections. **d** Correlation between RNA expression and GAS/CSS for the cortex-specific (SpaMosaic’s cluster 8) genes across all sections. **e** Upset plots of three region-specific gene sets in quadrants I (top) and III (bottom) based on correlations between measured RNA and measured ATAC, imputed H3K4me3, and imputed H3K27ac for section 4.

**Supp. Fig. S12.**
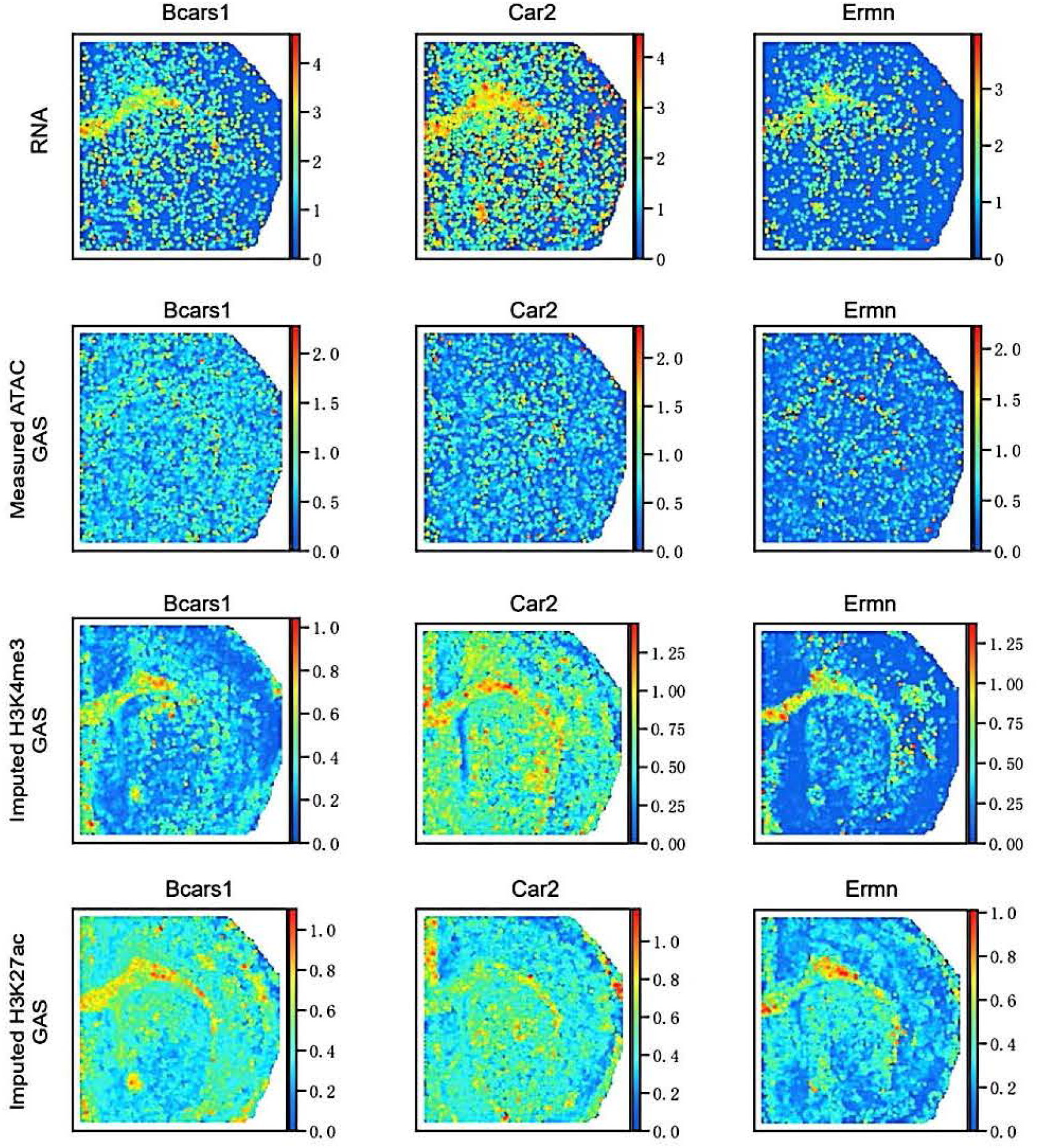
Spatial heatmap of measured and imputed gene expression/gene activity scores. Spatial heatmap of measured RNA, measured ATAC GAS, imputed H3K4me3 GAS and imputed H3K27ac GAS for CCG-specific genes Beas I, Car2 and Ermn.

**Supp. Fig. S13.**
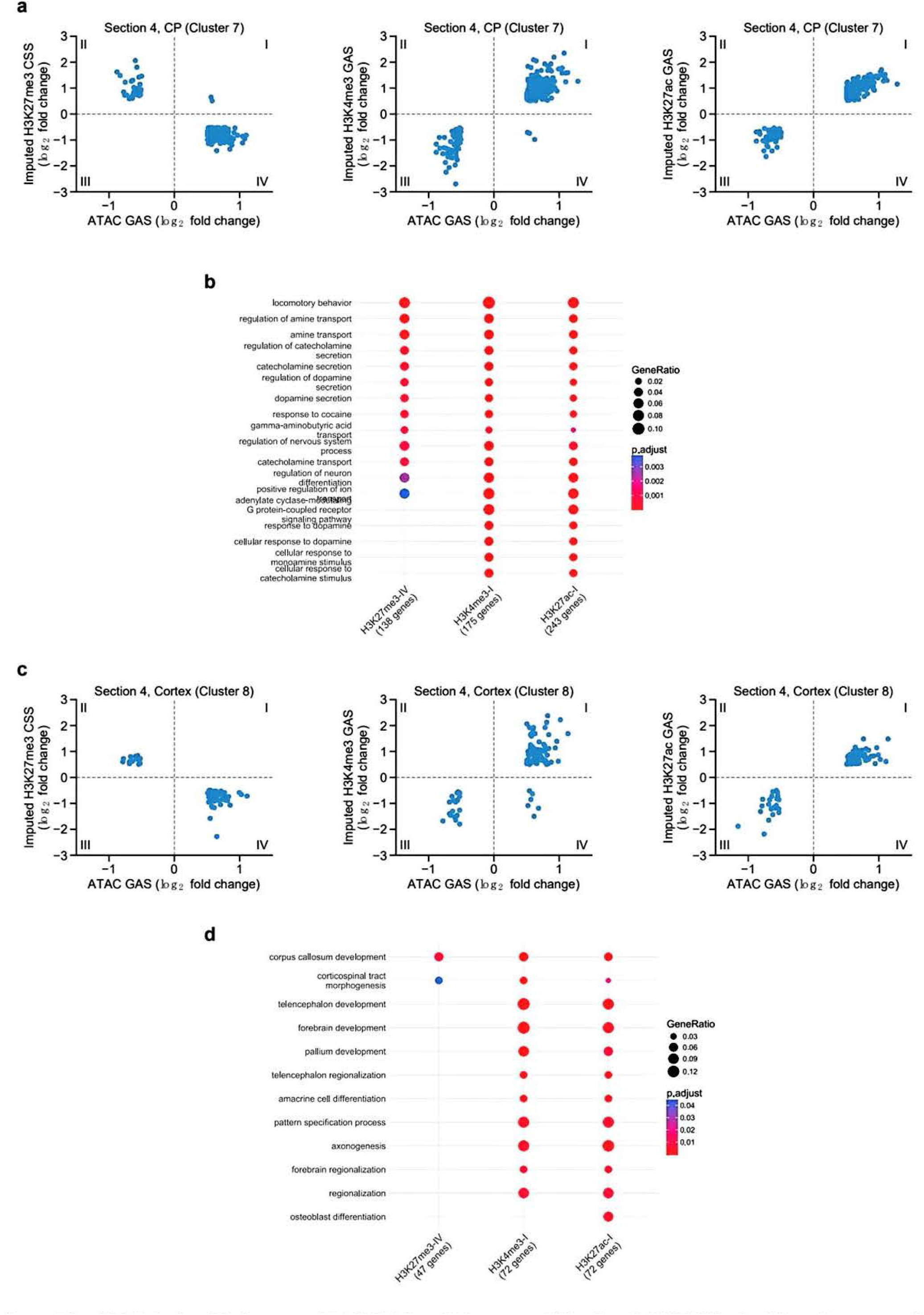
Relationship between ATAC GAS and histone modification GAS/CSS in the CP and cortex regions. **a** Relationship between imputed GAS/CSS and measured ATAC GAS for CF-specific genes on the fourth section. **b** Enriched GO terms for CP-specific genes in quadrants I and IV of panel **a. c** Relationship between imputed GAS/CSS and measured ATAC GAS for cortex-specific genes on the fourth section. **d** Enriched GO terms for cortex-specific genes in quadrants I and IV in panel **c**.

**Supp. Fig. S14.**
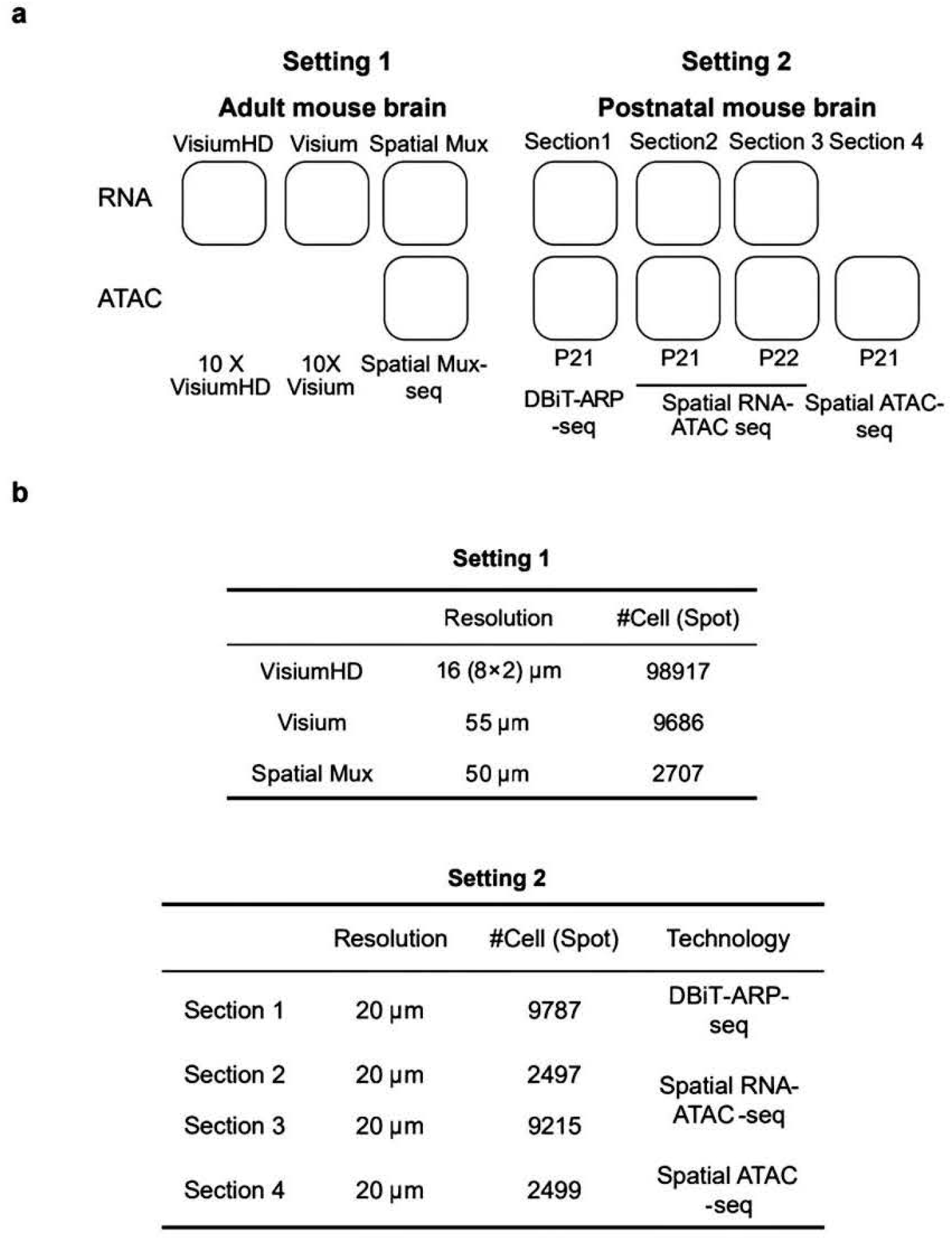
Dataset overview of scenarios 1 and 2 in cross-technology integration. **a** Configuration of data modalities for both integration scenarios. **b** Metadata for each section in the two scenarios, including resolutions (spot/pixel diameter) and spot counts.

**Supp. Fig. S15.**
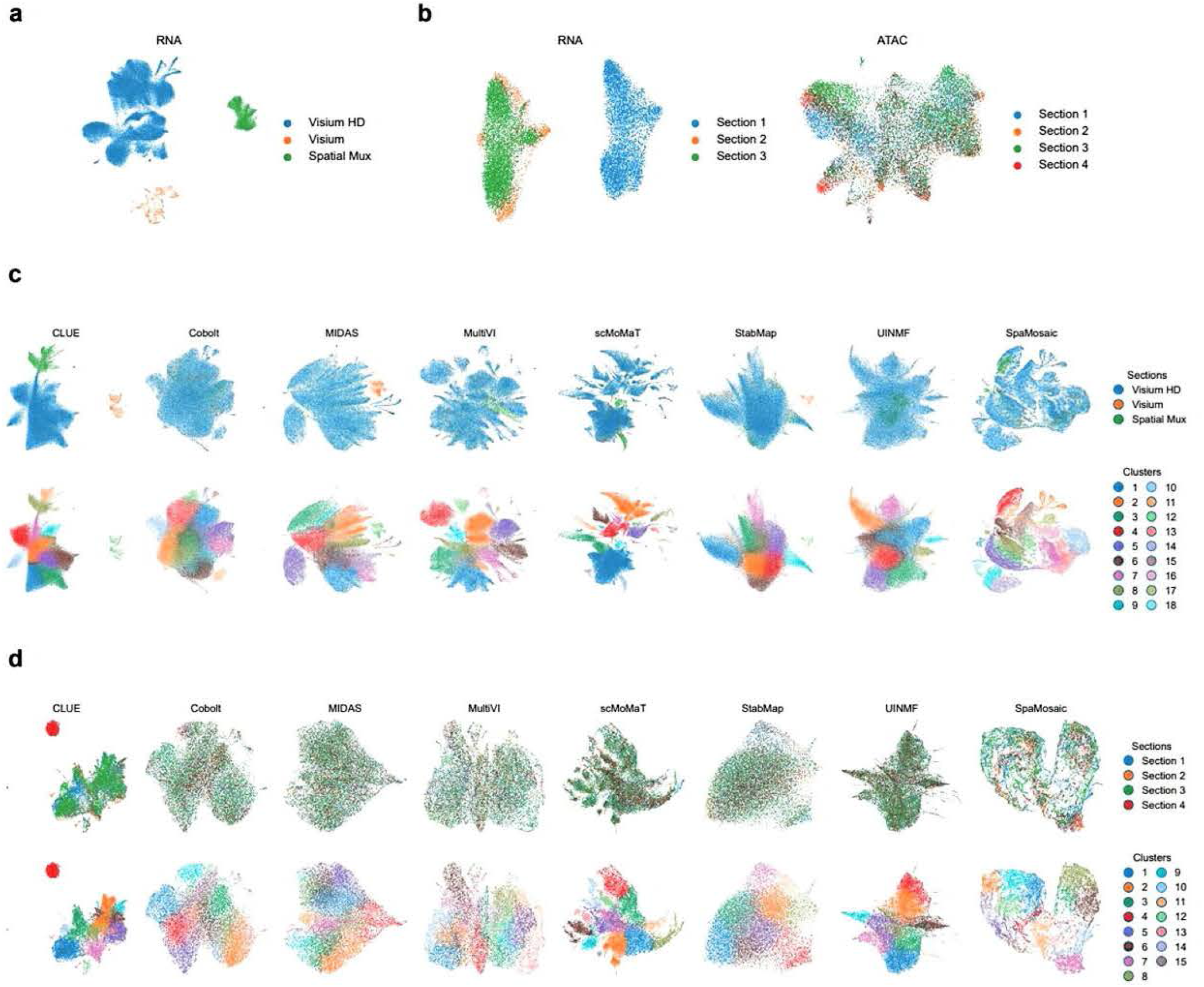
UMAP plots of cross-technology integration analysis. **a** UMAP plots of the unintegrated RNA profiles in scenario 1. Spots are colored by section labels. **b** UMAP plots of the unintegrated RNA and ATAC profiles in scenario 2. **c** UMAP plots of the embeddings from all mosaic integration methods in scenario l. **d** UMAP plots of the embeddings from all mosaic integration methods in scenario 2.

**Supp. Fig. S16.**
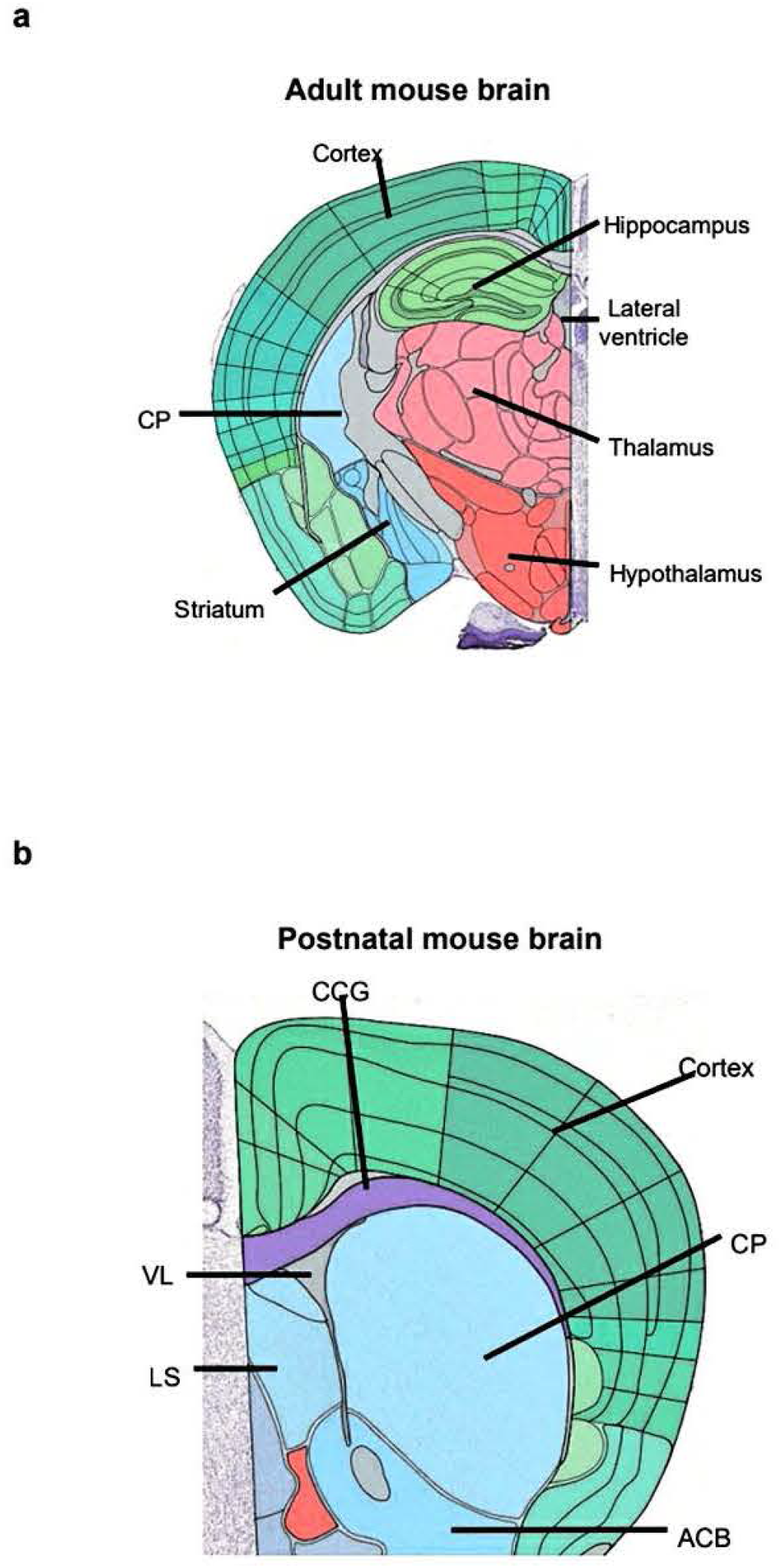
Reference atlas of mouse brain for cross-technology integration analysis. **a** Reference atlas of adult mouse brain structures. **b** Reference atlas of postnatal mouse brain structures. Both images were from the Allen Reference Atlas (https://atlas.brain-map.org/)

**Supp. Fig. S17.**
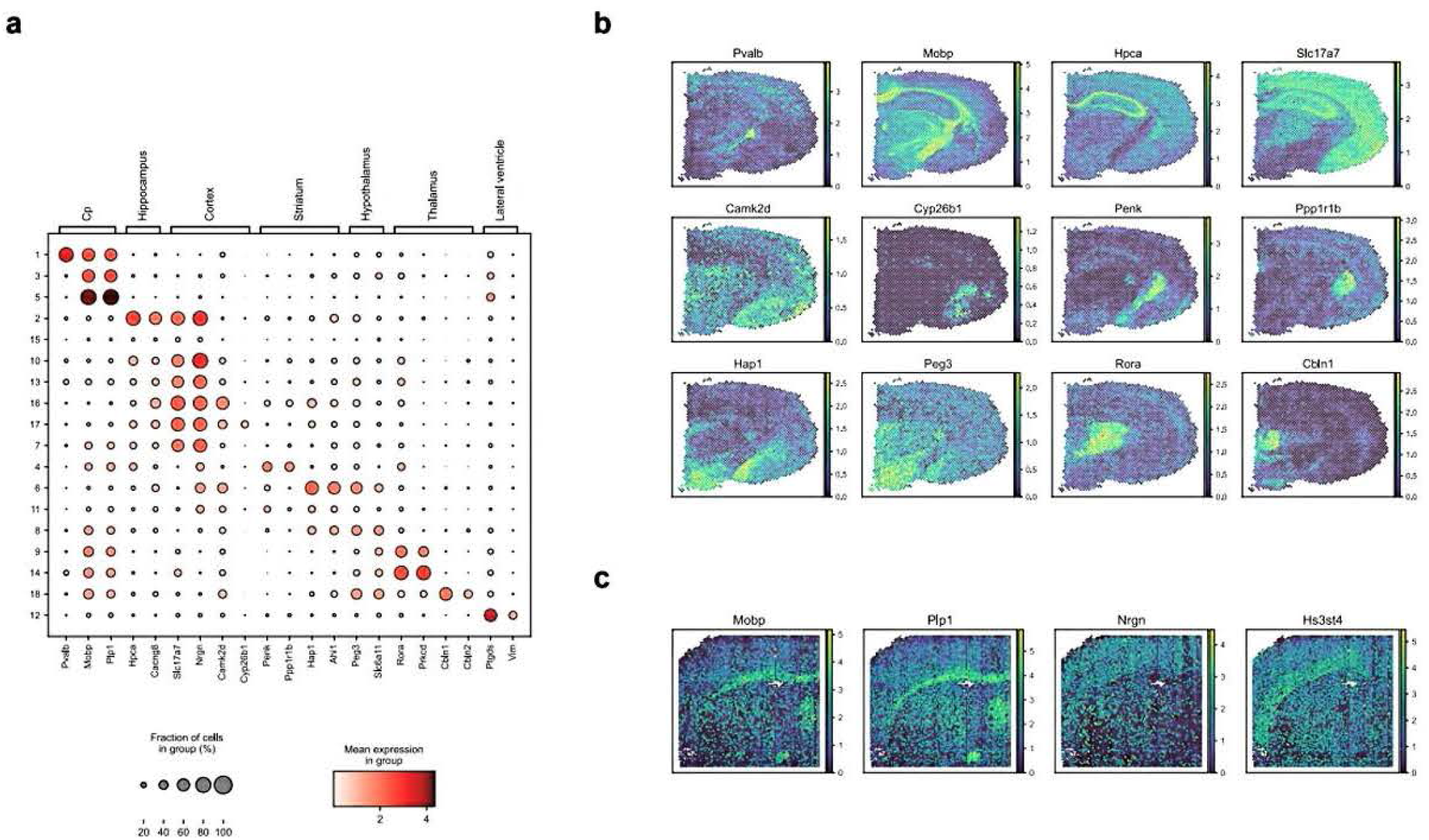
Marker gene validation in cross-technology integration (Scenario 1). **a** Dot plot of region-specific marker gene expression in SpaMosaic’s clusters. **b** Spatial heatmap of RNA expression for selected marker genes in the 10x Genomics Visium section. **c** Spatial plots of RNA expression for selected marker genes in the Spatial Mux-seq section.

**Supp. Fig. S18.**
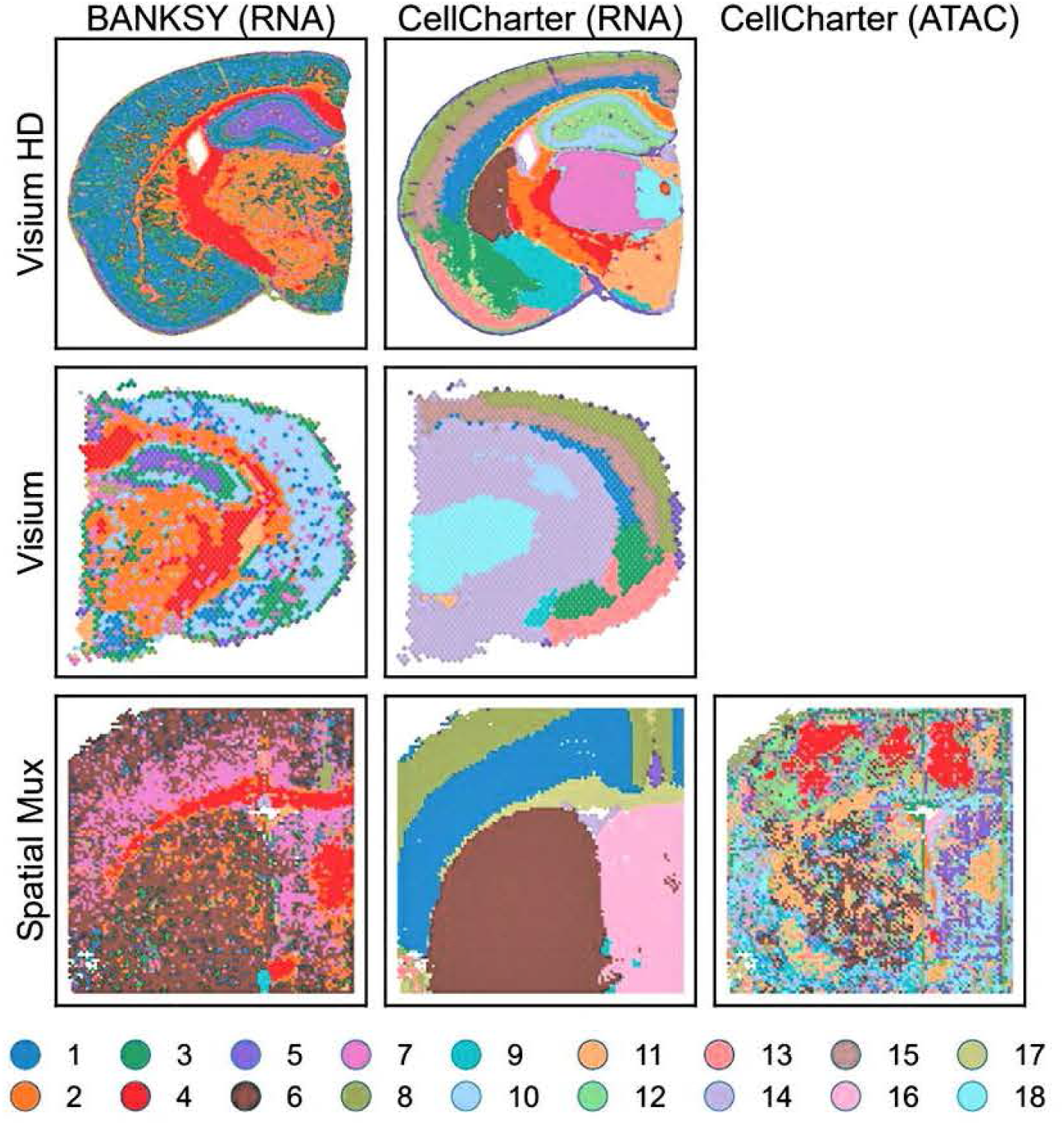
Spatial plots of clustering results from BANKSY and CellCharter in cross-technology integration (Scenario 1).

**Supp. Fig. S19.**
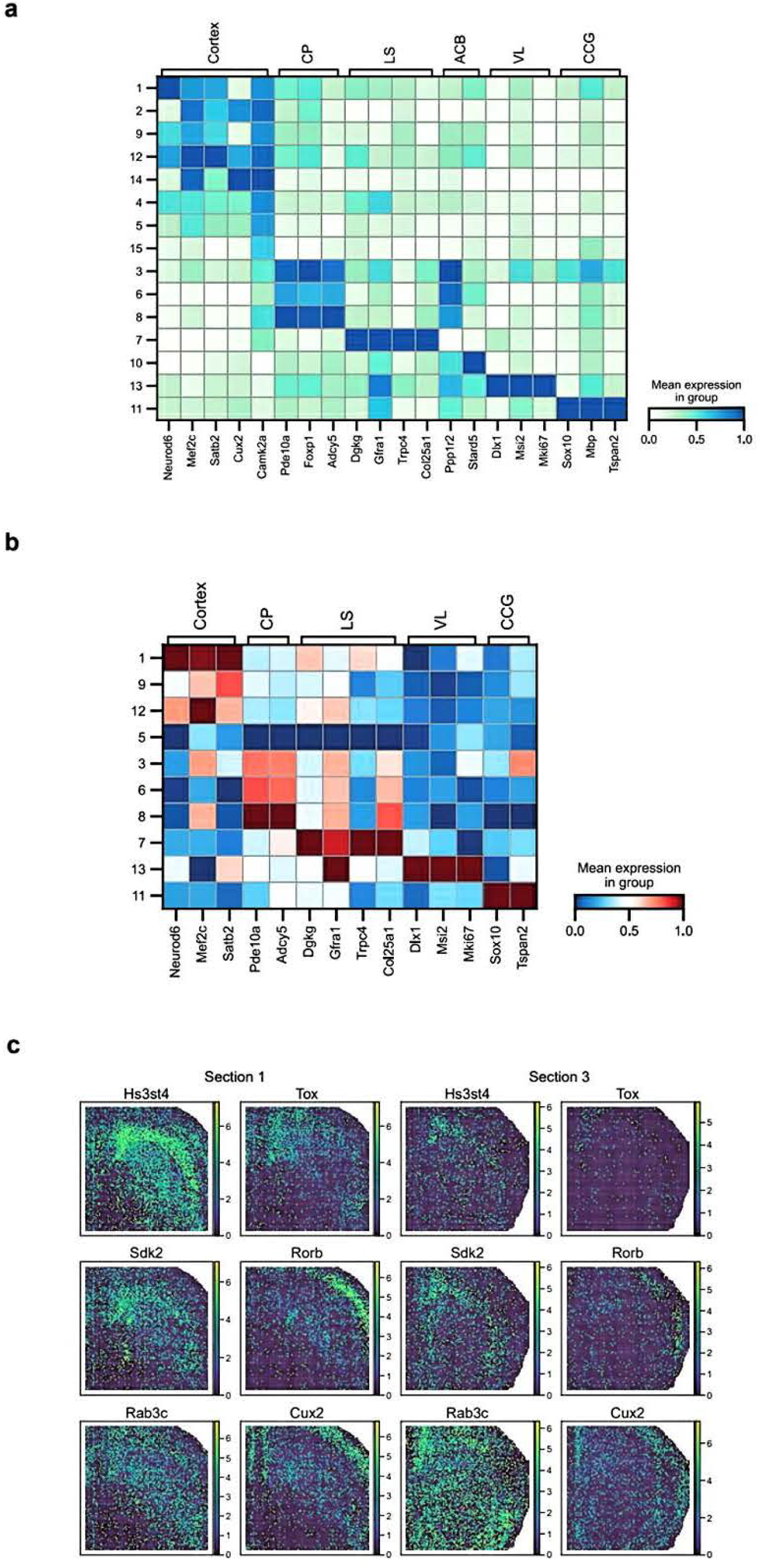
Marker gene expression and gene activity validation in cross-technology integration (Scenario 2). **a** Average RNA expression of region-specific marker genes in SpaMosaic’s clusters. **b** Average gene activity scores of region-specific marker genes in SpaMosaic’s clusters. **c** Spatial plots of RNA expression for cortical layer-specific markers *(Hs3st4, Tox, Sdk2* for deeper layers; *Rorb, Rab3c, Cux2* for upper layers) in sections 1 and 3.

**Supp. Fig. S20.**
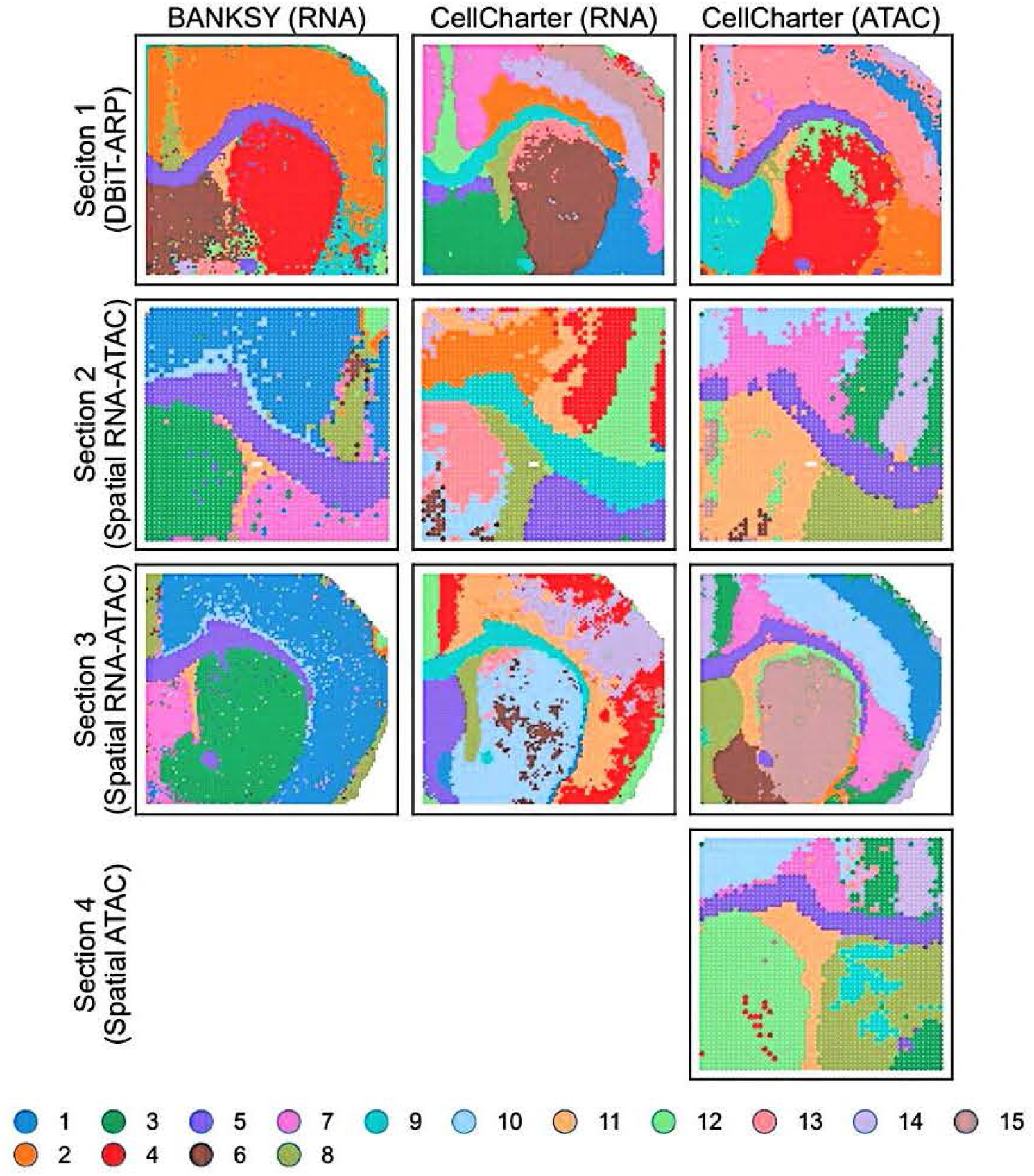
Spatial plots of clustering results from BANKSY and CellCharter in cross-technology integration (Scenario 2).

**Supp. Fig. S21.**
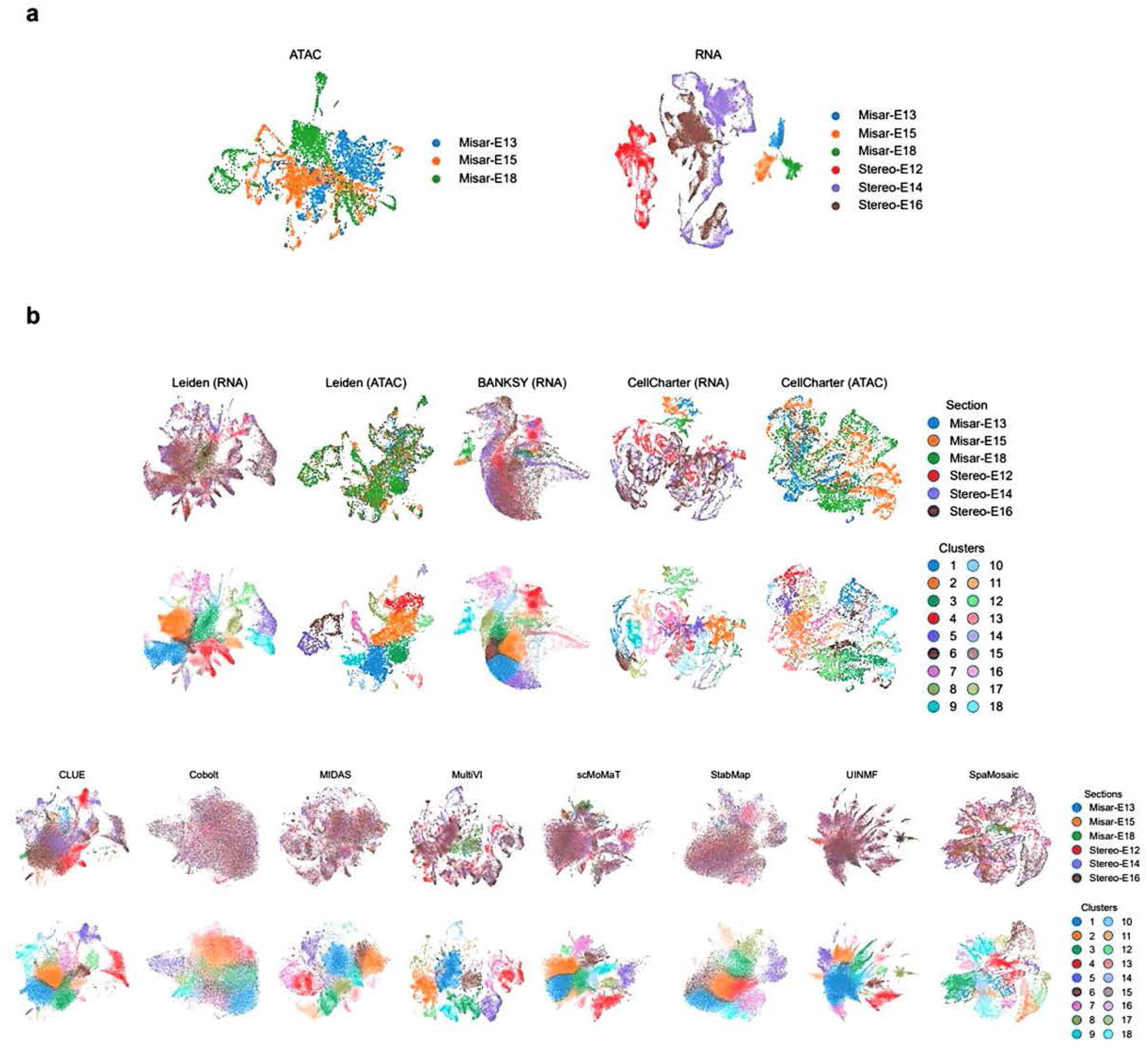
UMAP plots of embryonic mouse brain dataset (Misar+Stereo) integration analysis. **a** UMAP plots of the unintegrated ATAC and RNA profiles. Spots are colored by section labels. **b** UMAP plots of the embeddings from baseline methods and mosaic integration methods.

**Supp. Fig. S22.**
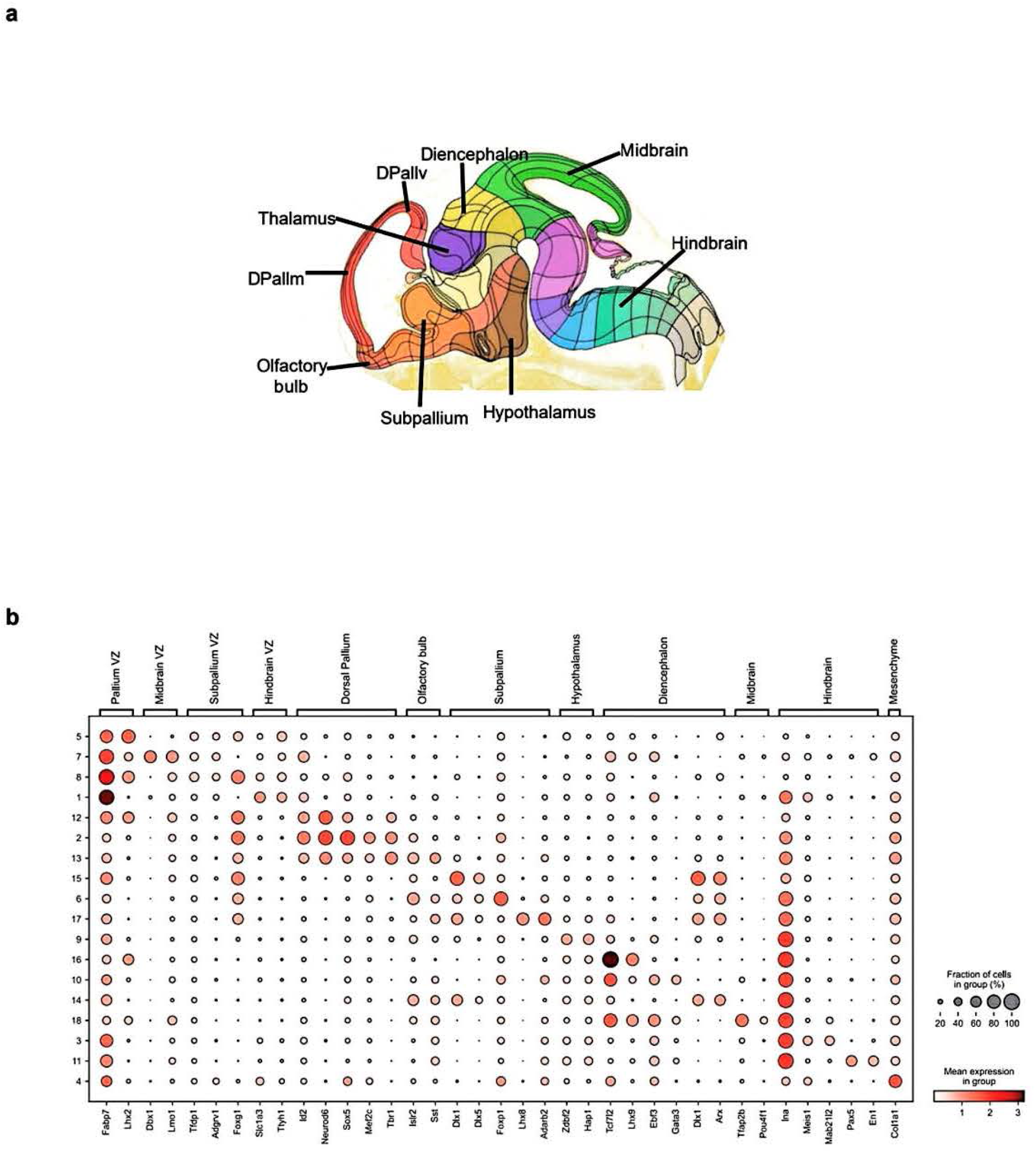
Reference structure and marker gene expression validation of SpaMosaic’s identified clusters in embryonic mouse brain (Misar+Stereo) dataset. **a** Reference atlas of developing mouse brain structures at the El3.5 stage. Image was taken from the Allen Reference Atlas (https://atlas.brain-map.org/). **b** Dot plot of region-specific marker gene RNA expression in SpaMosaic’s clusters.

**Supp. Fig. S23.**
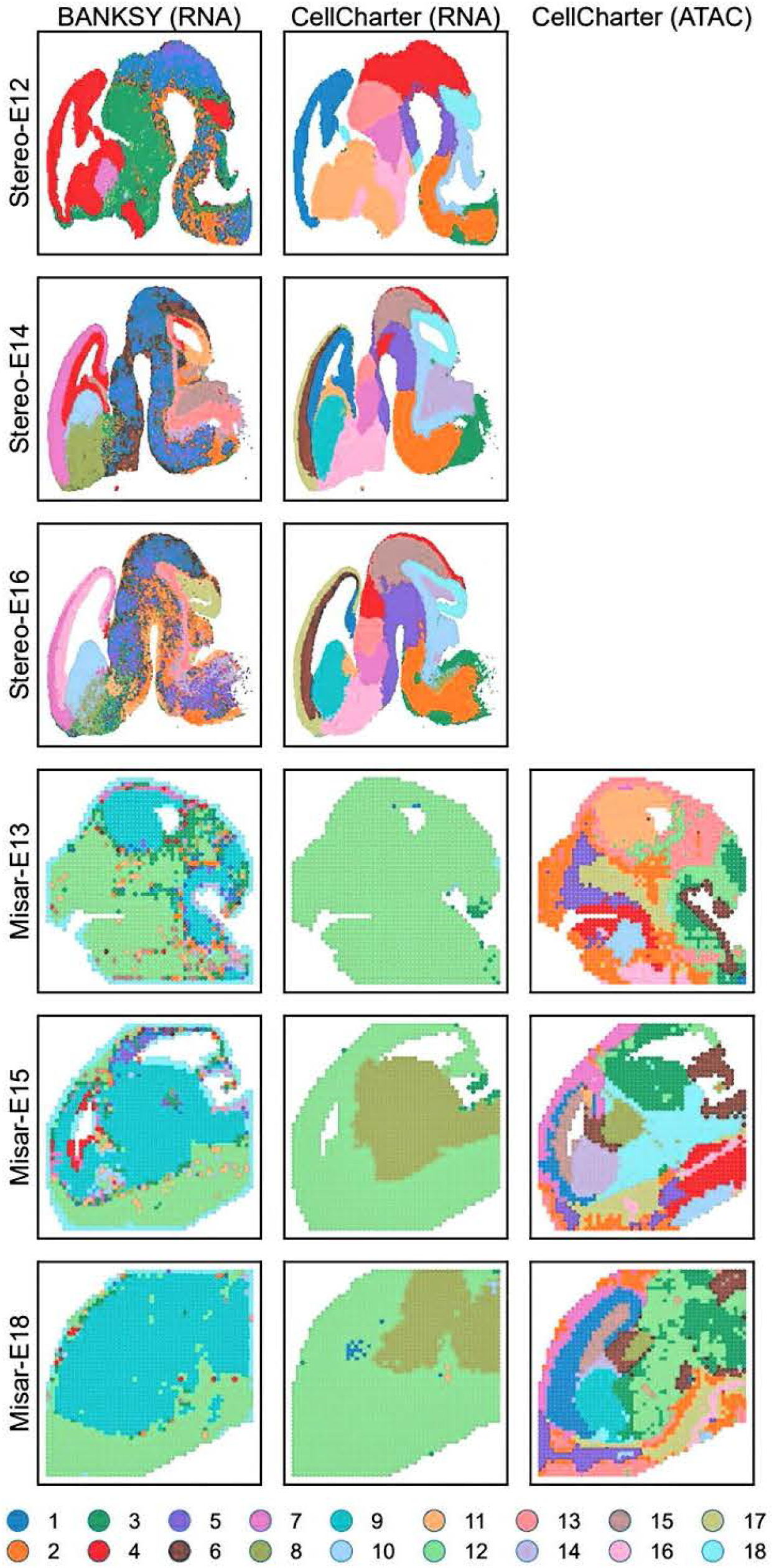
Spatial plots of clustering results from BANKSY and CellCharter on embryonic mouse brain (Misar+Stereo) dataset.

**Supp. Fig. S24.**
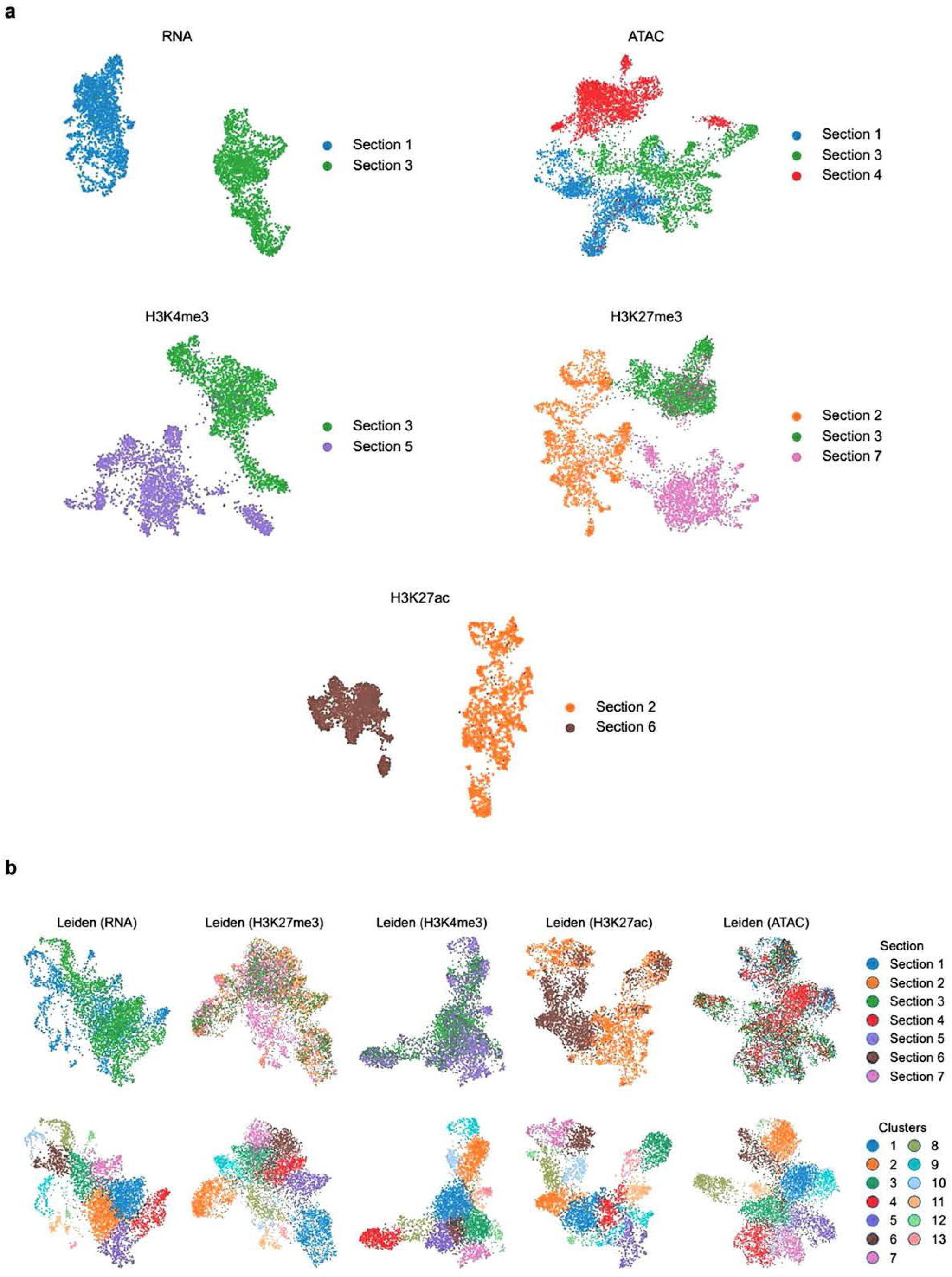
UMAP plots of mouse embryo dataset integration analysis. **a** UMAP plots of unintegrated RNA, ATAC, H3K4me3, H3K27me3, and H3K27ac histone modification profiles. Spots are colored by section labels. **b** UMAP plots of embeddings from baseline methods. Spots are colored by section labels in the first row and the second row by cluster labels.

**Supp. Fig. S25.**
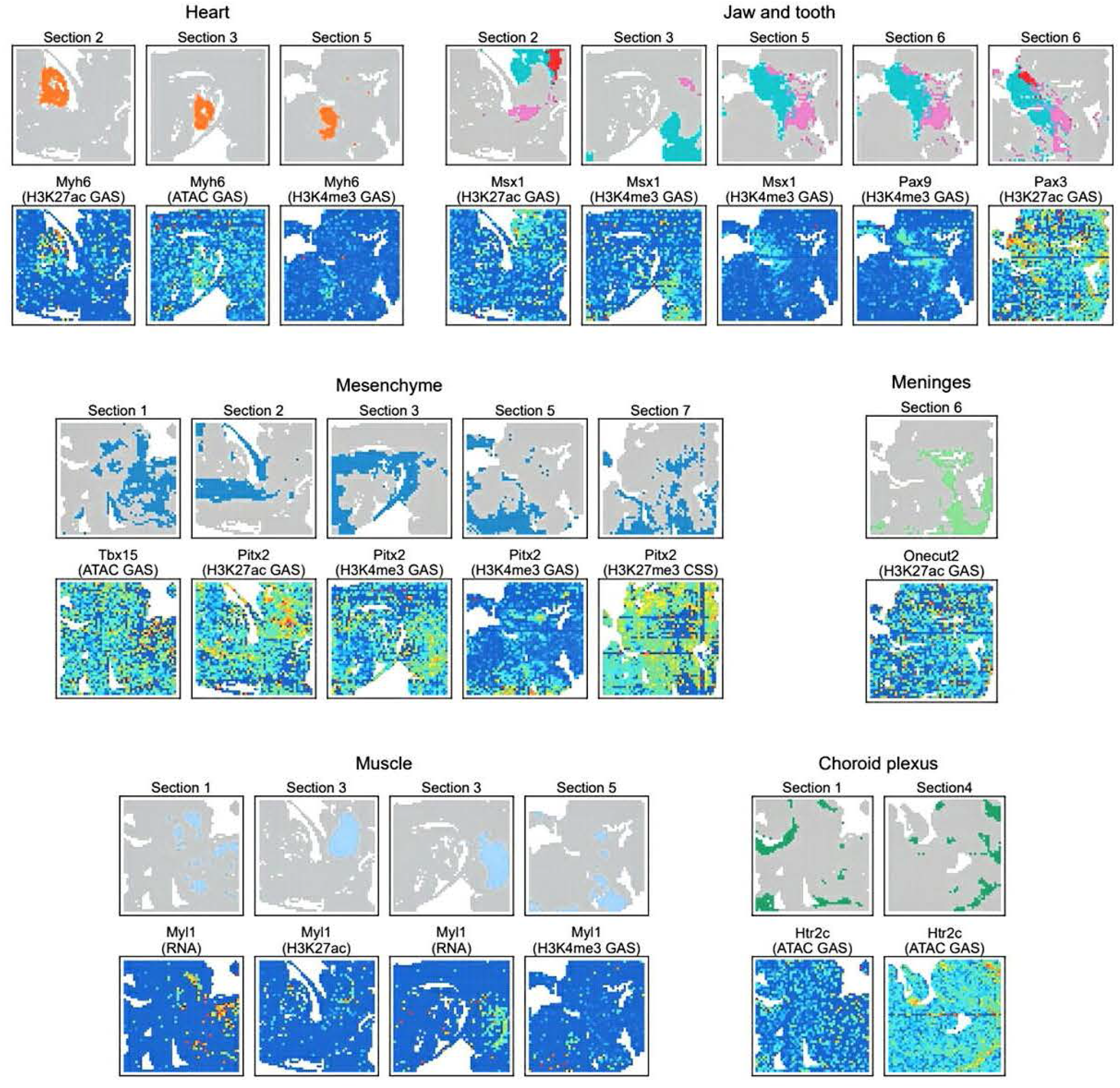
Spatial plots of selected clusters and gene activity scores/chromatin silencing scores of region-specific marker genes.

**Supp. Fig. S26.**
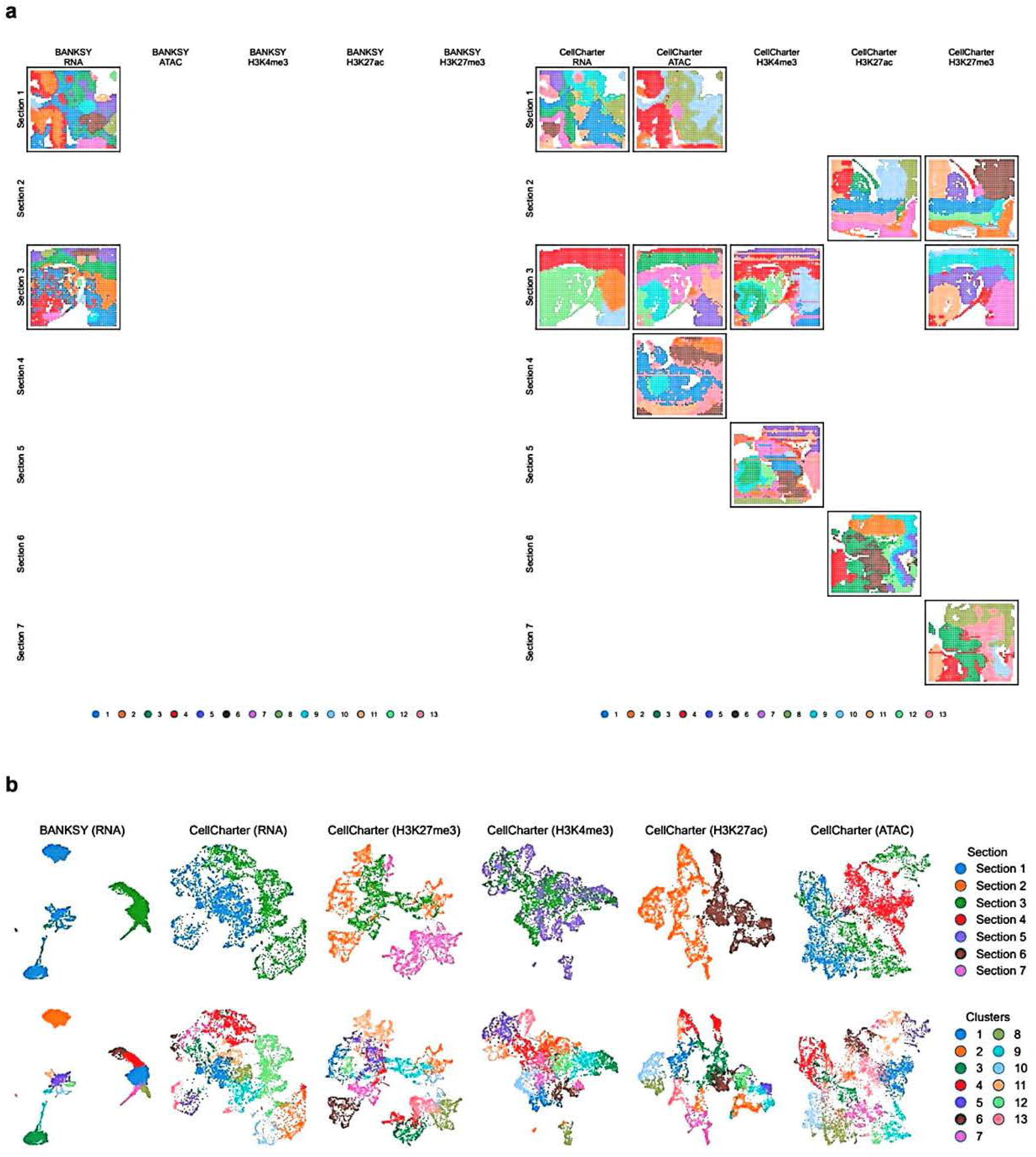
Spatial clustering plots and UMAP plots of baseline methods. **a** Spatial plots of clustering results from the baseline methods. **b** UMAP plots of embeddings from the baseline methods. Spots are colored by section labels in the first row and the second row by cluster labels.

**Supp. Fig. S27.**
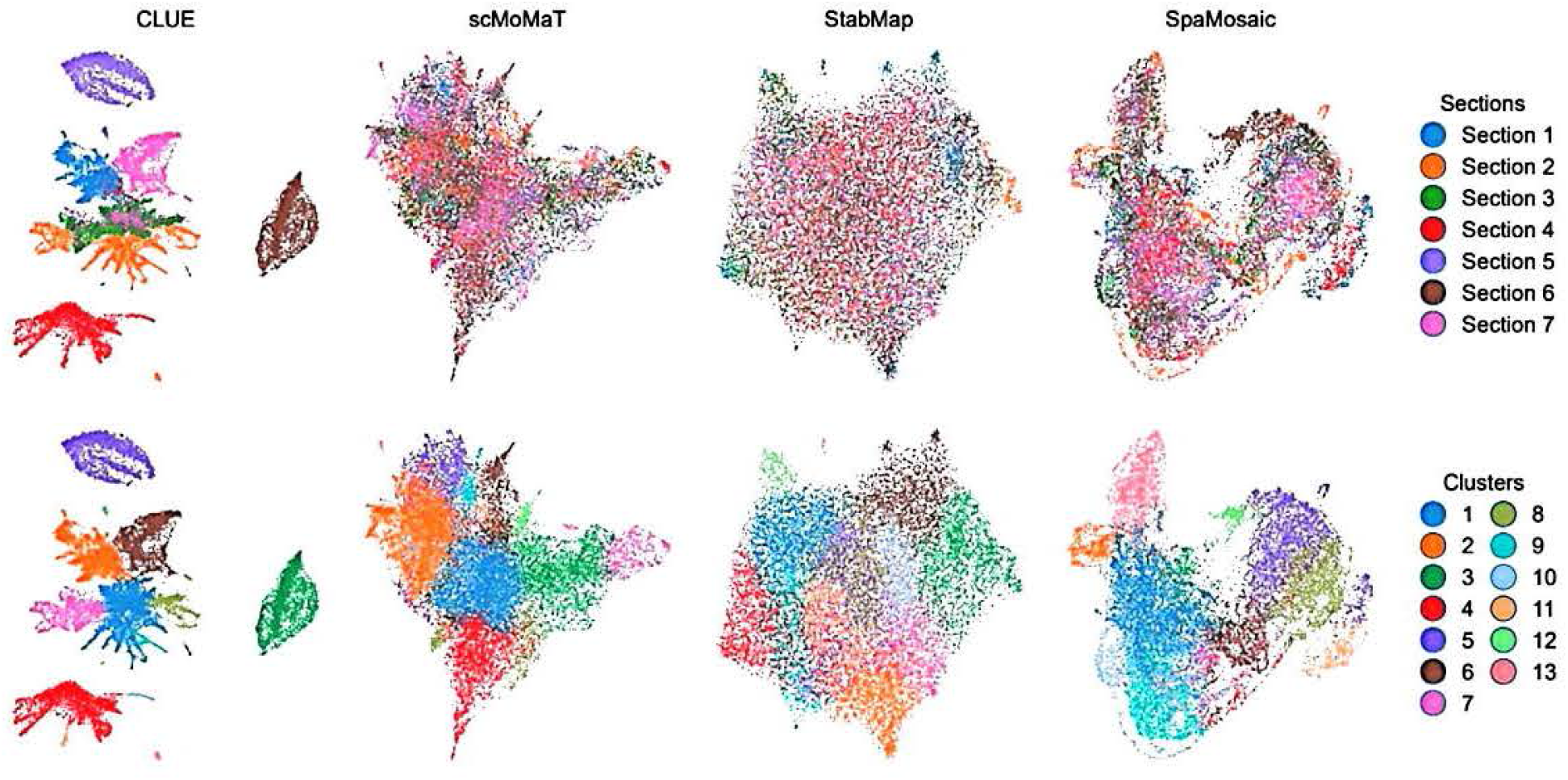
UMAP plots of embeddings from mosaic integration methods. Spots are colored by section labels in the first row and the second row by cluster labels.

**Supp. Fig. S28.**
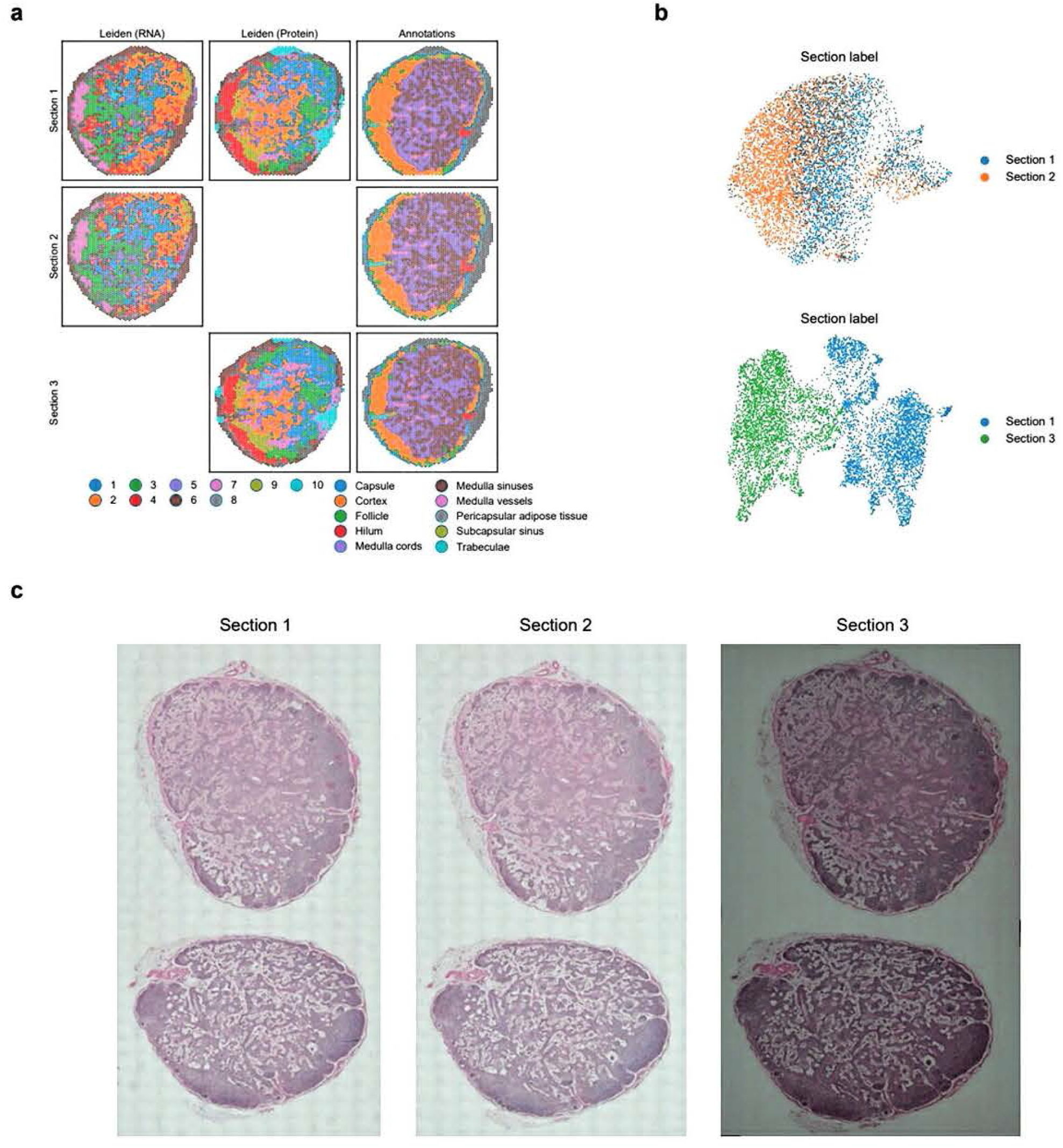
Human lymph node dataset setup and visualization. **a** Mosaic dataset configuration and clustering patterns for all three sections: section 1 with both RNA and protein expression profiles, section 2 without the protein expression modality, and section 3 without the RNA modality. Modality-specific clustering was performed jointly across relevant sections using the Leiden algorithm on batch-corrected representations. **b** UMAP plots of uncorrected RNA and protein profiles. **c** H&E staining images of the three sections.

**Supp. Fig. S29.**
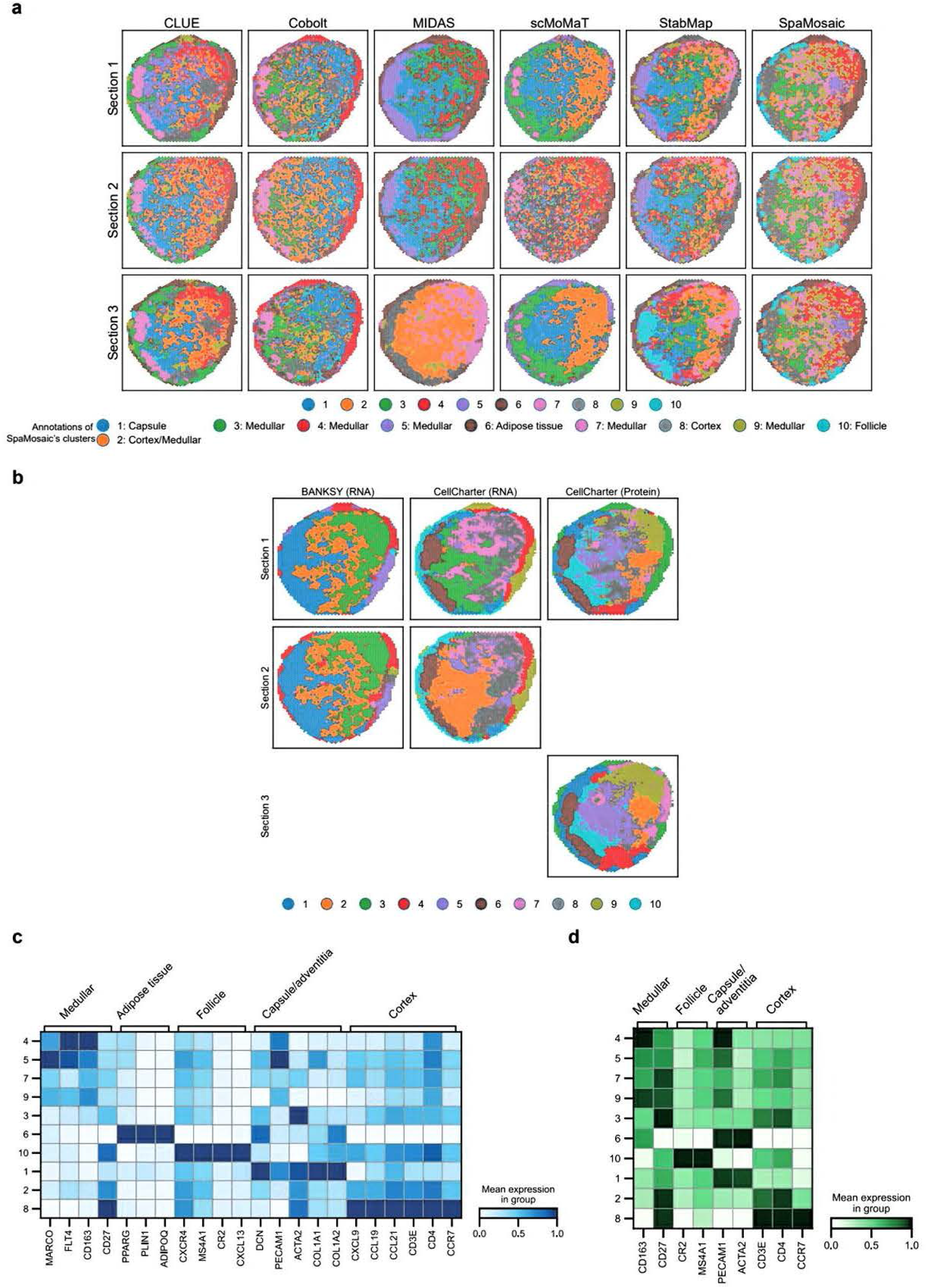
Spatial clustering plots and marker analysis on human lymph node dataset. **a** Spatial plots of clustering results from benchmarked methods. **b** Spatial plots of clustering results from **BANKSY** and CellCharter. **c** RNA expression of region-specific marker genes in SpaMosaic’s clusters. **d** Protein expression of region-specific markers in SpaMosaic’s clusters.

**Supp. Fig. S30.**
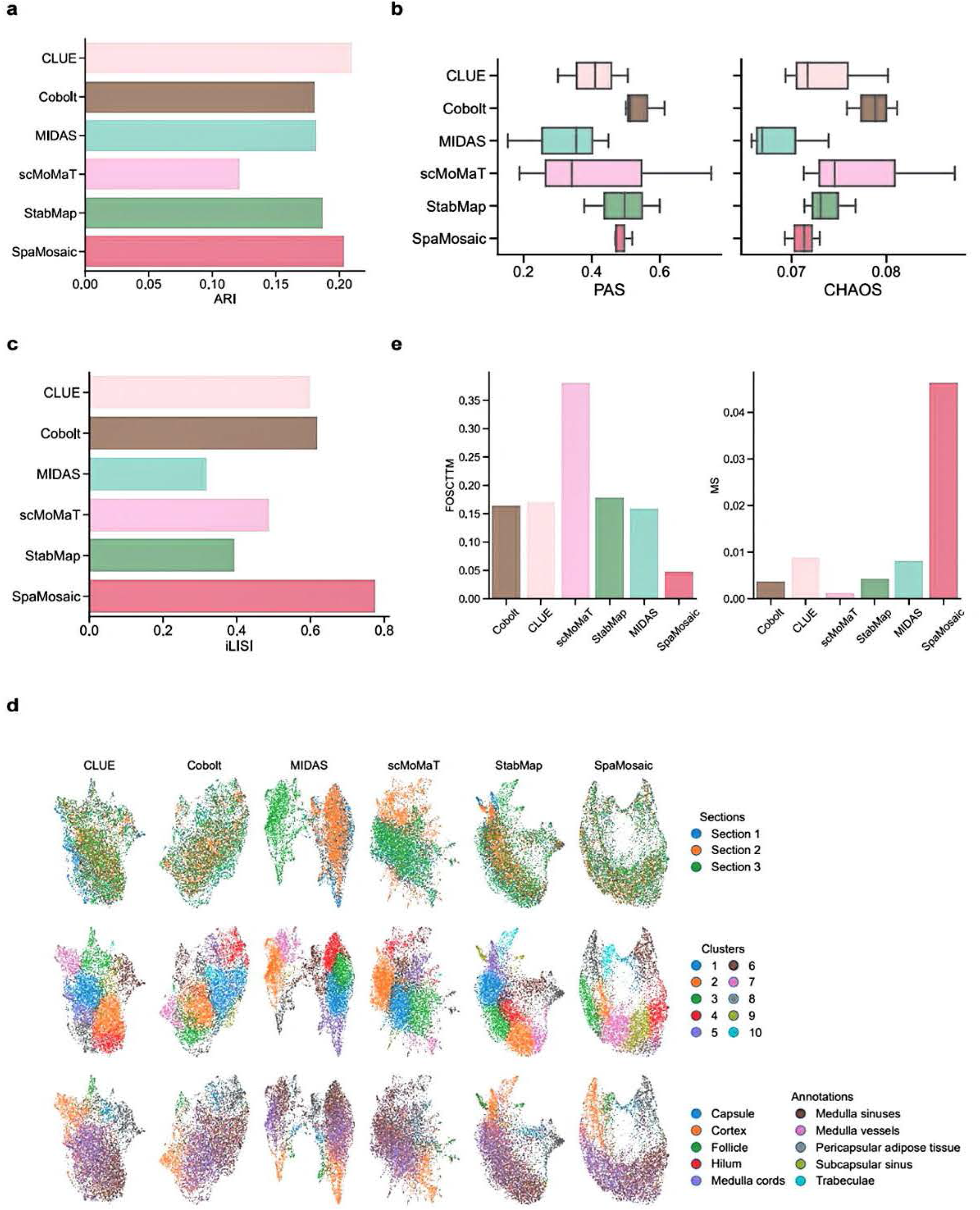
Quantitative benchmarking and UMAP plots of human lymph node dataset. **a** ARI scores of benchmarked methods. **b** PAS and CHAOS scores of benchmarked methods. **c** iLISI scores of benchmarked methods. **d** UMAP plots of embeddings from benchmarked methods. From top to bottom, spots are colored by section, cluster, and annotation, respectively. **e** FOSCTTM and MS scores of the benchmarked methods.

**Supp. Fig. S31.**
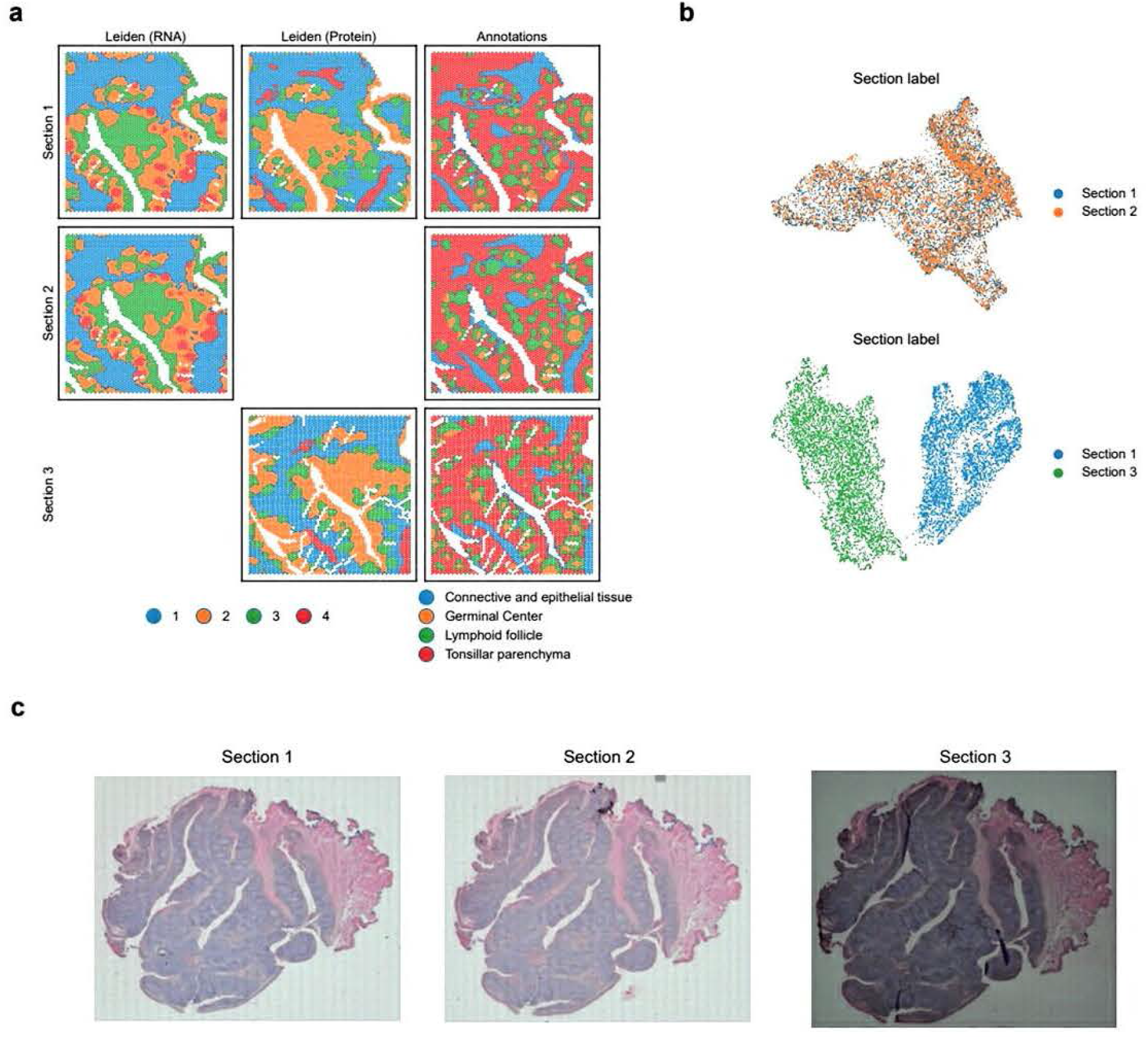
Human tonsil dataset setup and visualization. **a** Mosaic dataset configuration and clustering patterns for all three sections: section 1 with both RNA and protein expression profiles, section 2 without the protein expression modality, and section 3 without the RNA modality. Modality-specific clustering was performed jointly across relevant sections using the Leiden algorithm on batch-corrected representations. **b** UMAP plots of unintegrated RNA and protein profiles. **c** H&E staining images of the three sections.

**Supp. Fig. S32.**
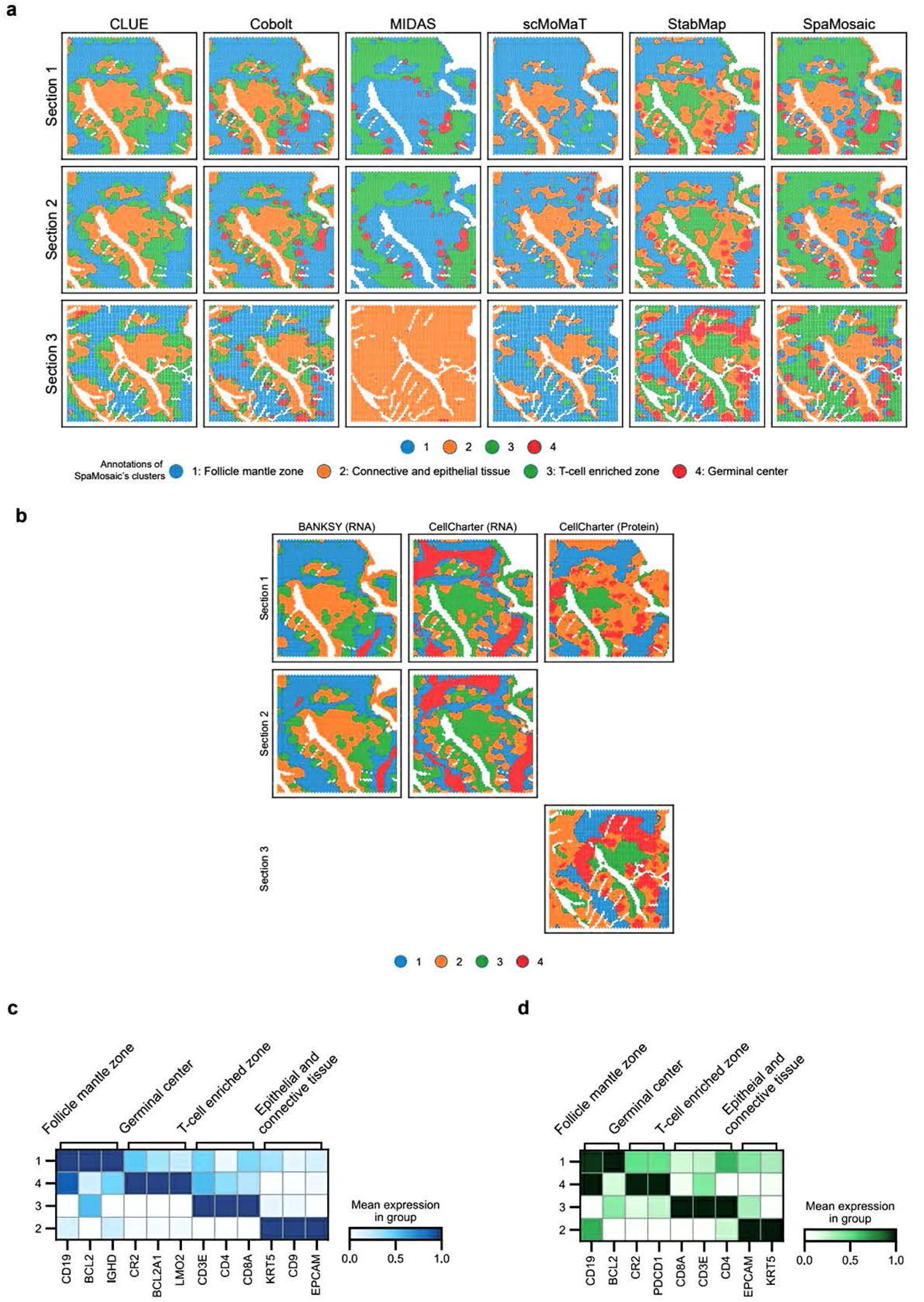
Spatial clustering plots and expression heatmap for human tonsil dataset. **a** Spatial plots of clustering results from benchmarked methods. **b** Spatial plots of clustering results from **BANKSY** and CellCharter. **c** RNA expression of region-specific marker genes in SpaMosaic’s clusters. **d** Protein expression of region-specific markers in SpaMosaic’s clusters.

**Supp. Fig. S33.**
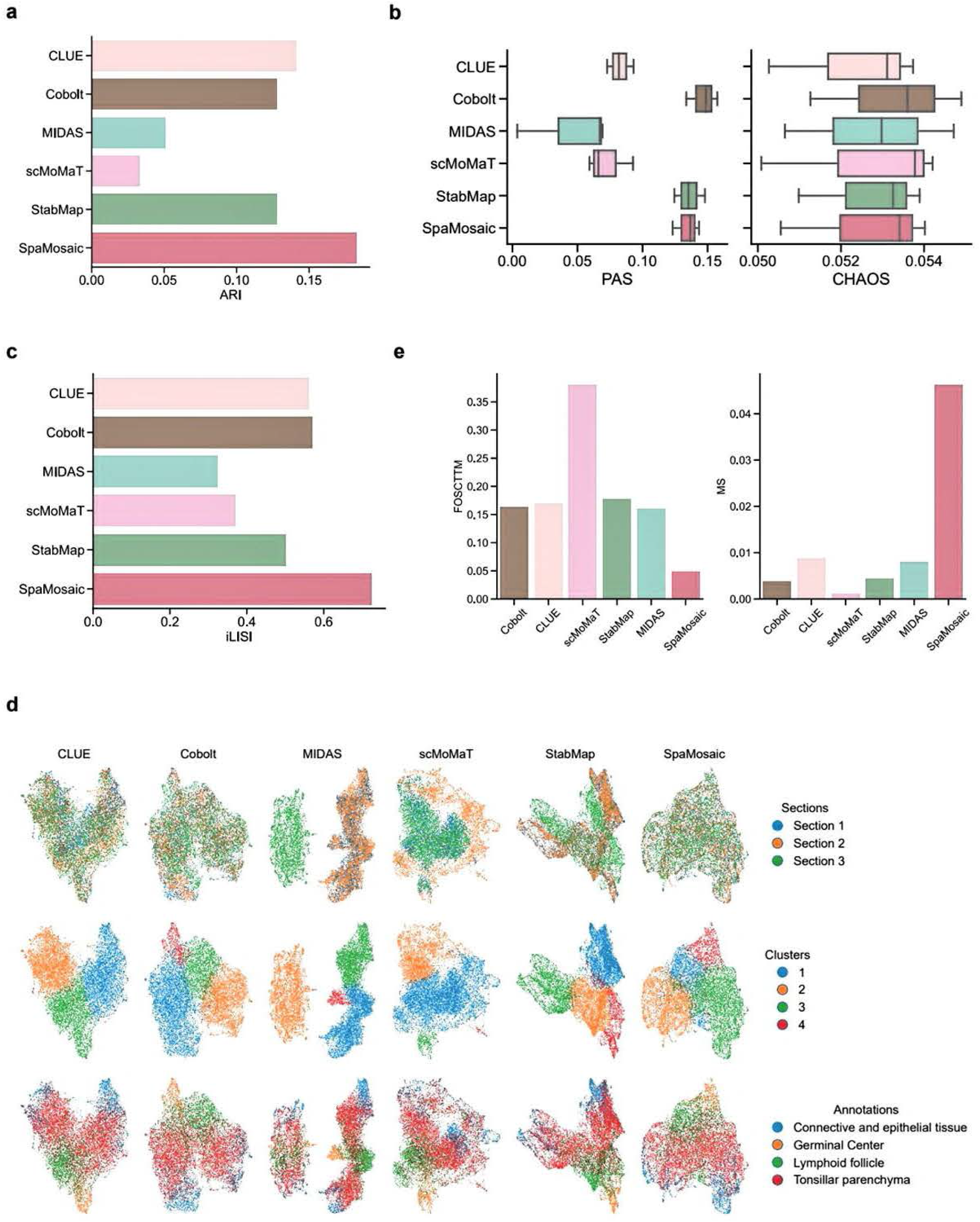
Quantitative benchmarking and UMAP plots on human tonsil dataset. **a** ARI scores of benchmarked methods. **b** PAS and CHAOS scores of benchmarked methods. **c** iLISI scores of benchmarked methods. **d** UMAP plots of embeddings from benchmarked methods. From top to bottom, spots are colored by section, cluster, and annotation, respectively. **e** FOSCTTM and MS scores of the benchmarked methods.

**Supp. Fig. S34.**
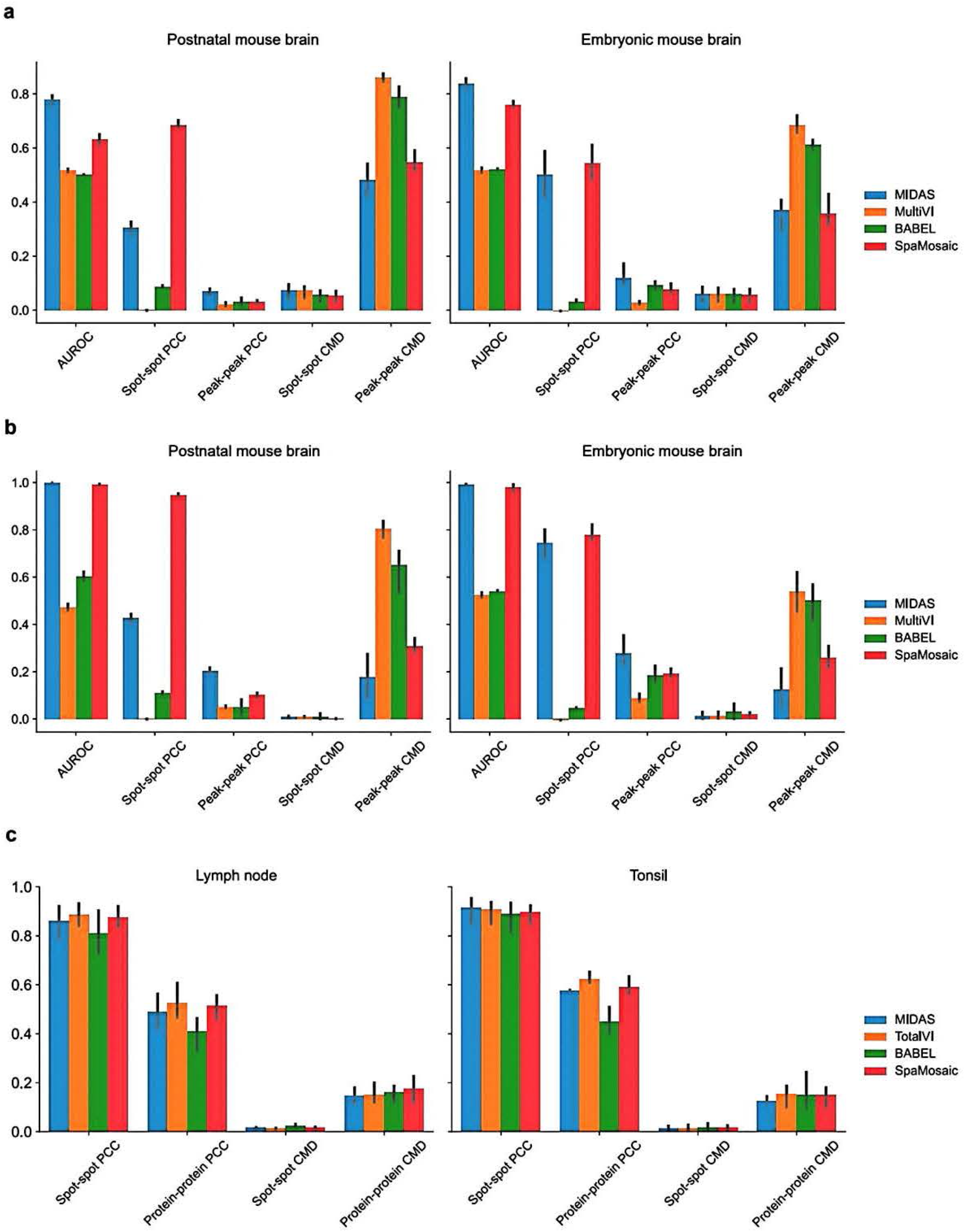
Quantitative benchmarking on imputation performance. **a** Benchmarking of ATAC imputation on the embryonic mouse brain (Misar) and postnatal mouse brain datasets. **b** Benchmarking of ATAC imputation with smoothed ground truth peak count matrices. **c** Benchmarking of protein imputation on the human lymph node and tonsil datasets.

**Supp. Fig. S35.**
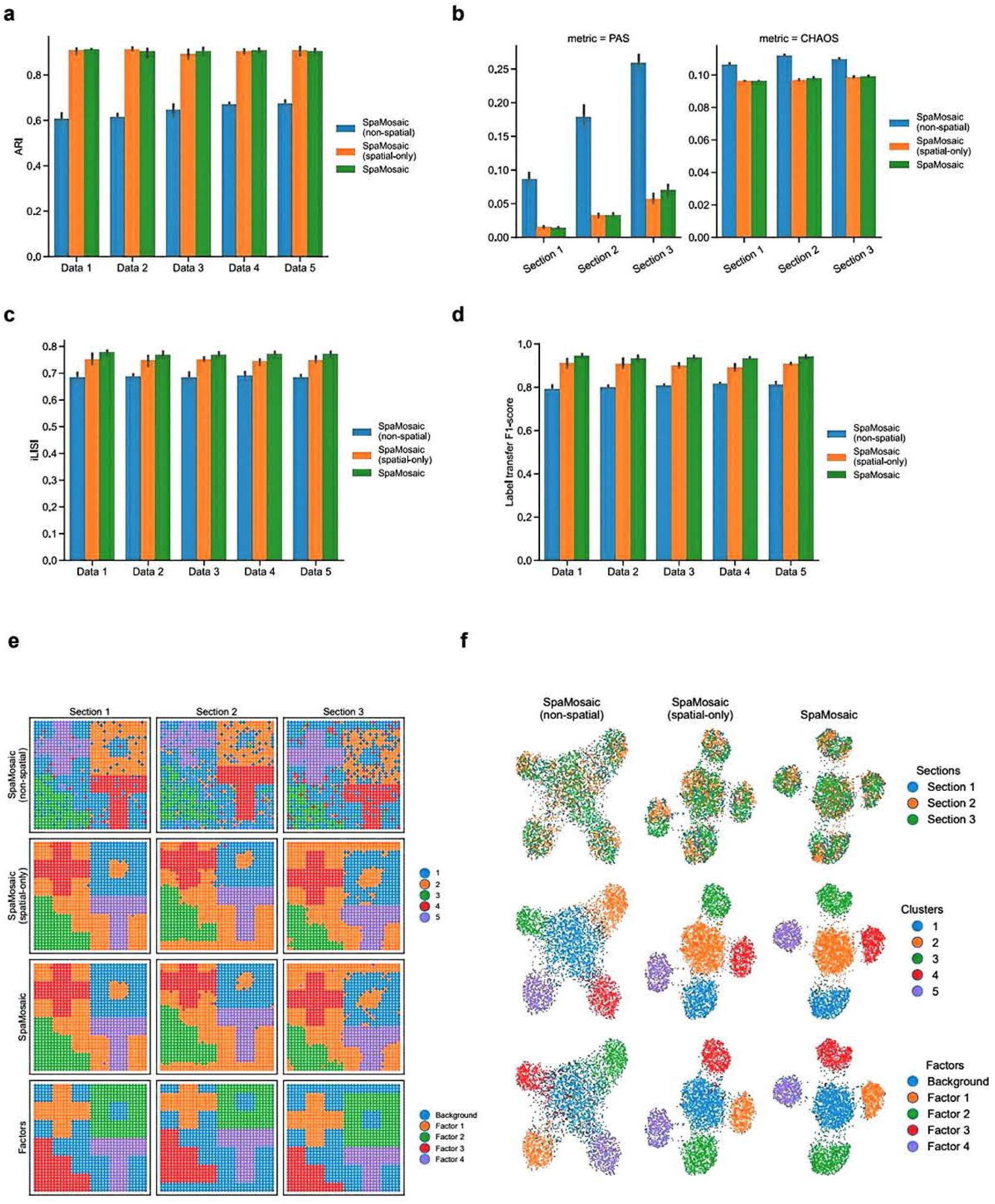
Ablation study on simulation datasets. **a** Comparison of ARI scores among the three model variants on the simulation datasets. **b** PAS and CHAOS scores of model variants. **c** iLISI scores of model variants. **d** Label transfer FI-scores of model variants. **e** Spatial plots of clustering results. The last row represents the ground truth of factor patterns. **f** UMAP plots of embeddings from the variants. From top to bottom, spots are colored by section, cluster, and factor labels.

**Supp. Fig. S36.**
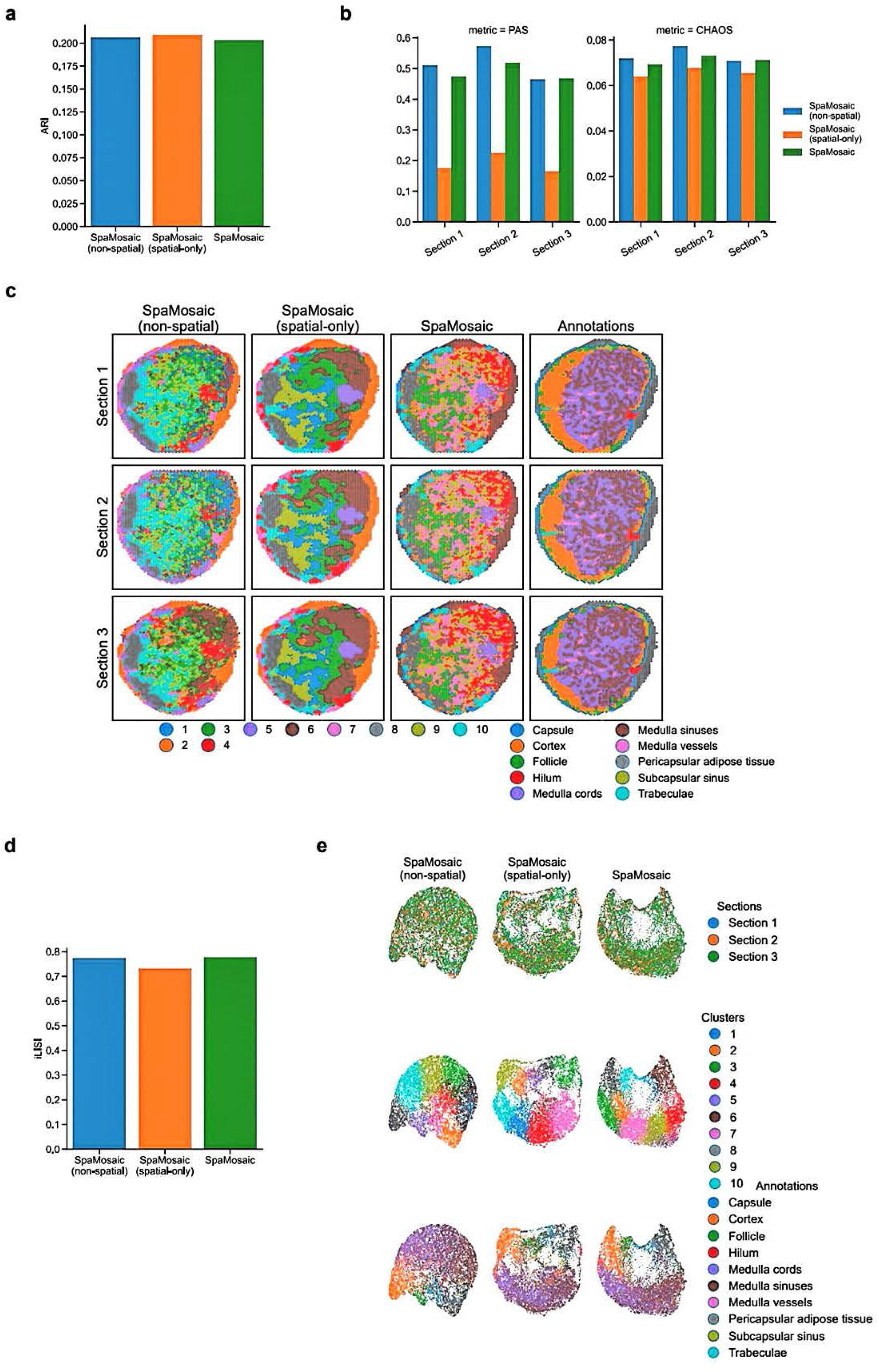
Ablation study on the human lymph node dataset. **a** Comparison of ARI scores among the three model variants on the human lymph node dataset. **b** PAS and CHAOS scores of model variants. **c** Spatial plots of clustering results. The last column depicts the manual annotation. **d** iLISI scores of model variants. **e** UMAP plots of embeddings from the variants. Spots are colored by section (top) and cluster (bottom).

**Supp. Fig. S37.**
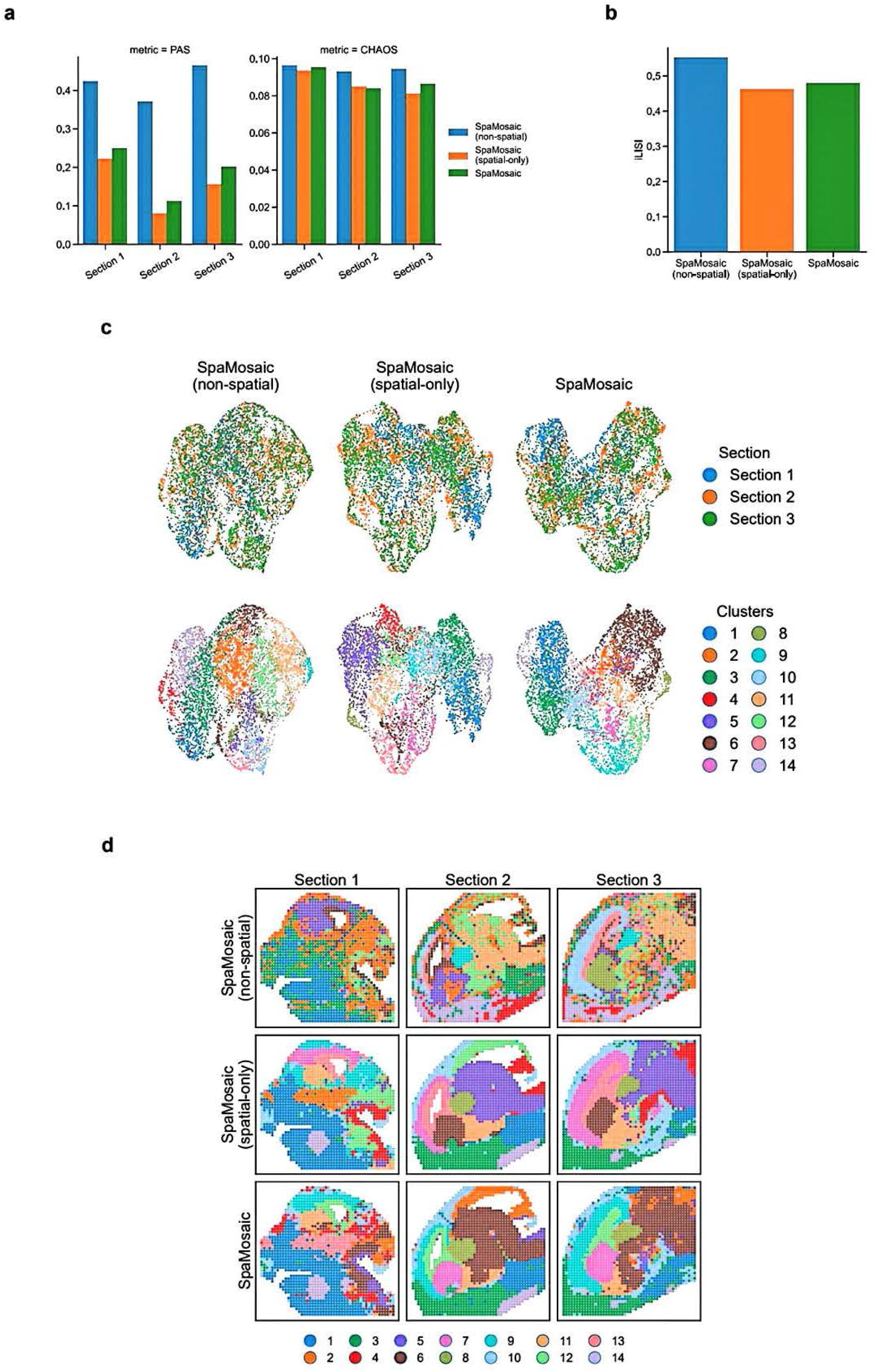
Ablation study on the embryonic mouse brain dataset (Misar). **a** PAS and CHAOS scores of the three model variants on the embryonic mouse brain dataset (Misar). **b** iLISI scores of model variants. **c** UMAP plots of embeddings from the variants. Spots are colored by section (top) and cluster (bottom). **d** Spatial plots of clustering results.

**Supp. Fig. S38.**
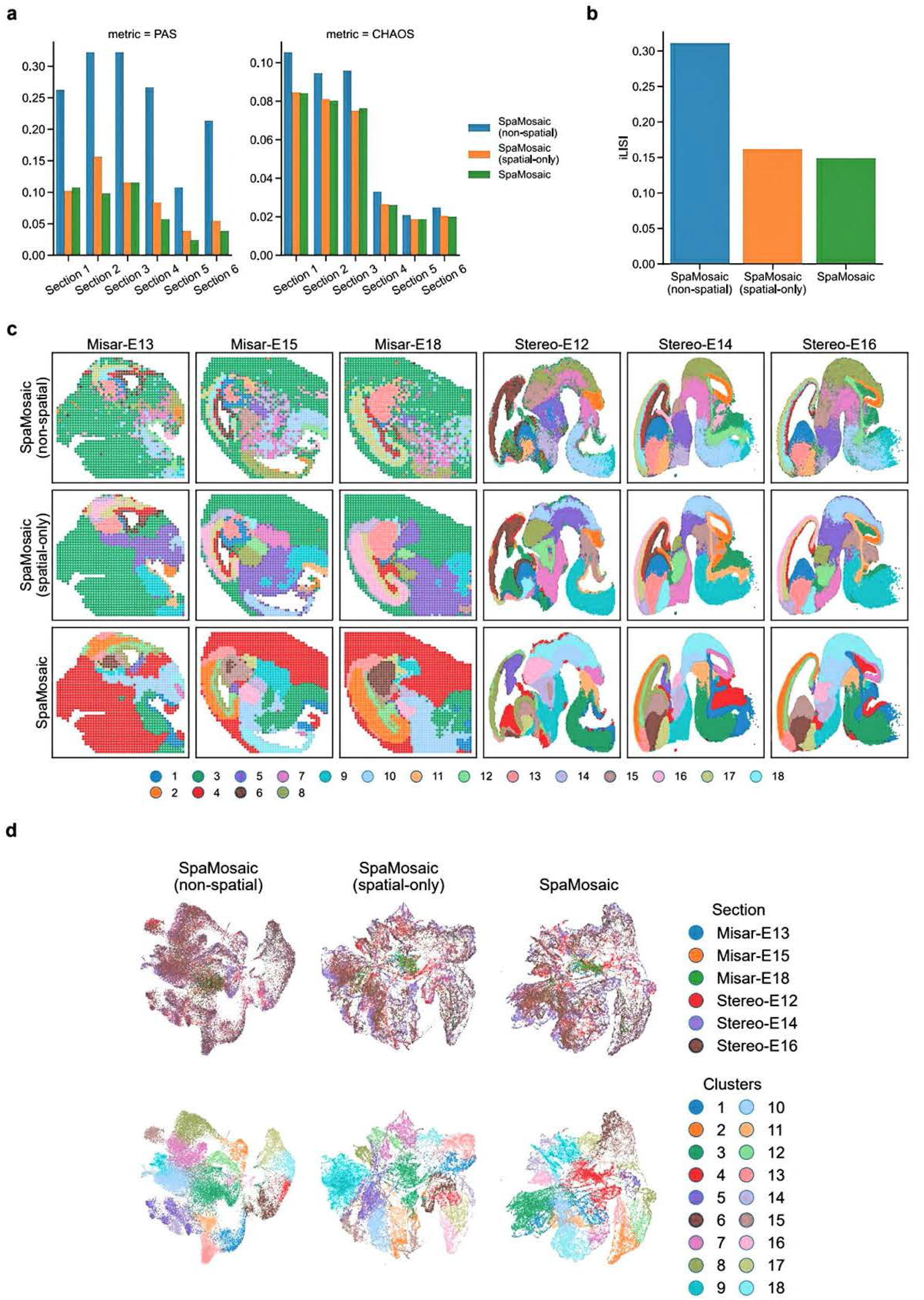
Ablation study on the embryonic mouse brain dataset (Misar+Stereo). **a** PAS and CHAOS scores of the three model variants on the embryonic mouse brain dataset (Misar+Stereo). **b** iLISI scores of model variants. **c** Spatial plots of clustering results. **d** UMAP plots of embeddings from the variants. Spots are colored by section (top) and cluster (bottom).

**Supp. Fig. S39.**
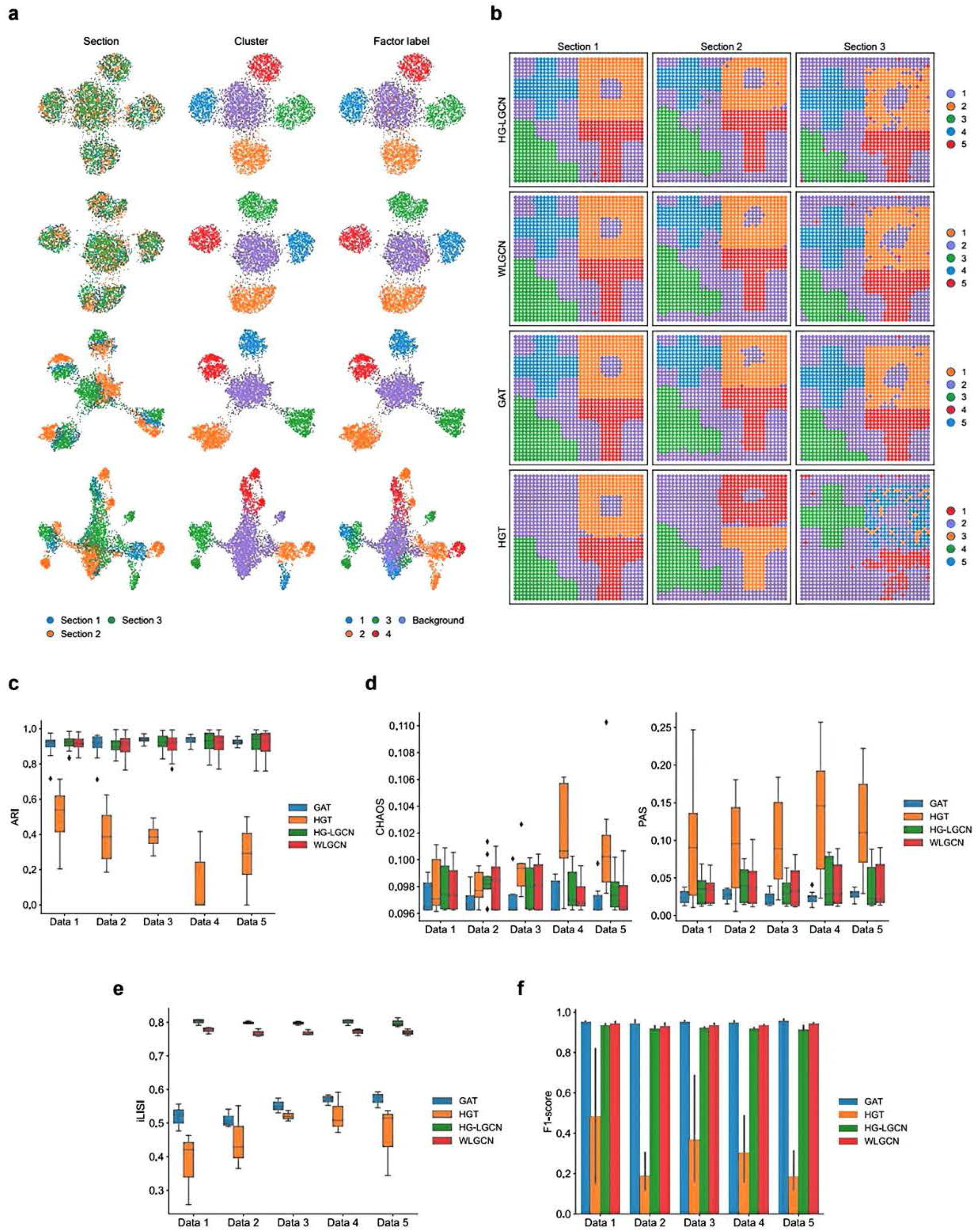
Benchmarking different graph neural network architectures. **a** UM.AP plots of embeddings generated with the different GNN architectures on the first replicate dataset of the first simulation setting. From left to right, the spots are colored by section, cluster and ground truth factor labels. **b** Spatial plots of clusters obtained with the different GNN architectures. The spot colors are matched with the cluster colors in **a. c** Box plots showing the ARI scores with different GNN architectures. In the box plot, the center lines indicate the median, boxes indicate the interquartile range, and whiskers indicate l.5x interquartile range. **d** PAS and CHAOS scores with different GNN architectures. **e** iLISI scores with different GNN architectures. **f** Label transfer Fl-scores with different GNN architectures.

**Supp. Fig. S40.**
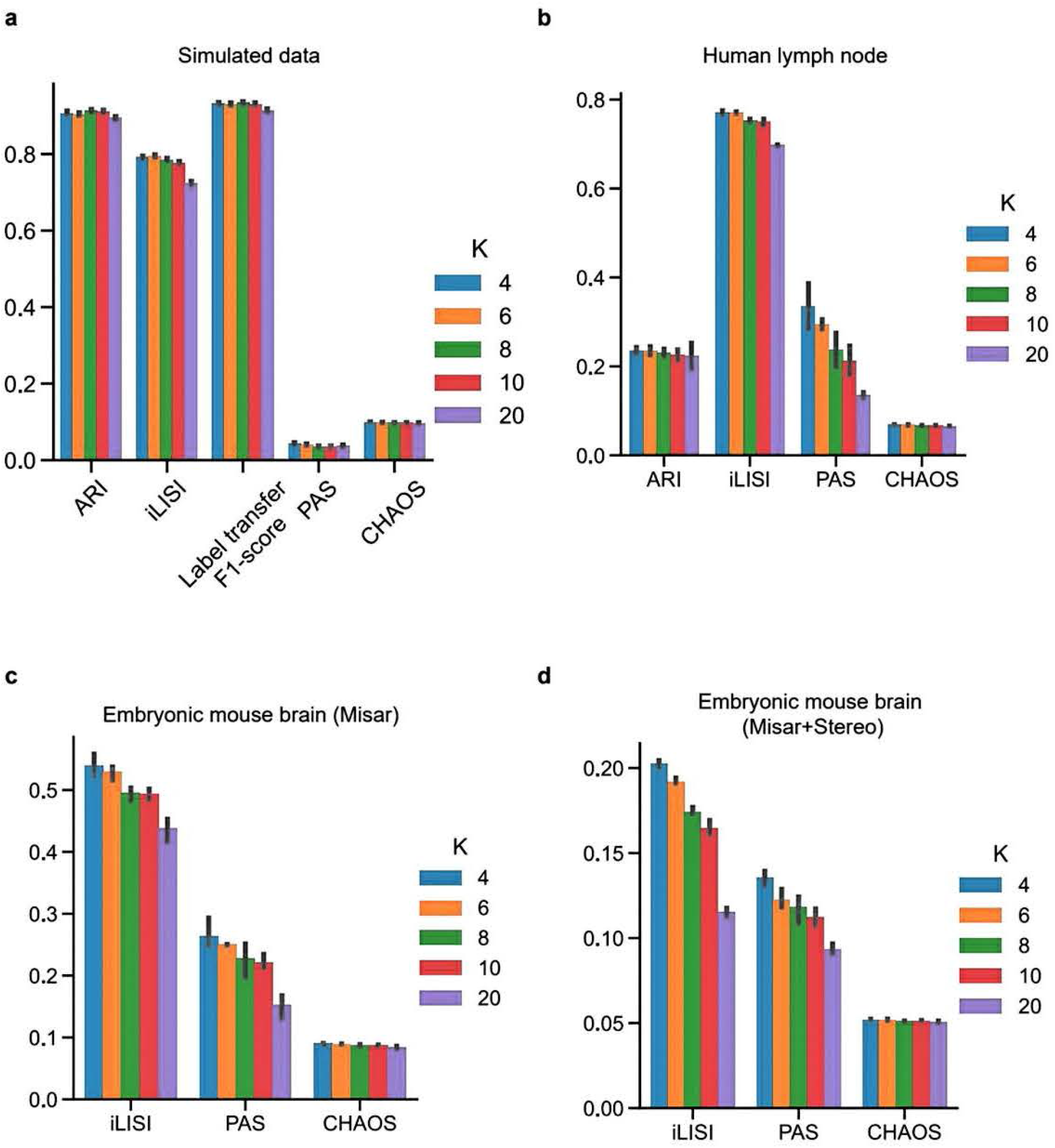
Quantitative evaluation of parameter k sensitivity in spatial adjacency graph construction. **a** Benchmarking metrics computed on the simulation datasets, ARI, iLISI, label transfer Fl-scores, PAS and CHAOS. **b** Benchmarking metrics computed on the human lymph node dataset. **c** Benchmarking metrics computed on the embryonic mouse brain dataset (Misar). **d** Benchmarking metrics computed on the embryonic mouse brain dataset (Misar+Stereo).

**Supp. Fig. S41.**
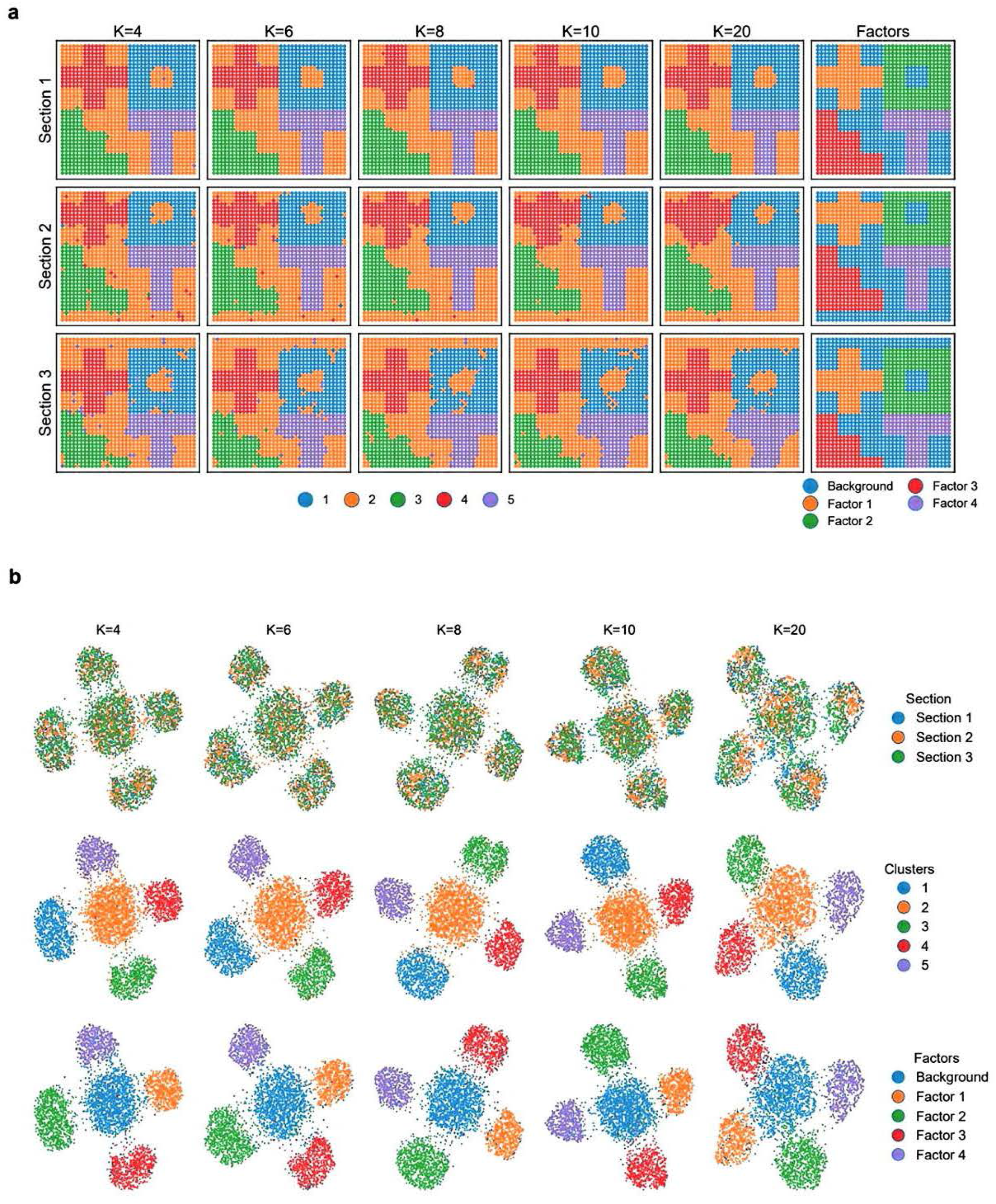
Visualization of parameter k impact on the first replicate of simulated datasets. **a** Spatial plots of clustering results from SpaMosaic with different k values. The rightmost column shows the ground truth of factor patterns. **b** UMAP plots of embeddings from SpaMosaic with different k values. From top to bottom, spots are colored by section, cluster and factor labels.

**Supp. Fig. S42.**
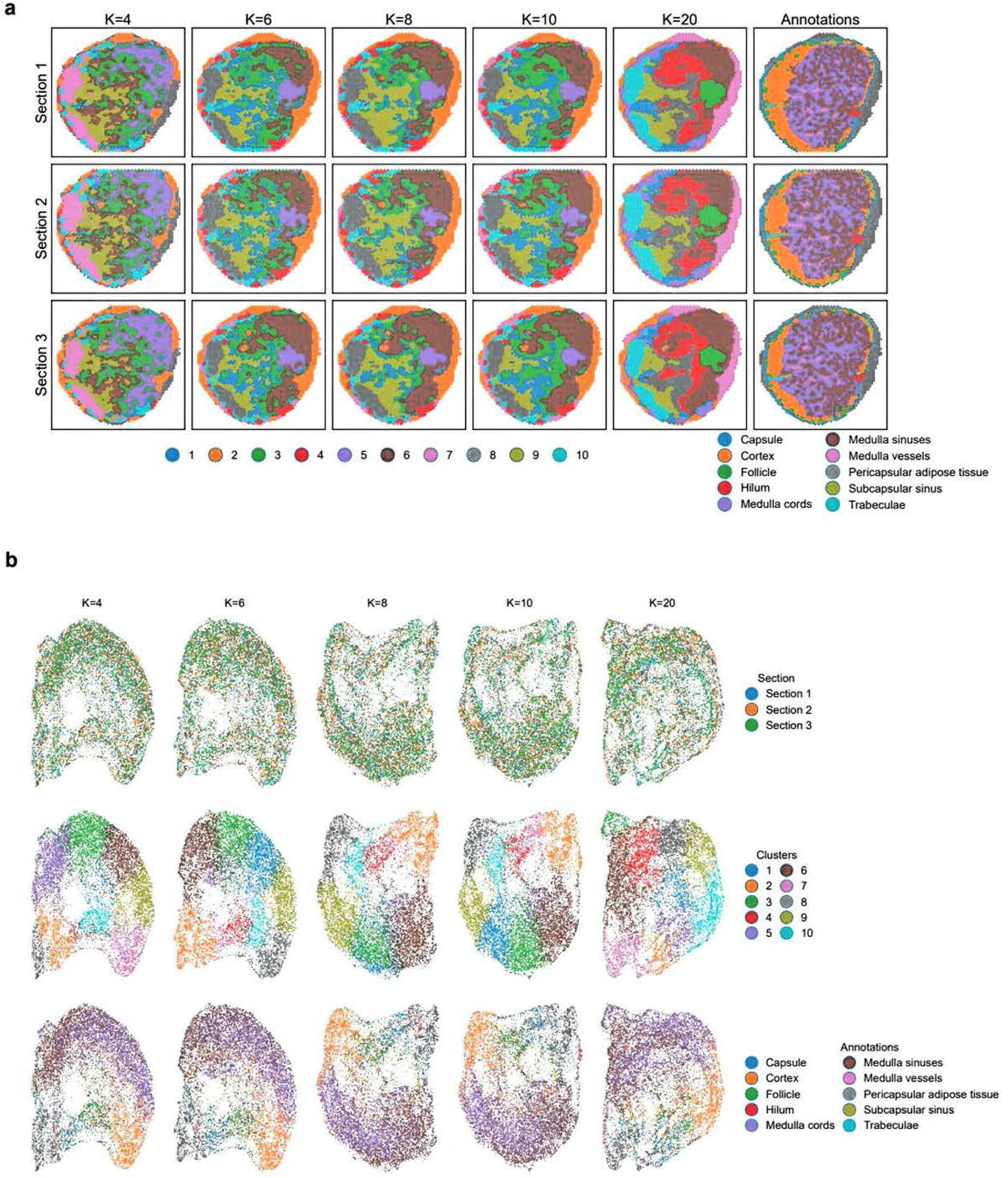
Visualization of parameter k impact on the human lymph node dataset. **a** Spatial plots of clustering results from SpaMosaic with different k values. The rightmost column shows the expert manual annotations. **b** UMAP plots of embeddings from SpaMosaic with different k values. From top to bottom, spots are colored by section, cluster and annotation labels.

**Supp. Fig. S43.**
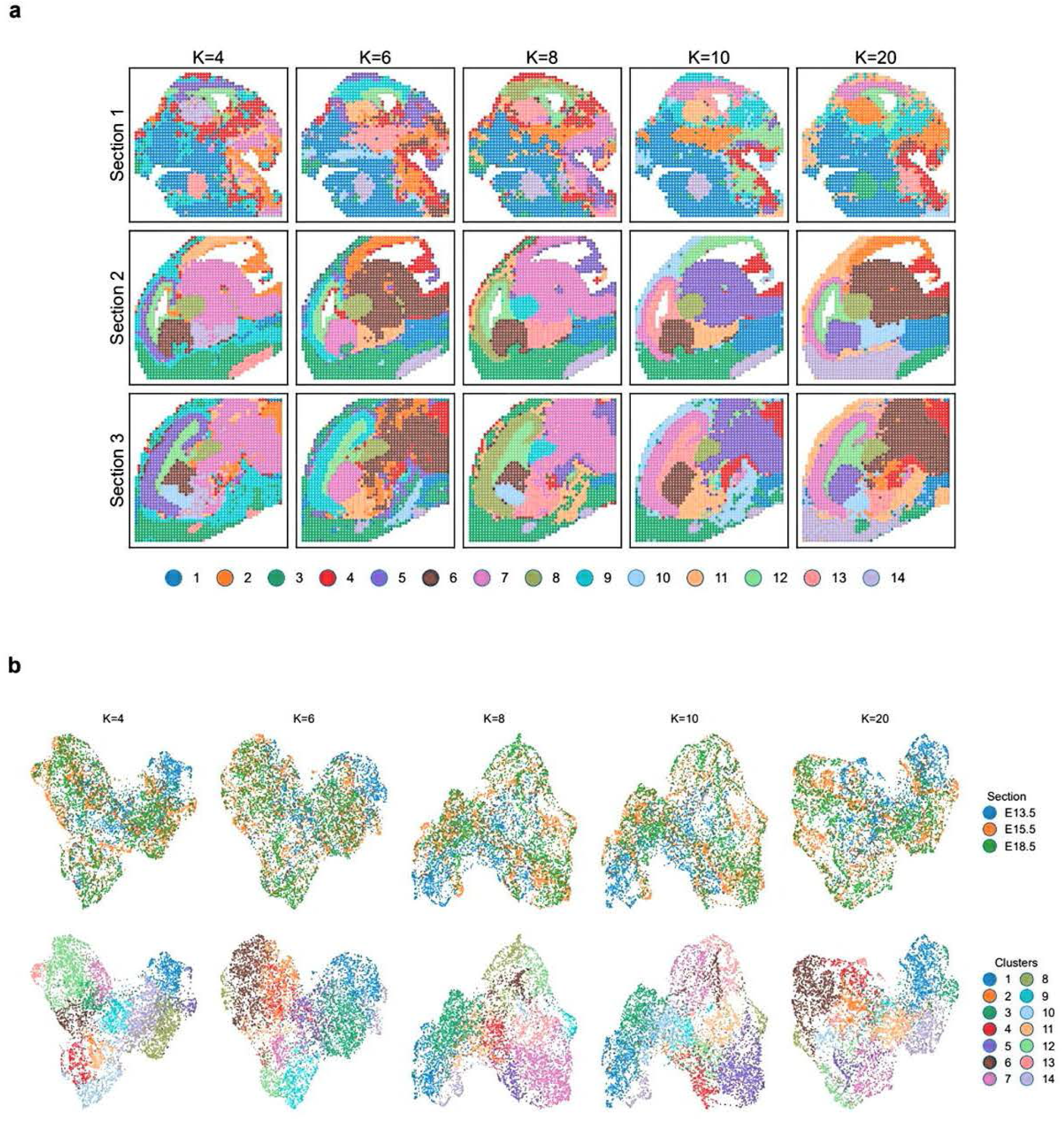
Visualization of parameter k impact on the embryonic dataset (Misar). **a** Spatial plots of clustering results from SpaMosaic with different k values. **b** UMAP plots of embeddings from SpaMosaic with different k values. From top to bottom, spots are colored by section and cluster labels.

**Supp. Fig. S44.**
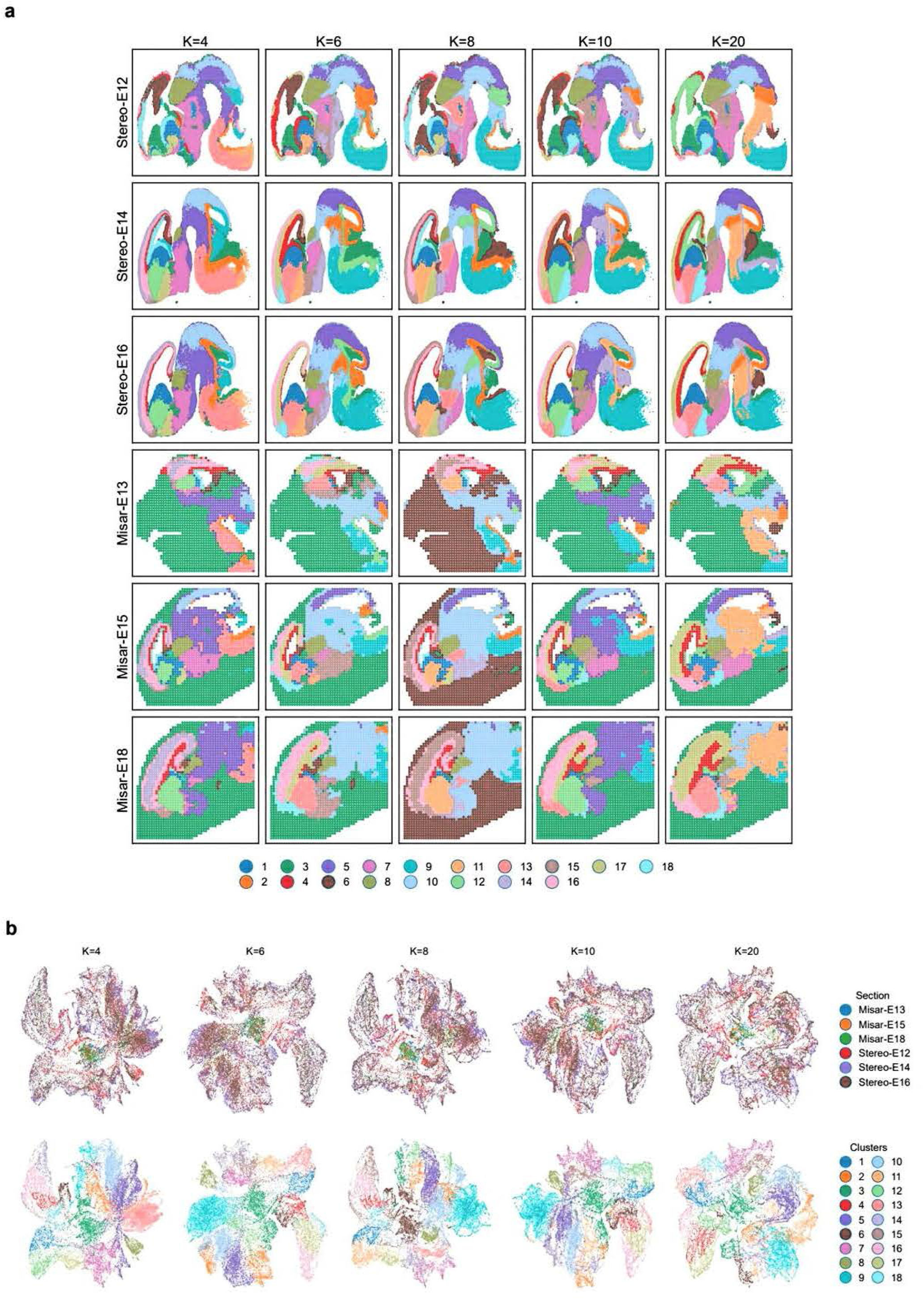
Visualization of parameter k impact on the embryonic dataset (Misar+Stereo). **a** Spatial plots of clustering results from SpaMosaic with different k values. **b** UMAP plots of embeddings from SpaMosaic with different k values. From top to bottom, spots are colored by section and cluster labels.

**Supp. Fig. S45.**
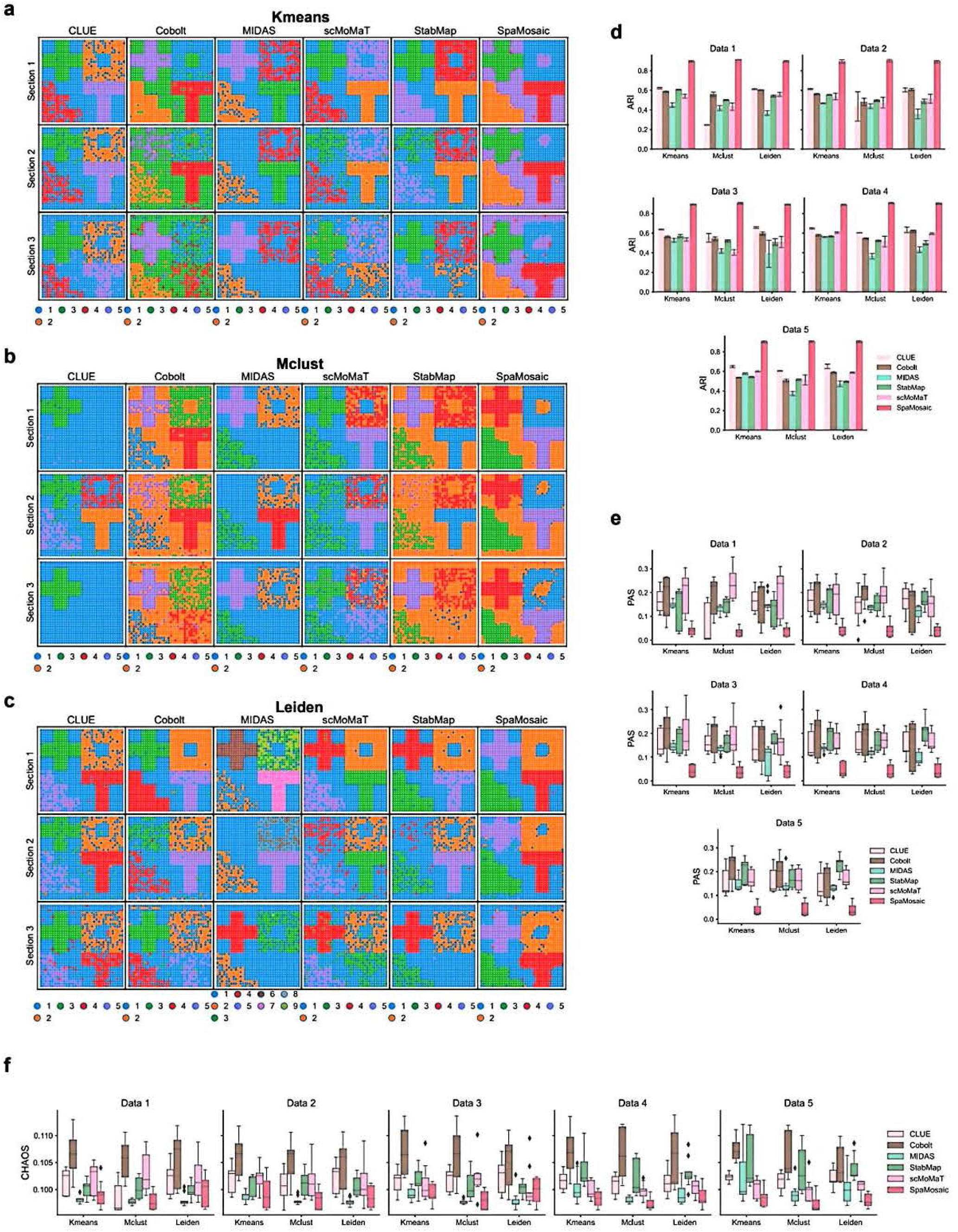
Benchmarking of mosaic integration methods using different clustering algorithms on simulated datasets. **a-c** Spatial plots of clustering results from benchmarked mosaic integration methods using Kmeans **(a)**, Mclust **(b)**, and Leiden **(c)** for clustering. **d-f** Performance comparison using ARI **(d)**, PAS **(e)** and CHAOS **(f)** metrics.

**Supp. Fig. S46.**
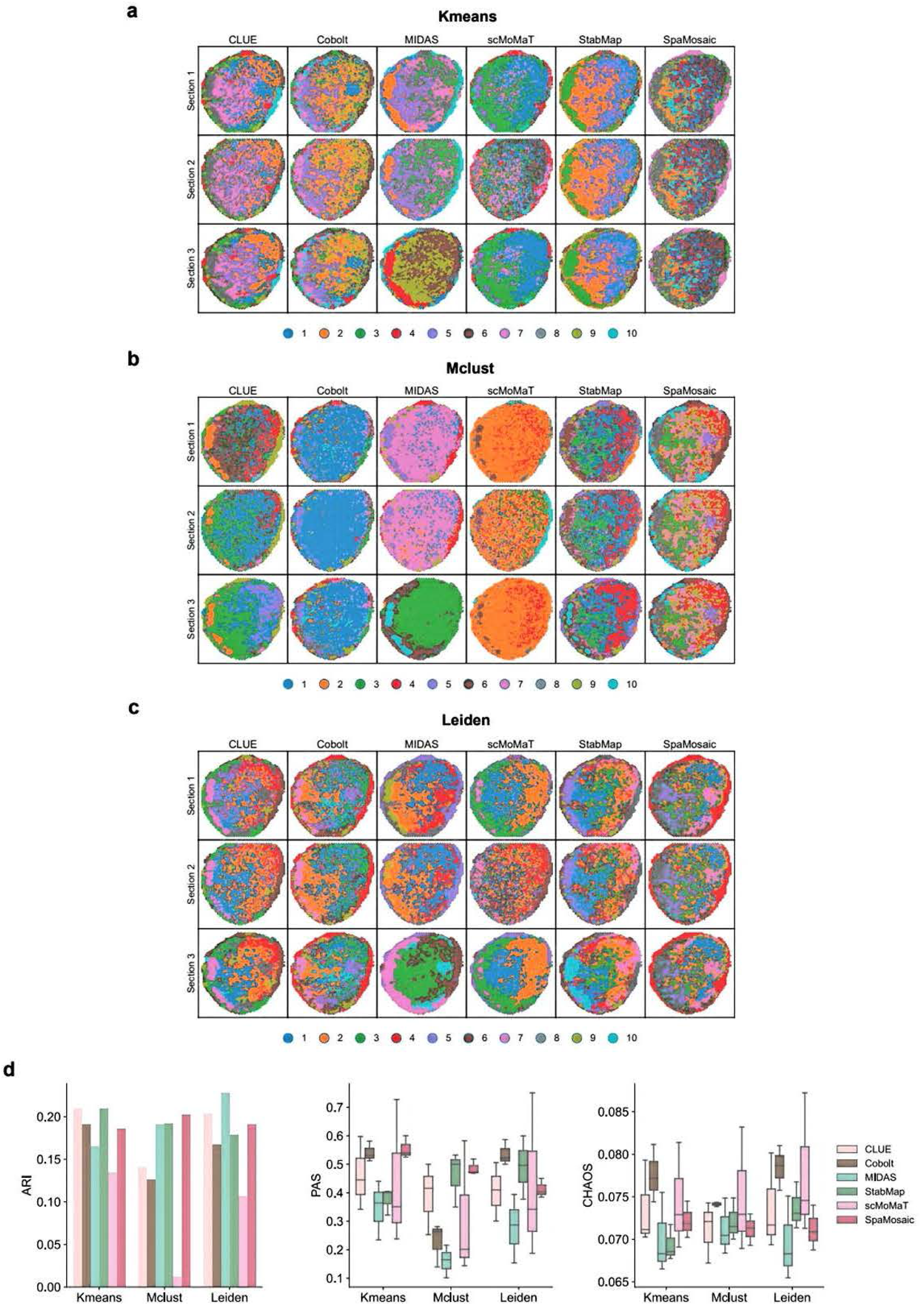
Benchmarking of mosaic integration methods using different clustering algorithms on the human lymph node dataset. **a-c** Spatial plots of clustering results from benchmarked mosaic integration methods using Kmeans **(a)**, Mclust **(b)**, and Leiden (c) for clustering. **d** Performance comparison using ARI, PAS and CHAOS metrics.

**Supp. Fig. S47.**
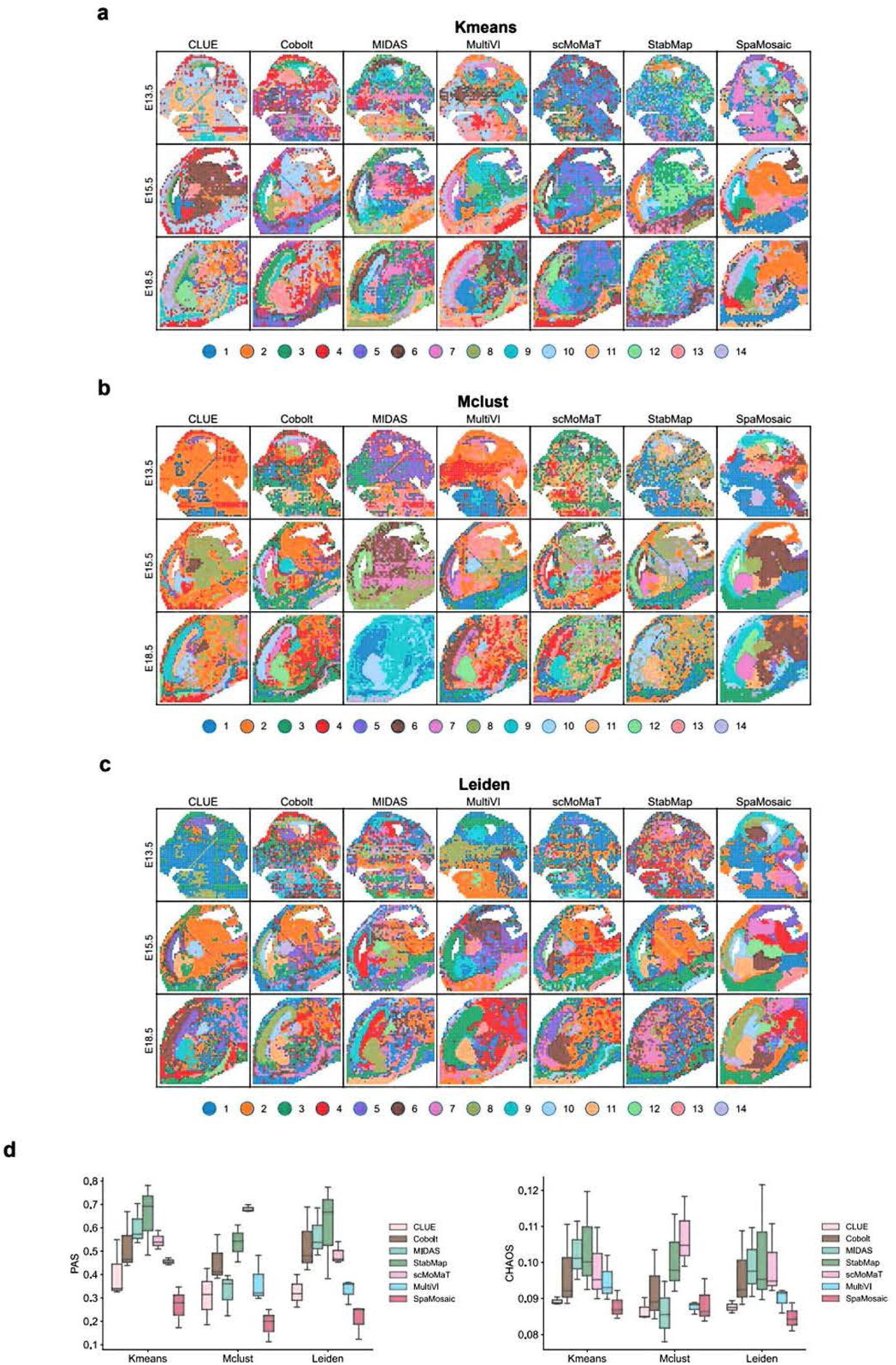
Benchmarking of mosaic integration methods using different clustering algorithms on the embryonic mouse brain dataset (Misar). **a-c** Spatial plots of clustering results from benchmarked mosaic integration methods using Kmeans **(a)**, Mclust **(b)**, and Leiden **(c)** for clustering. **d** Performance comparison using PAS and CHAOS metrics.

**Supp. Fig. S48.**
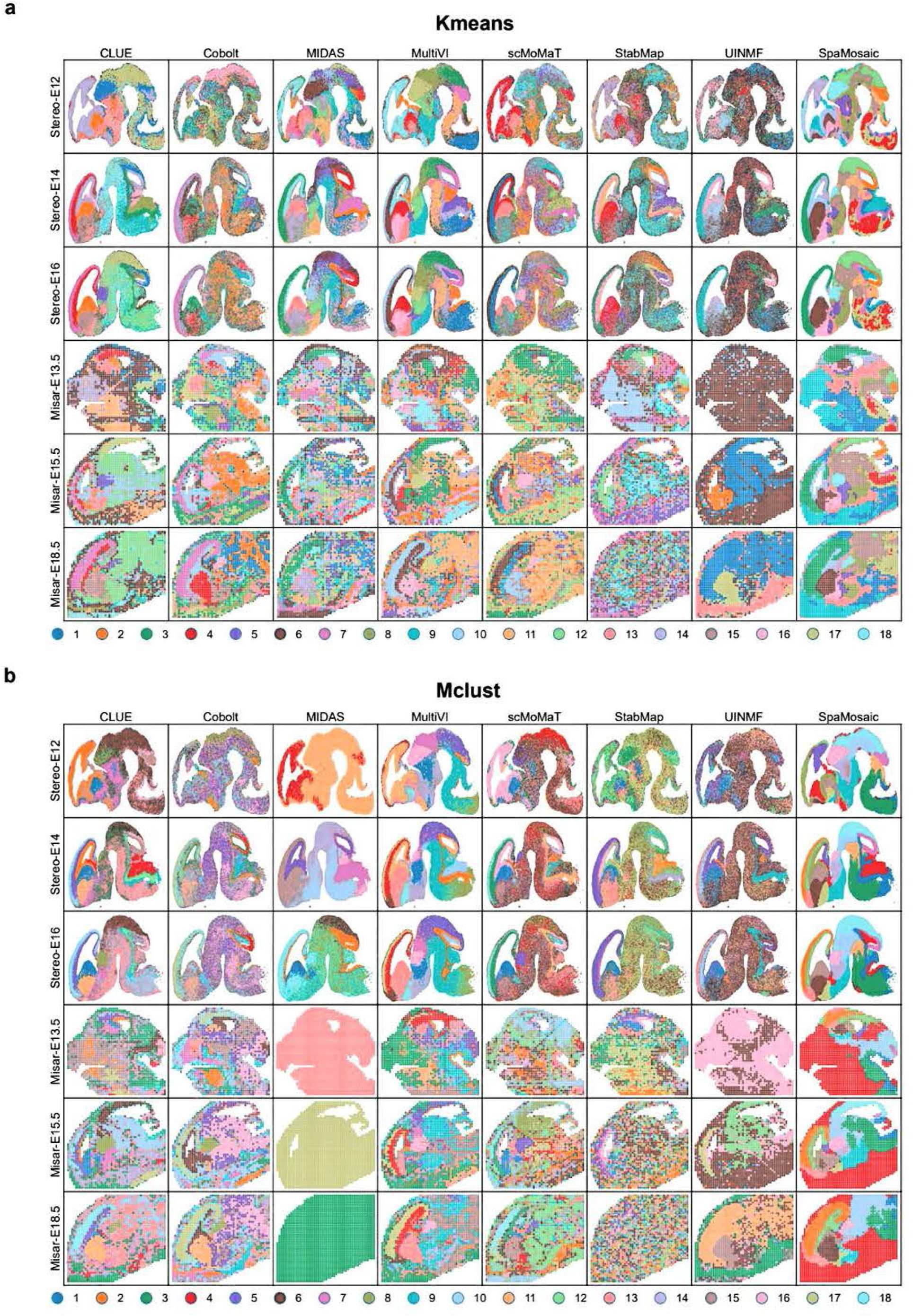
Benchmarking of mosaic integration methods using different clustering algorithms on the embryonic mouse brain dataset (Misar+Stereo). **a-b** Spatial plotsof clustering results from all compared mosaic integration methods using Kmeans **(a)** and Mclust **(b)** for clustering.

**Supp. Fig. S49.**
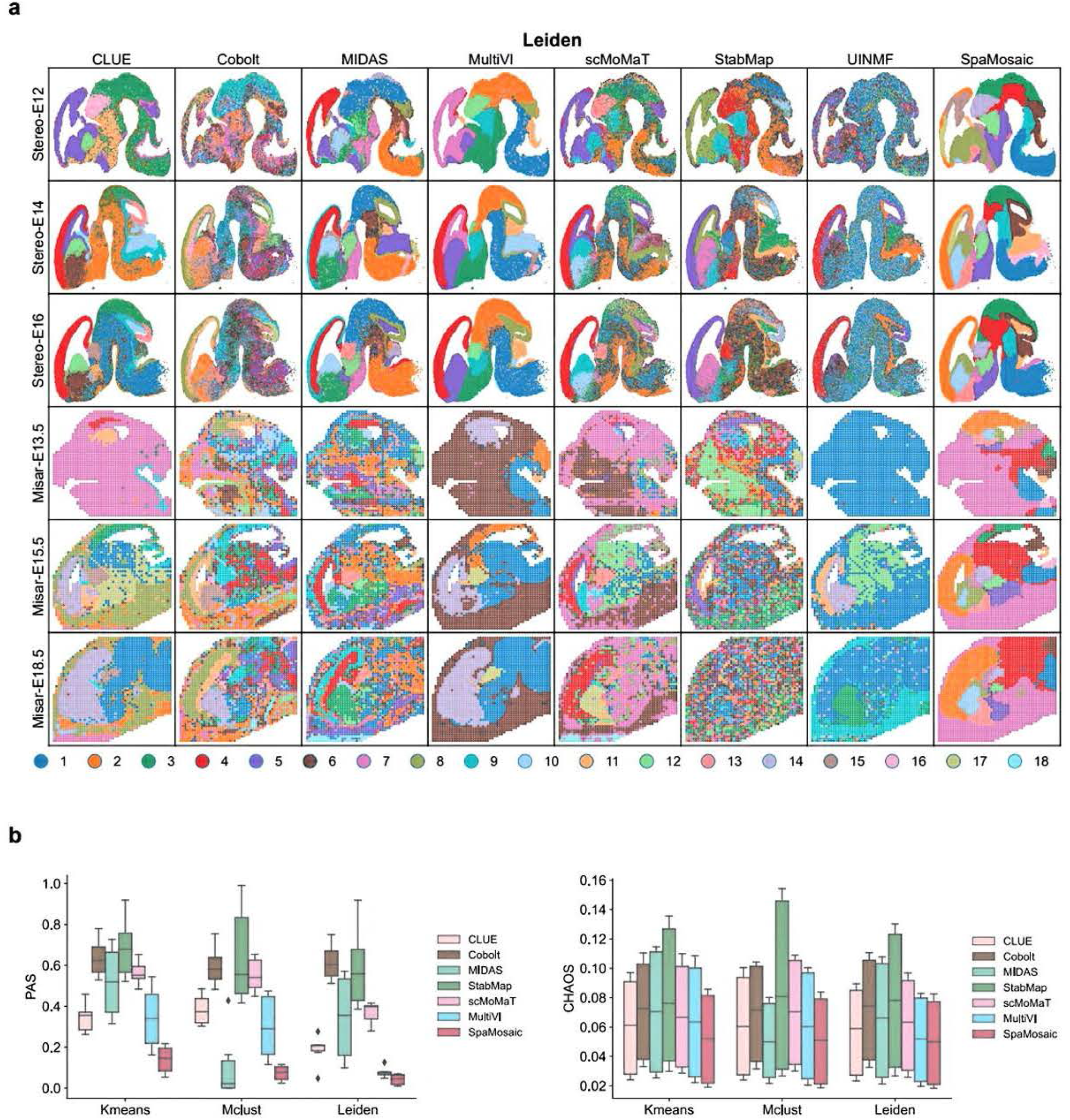
Benchmarking of mosaic integration methods using different clustering algorithms on the embryonic mouse brain dataset (Misar+Stereo). **a** Spatial plots of clustering results from benchmarked mosaic integration methods using the Leiden clustering algorithm. **b** Performance comparison using PAS and CHAOS metrics.

**Supp. Fig. S50.**
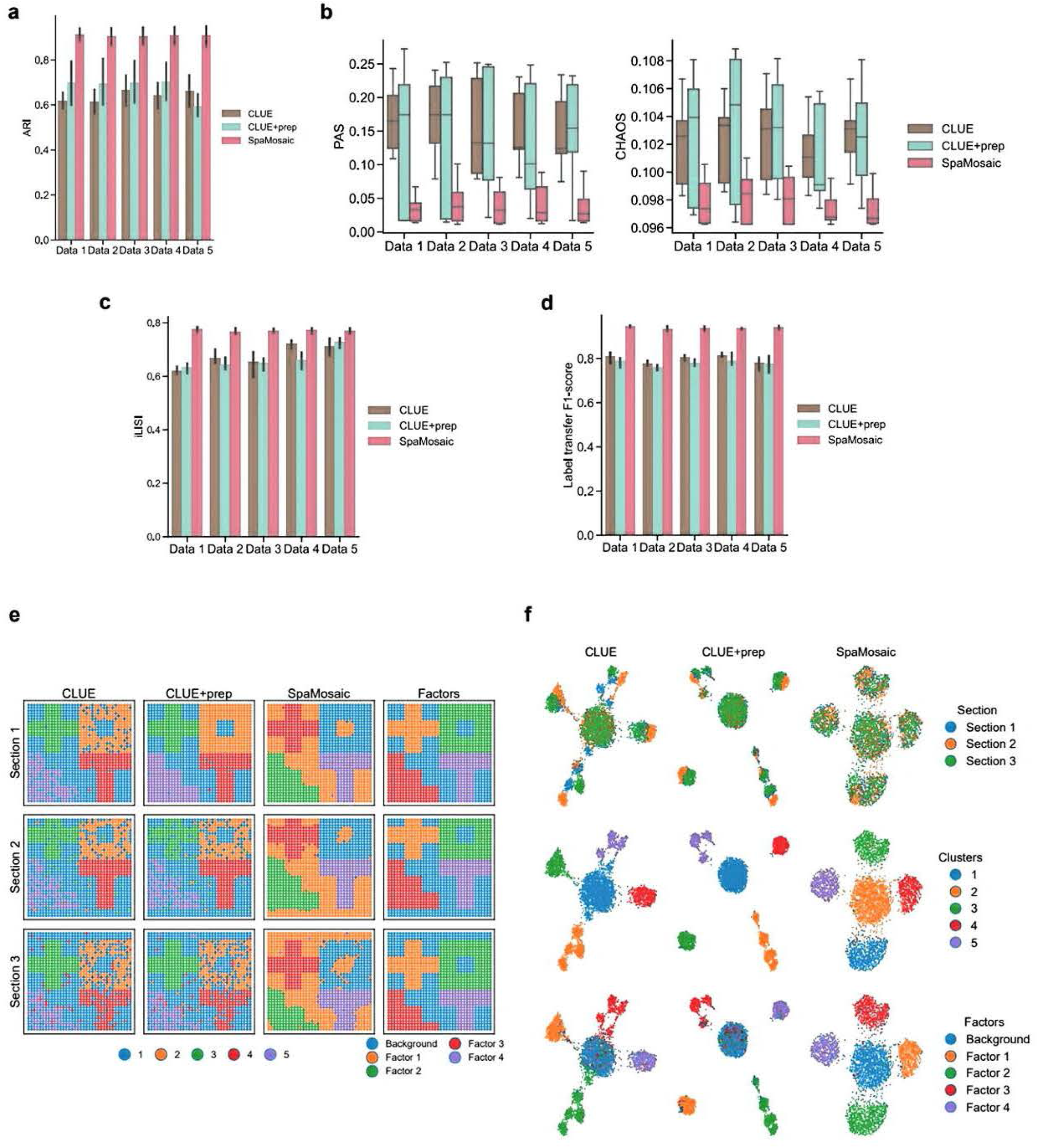
Benchmarking results of SpaMosaic, CLUE, and CLUE+prep on the simulated data. **a** ARI scores of the three configurations on five simulation datasets. **b** PAS and CHAOS scores of the three configurations. **c** iLISI scores of the three method configurations. **d** Label transfer F1-scores of the three configurations. **e** Spatial plots of clustering results of the first replicate of the simulation datasets. The rightmost column shows the ground truth of factor patterns. **f** UMAP plots of embeddings of the first replicate of simulation datasets. From top to bottom, the spots are colored by section, cluster and factor labels.

**Supp. Fig. S51.**
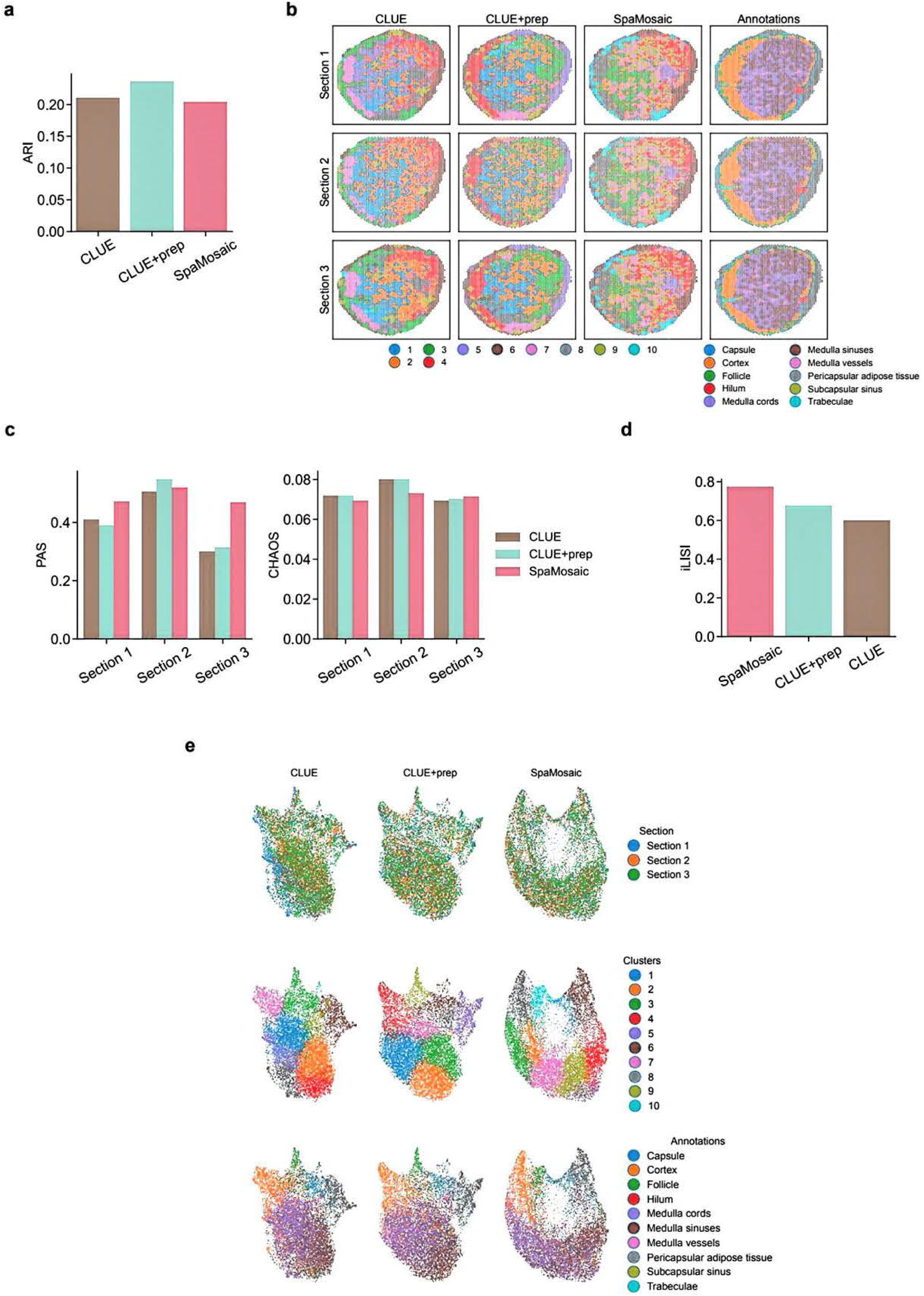
Benchmarking results of SpaMosaic, CLUE, and CLUE+prep on the human lymph node dataset. a ARI scores of the three method configurations. **b** Spatial plots of clustering results. The rightmost column shows the expert 1nanual annotations. **c** PAS and CHAOS scores of the three configurations on individual sections. **d** iLISl scores of the three configurations. **d** Label transfer F1-scores of the three configurations. **e** UMAP plots of embeddings from the three configurations. From top to bottom, the spots are colored by section, cluster and annotation labels.

**Supp. Fig. S52.**
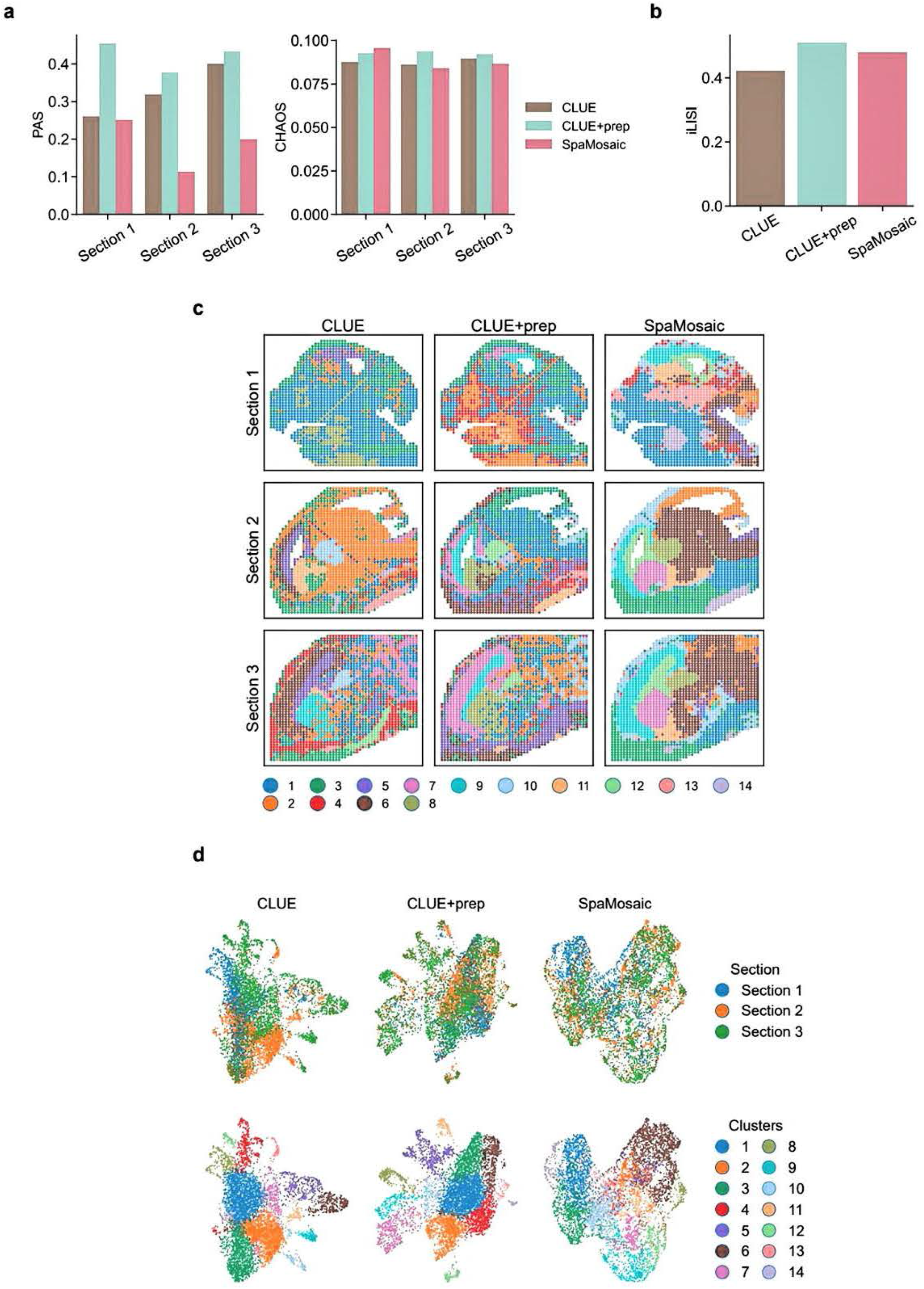
Benchmarking results of SpaMosaic, CLUE, and CLUE+prep on the embryonic mouse brain dataset (Misar). **a** PAS and CHAOS scores of the three configurations on individual sections. **b** iLISI scores of the three configurations. **c** Spatial plots of clustering results from the three configurations. **d** UMAP plots of embeddings from three methods. From top to bottom, the spots are colored by section and cluster labels.

**Supp. Fig. S53.**
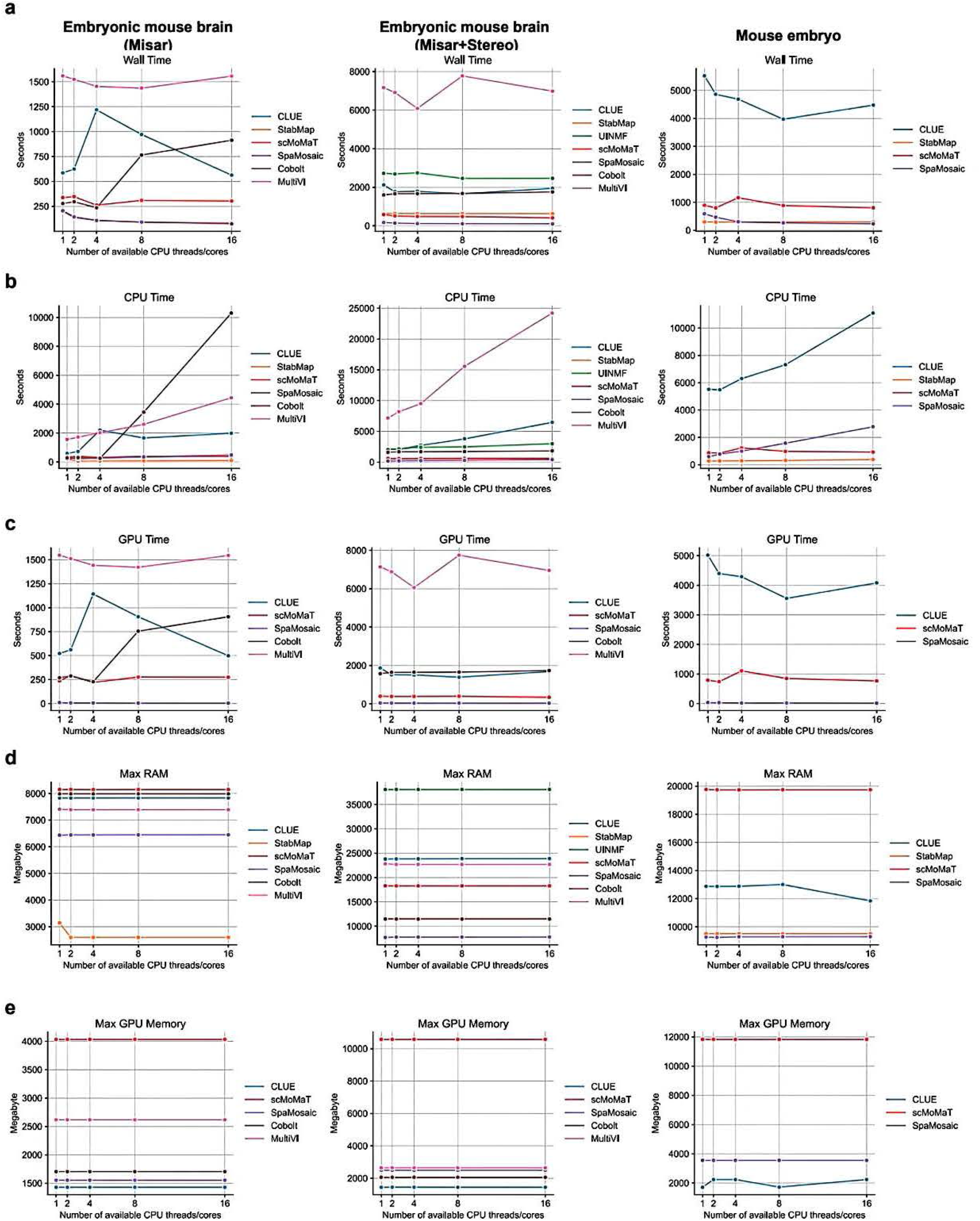
Computational cost comparison of mosaic integration methods across different CPU core counts. **a-e** Computational performance comparison of different mosaic integration methods on the embryonic mouse brain (Misar), embryonic mouse brain (Misar+Stereo), and mouse embryo datasets, measuring the wall time (**a**), CPU time (**b**), GPU time (**c**), maximum memory consumption (**d**), and maximum GPU memory consumption (**e**).

**Supp. Fig. S54.**
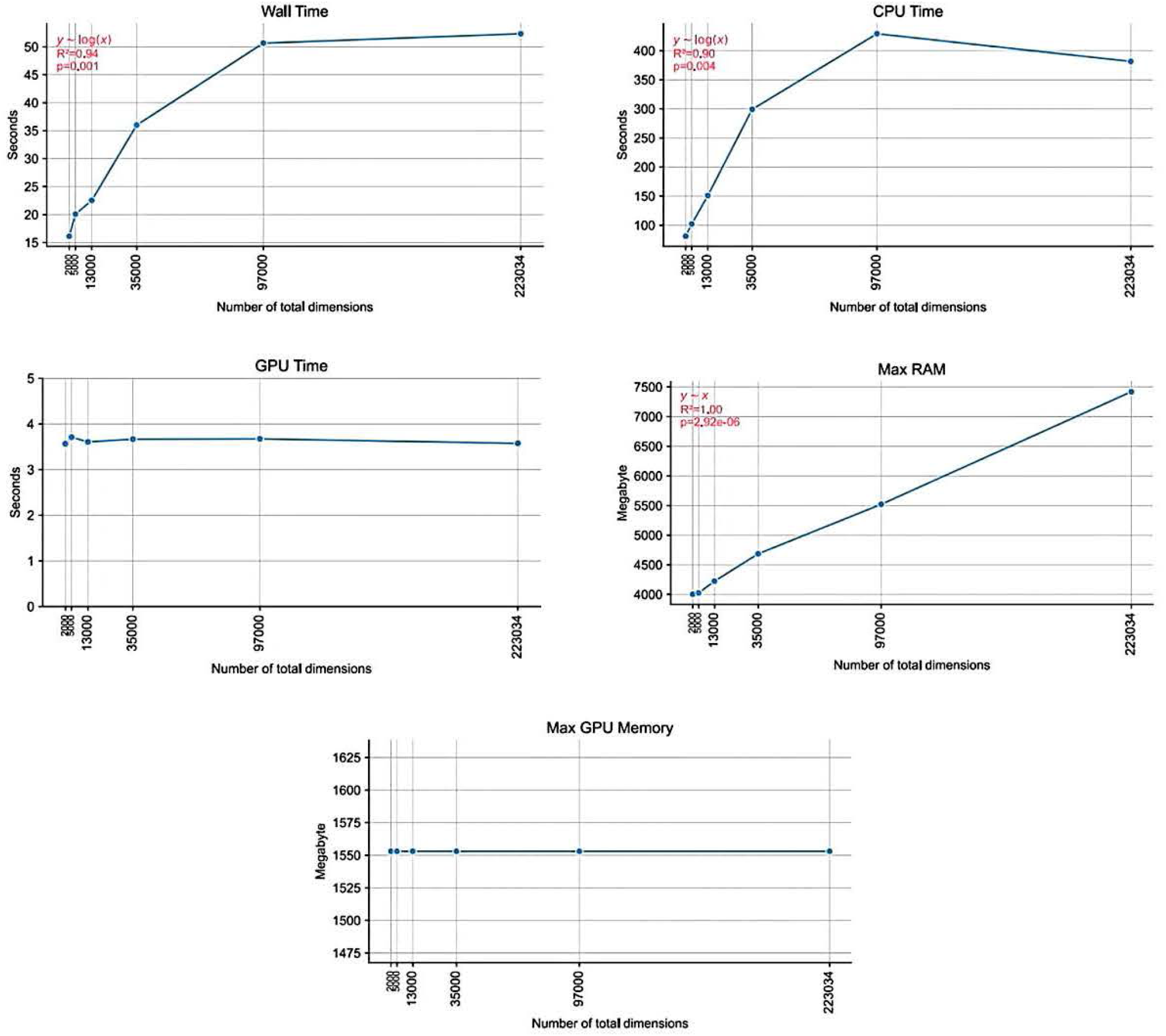
Computational cost of SpaMosaic with different input data dimensions. To assess the impact of input feature dimensionality on computational performance, we conducted experiments using the embryonic mouse brain (Misar) dataset, which contains two modalities (RNA and ATAC) across three batches: one batch with both modalities (1949 spots), one with RNA only (1777 spots), and one with ATAC only (2129 spots). We randomly sampled varying numbers of features from the RNA and ATAC modalities. For RNA, 1,000; 2,000; 4,000; 8,000; 16,000; and 32,000 features were sampled; for ATAC, 1,000; 3,000; 9,000; 27,000; 81,000; and 191,034 features were sampled. Each RNA-ATAC feature pair was used to evaluate SpaMosaic’s runtime and memory consumption, enabling a systematic analysis of how increasing data dimensionality affects computational cost. The annotations (red) indicate the fitted functional form, coefficient of determination **(R**^**2**^) and significance level (*p*).

**Supp. Fig. S55.**
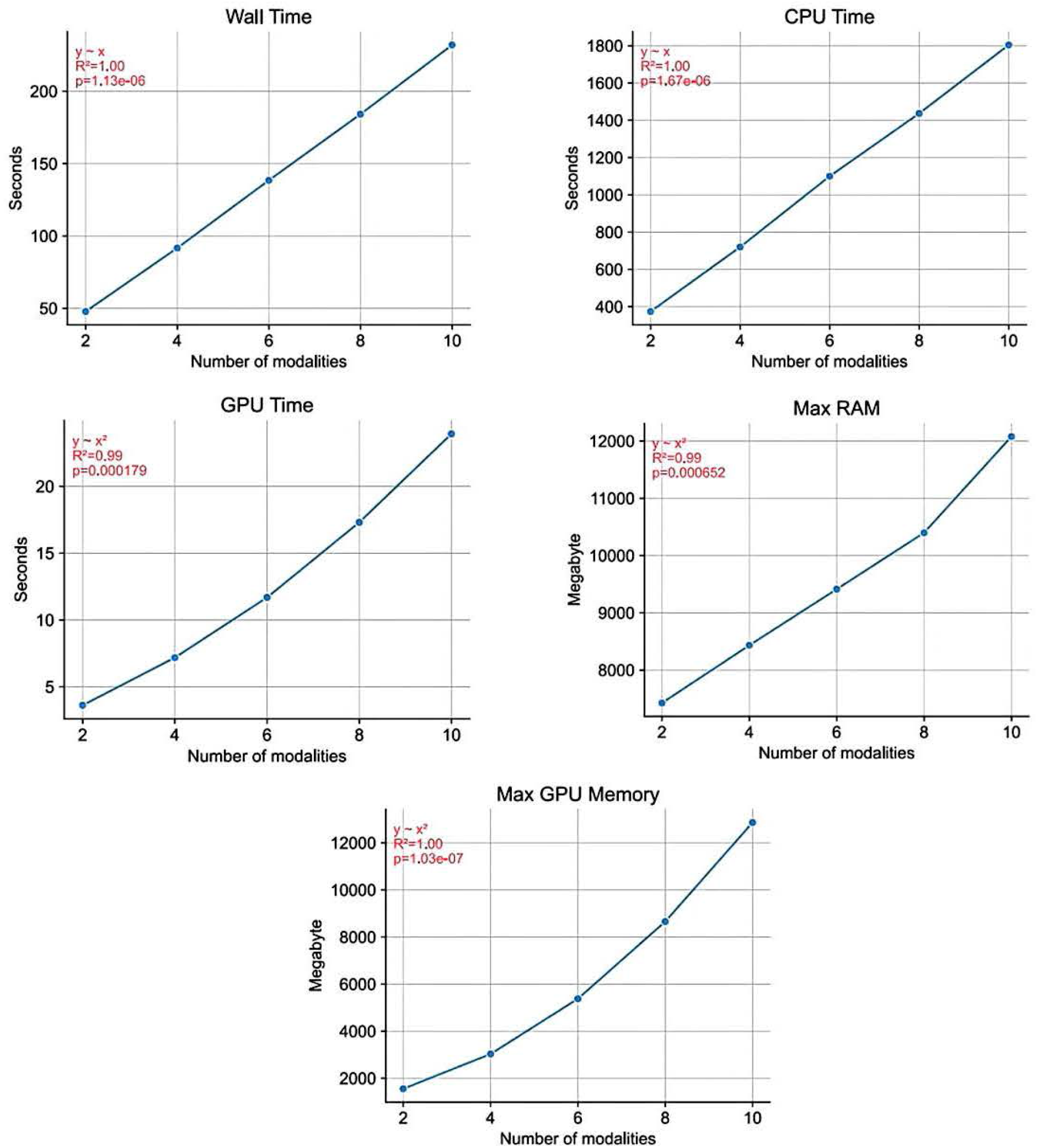
Computational cost of SpaMosaic with different numbers of input data modalities. To evaluate how the number of input modalities affects computational performance, we generated simulated datasets from the embryonic mouse brain (Misar) dataset, which originally contains two modalities (RNA and ATAC) across three batches: one batch with both modalities (1949 spots), one with RNA only (1777 spots), and one with ATAC only (2129 spots). Additional modalities were created by replicating each original modality within each batch by a specified multiplication factor and treating each copy as a distinct modality. For example, with a factor of 2, the batch containing both RNA and ATAC becomes {RNA-I, RNA-2, ATAC-1, ATAC-2}, whereas the RNA-only and ATAC-only batches become {RNA-I, RNA-2} and {ATAC-1, ATAC-2}, respectively, yielding 4 modalities in total. Multiplication factors from 1 to 5 were tested, corresponding to 2, 4, 6, 8, and 10 modalities. The annotations (red) indicate the fitted functional form, coefficient of determination (R^2^) and significance level (*p*).

**Supp. Fig. S56.**
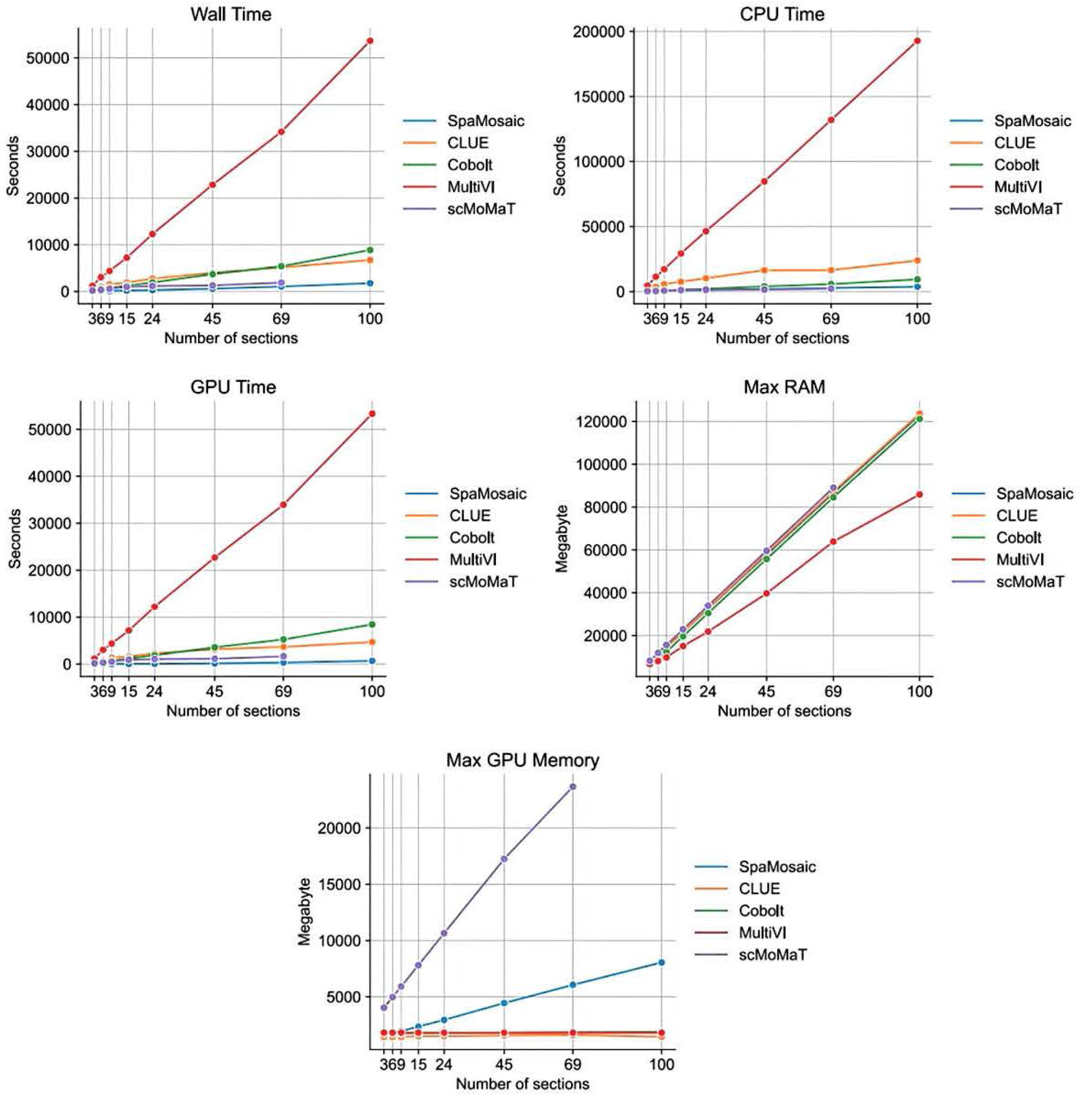
Computational cost comparison of mosaic integration methods across different numbers of input sections. To assess how the number of sections influences computational performance, we generated simulated datasets based on the embryonic mouse brain (Misar) dataset, which originally contains three sections (one multimodal section with 1949 spots, one RNA-only section with 1777 spots, and one ATAC-only section with 2129 spots). Additional sections were created by replicating each original section and its corresponding modalities according to predefined multiplication factors. For example, with a factor of 2, each of the three original sections was duplicated once, yielding a dataset with 2 multimodal sections, 2 RNA-only sections, and 2 ATAC-only sections. Replication settings were selected to produce total section counts of 3, 6, 9, 15, 24, 45, 69, and 100. For all configurations except the 100-section setting, the three original sections were replicated equally; the 100-section dataset was generated by replicating the first two sections 33 times and the third section 34 times. Only GPU-based methods were compared.

**Supp. Fig. S57.**
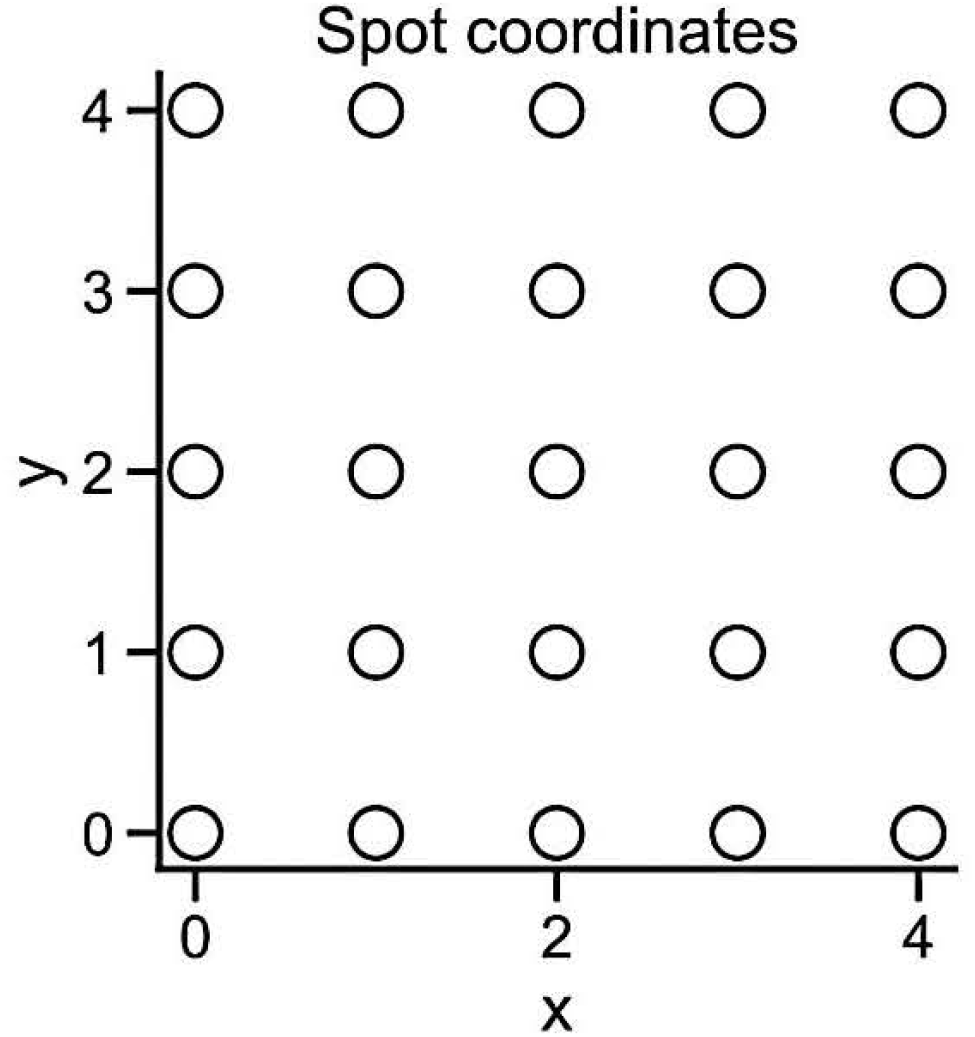
Illustration of spatial coordinates from the largest simulated section for measuring computational cost.

**Supp. Fig. S58.**
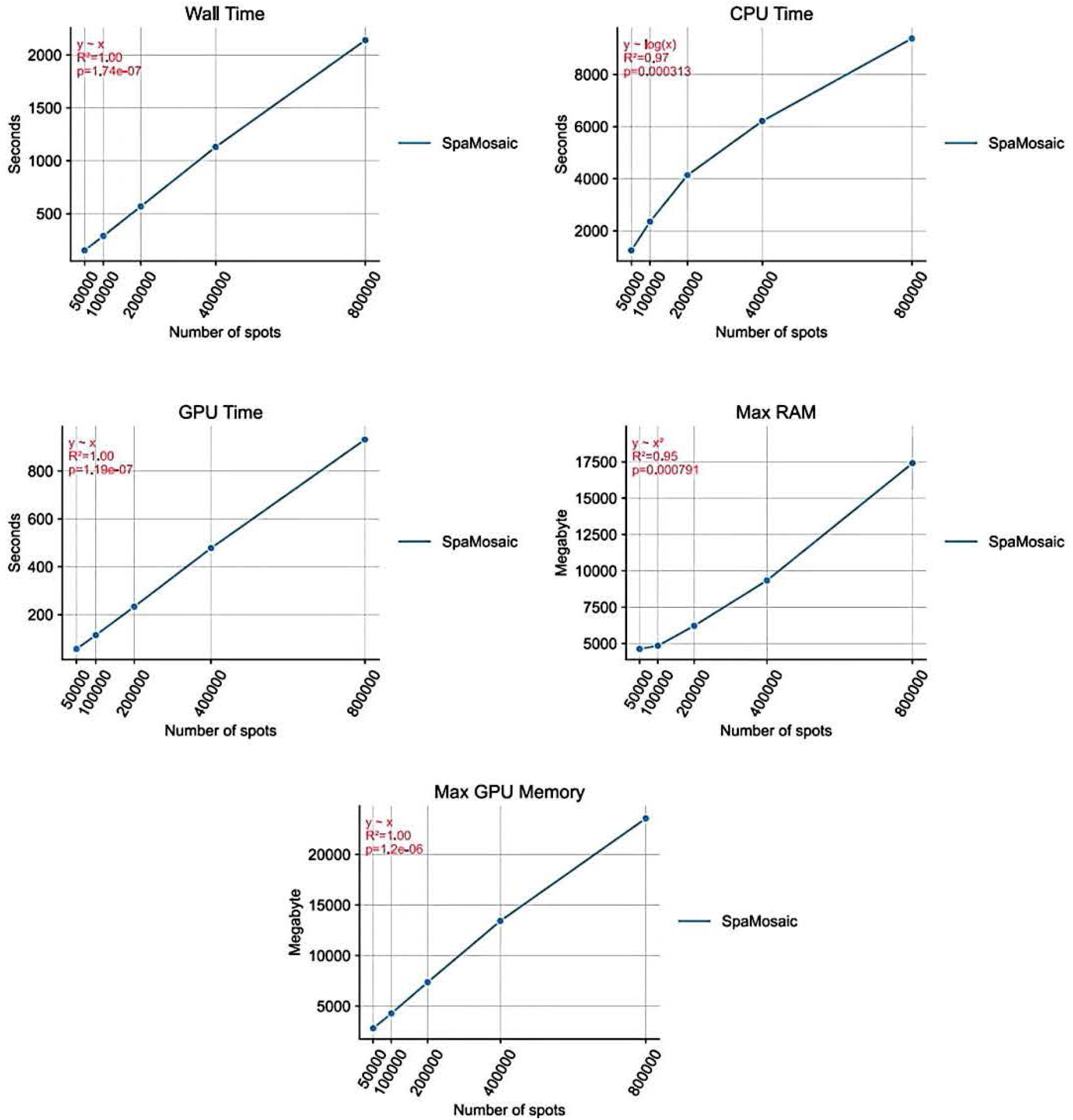
Computational scalability of SpaMosaic with increasing number of spots per input section. To evaluate SpaMosaic’s scalability with respect to spot density, we generated simulated datasets using the embryonic mouse brain (Misar) dataset as a template. Each dataset configuration consisted of three sections: one multimodal section (RNA+ATAC) and two single-modality sections (RNA-only and ATAC-only), with all three sections containing the same number of spots. RNA profiles were simulated with 10,000 genes using a Poisson distribution with parameter *λ* = 1, and ATAC profiles were simulated with 100,000 peaks encoded as binary values with 99.9% sparsity. We created datasets with 50,000, 100,000, 200,000, 400,000, and 800,000 spots per section. The annotations (red) indicate the fitted functional form, coefficient of determination (R^2^) and significance level *(p)*.

## Reference

1. Deng, Y. et al. Spatial-CUT&Tag: Spatially resolved chromatin modification profiling at the cellular level. Science 375, 681–686 (2022).

2. Zhang, D. et al. Spatial epigenome-transcriptome co-profiling of mammalian tissues. Nature 616, 113–122 (2023).

3. Ben-Chetrit, N. et al. Integration of whole transcriptome spatial profiling with protein markers. Nature biotechnoloy 41, 788–793 (2023).

4. Zhang, D. et al. Spatial dynamics of mammalian brain development and neuroinflammation by multimodal tri-omics mapping. bioRxiv, 2024.07.28.605493 (2024).

5. Guo, P. et al. Multiplexed spatial mapping of chromatin features, transcriptome and proteins in tissues. Nat Methods 22, 520–529 (2025).

6. Long, Y. et al. Deciphering spatial domains from spatial multi-omics with SpatialGlue. Nature methods 21, 1658–1667 (2024).

7. Argelaguet, R., Cuomo, A.S.E., Stegle, O. & Marioni, J.C. Computational principles and challenges in single-cell data integration. Nature biotechnology 39, 1202–1215 (2021).

8. Zhang, Z. et al. scMoMaT jointly performs single cell mosaic integration and multi-modal bio-marker detection. Nature communications 14, 384 (2023).

9. Gong, B., Zhou, Y. & Purdom, E. Cobolt: integrative analysis of multimodal single-cell sequencing data. Genome biology 22, 351 (2021).

10. Ghazanfar, S., Guibentif, C. & Marioni, J.C. Stabilized mosaic single-cell data integration using unshared features. Nature biotechnology 42, 284–292 (2024).

11. He, Z. et al. Mosaic integration and knowledge transfer of single-cell multimodal data with MIDAS. Nature biotechnology 42, 1594–1605 (2024).

12. Singhal, V. et al. BANKSY unifies cell typing and tissue domain segmentation for scalable spatial omics data analysis. Nature Genetics 56, 431–441 (2024).

13. Xu, H. et al. Unsupervised spatially embedded deep representation of spatial transcriptomics. Genome medicine 16, 12 (2024).

14. Long, Y. et al. Spatially informed clustering, integration, and deconvolution of spatial transcriptomics with GraphST. Nature communications 14, 1155 (2023).

15. Zhou, X., Dong, K. & Zhang, S. Integrating spatial transcriptomics data across different conditions, technologies and developmental stages. Nature computational science 3, 894–906 (2023).

16. Wang, G. et al. Construction of a 3D whole organism spatial atlas by joint modelling of multiple slices with deep neural networks. Nature machine intelligence 5, 1200–1213 (2023).

17. Varrone, M., Tavernari, D., Santamaria-Martinez, A., Walsh, L.A. & Ciriello, G. CellCharter reveals spatial cell niches associated with tissue remodeling and cell plasticity. Nature Genetics 56, 74–84 (2024).

18. Li, Z. et al. Cross-modality representation and multi-sample integration of spatially resolved omics data. bioRxiv (2025).

19. Chen, T., Kornblith, S., Norouzi, M. & Hinton, G. A simple framework for contrastive learning of visual representations. in International conference on machine learning 1597–1607 (PMLR, 2020).

20. Korsunsky, I. et al. Fast, sensitive and accurate integration of single-cell data with Harmony. Nature methods 16, 1289–1296 (2019).

21. Stuart, T. et al. Comprehensive Integration of Single-Cell Data. Cell 177, 1888–1902 e21 (2019).

22. Haghverdi, L., Lun, A.T.L., Morgan, M.D. & Marioni, J.C. Batch effects in single-cell RNA-sequencing data are corrected by matching mutual nearest neighbors. Nature biotechnology 36, 421–427 (2018).

23. Velickovic, P. et al. Graph Attention Networks. ArXiv abs/1710.10903(2017).

24. Dong, K. & Zhang, S. Deciphering spatial domains from spatially resolved transcriptomics with an adaptive graph attention auto-encoder. Nature communications 13, 1739 (2022).

25. Hu, Z., Dong, Y., Wang, K. & Sun, Y. Heterogeneous Graph Transformer. Proceedings of The Web Conference 2020 (2020).

26. He, X. et al. LightGCN: Simplifying and Powering Graph Convolution Network for Recommendation. Proceedings of the 43rd International ACM SIGIR Conference on Research and Development in Information Retrieval (2020).

27. Tu, X., Cao, Z.-J., Xia, C.-R., Mostafavi, S. & Gao, G. Cross-linked unified embedding for cross-modality representation learning. in Advances in Neural Information Processing Systems (2022).

28. Townes, F.W. & Engelhardt, B.E. Nonnegative spatial factorization applied to spatial genomics. Nature methods 20, 229–238 (2023).

29. Shang, L. & Zhou, X. Spatially aware dimension reduction for spatial transcriptomics. Nature Communications 13, 7203 (2022).

30. Luecken, M.D. et al. Benchmarking atlas-level data integration in single-cell genomics. Nature methods 19, 41–50 (2022).

31. Traag, V.A., Waltman, L. & Van Eck, N.J. From Louvain to Leiden: guaranteeing well-connected communities. Scientific reports 9, 1–12 (2019).

32. Becht, E. et al. Dimensionality reduction for visualizing single-cell data using UMAP. Nature biotechnology (2018).

33. Jiang, F. et al. Simultaneous profiling of spatial gene expression and chromatin accessibility during mouse brain development. Nature methods 20, 1048–1057 (2023).

34. Ashuach, T. et al. MultiVI: deep generative model for the integration of multimodal data. Nature methods 20, 1222–1231 (2023).

35. Science, A.I.f.B. Allen Reference Atlas – Mouse Brain. (2011).

36. Chen, A. et al. Spatiotemporal transcriptomic atlas of mouse organogenesis using DNA nanoball-patterned arrays. Cell 185, 1777–1792 e21 (2022).

37. Simeone, A. & Acampora, D. The role of Otx2 in organizing the anterior patterning in mouse. Int J Dev Biol 45, 337–45 (2001).

38. Bishop, K.M., Rubenstein, J.L. & O’Leary, D.D. Distinct actions of Emx1, Emx2, and Pax6 in regulating the specification of areas in the developing neocortex. J Neurosci 22, 7627–38 (2002).

39. Bertacchi, M. et al. NR2F1 regulates regional progenitor dynamics in the mouse neocortex and cortical gyrification in BBSOAS patients. EMBO J 39, e104163 (2020).

40. Yun, K., Mantani, A., Garel, S., Rubenstein, J. & Israel, M.A. Id4 regulates neural progenitor proliferation and differentiation in vivo. Development 131, 5441–8 (2004).

41. Lein, E.S. et al. Genome-wide atlas of gene expression in the adult mouse brain. Nature 445, 168–76 (2007).

42. Singh, R. et al. Unsupervised manifold alignment for single-cell multi-omics data. ACM BCB 2020, 1–10 (2020).

43. Lücken, M. et al. A sandbox for prediction and integration of DNA, RNA, and proteins in single cells. in NeurIPS Datasets and Benchmarks (2021).

44. Khandelwal, N. et al. FOXP1 regulates the development of excitatory synaptic inputs onto striatal neurons and induces phenotypic reversal with reinstatement. Sci Adv 10, eadm7039 (2024).

45. Zeisel, A. et al. Molecular Architecture of the Mouse Nervous System. Cell 174, 999–1014 e22 (2018).

46. Kriebel, A.R. & Welch, J.D. UINMF performs mosaic integration of single-cell multi-omic datasets using nonnegative matrix factorization. Nature Communications 13, 780 (2022).

47. Puighermanal, E. et al. Functional and molecular heterogeneity of D2R neurons along dorsal ventral axis in the striatum. Nat Commun 11, 1957 (2020).

48. Diamant, I., Clarke, D.J.B., Evangelista, J.E., Lingam, N. & Ma’ayan, A. Harmonizome 3.0: integrated knowledge about genes and proteins from diverse multi-omics resources. Nucleic Acids Res 53, D1016–D1028 (2025).

49. Lopez, R., Regier, J., Cole, M.B., Jordan, M.I. & Yosef, N. Deep generative modeling for single-cell transcriptomics. Nature Methods 15, 1053–1058 (2018).

50. Wu, K.E., Yost, K.E., Chang, H.Y. & Zou, J. BABEL enables cross-modality translation between multiomic profiles at single-cell resolution. Proc Natl Acad Sci U S A 118(2021).

51. Gayoso, A. et al. Joint probabilistic modeling of single-cell multi-omic data with totalVI. Nature Methods 18, 272–282 (2021).

52. Hu, Y. et al. WEDGE: imputation of gene expression values from single-cell RNA-seq datasets using biased matrix decomposition. Brief Bioinform 22(2021).

53. Hu, Y. et al. Benchmarking algorithms for single-cell multi-omics prediction and integration. Nat Methods 21, 2182–2194 (2024).

54. Wu, S.J. et al. Single-cell CUT&Tag analysis of chromatin modifications in differentiation and tumor progression. Nat Biotechnol 39, 819–824 (2021).

55. Yu, G., Wang, L.G., Han, Y. & He, Q.Y. clusterProfiler: an R package for comparing biological themes among gene clusters. Omics: a journal of integrative biology 16, 284–7 (2012).

56. Zhang, S. et al. Sox2 Is Essential for Oligodendroglial Proliferation and Differentiation during Postnatal Brain Myelination and CNS Remyelination. J Neurosci 38, 1802–1820 (2018).

57. Sock, E. & Wegner, M. Using the lineage determinants Olig2 and Sox10 to explore transcriptional regulation of oligodendrocyte development. Dev Neurobiol 81, 892–901 (2021).

58. Yang, F. et al. Proteomics of the corpus callosum to identify novel factors involved in hypomyelinated Niemann-Pick Type C disease mice. Mol Brain 12, 17 (2019).

59. Quintes, S. et al. Zeb2 is essential for Schwann cell differentiation, myelination and nerve repair. Nat Neurosci 19, 1050–1059 (2016).

60. Zhao, C. et al. Dual Requirement of CHD8 for Chromatin Landscape Establishment and Histone Methyltransferase Recruitment to Promote CNS Myelination and Repair. Dev Cell 45, 753–768 e8 (2018).

61. Farrell, K. et al. Genetic, transcriptomic, histological, and biochemical analysis of progressive supranuclear palsy implicates glial activation and novel risk genes. Nat Commun 15, 7880 (2024).

62. Brockschnieder, D., Sabanay, H., Riethmacher, D. & Peles, E. Ermin, a myelinating oligodendrocyte-specific protein that regulates cell morphology. J Neurosci 26, 757–62 (2006).

63. Morita, J. et al. Structure and biological function of ENPP6, a choline-specific glycerophosphodiester-phosphodiesterase. Sci Rep 6, 20995 (2016).

64. Chen, X., Lu, W. & Wu, D. Sirtuin 2 (SIRT2): Confusing Roles in the Pathophysiology of Neurological Disorders. Front Neurosci 15, 614107 (2021).

65. Edvardson, S. et al. Mutations in the fatty acid 2-hydroxylase gene are associated with leukodystrophy with spastic paraparesis and dystonia. Am J Hum Genet 83, 643–8 (2008).

66. Voskuhl, R.R. et al. Gene expression in oligodendrocytes during remyelination reveals cholesterol homeostasis as a therapeutic target in multiple sclerosis. Proc Natl Acad Sci U S A 116, 10130–10139 (2019).

67. Sharma, K., Singh, J. & Pillai, P.P. MeCP2 Differentially Regulate the Myelin MBP and PLP Protein Expression in Oligodendrocytes and C6 Glioma. J Mol Neurosci 65, 343–350 (2018).

68. Genomics, x. Visium HD CytAssist Gene Expression Libraries of Mouse Brain (H&E). (2024).

69. Genomics, x. Mouse Brain Section (Coronal) - 1 Standard. (2022).

70. Deng, Y. et al. Spatial profiling of chromatin accessibility in mouse and human tissues. Nature 609, 375–383 (2022).

71. Yao, Z. et al. A transcriptomic and epigenomic cell atlas of the mouse primary motor cortex. Nature 598, 103–110 (2021).

72. Nieto, M. et al. Expression of Cux-1 and Cux-2 in the subventricular zone and upper layers II-IV of the cerebral cortex. J Comp Neurol 479, 168–80 (2004).

73. Puelles, L. et al. Pallial and subpallial derivatives in the embryonic chick and mouse telencephalon, traced by the expression of the genes Dlx-2, Emx-1, Nkx-2.1, Pax-6, and Tbr-1. J Comp Neurol 424, 409–38 (2000).

74. Pla, R. et al. Dlx1 and Dlx2 Promote Interneuron GABA Synthesis, Synaptogenesis, and Dendritogenesis. Cereb Cortex 28, 3797–3815 (2018).

75. van Velthoven, C.T.J. et al. The transcriptomic and spatial organization of telencephalic GABAergic neuronal types. bioRxiv (2024).

76. Lipiec, M.A. et al. TCF7L2 regulates postmitotic differentiation programmes and excitability patterns in the thalamus. Development 147(2020).

77. Guo, Q. & Li, J.Y.H. Defining developmental diversification of diencephalon neurons through single cell gene expression profiling. Development 146(2019).

78. Lahti, L. et al. Differentiation and molecular heterogeneity of inhibitory and excitatory neurons associated with midbrain dopaminergic nuclei. Development 143, 516–29 (2016).

79. Isogai, E., Okumura, K., Saito, M., Tokunaga, Y. & Wakabayashi, Y. Meis1 plays roles in cortical development through regulation of cellular proliferative capacity in the embryonic cerebrum. Biomed Res 43, 91–97 (2022).

80. Asano, M. & Gruss, P. Pax-5 is expressed at the midbrain-hindbrain boundary during mouse development. Mech Dev 39, 29–39 (1992).

81. Stoykova, A., Treichel, D., Hallonet, M. & Gruss, P. Pax6 modulates the dorsoventral patterning of the mammalian telencephalon. J Neurosci 20, 8042–50 (2000).

82. Leggere, J.C. et al. NOVA regulates Dcc alternative splicing during neuronal migration and axon guidance in the spinal cord. Elife 5(2016).

83. Yang, M. et al. Contrastive learning enables rapid mapping to multimodal single-cell atlas of multimillion scale. Nature Machine Intelligence 4, 696–709 (2022).

84. He, K., Fan, H., Wu, Y., Xie, S. & Girshick, R.B. Momentum Contrast for Unsupervised Visual Representation Learning. 2020 IEEE/CVF Conference on Computer Vision and Pattern Recognition (CVPR), 9726–9735 (2019).

85. Radford, A. et al. Learning transferable visual models from natural language supervision. in International conference on machine learning 8748–8763 (PMLR, 2021).

86. Zheng, R. et al. A Flexible Data-Driven Framework for Correcting Coarsely Annotated scRNA-seq Data. Big Data Mining and Analytics (2025).

87. Waikhom, L. & Patgiri, R. Graph Neural Networks: Methods, Applications, and Opportunities. ArXiv abs/2108.10733(2021).

88. Hadsell, R., Rao, D., Rusu, A.A. & Pascanu, R. Embracing Change: Continual Learning in Deep Neural Networks. Trends in Cognitive Sciences 24, 1028–1040 (2020).

89. Weiss, K.R., Khoshgoftaar, T.M. & Wang, D. A survey of transfer learning. Journal of Big Data 3, 1–40 (2016).

90. Liu, F.T., Ting, K.M. & Zhou, Z.-H. Isolation Forest. 2008 Eighth IEEE International Conference on Data Mining, 413–422 (2008).

91. Xia, C.R., Cao, Z.J., Tu, X.M. & Gao, G. Spatial-linked alignment tool (SLAT) for aligning heterogenous slices. Nature communications 14, 7236 (2023).

92. Maas, A.L. Rectifier Nonlinearities Improve Neural Network Acoustic Models. (2013).

93. Ioffe, S. & Szegedy, C. Batch Normalization: Accelerating Deep Network Training by Reducing Internal Covariate Shift. ArXiv abs/1502.03167(2015).

94. Srivastava, N., Hinton, G.E., Krizhevsky, A., Sutskever, I. & Salakhutdinov, R. Dropout: a simple way to prevent neural networks from overfitting. Journal of machine learning research 15, 1929–1958 (2014).

95. Fey, M. & Lenssen, J.E. Fast Graph Representation Learning with PyTorch Geometric. ArXiv abs/1903.02428(2019).

96. Kingma, D.P. & Ba, J. Adam: A Method for Stochastic Optimization. CoRR abs/1412.6980(2014).

97. Pedregosa, F. et al. Scikit-learn: Machine learning in Python. the Journal of machine Learning research 12, 2825–2830 (2011).

98. Cao, Z.J. & Gao, G. Multi-omics single-cell data integration and regulatory inference with graph-linked embedding. Nature Biotechnology 40, 1458–1466 (2022).

99. Clevert, D.-A. Fast and accurate deep network learning by exponential linear units (elus). arXiv preprint arXiv:1511.07289 (2015).

100. Lotfollahi, M. et al. Mapping single-cell data to reference atlases by transfer learning. Nat Biotechnol 40, 121–130 (2022).

101. Fraley, C. & Rafter, A.E. MCLUST: Software for Model-Based Cluster and Discriminant Analysis. Journal of Classification (1998).

102. Blondel, V.D., Guillaume, J.-L., Lambiotte, R. & Lefebvre, E. Fast unfolding of communities in large networks. Journal of statistical mechanics: theory and experiment 2008, P10008 (2008).

103. Wolf, F.A., Angerer, P. & Theis, F.J. SCANPY: large-scale single-cell gene expression data analysis. Genome Biology 19, 15 (2018).

