## supplementary notes for the main text for "Mosaic integration of spatial multi-omics with SpaMosaic"

**Supplementary Information**

### Application of SpaMosaic to human lymph node and human tonsil datasets.

**Lymph node (RNA+protein, Supp. Fig. S28-S30)**

Here we benchmarked SpaMosaic and competing methods on an in-house generated human lymph node dataset. Lymph nodes are small organs found throughout the human body as part of the lymphatic system. Within the lymph node, the cortex and medulla harbors concentrations of different immune cells like T and B cells that become activated during an infection. In particular, the B cells congregations in the cortex are called follicles. The organ is surrounded by a protective capsule of fibrous tissue, followed by the subscapsular sinus, a space through which lymphatic fluids flow through. The dataset has three tissue sections where the first and second sections are sequential. The 10x Genomics Visium RNA and protein co-profiling technology was used to acquire the transcriptome and protein profiles for each section. To create a mosaic dataset, we removed the protein and RNA profiles of the second and third section, respectively (**Supp. Fig. S28a**). Prior to integration, we examined the UMAP plots for batch effects. In the RNA modality, we could observe the presence of mild batch effects, but much stronger batch effects were visible in the protein modality between sections 1 and 3 (**Supp. Fig. S28b**). Such levels of batch effects may be attributed to sections 1 and 2 being sequential and sections 1 and 3 being non-sequential sections. For performance assessment, we had the accompanying H&E images (**Supp. Fig. S28c**) annotated by an expert to provide the ground truth. Unlike the other datasets used for benchmarking in the main text, this lymph node dataset presents a unique challenge due to its intricate cell type-centric structures and a different combination of data modalities, namely RNA and protein.

We first visually inspected the results obtained from the different methods. For SpaMosaic, it captured many of the expected lymph node structures across the three sections (**Supp. Fig. S29a**). This includes the follicles (cluster 10), cortex (cluster 8), adipose tissue (cluster 6) and capsule (cluster 1), which we matched to the manual annotations. For the remaining clusters, they primarily matched the annotated medullar regions. Cobolt and StabMap produced similar outputs as SpaMosaic, capturing the follicles (clusters 4 and 9), cortex (clusters 7 and 5) and adipose tissue (clusters 4 and 8). CLUE also detected similar structures, except for the follicles that were incorrectly grouped with other structures (cluster 3). scMoMaT demarcated the cortex (cluster 7) across three sections but was less effective at capturing other structures. MIDAS captured the cortex of each individual section but failed to link the clusters across all three sections. In addition to the mosaic integration methods, we also tested the spatial clustering algorithms. Although BANKSY captured more coherent spatial domains compared to the competing single-cell mosaic integration methods, we could only annotate two of its clusters (clusters 4 and 5), which match the ground truth’s adipose tissue (**Supp. Fig. S29b**). In contrast, CellCharter not only produced coherent spatial domains but also identified a broader range of structures. These included the cortex (cluster 6) and adipose tissue (cluster 9 in the protein modality and cluster 3 in the RNA modality) (**Supp. Fig. S29b**). However, it demarcated the follicles (cluster 1) in the RNA modality of section 1 but failed to capture them in the RNA modality of section 2 or in the protein modality.

In addition to comparing against the expert annotation, we further validated SpaMosaic’s clusters by visualizing the RNA and protein expression of known tissue specific markers (**Supp. Fig. S29c)**. *MARCO* and *FLT4*, which are markers of lymphatic endothelial cells that are enriched in the human lymph medullary region^1^, had high RNA expression in clusters 4 and 5. CD163 and CD27, markers of macrophages and plasma cells that are enriched in medullary region^2^, had high protein expression in clusters, 3, 7, and 9. *PPARG*, *PLIN1*, and *ADIPOQ*, which are highly expressed in adipocytes^3–5^, had high RNA expression in cluster 6. The markers of follicular cells^6–8^, *CR2*, *MS4A1*, *CXCR4*, and *CXCL13*, had high RNA and protein expression in cluster 10. *PECAM1* and *ACTA2*, with reported expression in the capsule or adventitia^9^, had high RNA and protein expression in cluster 1. We also found markers of T-zone reticular cells^9^, *CXCL9*, and *CCL19*, to have high RNA expression in cluster 8. Finally, T cell markers CD3E, CD4, and CCR7 had high protein expression in clusters 2 and 8.

We next quantitatively evaluated SpaMosaic’s performance in three aspects: spatial domain identification and continuity (ARI, PAS, and CHAOS), batch mixing (iLISI), and modality alignment (FOSCTTM and MS). In terms of spatial domain identification, CLUE achieved the highest ARI, while SpaMosaic was the second highest (**Supp.** **Fig. S30a**). SpaMosaic ranked fourth in overall PAS score, while its overall CHAOS score ranked second (**Supp.** **Fig. S30b**). Unlike its performance on the other presented examples, SpaMosaic did not achieve a clear advantage in the spatial continuity metrics for this lymph node dataset. This may be primarily due to its noisy identification of the large medullary region located at the center of the sections. For batch mixing, SpaMosaic had the top iLISI score, followed by Cobolt (**Supp.** **Fig. S30c**). We also visually inspected the UMAPs colored by section, cluster, and manual annotation (**Supp.** **Fig. S30d**). SpaMosaic mixed the three sections uniformly, matching its iLISI score, while maintaining separation between the manually annotated clusters. In contrast, MIDAS and scMoMaT failed to mix the three sections with visible separation between the sections. Finally, we assessed modality alignment with FOSCTTM and MS. To calculate the FOSCTTM and MS metrics, we created a new mosaic dataset using the first and second sections, where we split the RNA and protein profiles of the second section to obtain two new sections. For both metrics, SpaMosaic was the top performer (**Supp.** **Fig. S30e**).

**Tonsil (RNA+protein, Supp. Fig. S31-S33)**

In this example, we benchmarked SpaMosaic and the competing methods on an in-house generated human tonsil dataset. Like lymph nodes, tonsils are lymphoid organs which play a key role in human immunity. They are irregularly structured with B cell follicles embedded in T cell enriched zones that are further surrounded by epithelial and connective tissues. In these follicles, transient germinal centers form to facilitate B cell activation, proliferation, and somatic hypermutation during an immune response. The dataset consists of three sections generated using the 10x Genomics Visium RNA and protein co-profiling technology. The first and second sections were two sequential sections, and transcriptome and protein (ADT) profiles were acquired for all three sections. To create a mosaic dataset, we removed the protein and RNA profiles of the second and third section, respectively (**Supp.** **Fig. S31a**). We first examined the UMAP plots for batch effects. We did not find obvious batch effects in the RNA modality, likely due to sections 1 and 2 being neighboring sections. However, strong batch effects were present in the protein modality for sections 1 and 3, which were non-sequential (**Supp. Fig. S31b**). We also employed expert annotation of the accompanying H&E images to assist in annotating the subsequent analyses (**Supp. Fig. S31a, c**).

Among the mosaic integration methods, Cobolt, StabMap, and SpaMosaic successfully identified the germinal centers (cluster 4), which were surrounded by follicle mantle zones (cluster 3 for Cobolt, cluster 1 for SpaMosaic and cluster 2 for StabMap in sections 1 and 2; **Supp.** **Fig. S32a**). This demonstrated that, despite applying spatial smoothing, SpaMosaic was still capable of capturing relatively small structures in the tissue section. In addition, SpaMosaic identified other key regions like the T-cell enriched zone (cluster 3, corresponding to the tonsillar parenchyma) and connective and epithelial tissues (cluster 2). Among the competing methods, Cobolt produced results most similar to those of SpaMosaic. For StabMap, its performance on section 3 was poor with the captured germinal centers merging with the surrounding regions. CLUE, MIDAS, and scMoMaT could capture only certain structures while missing out on the rest. For the spatial clustering methods MIDAS, BANKSY, and CellCharter, they also failed to detect the germinal centers, either by merging them with the surrounding tissues or by omitting them entirely (**Supp.** **Fig. S32b**).

To annotate SpaMosaic’s clusters, we plotted the relevant cell type marker expression in the RNA and protein modalities (**Supp.** **Fig. S32c**). With the B cell markers *CD19*^7^, we identified the B cell enriched regions, namely clusters 1 (follicles) and 4 (germinal centers). We also found high RNA expression of *IGHD* (naïve B cells)^7^ and BCL2 protein expression (naive and memory B cells)^10^ in cluster 1. For cluster 4, we found high RNA expression of *BCL2A1* and *LMO2* that mark germinal center B cells^6^, expression of PDCD1 protein that regulates B cell survival in germinal centers^6^, as well as both RNA and protein expression of *CR2* which is a marker for mature B cells and follicular dendritic cells^7^. To identify the T cell enriched zones, we plotted the canonical T cell markers of *CD3E*, *CD4*, and *CD8A*, and found their RNA and protein to be highly expressed in cluster 3. To annotate the epithelial tissues, we examined the expression of the canonical epithelial marker EPCAM, which was highly expressed in cluster 2. We also note in cluster 2 the RNA expression of *CD9*^7^, which identifies B cells and plasma cell subsets within tonsillar tissues like the crypts.

We next quantitatively evaluated SpaMosaic’s performance in three aspects: spatial domain identification and continuity (ARI, PAS, and CHAOS), batch mixing (iLISI), and modality alignment (FOSCTTM and MS). In terms of ARI, SpaMosaic was the highest and followed by CLUE, StabMap, and StabMap (**Supp.** **Fig. S33a**). For the spatial continuity metrics, CLUE, MIDAS, and scMoMaT scored the best in PAS, while MIDAS and CLUE were the best in terms of CHAOS. We believe their top performance in both metrics was due to their merging of tissue structures in their clusters (**Supp.** **Fig. S33b**). For batch mixing assessment with iLISI, SpaMosaic was the leading method (**Supp.** **Fig. S33c**). We also cross checked by examining the UMAP plots of the method outputs colored by the section, cluster, and manual annotation labels (**Supp. Fig. S33d**). SpaMosaic, Cobolt, and CLUE showed high levels of section mixing, which was also reflected in their high iLISI scores. To compute the FOSCTTM and MS metrics, we created a new mosaic dataset using the first and second sections, where we split the RNA and protein profiles of the second section to obtain two new sections. For both metrics, SpaMosaic was again the best performing method (**Supp.** **Fig. S33e**).

### Benchmarking imputation performance

**Imputation experiment settings**

After data integration, imputation of missing data modalities is a task that some mosaic data integration methods including SpaMosaic can accomplish. Here we assessed the performance of SpaMosaic, MIDAS, MultiVI, and BABEL at imputing ATAC and protein modalities based on the RNA profiles. Four datasets were used for assessment, the embryonic mouse brain (Misar) dataset (three sections at the E13.5, E15.5, and E18.5 stages, acquired with Misar-seq, RNA and ATAC modalities), postnatal mouse brain (three sections, RNA and ATAC modalities), human tonsil (three sections, RNA and protein modality) and human lymph node (three sections, RNA and protein modality) datasets. For each dataset, we performed three-fold cross validation. Specifically, we removed the ATAC or protein expression profiles from one section and used the other two sections as training datasets. We then evaluated the imputation performance on the section that had any modality removed. Each section was employed once as the test set.

**Benchmarking metrics**

To quantitatively assess the ATAC and protein imputation results, we used the Pearson correlation coefficients (PCC) and Correlation matrix distance (CMD)^11,12^. For ATAC data imputation, we additionally used the area under the receiver operating characteristic curve (AUROC).

*PCC.* For two vectors $x,y$*,* PCC is defined as:

$$PCC\left( x,y \right)=\frac{\sum_{1}^{m} (x_{i}-\bar{x})(y_{i}-\bar{y})}{\sqrt{\sum_{1}^{m} \left( x_{i}-\bar{x} \right)^{2}}\sqrt{\sum_{1}^{m} \left( y_{i}-\bar{y} \right)^{2}}}$$

For each modality, we calculated both spot-spot PCC and feature-feature PCC. For spot-spot PCC, $m$ is the number of feature dimensions, and $x,y$ denotes the feature profiles of two spots where $\bar{x}=\sum_{1}^{m} x_{i}$ and $\bar{y}=\sum_{1}^{m} y_{i}$. For feature-feature PCC, $m$ is the number of spots in the test set, and $x,y$ are features within all spots. We then summarized the results by computing the average value for the spot-spot PCC and feature-figure PCC across all spots and features, respectively. The PCC has a range of [-1, 1], where 1 indicates perfect positive correlation between two variables while -1 indicates perfect negative correlation and 0 indicates no correlation between the two vectors.

*CMD.* CMD measures the difference between two correlation matrices $X$ and $Y$, which is defined as^11,12^:

$$CMD\left( X,Y \right)=1-\frac{trace(XY)}{\left\| X \right\|_{F}\left\| Y \right\|_{F}}$$

where $\left\| \cdot\right\|_{F}$ denotes the Frobenius norm of a matrix, $trace(XY)$ is the sum of entries on the main diagonal of matrix $XY$. Similar to PCC, we calculated both spot-spot CMD and feature-feature CMD. For spot-spot CMD, $X$ and $Y$ denote the spot-spot PCC matrices calculated using the imputed profiles and ground truth, respectively. For feature-feature CMD, $X$ and $Y$ denote the feature-feature PCC matrices calculated using the imputed profiles and ground truth, respectively. CMD has a range of [0, 1], where 0 indicates perfect alignment of two correlation matrices.

*AUROC*. AUROC assesses performance in binary prediction tasks. Before computing the AUROC, we binarized the target peak count matrix using a threshold of 1. AUROC ranges from 0 to 1, where a value of 1 indicates perfect prediction.

*kNN smoothing for ATAC.* As the peak counts of ATAC data are noisy and sparse, its direct usage in performance assessment may result in underestimation of the imputation’s efficacy. Therefore, we adopted the approach of Tal et al.^13^ in *k*NN smoothing of the raw peak count matrix. Specifically, we took the top 50 principal components of the expression data and computed the *k*NN graph ($K = 50$). Thereafter, we computed the average of the neighbors’ expression values for each spot. We then computed the average values of the spot-spot/peak-peak PCC and AUROC metrics.

**Benchmarking methods for imputation**

*MIDAS.* We ran MIDAS using the procedure described for mosaic integration, except for setting the command parameter ‘--act’ to *translate*.

*TotalVI*^14^. We followed the procedure as described in TotalVI’s tutorials (scvi version 0.19.0) (https://docs.scvi-tools.org/en/stable/tutorials/notebooks/multimodal/totalVI.html). The raw count matrices were used as input to TotalVI. The configuration parameters were set as $latent\_distribution=’normal’$, $n\_layers\_decoder=2$, while the other training parameters were left at the defaults.

*MultiVI*^13^*.* We ran MultiVI by following the pipeline described in its tutorial (scvi version 0.19.0) (<https://docs.scvi-tools.org/en/stable/tutorials/notebooks/multimodal/MultiVI_tutorial.html>). Raw count matrices were used as input and the training epoch was set to 100 while the other parameters were left at their default values.

*BABEL*^15^*.* We ran BABEL by following the tutorial for the DANCE toolkit (version 1.0.1)^16^ (<https://github.com/OmicsML/dance/tree/main/examples/multi_modality/predict_modality/babel.py>). Raw count matrices were used as input and all parameters were set to default.

**Benchmarking results**

The quantitative benchmarking results on the four datasets are presented in **Supp. Fig. S34**. For the postnatal mouse brain dataset ATAC prediction task, MIDAS achieved the highest AUROC scores, followed by SpaMosaic (**Supp. Fig. S34a**). For spot-spot evaluation, SpaMosaic achieved the highest PCC and the lowest CMD (lower CMD scores indicate better performance) while for peak-peak evaluation, MIDAS achieved the highest PCC and the lowest CMD, followed by SpaMosaic and Babel. The embryonic mouse dataset prediction also showed similar results. These findings highlighted distinct strengths of different methods: SpaMosaic’s imputations better preserved spot-spot relations while MIDAS’s imputations better preserved the peak-peak relations. The reason is that SpaMosaic’s imputation is based on spatial-aware embeddings that better preserves spot-spot spatial relations, while MIDAS’s imputations are based on neural networks, which are not constrained by spatial information and can better adapt to feature patterns. Next, we recomputed the benchmarking metrics with the smoothed ground truth peak count matrices. With the smoothed ground truth, all methods’ performance improved substantially (**Supp. Fig. S34b**). For example, on the postnatal mouse brain dataset, SpaMosaic’s spot-spot PCC improved by 34.4% and AUROC improved by 52.3%, while MIDAS’s spot-spot PCC improved by 34.6% and AUROC improved by 32.4%. In this scenario, both SpaMosaic and MIDAS scored the highest AUROC, and SpaMosaic was still the top performer at preserving spot-spot relations while MIDAS remained the best in preserving peak-peak relations. For the protein modality prediction task with the lymph node and tonsil, the methods’ performance was broadly comparable (**Supp. Fig. S34c**). Only a few statistical differences were found, like BABEL’s poorer performance at protein-protein PCC.

### Ablation study on graph construction strategies

Here we present a study of SpaMosaic variants to evaluate the contribution of SpaMosaic’s model components towards its performance. The first variant is SpaMosaic (non-spatial), where the GNN component is replaced by a conventional multi-layer perceptron (MLP) while all other parts remain the same as the original SpaMosaic. The second variant is SpaMosaic (spatial-only), which retains the GNN but excludes the expression adjacency relationship when constructing the cell-cell graph, allowing us to investigate how the model performs when the GNN considers only spatial information. We assessed SpaMosaic and the two variants on both simulated data and three real datasets, the human lymph node, embryonic mouse brain (Misar) and embryonic mouse brain (Misar+Stereo) datasets.

For the simulation dataset, SpaMosaic and SpaMosaic (spatial-only) achieved similar ARI scores, both outperforming SpaMosaic (non-spatial) by a clear margin (**Supp. Fig. S35a**). This highlighted the importance of incorporating spatial information for improving clustering performance on this dataset. In the spatial domain continuity evaluation, SpaMosaic (spatial-only) obtained PAS scores comparable to SpaMosaic on sections 1 and 2, while scoring lower in PAS on section 3 (**Supp. Fig. S35b**). Both methods exhibited nearly identical CHAOS scores and substantially outperformed SpaMosaic (non-spatial) on both PAS and CHAOS metrics. In terms of batch integration, SpaMosaic attained the highest iLISI scores, followed by SpaMosaic (spatial-only) and SpaMosaic (non-spatial) (**Supp. Fig. S35c**). For modality alignment, SpaMosaic achieved the highest label transfer F1-scores, with SpaMosaic (spatial-only) ranking second (**Supp. Fig. S35d**). Consistent with these quantitative metrics, spatial plots of the clusters further demonstrated that the patterns identified by SpaMosaic and SpaMosaic (spatial-only) were more closely aligned with the factor labels (**Supp. Fig. S35e**). The UMAP plots also revealed that SpaMosaic produced clearer factor separation and more effective batch mixing than SpaMosaic (spatial-only), supporting its higher iLISI and F1-scores (**Supp. Fig. S35f**).

For the human lymph node dataset, all three models achieved similar ARI scores, with SpaMosaic (spatial-only) performing marginally better (**Supp. Fig. S36a**). Consistent with the previous dataset, SpaMosaic (spatial-only) attained the lowest PAS and CHAOS scores, followed by SpaMosaic (**Supp. Fig. S36b**). The improved scores of SpaMosaic (spatial-only) over the other two methods stemmed from its more coherent clustering within the medullary regions, whereas SpaMosaic and SpaMosaic (non-spatial) captured more dispersed clusters due to the noisy expression data (**Supp. Fig. S36c**). In terms of batch integration, SpaMosaic achieved the highest iLISI score, closely followed by SpaMosaic (non-spatial) (**Supp. Fig. S36d**). This suggested that incorporating cross-section proximity improved integration. Considering the results of all three SpaMosaic variants, it appeared that incorporating expression adjacency relationships helped to mitigate these batch effects. UMAP plots further confirmed that SpaMosaic and SpaMosaic (non-spatial) achieved more uniform mixing across sections, whereas SpaMosaic (spatial-only) exhibited some section-specific separation (**Supp. Fig. S36e**).

With the embryonic mouse brain (Misar) dataset, SpaMosaic (spatial-only) was again the best overall in terms of PAS and CHAOS, except for section 2’s CHAOS score (**Supp. Fig. S37a**). SpaMosaic was second in all except for section 2’s CHAOS score where it edged out SpaMosaic (spatial-only). Both SpaMosaic (spatial-only) and SpaMosaic outperformed SpaMosaic (non-spatial) by a clear margin in terms of PAS, again demonstrating that incorporating spatial information improves spatial clustering continuity. For batch integration, SpaMosaic (non-spatial) attained the highest iLISI score and demonstrated the most effective batch mixing in UMAP visualization, followed by SpaMosaic (**Supp. Fig. S37b, c**). However, despite this higher iLISI score, spatial plots of the clustering revealed that SpaMosaic (non-spatial) failed to consistently align key regions across the three sections. For example, cluster 11 (hindbrain) in sections 2 and 3 was absent in section 1, and cluster 10 (mantle zone of dorsal pallium, DPallm) in sections 2 and 3 did not appear in section 1 (**Supp. Fig. S37d**). In contrast, SpaMosaic produced more coherent and more consistent cluster patterns across all three sections.

For the embryonic mouse brain (Misar+Stereo) dataset, SpaMosaic achieved the lowest PAS scores on 4 out of 6 sections, with SpaMosaic (spatial-only) ranking second; both substantially outperformed SpaMosaic (non-spatial) (**Supp. Fig. S38a**). Both SpaMosaic and SpaMosaic (spatial-only) also achieved comparable CHAOS values, again outperforming SpaMosaic (non-spatial). In terms of batch integration, SpaMosaic (non-spatial) attained the highest iLISI score (**Supp. Fig. S38b**). We speculate that this is because spatial proximity encoding preserves spatial relationships by grouping nearby spots together, which reduces batch mixing. From the spatial plots, we observed that SpaMosaic (non-spatial) identified clustering patterns that appeared noisier compared to those from SpaMosaic and SpaMosaic (spatial-only) (**Supp. Fig. S38c**). For example, cluster 10 (hindbrain) was more dispersed across sections and lacked a clear spatial pattern. This was also reflected in its comparatively poorer PAS and CHAOS scores. The UMAP visualizations showed that SpaMosaic (non-spatial) indeed produced more uniform embeddings in terms of section and hence a higher iLISI score (**Supp. Fig. S38d**).

Overall, our testing with simulated and real datasets confirmed that incorporating spatial information can substantially enhance the spatial continuity of identified spatial domains and improve domain consistency across sections, making them more closely aligned with the underlying anatomical structure. However, we note that this can be accompanied by a trade-off in poorer batch integration on the local scale as measured by the iLISI metric. Meanwhile, considering cross-section expression adjacency in the cell-cell graph improves representation consistency across sections and thus batch integration, although it may slightly compromise the spatial clustering coherence within individual sections. This may be attributed to the fact that spatial-only information enforces local continuity, whereas integrating cross-section expression adjacency introduces a global structure at the cost of local smoothness.

### Benchmarking different graph network architectures

**Details of benchmarking model architecture**

In SpaMosaic’s model, a spot-spot graph is created for each modality to learn information on the relationships between spots in terms of expression similarity and spatial proximity. Prior to finalizing the model, we considered four different graph neural architectures. In the first architecture, we considered homogeneous graphs and employed the graph attention network (GAT)^17,18^ to automatically learn the different relationships between spots. We adopted the same approach of Dong et al.^18^, which is described as follows. The encoder layer in layer $k$is defined as:

$$z_{i}^{k}=\sigma(\sum_{j\in S_{i}} att_{ij}^{k}(W_{k}z_{j}^{k-1}))$$

where $W_{k}$ is the trainable weight matrix, $\sigma$ is the non-linear activation, $S_{i}$ denotes the neighbor set of spot $i$ (including itself), $att_{ij}^{k}$ is the edge weight between spot $i$ and $j$ which is the output of $k$-th graph attention layer and $z^{0}$ denotes the model inputs. We highlight here that the final layer does not employ any attention mechanism. The $k$-th attention layer is defined as^18^:

$$att_{ij}^{k}=\frac{exp(e_{ij}^{k})}{\sum_{j\in S_{i}} exp(e_{ij}^{k})}$$

where $e_{ij}^{k}=\mathrm{Sigmoid}\left( {v_{s}^{k}}^{T}\left( W_{k}z_{i}^{k-1} \right)+{v_{r}^{k}}^{T}(W_{k}z_{j}^{k-1}) \right)$ with $v_{s}^{k}$ and $v_{r}^{k}$ being trainable weight vectors^18^. We also followed the same approach of Dong et al. in setting up the decoder which shares the same weight matrices $W_{k}$ and attentions scores $att_{ij}^{k}$ as the encoder.

Alternatively, the spot-spot graphs can be heterogeneous. In this second approach, we considered two types of edges in the graph and employed the general heterogeneous graph transformer (HGT)^19^ to learn the heterogeneous edges. In brief, HGT uses node- and edge-type dependent parameters to characterize the heterogeneous attention over each edge, achieving dedicated representations for different types of nodes and edges^19^. We used the implementation of HGT in the Pytorch Geometric library^20^ and followed the example to build the whole network. Moreover, based on the framework of light graph convolution network (LGCN)^21,22^, we proposed a third model architecture. In traditional LGCNs, information is propagated and aggregated through:

$$\tilde{X}=LGCN\left( A, X \right)=Concat(X, \hat{A}X,\hat{A}^{2}X,\ldots,\hat{A}^{l}X)$$

where $\hat{A}=\tilde{D}^{-\frac{1}{2}}(A+I)\tilde{D}^{-\frac{1}{2}}$, $\tilde{D}$ is the diagonal degree matrix of $A+I$, and $l$ denotes the number of layers. The embeddings are finally transformed by a group of nonlinear transformations a multiple layer perceptron (MLP):

$$Z=f_{mlp}(\tilde{X})$$

Our modified HG-LGCN formulation then take the following form:

$$\tilde{X}=LGCN^{1}\left( A, X \right)=Concat(X, f\left( X \right), f\left( f\left( X \right) \right)\ldots,f^{l}\left( X \right))$$

where $f\left( X \right)={Concat(\hat{A}}^{intra}X,\hat{A}^{inter}X)$. The key idea of this modification is to construct two propagation paths between spots, and the output of current layer is recursively applied to subsequent layers. Thus, information propagation in this model architecture is structured like a complete binary tree. The fourth and last architecture is the weighted LGCN (WLGCN) that is described in the method section.

**Benchmarking results**

We first searched for the best combination of hyper-parameters for each architecture on a simulated dataset (simulation parameters: $b^{rna}=4$, $s^{rna}=1/15$, $p^{rna}=0$, $b^{pro}=2.0$, $s^{pro}=1/20$). The hyper-parameters optimized were the number of layers, the dimension of hidden layers, the output dimensions and the number of nearest neighbors used in constructing the spot-spot adjacency graph. We selected the parameter combination that achieved the highest ARI scores for the whole dataset.

Using the first replicate of simulation setting 1 as an example, we first examined the UMAP plots of the outputs from the different architectures (**Supp. Fig. S39a**). The output of all methods showed good separation of clusters except HGT. In terms of batch mixing, WLGCN and HG-LGCN achieved the best mixing visually, while local concentrations of batch specific spots were clearly visible for HGT and GAT. We hypothesized that attention-based architectures preferentially amplify signals within highly homogeneous, intra-section neighborhoods, thereby restricting information propagation across sections and leading to poorer batch mixing. We also examined the spatial plots of the outputs (**Supp.** **Fig. S39b**). HGT achieved the worst performance, being unable to capture factor 1 (orange) in sections 1 and 2, and factor 3 in section 3, while the other three methods were able to correctly capture the factors.

We further quantitatively evaluated the four architectures on the data generated with all five simulation settings in three aspects: spatial domain identification and continuity (ARI, PAS, and CHAOS), batch mixing (iLISI) and modality alignment (F1-score of label transfer between section 2 and 3). In terms of ARI, WLGCN, HG-LGCN, and GAT showed similar performance and were better than HGT (**Supp. Fig. S39c**). For the PAS and CHAOS metrics, the GAT, WLGCN, and HG-LGCN architectures were overall superior to HGT. Among these top performers, GAT achieved the best scores across most datasets, though its lead over WLGCN and HG-LGCN was relatively small (**Supp. Fig. S39d**). Assessing integration with the iLISI metric, HG-LGCN attained the highest iLISI scores with WLGCN as a close second (**Supp. Fig. S39e**). GAT and HGT lagged behind, and these were consistent with their UMAP plots. Finally, the label transfer F1-scores showed similarly good results for WLGCN, HG-LGCN, and GAT, while HGT again performed worse (**Supp. Fig. S35f**). In summary, both WLGCN and HG-LGCN achieved similarly superior performance. However, unlike HG-LGCN whose feature dimensionality grows exponentially with the number of layers ($2^{L}$), WLGCN increases only linearly ($L+1$), resulting in substantially lower time and memory costs for deeper networks. This efficiency makes WLGCN more practical for large graphs without sacrificing accuracy. Therefore, we selected WLGCN for our final model.

### Sensitivity analysis of *k* in spatial adjacency graph construction

The parameter *k* determines the neighborhood size in the spatial *k*-nearest neighbor (*k*NN) graph, directly controlling the extent of local information smoothing and propagation across spot embeddings. Intuitively, one might expect that the optimal *k* depends on the characteristics of specific technology, such as 10x Genomics Visium or Stereo-seq, where each spot is arranged in a hexagonal lattice with six direct neighbors. To investigate how the choice of *k* influences the model outcomes, we conducted evaluation on the simulated data, human lymph node, embryonic mouse brain (Misar), and embryonic mouse brain (Misar+Stereo) seq datasets with *k* set to 4, 6, 8, 10, and 20. We note that SpaMosaic performs random sampling of neighboring spots when the number of available neighbors is greater than the target *k*. While a fixed random seed could be set to ensure reproducibility, we deliberately varied the random seed to investigate whether different sampling results would substantially impact the outcomes. Therefore, for each *k* value, we repeated the experiment three times, each time using a randomly selected new seed.

We observed similar trends for the quantitative metrics across four datasets. The ARI scores remained relatively stable when *k* ranged from 4 to 10, but showed a noticeable decline at *k*=20 (**Supp. Fig. S40a, b**). The spatial clustering plots suggested an intuitive explanation: in the simulated dataset, increasing *k* causes each spatial factor to expand its cluster, particularly in sections 2 and 3, leading to a deviation from the ground-truth labels (**Supp. Fig. S41a**). In terms of spatial clustering continuity, larger *k* values led to lower PAS scores (**Supp. Fig. S40a-d**). This is expected, as a larger *k* introduces more neighboring spots into the smoothing process, making it more likely that most neighbors share the same cluster label. However, the CHAOS scores did not show substantial fluctuations (**Supp. Fig. S40a-d**). This is because PAS and CHAOS evaluate different aspects of the clustering results. Specifically, PAS assesses whether the majority of spots within a local spatial neighborhood belong to different clusters, thereby reflecting a broad spatial context around each spot. In contrast, CHAOS measures the nearest distance from each spot to another spot within the same cluster. As a result, CHAOS primarily reflects the presence of isolated points in the clustering outcome. From this perspective, varying *k* has a similar effect on reducing clustering outliers across four datasets. In terms of batch integration, a larger *k* generally led to lower iLISI scores across the four test datasets (**Supp. Fig. S40a-d**). The UMAP visualization further supported the result that increasing *k* led to reduced section mixing, and this trend was also observed in other datasets (**Supp. Fig. S41b and Supp. Fig. S42-S44**). These phenomena were likely due to the incorporation of too many neighbors in the *k*NN graph, which resulted in over-smoothing of nearby spot representations and increased inconsistencies across batches. Importantly, across the four datasets, triplicate runs with different random seeds for each *k* value produced highly consistent results, with only minor variation across all quantitative metrics (**Supp. Fig. S40**). This consistency demonstrated that the random sampling of neighbors inherent in SpaMosaic’s *k*NN graph construction did not introduce substantial variability to the experimental outcomes. Moreover, *k*=10 achieved a well-balanced performance in terms of clustering accuracy, spatial continuity, and batch integration across four datasets. Hence, we adopted *k*=10 for all examples presented in our manuscript.

### Impact of clustering method choice on benchmarking

The use of different clustering algorithms downstream of data integration leads to distinct sets of clusters which in turn can affect the benchmarking results. To validate the robustness of our benchmarking framework and its results, we examined each method’s performance with different downstream clustering algorithms (Kmeans, Mclust, and Leiden). We conducted these analyses on four datasets: simulated data, the human lymph node dataset, the embryonic mouse brain (Misar), and embryonic mouse brain (Misar+Stereo) datasets. To evaluate performance, we used the PAS and CHAOS metrics, and additionally reported ARI scores when ground truth labels were available.

For the simulated dataset, **Supp. Fig. S45a-c** panels show an example of spatial clustering results for SpaMosaic and competing methods applied with Kmeans, Mclust, and Leiden algorithms, respectively. SpaMosaic consistently identified the five spatial domains with high accuracy and superior spatial continuity across all clustering methods. This superior performance was reflected in its highest ARI and lowest PAS scores across all five simulated datasets, regardless of the clustering method used (**Supp. Fig. S45d-f**). Among the competing methods, Mclust did not work well with CLUE as only one spatial factor was captured in sections 1 and 3. On the other hand, CLUE paired with either Leiden and Kmeans was able to capture the factors and was second to SpaMosaic in terms of ARI. For the PAS metric, either CLUE or MIDAS were second to SpaMosaic. For CHAOS, SpaMosaic was consistently first across three clustering methods on datasets 4 and 5, and ranked in the top two with Kmeans and Mclust on datasets 1, 2 and 3.

For the human lymph node dataset, **Supp. Fig. S46a-c** panels illustrate the spatial clustering results for SpaMosaic and other methods using Kmeans, Mclust, and Leiden, respectively. CLUE, Cobolt, and scMoMaT all showed notable performance degradation with Mclust, as observable from the spatial plots and ARI scores, while the other methods were relatively robust to the choice of clustering method. When Kmeans was used, StabMap achieved the highest ARI and relatively low PAS and CHAOS values (**Supp. Fig. S46d**). With Mclust, SpaMosaic achieved the highest ARI score. When Leiden was used, MIDAS yielded the best performance in terms of both ARI and spatial continuity metrics (**Supp. Fig. S46d**).

For the embryonic mouse brain (Misar) dataset, **Supp. Fig. S47a-c** panels show that SpaMosaic demonstrated robust performance across all clustering strategies in capturing fine brain structures such as mantle zone of the dorsal pallium (DPallm) and ventricular zone of dorsal pallium (DPallv), and consistently producing coherent clustering results. The performance consistency is also observed in the metrics where SpaMosaic achieved the lowest PAS and CHAOS scores except for the CHAOS metric with Mclust where CLUE and MIDAS edge it out. Among the three clustering options, SpaMosaic with Mclust scored lowest for PAS while SpaMosaic with Leiden scored lowest for CHAOS (**Supp. Fig. S47d**). For the other methods, scMoMaT performed best with Leiden, while CLUE, Cobolt, MIDAS, StabMap, and MultiVI performed better when Mclust was used. Kmeans generally showed the worst overall performance for most methods with this dataset.

For the embryonic mouse brain (Misar+Stereo) dataset, as shown in **Supp. Fig. S48a, b and Supp. Fig. S49a**. SpaMosaic generated consistently coherent spatial domains across all clustering methods. SpaMosaic also scored well in the metrics with consistently the lowest or second lowest PAS and CHAOS scores across the three clustering algorithms (**Supp. Fig. S49b**). For the other methods, the choice of clustering algorithm had mixed impact. Taking MIDAS with Mclust as an example, it grouped the three Misar-seq slices into three large clusters, leading to suboptimal results. There was also no clear optimal clustering algorithm as measured with the metrics. Leiden was optimal for CLUE, scMoMaT, and MultiVI, while Cobolt was less sensitive to the choice of clustering algorithm.

In summary, this analysis demonstrated that while the choice of clustering algorithm can affect the performance of most methods, SpaMosaic remained robust and accurate regardless of the clustering method tested. As Mclust performed slightly better for SpaMosaic across datasets compared to the other clustering methods, we retained its use with SpaMosaic. For the competing methods, the impact of clustering algorithm choice was more varied. There was no consistently superior method and exhaustive testing of optimal clustering algorithms for each method and each dataset is out of the scope of this work.

### Effect of Harmony preprocessing on benchmarking results

Here we study the impact of preprocessing steps on the mosaic integration methods, specifically batch integration of the input data prior to learning a joint embedding. For SpaMosaic, Harmony is one of the preprocessing steps employed. A popular single-cell data integration tool, Harmony takes in low dimensional embeddings like PCA as input and outputs a batch corrected embedding of the same dimensions. Among the competing methods, only CLUE accepts such low dimension embeddings as input, while the other methods require raw counts or normalized expression matrices as input. Therefore, we could only additionally consider the combination of CLUE with Harmony (CLUE+prep). We thus conducted a separate comparison of SpaMosaic, default CLUE and CLUE with Harmony (CLUE+prep), to test the impact of Harmony as a preprocessing step. We applied the methods on three datasets, simulated data, the human lymph node, and embryonic mouse brain (Misar) datasets. We then quantitatively assessed the outputs with metrics that covered three aspects: spatial domain identification and continuity (using ARI, PAS, and CHAOS), batch correction (iLISI), and modality alignment (label transfer F1-score).

For the simulated data, SpaMosaic outperformed all other methods by a clear margin across all metrics and in all five simulation settings (**Supp. Fig. S50a-d**). Compared to CLUE, CLUE+prep achieved higher ARI scores on 4 out of 5 datasets, but its PAS and CHAOS scores were poorer on 3 out of 5, indicating reduced spatial continuity. Despite using batch-corrected low-dimensional inputs, CLUE+prep did not show noticeably improved iLISI scores over CLUE, indicating that simply applying batch correction to each modality is not sufficient for better data integration. In addition, CLUE+prep achieved lower label transfer F1-scores than CLUE across all datasets. Spatial clustering and UMAP plots also highlighted SpaMosaic’s superior performance in both clustering quality and data integration, with CLUE+prep and CLUE showing little difference (**Supp. Fig. S50e, f**).

For the human lymph node dataset, CLUE+prep achieved the highest ARI score, while SpaMosaic and CLUE obtained comparable scores (**Supp. Fig. S51a**). However, only SpaMosaic and CLUE+prep successfully delineated the follicle structures that are small regions embedded within the cortex (**Supp. Fig. S51b**). In terms of spatial continuity, SpaMosaic attained the lowest CHAOS scores on sections 1 and 2, with CLUE and CLUE+prep obtaining similar scores (**Supp. Fig. S51c**). However, the PAS metric revealed a different trend: CLUE achieved the lowest PAS scores across all three sections, whereas SpaMosaic obtained the highest PAS scores on sections 1 and 3. This discrepancy may be attributed to the spatial clustering within the medullary regions, where SpaMosaic’s clusters were more dispersed (**Supp. Fig. S51b**). In terms of batch integration, SpaMosaic achieved the highest iLISI score with CLUE+prep being second (**Supp. Fig. S51d**). UMAP plots also showed that SpaMosaic achieved good batch mixing of the three sections, while CLUE+prep achieved better mixing than CLUE (**Supp. Fig. S51e**).

For the embryonic mouse brain (Misar) dataset, SpaMosaic attained the lowest PAS scores across all sections and the lowest CHAOS scores on sections 2 and 3, suggesting improved spatial continuity over the other two methods (**Supp. Fig. S52a**). Meanwhile, CLUE+prep yielded higher PAS and CHAOS scores compared to CLUE. In terms of batch integration, CLUE+prep attained the highest iLISI score, with SpaMosaic achieving a close second (**Supp. Fig. S52b**). Spatial plots of the clustering results further demonstrated that spatial patterns in SpaMosaic’s clusters were more coherent and more consistent across three sections (**Supp. Fig. S52c**). For example, SpaMosaic consistently assigned the hindbrain and diencephalon regions to cluster 6 across all three sections, with this cluster exhibiting clear spatial domain boundaries. In contrast, CLUE assigned the hindbrain and diencephalon regions to different clusters (1, 2, and 7, respectively) that appeared dispersed, while CLUE+prep showed similar inconsistency (clusters 1, 1, and 2, respectively) with equally dispersed spatial patterns. Examination of the UMAP plots confirmed that the sections were better integrated in the results of SpaMosaic and CLUE+prep than in those of CLUE (**Supp. Fig. S52d**).

In summary, applying batch correction to each modality before integration presents a trade-off: it can improve performance in terms of clustering accuracy (ARI) and batch integration (iLISI), but often at the expense of spatial continuity as measured with metrics (PAS and CHAOS) and without clear benefits for modality alignment. In our comparative analysis of CLUE, CLUE+prep and SpaMosaic, SpaMosaic consistently outperformed the others on simulated data. On real tissue derived datasets, SpaMosaic achieved superior spatial domain coherence and cross-section clustering consistency, while maintaining batch mixing performance comparable to CLUE. Although CLUE+prep showed the highest iLISI scores in embryonic mouse brain (Misar) dataset, this came with noticeably poorer spatial coherence and cross-section clustering consistency.

### Evaluation of time and memory consumption

Here we present a study that evaluates the computational performance of SpaMosaic and competing methods. All evaluation tests were performed on a server equipped with a 36-core Intel Core i9-10980XE CPU, 256 GB of RAM and an NVIDIA GeForce RTX 3090 GPU with 24GB of GPU memory. We recorded the CPU time usage, GPU time usage, wall time, peak Random Access Memory (RAM) usage and peak GPU memory consumption of the methods while varying the number of CPU cores/threads. To assess performance in diverse mosaic integration scenarios, we used three datasets: embryonic mouse brain (Misar), embryonic mouse brain (Misar+Stereo) and mouse embryo datasets. The embryonic mouse brain dataset consists of one multimodal section and two single-modality sections with different modalities. The embryonic mouse brain dataset contains six slices, all with RNA profiles, but only three of them also include an ATAC modality. The mouse embryo presents a more complex setting where no section contains all modalities and no modality is shared across all sections. For each scenario, we ran each method under different settings of CPU cores and maximum number of threads allowed, namely, 1, 2, 4, 8, and 16. We note that the server was shared with other users and therefore the performance metrics may have been subject to some variability.

We first considered CPU usage by the different methods. **Supp. Fig. S53a** illustrates the wall time usage of each method as the number of CPU cores/threads increases. SpaMosaic consistently achieves the lowest wall time, followed by StabMap and scMoMaT. In contrast, MultiVI and CLUE exhibit the highest wall times across all datasets. As the number of CPU cores/threads increased, the wall time for SpaMosaic, scMoMaT, and StabMap generally decreased, indicating effective parallelization and scalability. In comparison, the wall time of the remaining methods either remain relatively unchanged or fluctuated with increasing cores, suggesting limited multi-threading support**. Supp. Fig. S53b** presents the CPU time comparison across all methods. SpaMosaic and StabMap achieved the lowest CPU times, while MultiVI and CLUE remained among the most computationally expensive in terms of CPU usage. For GPU time usage, the trends largely mirrored those observed for wall time (**Supp. Fig. S53c**). Notably, SpaMosaic showed the lowest GPU runtime, indicating that most of its wall time was spent on preprocessing prior to model training. On RAM usage, StabMap and SpaMosaic once again had the lowest memory consumption across all datasets (**Supp. Fig. S53d**). On the other hand, scMoMaT exhibited the highest RAM usage on the embryonic mouse brain (Misar) and mouse embryo datasets, while UINMF consumed the highest amount of RAM on the embryonic mouse brain (Misar+Stereo) dataset. As for GPU memory consumption, CLUE consistently ranked lowest across all three datasets (**Supp. Fig. S53e**). scMoMaT also had the highest GPU memory usage, while SpaMosaic maintained moderate usage close to CLUE’s.

To investigate how SpaMosaic scales with the number of features, modalities and samples, we conducted additional experiments with the embryonic mouse brain (Misar) dataset under a maximum CPU core/thread limit of 16. Specifically, to evaluate the impact of varying feature numbers on SpaMosaic’s computational performance, we randomly sampled a range of features from the RNA and ATAC modalities. For RNA, the number of sampled features employed was 1,000; 2,000; 4,000; 8,000; 16,000; 32,000. For ATAC, it was 1,000; 3,000; 9,000; 27,000; 81,000; 191,034. Each RNA-ATAC feature pair was used to test the performance of SpaMosaic, as shown in **Supp. Fig. S54.** Overall, as the number of features increased, both wall time and CPU time showed a logarithmic growth, though wall time remained under 60 seconds across all settings. GPU time remained under 4 seconds. This is expected because increasing the number of input features primarily affects the preprocessing stage which entails little GPU processing, while the GPU-intensive neural network training is performed on the fixed-size low-dimensional representations and is thus largely unaffected by the original feature count. As for RAM usage, it increased linearly with the number of features, whereas GPU memory usage showed no clear change.

We next evaluated the impact of the number of modalities on SpaMosaic’s computational performance by using the embryonic mouse brain (Misar) to create simulated datasets with different modality counts. The embryonic mouse brain (Misar) originally contains two modalities, RNA and ATAC, and three batches where one batch contains both RNA and ATAC, while the other two contain only RNA and only ATAC, respectively. To simulate datasets with more modalities, we replicated each original modality per batch by a given multiplication factor and treated each copy as a distinct modality. For example, with a factor of 2: the first batch originally has RNA and ATAC; it becomes {RNA-1, RNA-2, ATAC-1, ATAC-2}; the second batch (originally RNA-only) becomes {RNA-1, RNA-2}; the third batch (originally ATAC-only) becomes {ATAC-1, ATAC-2}. This results in 4 distinct modalities. We tested multiplication factors from 1 to 5, corresponding to 2, 4, 6, 8, and 10 modalities.

The results obtained are shown in **Supp. Fig. S55a**, where we see that the wall time and CPU time increased approximately linearly with the number of modalities. However, even at the highest setting with 10 modalities, wall time remained under 4 minutes. While the curve for GPU time was steeper than that of the wall time’s, the maximum time required was around 30 seconds, contributing minimally to the overall runtime. In contrast, RAM and especially GPU memory usage increased more sharply, from approximately 2 GB to 12 GB. This steep growth was primarily due to SpaMosaic’s design, which required pairwise alignment between modalities and thus a quadratic increase in the number of modality pairs (**Supp. Fig. S55b**). Currently, datasets with four or more modalities are rare and therefore it is not a major limitation presently. For datasets with three or fewer modalities, GPU memory usage remained below 4 GB.

We also evaluated the impact of increasing the number of sections using simulated data created from the embryonic mouse brain (Misar) dataset. Specifically, we replicated the original three sections with different multiplication factors to increase the number of sections. For example, with a factor of 2, each original slice and its associated modalities were duplicated once, resulting in a dataset with 2 multimodal sections, 2 RNA-only sections, and 2 ATAC-only sections. We employed several replication settings that corresponded to total section counts of 3, 6, 9, 15, 24, 45, 69, and 100. For all cases except the 100-section setting, the three original sections were replicated equally. The 100-section dataset was created by replicating the first two sections 33 times and the third section 34 times.

As shown in **Supp. Fig. S56**, SpaMosaic, CLUE and Cobolt (GPU-based integration methods) scaled up to 100 sections with reasonable runtimes (<10,000s). While MultiVI was able to scale up to 100 sections, it required much longer runtime (>50,000s) than others. For scMoMaT, it was unable to handle 100 sections due to high GPU memory usage. Regarding specific resource usage, SpaMosaic achieved the lowest wall time, CPU time and GPU time consumption. For CPU memory usage, all methods showed a linear increase with the number of sections, with MultiVI exhibiting the slowest growth and the others displaying similar rates. For GPU memory usage, CLUE, Cobolt and MultiVI showed the slowest increase, likely due to their mini-batch training strategies. scMoMaT exhibited the highest memory growth, exceeding 23 GB at 69 sections, while SpaMosaic followed, though with a substantially lower growth rate.

Finally, we evaluated the maximum number of spots per section that SpaMosaic can handle. Using the embryonic mouse brain (Misar) dataset as template, we simulated data of multiple sets of three sections: one multimodal section (RNA+ATAC) and two single-modality sections (RNA or ATAC), with different numbers of spots. All three sections per set had the same number of spots. The spatial distribution of spots in each section is shown in **Supp. Fig. S57**. The RNA modality contained 10,000 genes with expression values drawn from a Poisson distribution with parameter $\lambda=1$, while the ATAC modality consisted of 100,000 accessibility peaks encoded as binary values (0 or 1) with 99.9% sparsity. We generated multiple dataset configurations in which each section contained 50,000, 100,000, 200,000, 400,000, or 800,000 spots. In the computation resource consumption testing, SpaMosaic was able to scale up to 800,000 spots per section (approximately an 895×895 spot grid) on the test server (**Supp. Fig. S58)**. Our future work will focus on optimizing the model implementation to enable mini-batch training, with the goal of further reducing GPU memory usage and thus further improving scalability.

In summary, we benchmarked SpaMosaic against existing mosaic integration methods across three distinct integration scenarios and found that SpaMosaic consistently achieved lower wall time, CPU time and GPU time, along with relatively low GPU memory and RAM usage. Even on the largest dataset, embryonic mouse brain (Misar+Stereo) (with 50,000 spots across six sections), SpaMosaic completed the integration within one minute, while the slowest method, MultiVI, required over 100 minutes. Using a fixed dataset, increasing the number of input features primarily affected preprocessing time and RAM usage, but had minimal impact on model training and inference speed. In contrast, increasing the number of modalities led to a quadratic increase in GPU time and memory usage, as SpaMosaic performs pairwise alignment across all modality combinations. Nevertheless, even with 10 modalities, SpaMosaic’s GPU runtime remained below 30 seconds. For increases in the number of sections, it entailed a linear rise in GPU time and memory usage by SpaMosaic. However, on the test server, SpaMosaic could handle up to 100 sections or more. Finally, with a mosaic dataset (one RNA+ATAC section, one RNA section, and one ATAC section), SpaMosaic successfully processed data with up to 800,000 spots per section.

### Reference

1.	Xiang, M. *et al.* A Single-Cell Transcriptional Roadmap of the Mouse and Human Lymph Node Lymphatic Vasculature. *Front Cardiovasc Med* **7**, (2020).

2.	Elmore, S. A. Histopathology of the Lymph Nodes. *Toxicol Pathol* **34**, 425–454 (2006).

3.	Hernandez-Quiles, M., Broekema, M. F. & Kalkhoven, E. PPARgamma in Metabolism, Immunity, and Cancer: Unified and Diverse Mechanisms of Action. *Front Endocrinol (Lausanne)* **12**, (2021).

4.	Najt, C. P., Devarajan, M. & Mashek, D. G. Perilipins at a glance. *J Cell Sci* **135**, (2022).

5.	Fernandez, L. F. A. & Pineda-Cortel, M. R. B. ADIPOQ gene (T45G and G276T) single nucleotide polymorphisms and their association with gestational diabetes mellitus in a Filipino population. *BMC Endocr Disord* **23**, 248 (2023).

6.	Russell, A. J. C. *et al.* Slide-tags enables single-nucleus barcoding for multimodal spatial genomics. *Nature* **625**, 101–109 (2024).

7.	Liu, Y. *et al.* High-plex protein and whole transcriptome co-mapping at cellular resolution with spatial CITE-seq. *Nat Biotechnol* **41**, 1405–1409 (2023).

8.	Husson, H. *et al.* CXCL13 (BCA‐1) is produced by follicular lymphoma cells: role in the accumulation of malignant B cells. *Br J Haematol* **119**, 492–495 (2002).

9.	Abe, Y. *et al.* A single-cell atlas of non-haematopoietic cells in human lymph nodes and lymphoma reveals a landscape of stromal remodelling. *Nat Cell Biol* **24**, 565–578 (2022).

10.	Martinez-Valdez, H. *et al.* Human germinal center B cells express the apoptosis-inducing genes Fas, c-myc, P53, and Bax but not the survival gene bcl-2. *J Exp Med* **183**, 971–977 (1996).

11.	Hu, Y. *et al.* Benchmarking algorithms for single-cell multi-omics prediction and integration. *Nat Methods* **21**, 2182–2194 (2024).

12.	Hu, Y. *et al.* WEDGE: imputation of gene expression values from single-cell RNA-seq datasets using biased matrix decomposition. *Brief Bioinform* (2021) doi:10.1093/bib/bbab085.

13.	Ashuach, T. *et al.* MultiVI: deep generative model for the integration of multimodal data. *Nat Methods* **20**, 1222–1231 (2023).

14.	Gayoso, A. *et al.* Joint probabilistic modeling of single-cell multi-omic data with totalVI. *Nat Methods* **18**, 272–282 (2021).

15.	Wu, K. E., Yost, K. E., Chang, H. Y. & Zou, J. BABEL enables cross-modality translation between multiomic profiles at single-cell resolution. *Proceedings of the National Academy of Sciences* **118**, (2021).

16.	Ding, J. *et al.* DANCE: a deep learning library and benchmark platform for single-cell analysis. *Genome Biol* **25**, 72 (2024).

17.	Veličković, P. *et al.* Graph attention networks. *arXiv preprint arXiv:1710.10903* (2017).

18.	Dong, K. & Zhang, S. Deciphering spatial domains from spatially resolved transcriptomics with an adaptive graph attention auto-encoder. *Nat Commun* **13**, 1739 (2022).

19.	Hu, Z., Dong, Y., Wang, K. & Sun, Y. Heterogeneous graph transformer. in *Proceedings of the web conference 2020* 2704–2710 (2020).

20.	Fey, M. & Lenssen, J. E. Fast graph representation learning with PyTorch Geometric. *arXiv preprint arXiv:1903.02428* (2019).

21.	He, X. *et al.* Lightgcn: Simplifying and powering graph convolution network for recommendation. in *Proceedings of the 43rd International ACM SIGIR conference on research and development in Information Retrieval* 639–648 (2020).

22.	Xia, C.-R., Cao, Z.-J., Tu, X.-M. & Gao, G. Spatial-linked alignment tool (SLAT) for aligning heterogenous slices. *Nat Commun* **14**, 7236 (2023).
